# Design-Evolution Synergy Enables New-to-Nature Enzyme Chemistry

**DOI:** 10.64898/2026.10.01.754751

**Authors:** Zi-Qi Li, Ruijie K. Zhang, Pei-Pei Xie, Jacob Kelly, Yu Zhang, Zi-Yang Qin, Theophile Lambert, Orna Mukhopadhyay, Deirdre Hanley, Amy E. Danson, Josh Abramson, Sukhdeep Singh, Zachary Wu, Pushmeet Kohli, Peng Liu, Jue Wang, Frances H. Arnold

**Affiliations:** Division of Chemistry and Chemical Engineering, California Institute of Technology, Pasadena, California 91125, United States; Google DeepMind, London, United Kingdom; Department of Chemistry, University of Pittsburgh, 219 Parkman Avenue, Pittsburgh, Pennsylvania 15260, United States

## Abstract

Directed evolution and machine-learning-assisted evolution have transformed biocatalysis by enabling rapid optimisation of catalytic performance within compatible protein scaffolds. In parallel, advances in deep-learning-driven *de novo* protein design have expanded accessible protein architectural space, enabling the generation of functional catalysts for increasingly challenging transformations. We herein report a synergistic design-evolution strategy to encode new-to-nature regio- and enantioselective C–H amination reactions using engineered *de novo* haemproteins to synthesise chiral piperidines, a privileged pharmacophore in drug discovery. By integrating motif scaffolding, computational re-design, and directed evolution, we demonstrate *de novo* enzyme design can enable challenging, high-energy-barrier enzyme chemistry with programmable regio- and enantioselectivity, high catalytic efficiency, excellent enzyme stability, and scaffold diversity.

## Main

Nature’s catalytic machinery, enzymes, mediate an extraordinary range of chemical transformations with exceptional selectivity and efficiency. Yet naturally evolved enzymes remain confined to the reaction manifolds explored during biological evolution. Over the past decade, directed evolution (DE) *(1*) and machine-learning-assisted directed evolution (MLDE) (*2–3*) have transformed biocatalysis by enabling rapid optimisation of enzyme activity, selectivity and stability for increasingly diverse new-to-nature transformations (*4*). Within compatible protein architectures, these approaches efficiently amplify the “promiscuous” activities of proteins through iterative sequence diversification and selection. However, because DE and MLDE fundamentally operate within the sequence landscape of a fixed protein scaffold (Fig. 1a), catalytic outcomes requiring alternative active-site geometries may remain inaccessible regardless of sequence optimisation depth. Consequently, the central question in modern enzyme engineering is no longer whether evolution can optimise catalytic performance, but rather how to identify or create evolutionary starting points toward fundamentally new catalytic functions.

**Fig. 1:**
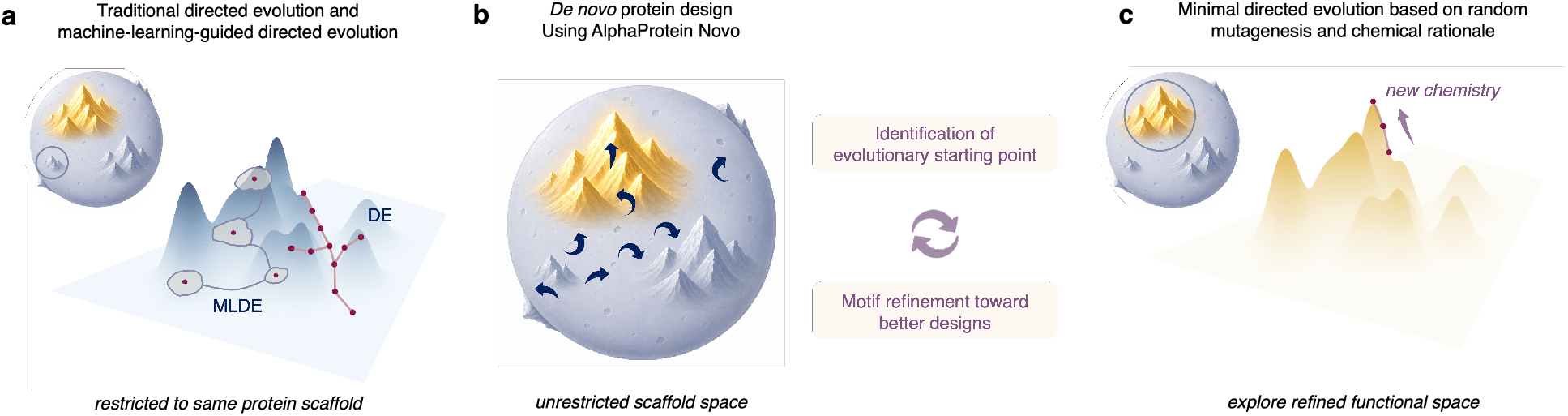
Background and Project Synopsis. **a**, Traditional directed evolution (DE) and machine-learning-guided directed evolution (MLDE) covers the *sequence-function* landscape within a specific protein scaffold. **b**, *De novo* protein design using AP Novo allows expansion of accessible architectural scaffold space. **c**, A design-evolution synergy allows interrogation of three-dimensional *sequence-structure-function* space by integrating discovery of evolutionary starting point, target-specific directed evolution, and evolution-guided expansion of scaffold diversity.

In the meantime, advances in deep-learning-driven *de novo* protein design (*5–6*) and structure prediction (*7–8*) have, in principle, opened vast regions of protein architectural space (Fig. 1b), enabling the generation of high-affinity small-molecule and protein binders (*9–15*). By generating proteins around specified catalytic motifs, ligands, cofactors, or transition-state geometries, *de novo enzyme* design is an area of growing interest, progressing from enzymes with natural functions (*16–20*), such as luciferases (*16*) and hydrolases (*17–20*), to enzymes catalysing new-to-nature transformations that are widely used in chemistry (*21–33*), including carbene transfer (*21– 23*), Kemp elimination (*24–25*), a retro-Aldol reaction (*26–27*), and Morita–Baylis–Hillman reactions (*27–28*). Collectively, these advances have established *de novo* enzyme design as a viable strategy for constructing functional catalysts with activities approaching, and in some cases surpassing (*18*), those of small-molecule catalysts or engineered natural enzymes. We now raise the bar and ask: can *de novo* enzyme design access transformations that require selective stabilisation of high-energy transition states and reactive intermediates that are beyond the scope of current small-molecule and biocatalytic paradigms?

Addressing this challenge requires understanding of the distinct strengths of each engineering strategy. *De novo* design expands accessible architectural scaffold space by encoding catalytic geometries, substrate-binding modes, and conformational landscapes that may not exist in naturally evolved proteins (*29–33*). However, for complex transformations, rationally designed stationary binding geometries alone rarely achieve the desired reactivity. The catalytic environment governing substrate activation, transition-state stabilisation, and product inhibition along the reaction trajectory is frequently incompletely understood or difficult to predict from first principles. Directed evolution, in contrast, explores catalytic solutions through iterative diversification and selection within experimentally interrogated sequence-fitness landscapes (*34*). Rather than functioning as distinct engineering strategies, *de novo* design and directed evolution are highly synergistic, and operate across complementary dimensions of the protein sequence– structure–function landscape. Specifically, while *de novo* design identifies evolutionary starting points (*35*) beyond natural scaffold and function space, directed evolution precisely evolves for the desired function and reciprocally leads to better designs through motif refinement. Both approaches converge on a common objective of not merely establishing catalytic feasibility, but efficiently accessing desired biocatalytic functions through strategically navigated design-evolution trajectories (Fig. 1b–c). As biocatalysis targets increasingly sophisticated chemistry, shortening the path from catalyst conception to functional implementation will define the future of enzyme engineering.

Recently, we described AlphaProtein Novo (AP Novo), a *de novo* enzyme design pipeline integrating diffusion-based motif-scaffolding with AlphaFold3-based filtering developed through iterative design–test–learn cycles (*36*). Building on this framework, we have developed a synergistic design-evolution strategy to encode new-to-nature regio- and enantioselective reactions, demonstrated here using *de novo* haemproteins for the synthesis of chiral piperidines, a privileged pharmacophore in drug discovery (*37*). Recent advances in *de novo* haem-binding helical bundles established catalytic feasibility within a specific designed scaffold using benchmark cyclopropanation and heteroatom–H bond insertion reactions (*21–23*); the present work expands both the accessible chemical space and the protein architecture of *de novo* haemprotein catalysis. By integrating motif scaffolding, computational re-design, directed evolution, and motif refinement, we demonstrate that *de novo* enzyme design can enable new-to-nature and new-to-chemistry biocatalysis with programmable regio- and enantioselectivity, high catalytic efficiency, and excellent enzyme stability and evolvability.

## Results and Discussion

### Reaction design and directed evolution from a natural protoglobin scaffold

Intramolecular C–H amination through metal-catalysed nitrene transfer has emerged as a powerful strategy for the synthesis of *N*-heterocycles (*38–43*). Substantial advances in both small-molecule catalysis (*38–42*) and biocatalysis (*44–51*) have been achieved in the construction of cyclic sultams (*44–47*), sulfamides (*48*), carbamates (*49*), lactams (*50*), and indolines (*51*) from electronically stabilised nitrenoid intermediates. By contrast, synthesis of chiral saturated *N*-heterocycles prevalent in pharmaceuticals and bioactive molecules (*37*) *via alkyl* nitrenoid C–H amination remains underdeveloped (*43, 52–57*), with asymmetric catalysis strategies limited to benzylic C– H amination leading to thermodynamically and kinetically favoured 2-arylpyrrolidines (*51, 58– 60*).

Extending alkyl nitrene chemistry beyond systems with unambiguous regioselectivity to those involving competing C–H amination pathways (Fig. 2) presents several fundamental challenges. Beyond the intrinsic instability of alkyl nitrenoid intermediates (*61–62*), regioselective amination of the benzylic versus homobenzylic C–H bond *via* competing 1,6-versus 1,5-hydrogen atom transfer, respectively, is complicated by the conformational flexibility of the alkyl chain and the absence of directing functionalities (*63–66*). Consequently, no precedents exist for catalyst-controlled regiodivergent C–H amination *via* an alkyl nitrenoid intermediate, nor for the enantioselective synthesis of either regioisomer. Surveying first-row transition-metal catalysts capable of intramolecular alkyl nitrene transfer (Scheme S3) further revealed that most systems adopt relatively planar coordination geometries with open axial access to the reactive intermediate. Unlike small-molecule catalysts, however, enzymes operate within DNA-encoded three-dimensional catalytic architectures capable of exerting precise control over substrate positioning and reactive-intermediate geometry. We therefore envisioned that a tailored enzyme scaffold could simultaneously govern regioselectivity and enantioselectivity in alkyl nitrenoid-mediated C–H amination.

**Fig. 2:**
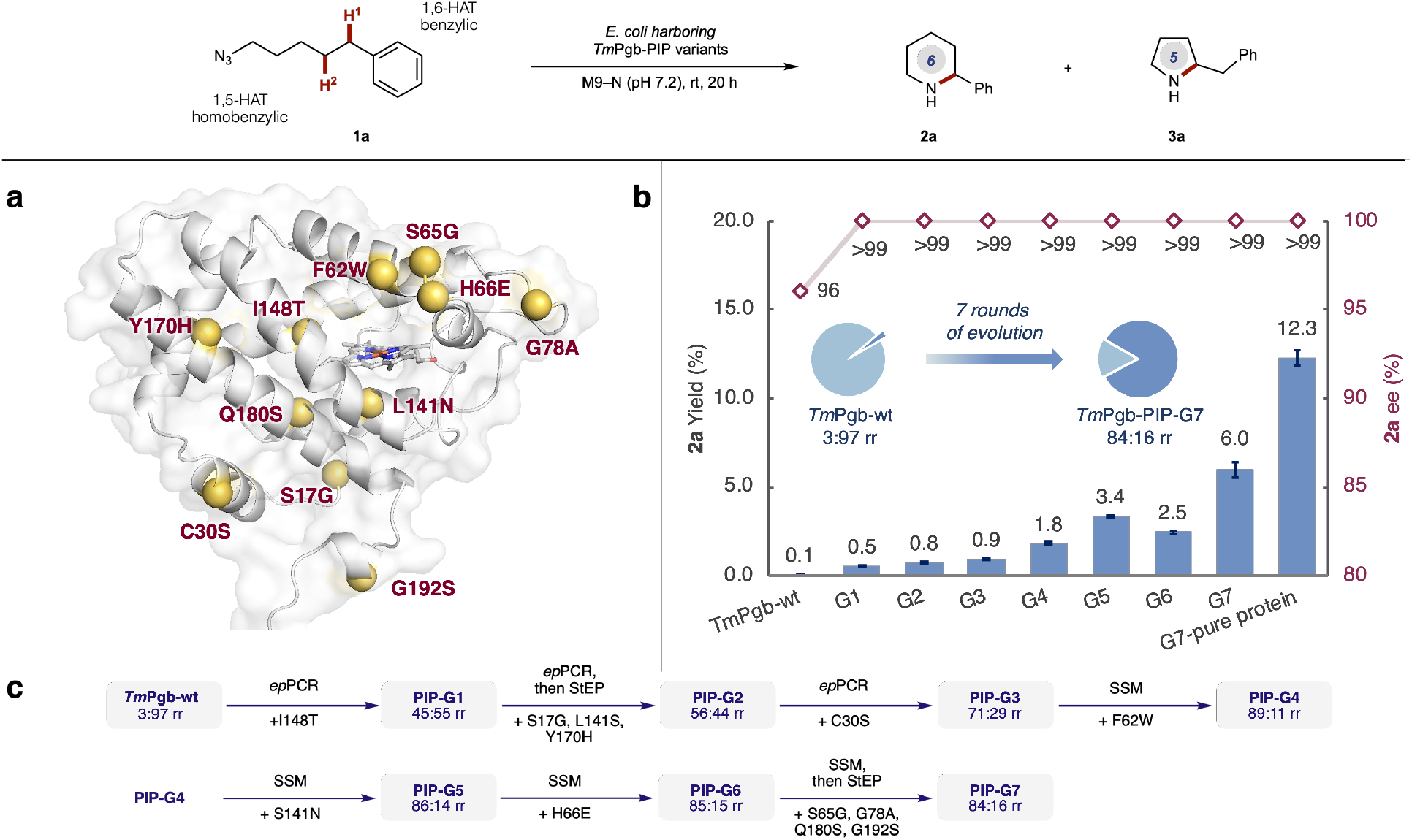
Directed evolution of *Tm*Pgb-PIP-G7-5448. **a**, Mutated residues of ***Tm*Pgb-PIP-G7-5448** compared to ***Tm*Pgb-wt**. Illustration is made on an AlphaFold 3-generated model. **b**, Evolutionary trajectory of ***Tm*Pgb-PIP-G7-5448**. Reaction performed at analytical scale using *E. coli* harbouring ***Tm*Pgb-PIP** variants (OD_600_ = 30) with **1a** (5.0 mM) as model substrate, in M9– N buffer (pH 7.2) with 5% v/v EtOH as co-solvent at room temperature under anaerobic conditions for 20 h. All yields represent assay yield of **2a** based on a calibration curve made on the SIM channel. **c**, Methods used for library generation, and mutations accumulated in each round of directed evolution.

Using (5-azidopentyl)benzene (**1a**, Fig. 2) as a model substrate, we first screened a panel of wild-type and engineered protoglobins to identify evolutionary starting points for regio- and enantioselective C–H amination (*67*). Several protoglobin homologues catalysed the desired intramolecular cyclisation, with the highest combined activity toward both regioisomeric products **2a** and **3a** (Fig. 2) observed in a collection of wild-type protoglobins found in various thermophilic archeae and bacteria. Starting from ***Tm*Pgb-wt** from ***T****hermus a****m****yloliquefaciens*, directed evolution afforded pyrrolidine synthase ***Tm*Pgb-PYR-G2-5441**, which delivered pyrrolidine **3a** with high regio- and enantioselectivity (*67*). During the initial round of evolution (Fig. 2a–c), mutation I148T emerged as a key determinant of regioselectivity, shifting the product distribution from 3:97 to 45:55 (**2a**:**3a**). The pronounced evolvability of the ***Tm*Pgb** scaffold prompted us to investigate whether directed evolution could further access regiodivergent C–H amination pathways. After seven rounds of evolution, ***Tm*Pgb-PIP-G7-5448**, bearing 11 mutations (S17G, C30S, F62W, S65G, H66E, G78A, S141N, I148T, Y170H, Q180S and G192S), was obtained.

This variant catalysed formation of piperidine **2a** in 6% assay yield with 84:16 rr and >99% ee under whole *E. coli* cell conditions, corresponding to an ∼80-fold improvement in activity together with reversal of regioselectivity toward **2a**. Further optimisation under purified-enzyme conditions increased the yield to 12.3%.

Together, these results demonstrate that directed evolution can steer the regioselectivity from an intrinsically biased starting point toward a complementary reaction pathway. This raises a deeper question: how do we choose the optimal starting point? Selecting a starting scaffold is essentially a hypothesis about its evolvability rather than an assessment of its present catalytic capabilities. The objective is therefore not to exhaustively search existing libraries for a single optimal scaffold, but to use target-informed criteria to identify or create promising starting points across natural and designed protein space.

### De novo enzyme design of nitrene transferase for regio- and enantioselective C–H amination reaction using AP Novo

Building on the evolutionary trajectory of ***Tm*Pgb-PIP-G7**, we surmised that the observed regioselectivity is primarily dictated by substrate positioning within scaffold-dependent active-site geometries. Although enzymes are inherently programmable through their genetic encoding, the locally accessible catalytic landscape could be constrained by the structural scaffolds available from natural evolution. *De novo* enzyme design aims to explore beyond local optima by constructing tailored evolutionary starting points that access properties beyond those readily attainable from nature.

*De novo* haemoprotein design has served as a benchmark for evaluating emergent catalytic function in artificial enzymes. Early studies largely focused on transformations with established small-molecule catalytic precedents. These reactions, such as styrene cyclopropanation (*21–22*), are already catalysed to an appreciable extent by free haem. In contrast, the desired regioselective C–H amination *via* biocatalytic nitrene transfer is an enzyme-enabled reaction, and represents a substantially more demanding reaction manifold involving high-energy reactive intermediates and stringent control over competing pathways. Such reactivity and selectivity have not been explored by *de novo* enzyme design, nor have they been addressed by small-molecule catalysis. This reaction can thus serve as a cornerstone demonstration of how *de novo* enzymes can extend beyond benchmark reactivity toward sophisticated abiological transformations representing real-world synthetic challenges (Fig. 3a).

**Fig. 3:**
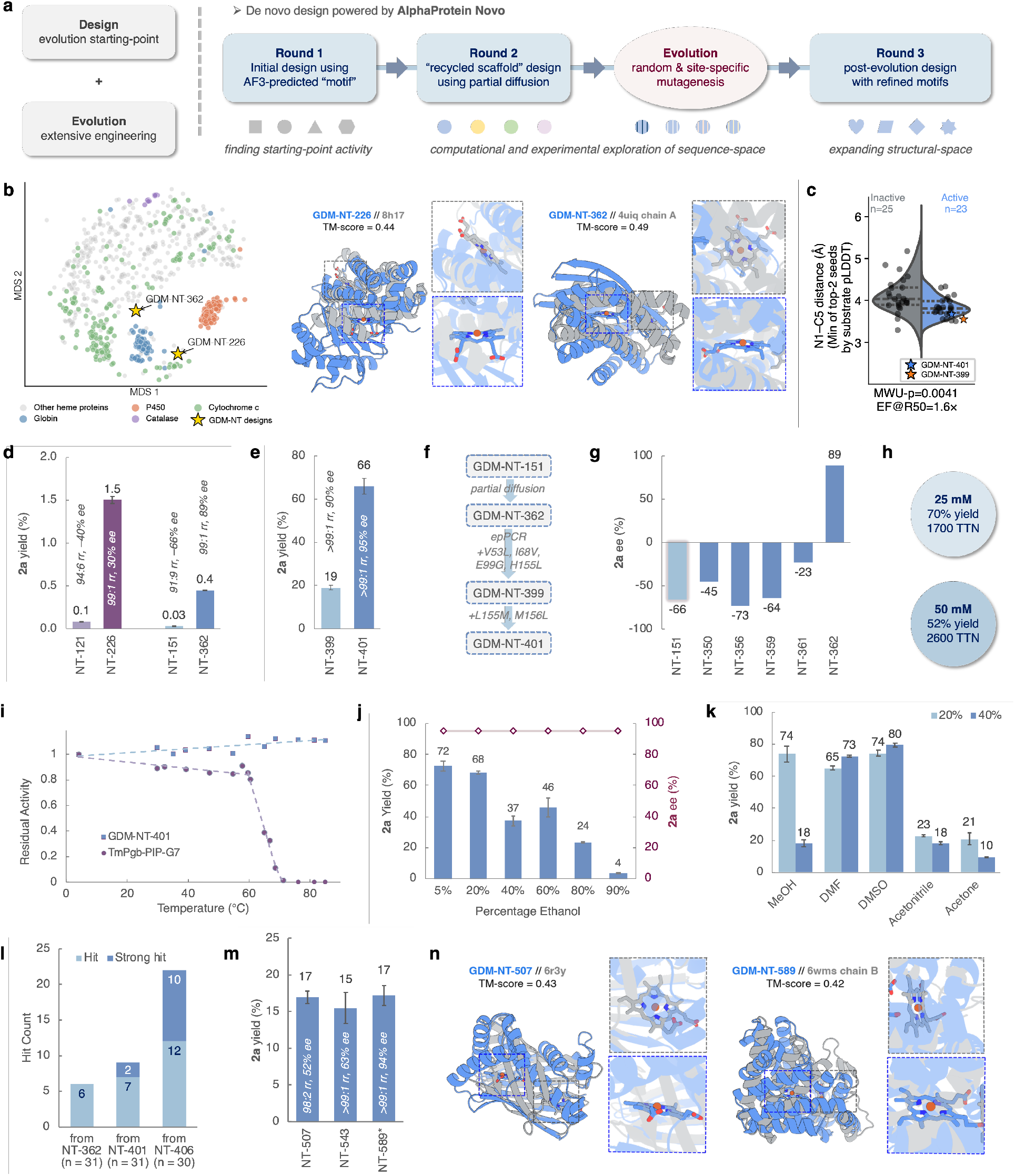
Design-evolution synergy toward architecturally orthogonal nitrene transferases. **a**, Left: previous *de novo* design campaigns often require extensive directed evolution after identifying starting-poiunt activity. Right: this work demonstrates a design-evolution synergy toward architecturally orthogonal nitrene transferases for the synthesis of chiral piperidines. **b**, MDS projection of all-pairs TM-align distances across non-redundant haem-binding chains from the PDB (March 2026), clustered at 70% sequence identity (n=1,016). Each point represents a cluster representative, coloured by family, with **GDM-NT** designs projected into the same space. Note: the cytochrome *c* label is a heterogeneous grouping defined by PDB annotation. (bottom) TM-align superposition of each design (blue) with its closest structural match (gray), showing folds distinct from known haem proteins. Insets show the haem binding pocket of designs and natural proteins. **c**, Distribution of AF3-LA modeled N1–C5 distance (Å) between inactive (n=25) and active (n=23) designs generated via partial diffusion of **GDM-NT-151** and specific mutated variants **(GDM-NT-399** and **GDM-NT-401)**. Sequences are classified as active if they exhibit at least 5% total C–H amination activity relative to wild-type protoglobin from ***T****hermus a****m****yloliquefaciens* (***Tm*Pgb-wt**). Structures of the protein in complex with the haem and alkyl chain were predicted using AF3-LA (*36*) with 8 seeds, with the ligand complex encoded *via* a single SMILES string. Distances are measured between the substrate nitrogen and the fifth carbon (C5), reporting the minimum distance observed within the top two structures as ranked by substrate-atom pLDDT. Active designs demonstrate a statistically significant reduction in N1–C5 distance (Mann–Whitney U test, p = 0.0041), yielding an enrichment factor at 50% recall (EF@R50) of 1.6×. **d**, Redesign using partial diffusion on recycled scaffold leading to improved activity and selectivity. **e**, Minimal engineering effort toward high regio- and enantioselective piperidine synthase **GDM-NT-401. f**, Engineering trajectory toward **GDM-NT-401. g**, Enantiodivergence observed during redesign using partial diffusion. **h**, High concentration experiments using purified **GDM-NT-401. i**, T_50_ (temperature at which 50% activity remains after incubation for 1 h) analysis of **GDM-NT-401** and ***Tm*Pgb-PIP-G7. j**, Stability of **GDM-NT-401** under high concentrations of ethanol. **k**, Stability of **GDM-NT-401** in other water-miscible organic co-solvents. **l**, Comparison of third-round design hit rates using motifs extracted from **GDM-NT-362, GDM-NT-401**, and **GDM-NT-406**. A hit is defined as having >5% of the combined **2a** and **3a** ion count on the SIM channel provided by ***Tm*Pgb-wt** and >1:1 (**2a**:**3a**) regioselectivity. A strong hit is defined as >100% of the combined **2a** and **3a** ion count on the SIM channel and >1:1 (**2a**:**3a**) regioselectivity. **m**, Yield, regioselectivity and enantioselectivity of the three top-performing designs from the third round of design. **n**, TM-align superposition of each design (blue) with its closest structural match (gray), showing folds distinct from known heme proteins. Insets show the haem binding pocket of designs and natural proteins. Reaction conditions: Reactions in panel d, e, h, i, j, k, m were run in duplicate or triplicate. Yields represent average yields between parallel reactions. Error bars represent standard deviation. Reaction conditions used in Fig 3c, e, l : alkyl azide **1** (5.0 mM), whole *E. coli* cells (OD_600_ = 10–15) expressing **GDM-NT** variants, 50 mM M9–N buffer (pH 7.2) on 96-well plates. Reaction conditions used in Fig 3d, g, m: alkyl azide **1** (5.0 mM), whole *E. coli* cells (OD_600_ = 30) expressing **GDM-NT** variants, 50 mM M9–N buffer (pH 7.2) on analytical scale. \***GDM-NT-589** can only be expressed at low cell density, resulting in OD_600_ = 10. Reaction conditions used in Fig 3h–k: alkyl azide **1** (5.0 mM, 50 mM or 100 mM), purified **GDM-NT** variants (haem concentration 10.0 μM), Na_2_S_2_O_4_ (2.5 mM), 50 mM M9–N buffer (pH 7.2) with various co-solvents. Reductant is required for reactions conducted using purified protein (see Supplementary Information, Table S16–S19).

With these considerations in mind, we selected this transformation as an ideal application of AP Novo (*36*). AP Novo improves upon the *in silico* benchmark performance of other generative models and the resulting enzymes demonstrate high *in vitro* catalytic efficiency on model reactions such as Kemp elimination and ester hydrolysis. Using AP Novo, piperidine synthases with novel scaffolds and starting-point activities were generated. Notably, despite modest activity (up to 22 TTN, 99:1 rr, 94% ee), multiple *de novo* enzymes achieved higher regioselectivity for **2a** than the active wild-type protoglobin starting points (*36*). To address catalytic activity limitations, we selected two initial designs, **GDM-NT-121** (1.6 TTN, 94:6 rr, –40% ee) and **GDM-NT-151** (0.9 TTN, 91:9 rr, 66% ee), for refinement *via* partial diffusion, a process that slightly re-noises and then denoises a structure to sample the local structural landscape (Fig. 3b–3d). We term the outputs of this process “recycled-scaffold” designs, as they are generated by optimizing parent designs with the same scaffold.

The set of “recycled-scaffold” designs exhibited markedly improved activity, including **GDM-NT-226** (1.5% yield, 99:1 rr, 29% ee), representing an 18-fold increase in yield relative to its parent **GDM-NT-121**, and **GDM-NT-362** (0.45% yield, 99:1 rr, 88.5% ee) with 15-fold greater yield than **GDM-NT-151** (Fig. 3d). In this design round, we introduced new filters which were predictive of activity in the first round (*36*), including one which evaluates the near-attack positions of the bond-forming carbon and nitrogen atoms (see Supplementary Information). Retrospectively, the distance between the nitrenoid nitrogen center (N1) and the benzylic carbon (C5) was shown to discriminate hits from less optimal designs in the ‘recycled-scaffold’ set based on **GDM-NT-151** (Fig. 3c), consistent with the proximity of these atoms in the hydrogen atom transfer (HAT) transition state (TS) of the proposed mechanism (see Fig. 5a and Supplementary Information, Fig. S14). Compared to natural haemoproteins, including those extensively explored for new-to-nature catalysis (cytochromes P450, cytochromes *c*, and globins (*4*)), these designs adopt architecturally distinct folds with differently positioned haem-binding pockets, with the closest structural match exhibiting a TM-score <0.5 with any protein in the PDB (Fig. 3b). With the discovery of these improved variants situated in a different structural space, we hypothesized that further improvements could be found by exploring this new local landscape by directed evolution.

We thus evaluated the evolvability of the *de novo* enzymes through random mutagenesis and high-throughput plate-based activity screening under whole-cell conditions. Libraries derived from both **GDM-NT-226** and **GDM-NT-362** yielded multiple variants exhibiting greater than two-fold improvements in activity (see Supplementary Information, Fig. S10, S11), suggesting that these *de novo* scaffolds are readily evolvable. Among these lineages, directed evolution of **GDM-NT-362** proved particularly productive (Fig. 3e–f), furnishing variants with more than two orders of magnitude enhancement in normalised activity, with **GDM-NT-399** (**GDM-NT-362**_V53L-I68V-E99G-H155L) giving the best result (19% yield, >99:1 rr, 90% ee). Sequence-structure analysis of the highest activity variants revealed a consistent mutation pattern at the axial His155 residue from the input motif for design. Despite this, the haem cofactor’s predicted position was unchanged, consistent with the sustained high regioselectivity. Prompted by the robustness to mutation in the proximal haem pocket, we further engineered site 155 and adjacent position 156.

To our delight, a collection of variants with different combinations of site 155/156 residues all yielded high activity toward **2a** production (see Supplementary Information, Table S15). On a plate-based screen, **GDM-NT-400** (**GDM-NT-362**_H155M-M156L) featuring methionine as the presumed axial ligand gave **2a** in 26% yield, while the best performing variant, **GDM-NT-401** (**GDM-NT-362**_V53L-I68V-E99G-H155M-M156L) bearing three additional mutations incorporated from random mutagenesis further increased the yield to 36%. This result was further optimised under normalized whole-cell conditions, giving **2a** in 66% yield, 1200 TTN, >99:1 rr, and 95% *ee*. This brief directed evolution campaign from a *de novo* enzyme scaffold suggests that building on a well-defined evolution starting point, in this case a highly regioselective parent design, directed evolution can be drastically accelerated.

Although enantioselectivity was not an explicit design objective, partial diffusion revealed pronounced enantiodivergence, with catalysts favouring both product enantiomers emerging in the second round of design (Fig. 3g). In contrast to small-molecule catalysis, where reversed enantioselectivity can be achieved through straightforward application of a mirror-image catalyst, enantiodivergence in enzyme engineering is considerably more challenging and typically arises through serendipitous mutations identified during directed evolution. The emergence of complementary stereochemical outcomes through local exploration of *de novo* sequence-structure space highlights the unique potential of partial-diffusion-based redesign for optimizing not only activity but otherwise elusive enantioselectivity.

### Stability and synthetic applicability of GDM-NT-401

We next evaluated **GDM-NT-401** under a range of stress conditions. To our delight, this enzyme remains highly productive under high substrate concentrations similar to those used for small-molecule transition-metal catalysts, giving **2a** in 70% yield, 1700 TTN at 50 mM substrate loading and 52% yield, 2600 TTN at 100 mM substrate loading (Fig. 3h). T_50_ measurements were conducted to demonstrate the high thermostability of the designed enzyme. No detectable loss of catalytic activity was observed for **GDM-NT-401** across the tested temperature range (30–85 °C), whereas ***Tm*Pgb-PIP-G7** lost half of its activity at around 65 °C (Fig. 3i). **GDM-NT-401** also demonstrated remarkable tolerance toward organic solvents. Reactions containing up to 60% ethanol still afforded **2a** in 46% yield while maintaining high regio- and enantioselectivity (Fig. 3j). Other commonly used water-miscible co-solvents, including methanol, acetonitrile, dimethyl sulfoxide, *N,N*-dimethylformamide, and acetone, were likewise tolerated at concentrations as high as 40% v/v (Fig. 3k). These properties highlight the capacity of *de novo* protein design to access physicochemical properties beyond those typically associated with naturally evolved haemoproteins.

### Design-evolution synergy

*From design to evolution*: Approximately 90% of natural haem-binding proteins use histidine or cysteine as the protein-derived axial ligand (*68*, see Supplementary Information, Figure. S4). On the other hand, noncanonical replacements of axial ligands are known to arise in nature (*69*) and, more importantly, have proved critical in previous directed evolution campaigns for new-to-nature enzyme activities (*68, 70–71*). To test whether the *de novo* design pipeline is able to propose these unconventional solutions, we generated “no-protein motif” designs with input motifs consisting only of the haem cofactor and *N*-heterocyclic products, removing the explicit axial constraint and found that the designed residues at the conventional axial position strongly favored histidine, reflecting its dominance in natural sequences (see Supplementary Information, Figure. S4). Thus, without further conditioning, the design model remains biased towards the distribution of the natural scaffolds it was trained on. This reveals a complementary relationship between design and directed evolution: design establishes architecturally permissive starting scaffolds encoding the desired regioselectivity, while directed evolution experimentally interrogates solutions outside the training distribution of current design methods.

*From evolution to design*: Moreover, the design-evolution relationship is not a one-way pipeline for generating new evolution starting points, but rather offers reciprocal synergy. We surmised that leveraging this reverse flow of information could be vital for improving initial hit rates, increasing starting-point activities, and generating scaffold diversity where new-to-nature chemistry may emerge. We therefore initiated a third round of novel-scaffold design using “refined motifs” extracted from empirically identified endpoints of directed evolution. Haem-bound piperidine poses and their corresponding proximal axial residues were extracted from AF3-LA predictions of two post-evolution designs **GDM-NT-401** (methionine) and **GDM-NT-406** (leucine), as well as pre-evolution parent scaffold **GDM-NT-362** (histidine) as a control motif (see Supplementary Information, Table S15). While designs derived from **GDM-NT-362** control motif yielded standard hit rates comparable to our previous second round of design (*36*), the evolution-refined-motif approach proved highly effective. Among the designs with refined motifs, 73% derived from **GDM-NT-406** and 29% from **GDM-NT-401** were active. Under a stringent threshold requiring greater than ***Tm*Pgb-wt** total C–H amination activity, the rates were 33% and 6.5%, respectively (Figure 3l). As a control experiment, simply mutating the axial histidine ligand in protoglobins or earlier designed scaffolds (such as **GDM-NT-121** and **GDM-NT-270**) failed to yield similar improvements, confirming that this activity boost stemmed from generating custom proteins around the refined motif rather than simple ligand substitution. Ultimately, even without further directed evolution, the leading designs **GDM-NT-507, GDM-NT-543** and **GDM-NT-589** already achieved high activity with strong regio- and enantioselectivity (Fig. 3m) and substantial scaffold diversity compared to natural enzymes (Fig. 3n and Supplementary Information, Figure S2).

Notably, 13 out of the 28 initial designs described in the AP Novo paper (*36*), including the most active designs **GDM-NT-121** and **GDM-NT-151**, adopted mixed α/β architectures. This enrichment was also observed in subsequent rounds of designs (see Supplementary Information, Figure S3), expanding beyond the predominantly alpha-helical folds of previous designed (*7, 21*) and natural haemoproteins, including globins, cytochromes, and peroxidases. This further highlights the capacity of *de novo* enzyme design to access structural solutions beyond the architectural constraints typically observed in natural haemoproteins, particularly in the context of demanding new-to-nature transformations (*21–23*). Although the origins of AP Novo’s enrichment of α/β folds remain unclear, the comparatively rigid structural organisation afforded by β-sheets (*72*) may contribute to the precise substrate positioning required for high regioselectivity.

Together, these results demonstrate a reciprocal design-evolution synergy in which *de novo* design searches broad scaffold space to create well-positioned evolutionary starting points, directed evolution interrogates their local sequence-fitness landscapes with only minimal steps required, and the solutions uncovered through evolution, in turn, guide subsequent design campaigns toward better hit rate, higher initial activity, and better scaffold diversity.

### Synthetic application of GDM-NT-401 in biocatalytic regio- and enantioselective C–H amination reactions

Having identified **GDM-NT-401** as an optimal catalyst, we next evaluated the scope and limitations of the biocatalytic piperidine synthesis under whole-cell conditions (Fig. 4a). A range of aryl substituents with diverse steric and electronic properties were investigated. Under standard conditions, 2-phenylpiperidine **2a** was obtained in 66% assay yield, corresponding to 1200 TTN, with >99:1 rr and 95% ee. The yield could be further increased to 88% using purified enzyme at 10 μM catalyst loading. Preparative-scale synthesis on 1-mmol scale afforded **2a** in 84% isolated yield while maintaining excellent regio- and stereoselectivity (see Supplementary Information for details). The modular substrate synthesis route using NaN_3_ as nitrogen source also enabled single-atom radiolabeling, furnishing the ^15^N-labelled product **2b** in 59% yield with preserved regio- and enantioselectivity. Substrates bearing *para*-substitutions gave variable yields from 5% to 55% (**2c**– **2g**). We attribute this variation primarily to steric interactions between substituted aryl groups and the protein scaffold, which were tailored for an unsubstituted substrate **1a** during the motif scaffolding step. Nevertheless, further engineering in this region could address a specific synthetic demand when desired. Meanwhile, *meta*-substituted substrates were also tolerated, affording products **2h**–**2k** in moderate yields, while sterically demanding 2-naphthyl substrate furnished piperidine **2l** in diminished yield. Notably, excellent regioselectivities were obtained despite substantial variation in steric and electronic properties, highlighting the reliable control over substrate positioning inside the enzyme active site. Good to excellent enantioselectivities were likewise observed throughout the substrate scope, demonstrating great potential in synthetic applicability. As a direct comparison, we performed parallel experiments using purified ***Tm*Pgb-PIP-G7**, the product of directed evolution of a natural protein. Across the examined substrate series, ***Tm*Pgb-PIP-G7** delivered the corresponding piperidines with consistently excellent enantioselectivities (>99% ee) favouring the opposite (*S*)-enantiomers, albeit exhibiting overall diminished substrate-dependent yields and regioselectivities. Although additional directed evolution could likely further enhance the activity and substrate promiscuity of this lineage, these results underscore the critical role of scaffold selection in biocatalyst development. As directed evolution remains inherently resource- and time-intensive, the accessible catalytic landscape is often strongly biased by the structural and mechanistic predispositions of the starting scaffold. Consequently, identifying privileged scaffolds capable of supporting productive reactivity at early stages can have profound impact for the eventual catalytic outcome, influencing the attainable levels of activity, selectivity, and substrate compatibility.

**Fig. 4:**
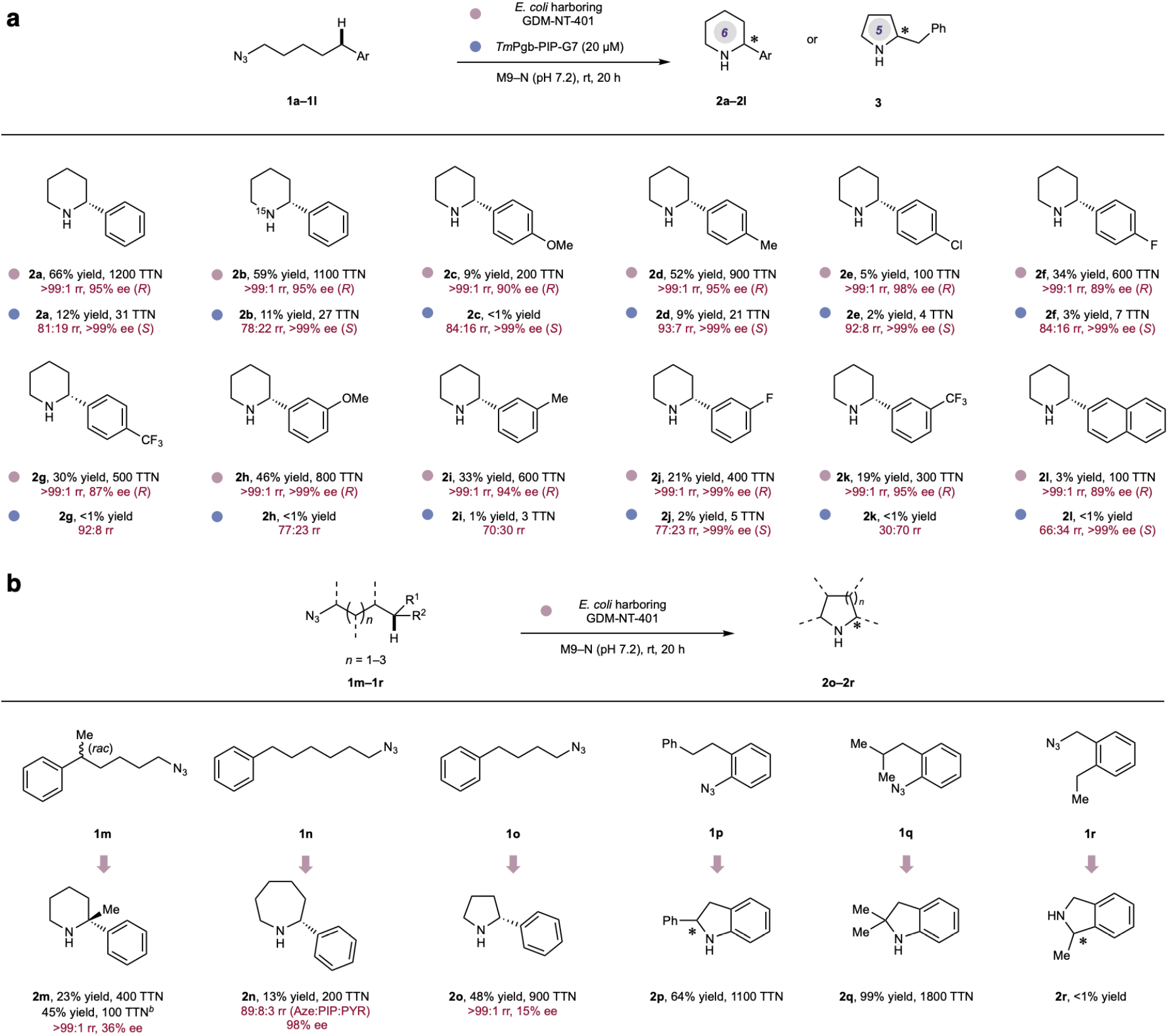
Substrate Scope of Intramolecular C–H Amination Reactions for the Preparation of Chiral *N*-Heterocycles. **a**, Substrate scope study for the synthesis of chiral 2-arylpiperdines. **b**, Substrate scope study with backbone diversity. Reaction conditions using **GDM-NT-401**: alkyl azide **1** (5.0 mM), whole *E. coli* cell (OD_600_ = 30) expressing **GDM-NT-401** (haem concentration 2.8 μM), 50 mM M9–N buffer (pH 7.2). Reaction conditions using ***Tm*Pgb-PIP-G7**: alkyl azide **1** (5.0 mM), purified ***Tm*Pgb-PIP-G7** (haem concentration 10.0 μM), 50 mM M9–N buffer (pH 7.2).

The high catalytic efficiency of **GDM-NT-401** prompted us to further evaluate its performance against structurally diverse alkyl and aryl azide substrates (Fig. 4b). Despite these substrates deviating substantially from the original design target, the enzyme retained appreciable catalytic activity across multiple substrate classes, highlighting that *de novo* enzymes can exhibit significant breadth in substrate range rather than being restricted to an exquisitely crafted model reaction. We speculate that the high intrinsic reactivity of **GDM-NT-401** contributes to this behaviour by enabling productive turnover even when substrate-binding geometries are not fully optimised. Using benzylic-substituted substrate **1m**, piperidine **2m** bearing an α-to-nitrogen quaternary stereocentre was obtained in 23% yield under whole-cell conditions and 45% yield using purified enzyme, albeit with modest enantioselectivity (36% ee). We next challenged the catalyst with substrate **1n**, where multiple cyclisation manifolds leading to azepane, piperidine, or pyrrolidine products are all mechanistically feasible. Prior to experimentation, analysis of the AF3-LA model of the putative alkyl nitrene intermediate within the enzyme active site suggested preferential formation of the seven-membered azepane product (see Supplementary Information, Figure S8). Consistent with this prediction, wet-lab experiments furnished 2-phenylazepane **2n** with high regioselectivity (89:8:3 azepane:piperidine:pyrrolidine) and 98% ee, demonstrating that the engineered active site can exert substantial control over competing radical cyclisation trajectories even in structurally flexible systems. Shortening the alkyl tether redirected cyclisation toward formation of 2-phenylpyrrolidine **2o**, which was obtained in 48% yield and 15% ee. Moreover, the reaction is not restricted to alkyl azides. Substituted aryl azides underwent efficient intramolecular C–H amination to furnish indolines **2p**–**2q** in high yields, whereas formation of the analogous isoindoline **2r** from benzyl azide was observed only in trace amounts.

### Computational analysis of TmPgb-PIP-G7 and GDM-NT-401 for the origin of regioselectivity

To better understand the origin of the high regioselectivity obtained using the *de novo*-designed **GDM-NT-401**, we computationally investigated how **GDM-NT-401** and the evolved ***Tm*Pgb-PIP-G7** variant each control regioselectivity. Although the benzylic ε-C–H bond is 10 kcal mol^−1^ weaker than the homobenzylic δ-C–H bond (Fig. 5a), previous catalytic systems generally provide only modest regioselectivity (*52, 54–55*), suggesting that bond dissociation energy (BDE) is not the only factor determining the outcome of the competing 1,6- and 1,5-HAT pathways. Consistent with this view, our density functional theory (DFT) calculations using a minimal iron porphyrin model revealed that the BDE difference is substantially attenuated at the corresponding transition states (ΔΔ*G*^‡^ = 3.2 kcal/mol, Fig. 5a). More importantly, the two pathways proceed through markedly different transition-state geometries. In the 1,6-HAT TS, the Ph group is placed in proximity to the haem cofactor, whereas the same Ph is allowed to move further away from the haem and adopt a more extended conformation in the homobenzylic 1,5-HAT TS. These distinct geometries provide the protein environment with a structural handle to differentiate the competing pathways through enzyme engineering or *de novo* design.

**Fig. 5:**
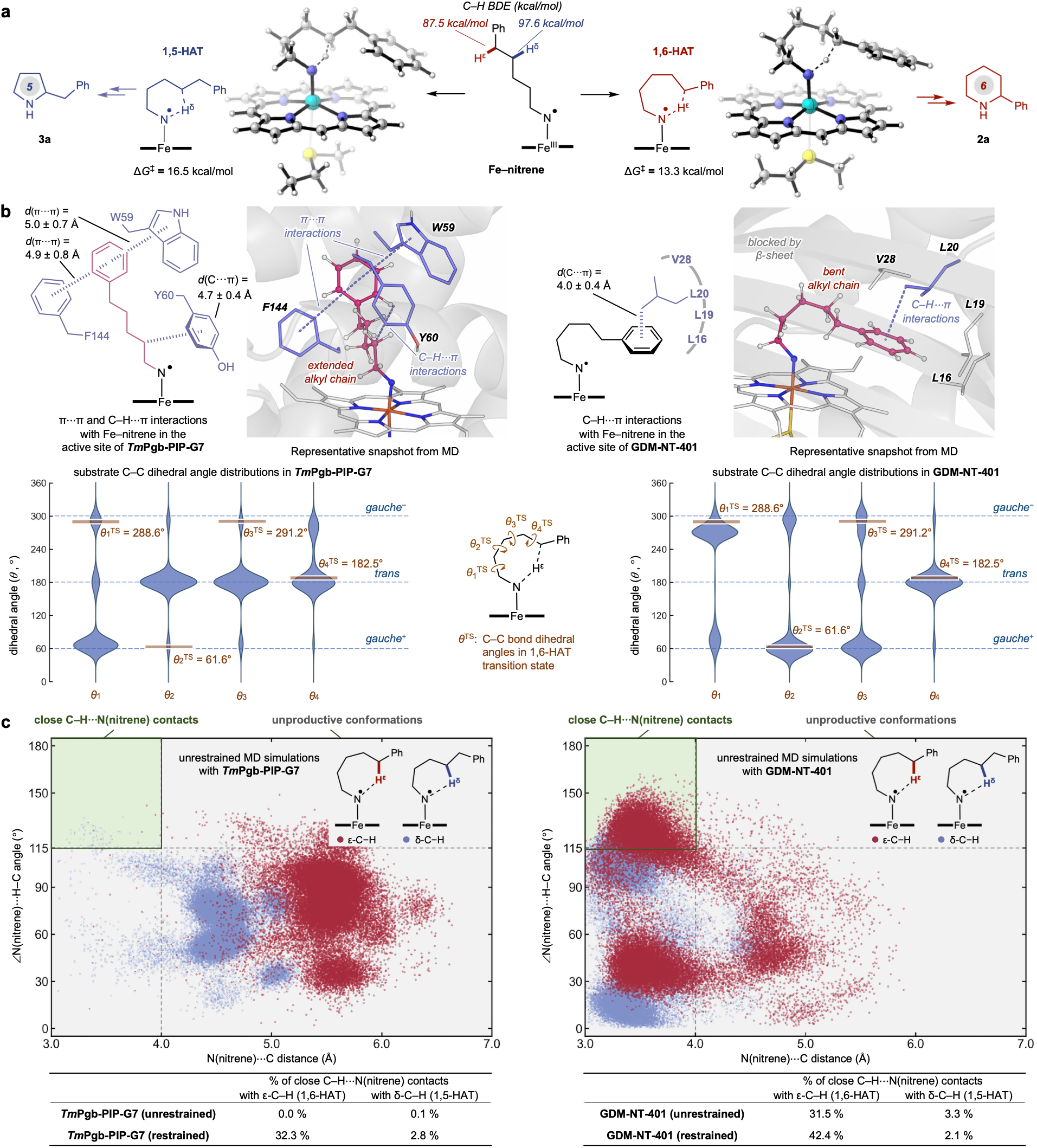
Computational analysis of enzymatic control of regioselectivity in C–H amination with *Tm*Pgb-PIP-G7 and GDM-NT-401. **a**, DFT-computed intrinsic regioselectivity of C–H amination and regioselectivity-determining 1,5-HAT and 1,6-HAT transition states; **b**, Preferred binding poses of the Fe–nitrene intermediate in the active sites of the evolved ***Tm*Pgb-PIP-G7** and the *de novo*-designed **GDM-NT-401**. Dihedral angle distributions from unrestrained MD simulations were compared with those in DFT-optimised 1,6-HAT transition state structure; **c**, Close contacts between the nitrene N atom and the ε- and δ-C–H bonds observed in the MD simulations.

To understand how the active site pockets affect the regioselectivity, three independent replicas of 500 ns classical molecular dynamics (MD) simulations of the Fe(III)–nitrene intermediate were performed in the evolved ***Tm*Pgb-PIP-G7** and in the *de novo*-designed **GDM-NT-401**. In ***Tm*Pgb-PIP-G7**, the alkyl chain of the nitrene favours an extended geometry with most of the rotatable C(*sp*^3^)–C(*sp*^3^) bonds adopting the *trans* conformation (*θ*_2_, *θ*_3_, *θ*_4_ ≈ 180°, defined in Fig. 5b). This places the Ph group on the alkyl chain in the hydrophobic pocket surrounded by Y60, W59, and F144 residues, forming stabilising π⋯π and C–H· ·π interactions. By contrast, in **GDM-NT-401**, the nitrene alkyl chain favours a bent geometry with most C–C bonds (*θ*_1_, *θ*_2_, *θ*_3_) in either *gauche*^+^ or *gauche*^−^ conformation and the terminal C^δ^–C^ε^ bond (*θ*_4_) in the *trans* conformation. These dihedral distributions closely match those of the DFT-computed 1,6-HAT TS, the exception being *θ*_3_, which remains flexible and samples multiple conformations in MD simulations. The resemblance between the preferred conformation of the Fe–nitrene intermediate in **GDM-NT-401** and the 1,6-HAT TS indicates that the designed pocket, specifically using **2a** as part of the input motif, effectively preorganises the intermediate for 1,6-HAT. Additionally, this preorganised geometry is stabilised by the interactions of the Ph group on the alkyl chain in a hydrophobic pocket surrounded by residues V28, L20, L19, and L16, which was first generated by the initial round of AP Novo design (**GDM-NT-151**) and subsequently refined during partial diffusion (**GDM-NT-362**, adding L28V mutation), highlighting the complementary benefit of both design strategies.

To investigate how the binding poses affect the interactions of the nitrene centre with the ε- and δ-C–H bonds, we analysed the close contacts between the nitrene N atom and these C–H bonds observed in the MD simulations (Fig. 5c). Here, a close contact is defined as when the N⋯C distance is shorter than 4.0 Å and the ∠N⋯H–C angle is greater than 115°. In the unrestrained simulations with ***Tm*Pgb-PIP-G7**, close contacts were essentially not observed with ε-C–H and only observed in 0.1% of the MD frames with δ-C–H, indicating substantial conformational changes would be needed to achieve either 1,6- or 1,5-HAT TS geometry. In contrast, 31.5% of unrestrained MD frames with **GDM-NT-401** show close contacts with ε-C–H, whereas only 3.3% have close contacts with δ-C–H. Therefore, the nitrene binding pose in **GDM-NT-401** not only enables a bent alkyl chain geometry necessary for the HAT transition state, but also positions the benzylic ε-C–H closer to the reactive nitrene centre, thereby enhancing the kinetic regioselectivity for 1,6-HAT. To further mimic Fe–nitrene/C–H interactions near the HAT transition states, a distance restraint (2.9–3.5 Å, force constant 100 kcal mol^−1^ Å^−2^) was applied between the nitrene nitrogen and the centroid of the δ- and ε-carbons. In these restrained MD simulations, ***Tm*Pgb-PIP-G7** populated close contacts with ε-C–H (1,6-HAT) in 32.3% of the MD frames and with δ-C–H (1,5-HAT) in 2.8% of the frames, whereas **GDM-NT-401** gave a higher ratio favoring ε-C– H (42.4% compared with 2.1% for δ-C–H. See Supplementary Information Fig. S28 for more details), consistent with the observed regioselectivity.

Taken together, these results indicate that the *de novo*-designed active site pocket of **GDM-NT-401** enhances 1,6-HAT regioselectivity by conformational preorganization enabled by a rationally designed input motif. It enforces a bent nitrene conformation that preorganises the benzylic ε-C– H towards the nitrene centre, whereas the extended pose enforced by the evolved ***Tm*Pgb-PIP-G7** active site requires substantial reorganisation to reach either HAT transition state. This contrast highlights a central advantage of mechanism-guided *de novo* enzyme design. Directed evolution can efficiently interrogate local sequence space once a suitable scaffold has been identified, but its success depends on the accessibility of the pre-existing landscape. *De novo* design addresses the upstream problem by allowing a mechanism-informed motif to guide scaffold construction rather than relying exclusively on architectures represented in natural proteins.

## Conclusion

In conclusion, we demonstrate that the combination of *de novo* enzyme design and directed evolution can enable new-to-nature chemistry beyond the reach of existing small-molecule and biocatalytic paradigms. Through this integrated approach, we developed a regio- and enantioselective intramolecular C–H amination reaction for the synthesis of chiral piperidines, a transformation requiring precise control over highly reactive intermediates and competing reaction pathways. To our knowledge, this reactivity has not previously been achieved in either transition-metal catalysis or biocatalysis.

More broadly, this work highlights the synergistic relationship between *de novo* design and directed evolution. Whereas *de novo* design expands the accessible structural landscape by generating catalytically permissive protein architectures beyond those found in nature, directed evolution efficiently navigates local sequence space to optimise activity and selectivity. The real-world improvements from directed evolution can then directly benefit *de novo* design through motif refinement, as demonstrated by the substantially higher hit rate and better initial activity. The rapid emergence of a highly active and selective piperidine synthase after only two rounds of design and engineering illustrates the power of combining three-dimensional sequence-structure-function exploration with conventional sequence-function screening. We anticipate that this design-evolution paradigm will provide a general framework for developing new enzyme scaffolds and catalytic functions, enabling increasingly sophisticated transformations that remain inaccessible to both naturally occurring enzymes and existing synthetic catalysts.

## Data availability

All data supporting the findings of this study are available in the main text and the Supplementary Information. Additional data are available from the corresponding authors upon request.

## Acknowledgement

This work is supported by US Department of Energy, Office of Science, Office of Basic Energy Sciences (grant DE-SC0021141 to F.H.A.), and the National Institutes of Health (grant R35 GM128779 to P.L.). DFT and MD simulations were carried out at the University of Pittsburgh Center for Research Computing and Data, supported by NSF award number OAC-2117681, and the Advanced Cyberinfrastructure Coordination Ecosystem: Services & Support (ACCESS) through allocation CHE140139, which is supported by NSF grants OAC-2138259, OAC-2138286, OAC-2138307, OAC-2137603, and OAC-2138296. We thank Dr. Sabine Brinkman-Chen for careful proofreading of the manuscript and Dr. Scott C. Virgil for assistance with HRMS analysis.

## Author contribution

Z.Q.L. and F.H.A. conceived the project. Z.Q.L., Z.Y.Q., O.M., and D.H. performed the directed evolution campaign. R.K.Z., Z.W., and J.K. performed the *de novo* protein design, with contribution from J.A. and S.S. P.P.X. performed the molecular dynamics study. Y.Z. performed the DFT study. Z.Q.L. performed the experimental evaluation of designs, with contribution from T.L. and A.E.D. Z.Q.L. wrote the manuscript with substantial input from P.P.X. and contributions from all authors. J.W., P.K., and Z.W. supervised the *de novo* design effort. P.L. supervised the computational studies. F.H.A. supervised the overall project.

R.K.Z., P.P.X., J.K., Y.Z., and Z.Y.Q contributed equally and are free to list their name in the front.

## Competing interest

The authors declare no competing interest.

## Additional information

**Supplementary Information** is available in the online version.

## Notes

### Competing Interest Statement

The authors have declared no competing interest.

