## supplementary information for "Design-Evolution Synergy Enables New-to-Nature Enzyme Chemistry"

#### General Information

##### Analytical Instruments and Source of Commercial Reagents

Unless otherwise stated, all materials were used as received from commercial sources without further purification. Erlenmeyer flasks used for cell-culture and protein expression were purchased from VWR and autoclaved before use. Glassware used for analytical reaction set-up and HPLC analysis were purchased from Agilent and used as received. All solvents were purchased from MilliporeSigma, VWR, Thermofisher and used as received. Silica gel chromatography was carried out using Biotage® Sfär chromatography column. $^{1}H$ and $^{13}C$ NMR spectra were recorded on a Bruker Prodigy 400 MHz instrument (400 MHz for $^{1}H$ NMR, 101 MHz for $^{13}C$ NMR, and 376 MHz for $^{19}F$ NMR). $^{1}H$ NMR spectra were reported relative to $Me_4Si$ (δ 0.0) in $CDCl_3$ or residual solvent signals (DMSO-$d_5$ at δ 2.54 or HDO at δ 4.79) unless otherwise stated. $^{13}C$ NMR spectra were calibrated to residual solvent signals ($CDCl_3$ at δ 77.16 or DMSO-$d_6$ at δ 39.52). Data for $^{1}H$ NMR are reported as follows: chemical shift (δ ppm), multiplicity (s = singlet, d = doublet, t = triplet, q = quartet, p = pentet, sext = sextet, m = multiplet, dd = doublet of doublets, dt = doublet of triplets, ddd = doublet of doublet of doublets), coupling constant (Hz), integration. Sonication was performed using a Qsonica Q500 sonicator. High-resolution mass spectra were obtained at the California Institute of Technology Mass Spectral Facility. Samples were analyzed by field ionization (FI) using a JEOL AccuTOF GC-Alpha (JMS-T2000GC) mass spectrometer interfaced with an Agilent 8890 GC system. Ions detected by FI are radical cations.

Chemical reactions were monitored using thin layer chromatography (Merck 60 silica gel plates) and a UV lamp for visualization. Analytical reverse-phase high-performance liquid chromatography-mass spectroscopy (HPLC-MS) was carried out using an Agilent 1260 series instrument and an Agilent C18 column (InfinityLab Poroshell 120 EC-C18, 4.6 × 50 mm, 2.7 µm; Part Number: 699975-902T) with water and acetonitrile, both containing 0.1% acetic acid, as the mobile phase. Analytical chiral polar protic HPLC-MS was conducted using an Agilent 1260 series instrument with methanol (containing 0.2 w% ammonium formate) and water (pH 4.0, with 20 mM ammonium formate). Enantiomers were separated using *dual* Agilent InfinityLab Poroshell 120 Chiral-V, 2.7 µm, 2.1 × 150 mm.

*Escherichia coli* cells were grown using Luria-Bertani or Terrific Broth medium (Research Products International) with 0.1 mg/mL ampicillin ($LB_{amp}$ or $TB_{amp}$). Phusion polymerase, and *Taq* polymerase were purchased from New England Biolabs (NEB, Ipswich, MA). Plasmids were purified by miniprep (New England Biolabs). Sanger sequencing was performed by Laragen, Inc. (Culver City, CA) using the T7 promoter primer and T7 terminator primers. Full plasmid sequencing was performed by Plasmidsaurus Inc. (Monrovia, CA). M9–N minimal media (abbreviated as M9–N buffer, pH 7.2) were used as buffering systems for whole cells. M9–N buffer contains 47.7 mM $Na_2HPO_4$, 22.0 mM $KH_2PO_4$, 8.6 mM NaCl, 2.0 mM $MgSO_4$, and 0.1 mM $CaCl_2$.

##### Cloning and Mutagenesis

***Tm*Pgb-PIP** variants:

All genes were encoded with a C-terminal 6×His tag for purification and inserted between the *Nde*I and *Xho*I restriction sites in the pET22b(+) vector.

**GDM-NT** variants:

DNA for clonal genes was purchased from Twist Bioscience (CA, USA). Genes were inserted into a modified pTriEx-4 vector to contain an N-terminal 8×His tag and HRV-3C cleavage site. For all genes, codons were optimised for *E. coli* using Twist Bioscience.

**Table S1**. Primers used in site-saturation mutagenesis and error-prone PCR for ***Tm*Pgb-PIP** variants.

| Primers | Sequence (5' → 3') |
|---|---|
| **Universal Forward (UniF)** | CCA ACT TAC TTC TGA CAA CGA TCG GAG GAC CGA AGG AGC TAA CCG CTT TTT TGC |
| **Universal Reverse (UniR)** | CGA TCG TTG TCA GAA GTA AGT TGG CCG CAG TGT TAT CAC TCA TGG TTA TGG CAG |
| **pET-005** | GAA ATA ATT TTG TTT AAC TTT AAG AAG GAG ATA TAC ATA TG |
| **pET-006** | GCC GGA TCT CAG TGG TGG TGG TGG TGG TGC TCG AG |
| **pET-007** | CAT ATG TAT ATC TCC TTC TTA AAG TTA AAC AAA ATT ATT TC |
| **pET-008** | CTC GAG CAC CAC CAC CAC CAC CAC TGA GAT CCG GC |

**Table S2**. Primers used in error-prone PCR for **GDM-NT** variants.

| Primers | Sequence (5' → 3') |
|---|---|
| **pTriEx-005** | GGTCTGGAAGTGCTGTTTCAAGGCCCGGGATCCAGTGGC |
| **pTriEx-006** | CAGCCTGCACCTGAGGTTAATCACTTAGTGTGATC |
| **pTriEx-007** | ATTAACCTCAGGTGCAGGCTGCCTATCAGAAGG |
| **pTriEx-008** | AAACAGCACTTCCAGACCGCTACTGCCATGATGGTG |

*Site-saturation mutagenesis*

Site-saturation mutagenesis was performed using the 22-codon method (*1*). Primers including the three 22-codon trick codons (NDT, VHG, and TGG) and primers within the ampicillin resistance cassette (**UniF** and **UniR**) were used to amplify the plasmid of interest in two pieces, with an overlap for Gibson assembly[2] in the gene encoding the protein of interest (excluding where the mutations are introduced) and an overlap for Gibson assembly in the gene encoding β-lactamase (which confers ampicillin resistance). This two-piece assembly can prevent misassembled constructs from presenting antibiotic resistance. The PCR reactions were performed according to the protocol in **Table S3** and **Table S4**. The PCR products were digested with *Dpn*I (1 μL) at 37 ºC for at least 1 hour. The resulting PCR products with SYBR Gold nucleic acid stain were loaded on 1.5% agarose gels, purified via electrophoresis, and visualized on a blue transilluminator. The DNA bands of interest were excised, and the DNA was extracted with New England Biolabs DNA cleanup and gel extraction kit. The two linear PCR products for a given site-saturation library were assembled via Gibson assembly.

**Table S3.** PCR reaction components for SSM using Phusion polymerase.

| PCR Reaction Components | Amount (μL) |
|---|---|
| PCR water | 32 |
| 5× HF buffer | 10 |
| 10 mM dNTPs mix | 1 |
| DMSO | 0.5 |
| 10 μM Forward Primer | 2.5 |
| 10 μM Reverse Primer | 2.5 |
| 30 ng/μL Plasmid | 0.5 |
| Phusion® High-Fidelity polymerase | 0.7 |

**Table S4.** Thermocycler conditions for Phusion PCR amplification

| Step | Temperature (°C) | Time (min:s) | Number of Cycles |
|---|---|---|---|
| Initial Denaturation | 98 | 00:30 | 1 |
| Denaturation | 98 | 00:10 | |
| Annealing | 55 | 00:20 | 30 |
| Extension | 72 | 02:30 | |
| Extension | 72 | 10:00 | 1 |
| Hold | 10 | - | |

*Random mutagenesis*

Random mutations using error-prone PCR (*ep*PCR) were introduced using Taq polymerase with the addition of varying $MnCl_2$ concentrations (300–600 μM) following a reported protocol. Primers **005** and **006** were used to amplify the enzyme's cassette in the plasmid of interest. The PCR reactions were performed according to the protocol in **Table S5** and **Table S6**. The amplified fragments were purified by gel electrophoresis (1.5% agarose gel) with SYBR Gold nucleic acid stain. The DNA bands of interest were excised, and the DNA was extracted with New England Biolabs DNA cleanup and gel extraction kit. The PCR product was cloned into an empty pET22b (+) or pTriEx vector between primer sites **007** and **008** via Gibson assembly. The empty pET22b (+) or pTriEx vector can be generated from the parent plasmid using primers **007** and **008** following slightly modified protocol in **Table S3** and **Table S4** (4 min instead of 2.5 min extension time). Libraries generated at different $MnCl_2$ concentrations were tested to determine the optimal balance of high gene diversity and low rate of inactivation. The chosen library was further tested to find improved variants.

**Table S5:** PCR reaction components for *ep*PCR.

| **Reaction Components** (μL) | $MnCl_2$ Concentration (μM) | | | |
|---|---|---|---|---|
| | 300 | 400 | 500 | 600 |
| 10–30 ng/μL Plasmid | 1 | 1 | 1 | 1 |
| 10 mM Forward Primer **005** | 2 | 2 | 2 | 2 |
| 10 mM Reverse Primer **006** | 2 | 2 | 2 | 2 |
| 10 mM dNTPs mix | 4 | 4 | 4 | 4 |
| 1 mM $MnCl_2$ | 30 | 40 | 50 | 60 |
| 10× Standard Taq buffer | 10 | 10 | 10 | 10 |
| Taq DNA polymerase | 1.6 | 1.6 | 1.6 | 1.6 |
| PCR water | 49.4 | 39.4 | 29.4 | 19.4 |

**Table S6:** Thermocycler conditions for Taq PCR amplification

| Step | Temperature (°C) | Time (min:s) | Number of Cycles |
|---|---|---|---|
| Initial Denaturation | 95 | 05:00 | 1 |
| Denaturation | 95 | 00:30 | |
| Annealing | 55 | 00:30 | 30 |
| Extension | 72 | 01:00 | |
| Extension | 72 | 05:00 | 1 |
| Hold | 10 | - | |

*Staggered extension process (StEP) PCR*

Staggered extension process (StEP) PCR was performed using an equimolar mixture of plasmids of the variants to be recombined (*2*). This method recombines mutations that lead to improvements in function with high efficiency by performing PCR cycles with varying extension temperatures and shorter extension times. This protocol involves two PCR reactions. Primers **005** and **006** (1 μM) were used to amplify the region coding the enzyme of interest using the protocol in **Table S7** and **Table S8**. Multiple reactions are set up at varying annealing temperatures (50 → 70 ºC). The PCR product was digested with *Dpn*I (1 μL) at 37 ºC for at least 1 hour. The resulting product with SYBR Gold nucleic acid stain was loaded on 1.5% agarose gels and visualized on a blue transilluminator. The reaction that works at the lowest temperature is typically moved forward to the second PCR. The DNA band of interest was excised and extracted with New England Biolabs DNA cleanup and gel extraction kit. The resulting DNA product is further amplified using the protocol described for site-saturation mutagenesis (**Table S3** and **Table S4**) using primers **005** and **006** (10 μM). The resulting product with SYBR Gold nucleic acid stain was loaded on 1.5% agarose gels and visualized on a blue transilluminator. The DNA band of interest was excised, and the DNA was extracted with New England Biolabs DNA cleanup and gel extraction kit. The PCR product was cloned into a pET22b (+) vector between primer sites **007** and **008** via Gibson assembly (*3*).

**Table S7:** PCR reaction 1 components for StEP recombination.

| PCR reaction components | Amount (μL) |
|---|---|
| PCR water | 14.27 (adjustable) |
| 10× Standard Taq buffer | 2 |
| 10 mM dNTPs mix | 0.4 |
| Forward Primer (1 μM) | 1 |
| Reverse Primer (1 μM) | 1 |
| Mix Plasmids (250 ng total) | 1 (adjustable) |
| Taq Polymerase | 0.33 |

**Table S8:** Thermocycler conditions for PCR reaction 1 in StEP recombination.

| Step | Temperature (ºC) | Time (min:s) | Number of Cycles |
|---|---|---|---|
| Initial Denaturation | 95 | 5:00 | 1 |
| Denaturation | 95 | 00:10 | 120 |
| Annealing | 55 | 00:01 | |
| Extension | Gradient (50 → 72) | 00:01 | |
| Hold | 10 | - | |

#### Transformation of BL21 *E. coli* cells

*Transformation of individual variants:*

Plasmids constructed from Gibson assembly were used to transform *E. coli* T7 express BL21(DE3) electro-competent cells by electroporation. SOC medium was added to rescue the cells. Subsequently, the cells were incubated at 37 °C for 30 minutes before being plated on Luria Bertani medium (Research Products International) supplemented with carbenicillin (0.1 mg/mL) agarose plates. The agarose plates with the plated cells are incubated at 37 ºC for 12–16 h before the libraries are picked.

*Transformation of multiple variants on 96-well plate:*

100 μL of T7 Express chemically competent cells were aliquoted into a PCR strip on ice. Using a multichannel pipette, 5 μL of cells and 1 μL of DNA construct were added to each well of a sterile, cone-shaped 96-well plate. Following a 30-minute incubation on ice, the plate was heat-shocked at 42 °C for 10 seconds and immediately returned to ice for 5 minutes. Cells were rescued with 100 μL of SOC medium and incubated at 37 °C for 30 minutes for outgrowth. Following outgrowth, 10 μL of each culture was inoculated into a 96-deep well plate preloaded with 400 μL/well of LB medium supplemented with ampicillin (100 μL/mL) ($TB_{amp}$). These pre-cultures were incubated at 37 °C, 80% humidity, and 220 rpm shaking speed for 7–8 hours. Subsequently, 50 μL of each pre-culture was used to inoculate a secondary 96-deep well plate containing 950 μL/well of Terrific Broth supplemented with 100 μg/mL ampicillin ($TB_{amp}$) per well. This expression culture was shaken at 37 °C and 220 rpm until an $OD_{600}$ of 0.8–1.0 was reached (typically 2–2.5 hours). The plates were then chilled on ice for 30–45 minutes and induced with 0.5 mM β-d-1-thiogalactopyranoside (IPTG) and 1.0 mM 5-aminolevulinic acid (ALA) (final concentrations). Protein expression was conducted at 22 °C and 220 rpm for 16–20 hours.

#### Cell Culture and Protein Expression

*Analytical-scale protein expression on 96-Well Plate*

Single colonies from $LB_{carb}$ agar plates were picked using sterile toothpicks and cultured in deep-well 96-well plates containing $LB_{amp}$ (500 μL/well) at 37 ºC, 80% humidity, and 220 rpm shaking speed for 7–8 hours. Subsequently, $TB_{amp}$ (950 μL/well) in a deep-well plate was inoculated with an aliquot (50 μL/well) of these pre-cultures and allowed to shake for 2–2.5 hours at 37 ºC, 80% humidity, and 220 rpm until an optical density at 600 nm ($OD_{600}$) of 0.8–1.0. The plates were then rested on ice for 30–45 minutes and induced with 0.5 mM isopropyl *β*-D-1-thiogalactopyranoside (IPTG) and 1.0 mM 5-aminolevulinic acid (ALA) (final concentrations). Protein expression was conducted at 22 ºC, and 220 rpm for 16–20 hours.

*Mid-scale protein expression in Erlenmeyer flasks*

Single colonies from $LB_{carb}$ agar plates were picked using sterile toothpicks and cultured in a culture tube containing $LB_{amp}$ (5 mL/tube) at 37 ºC with 220 rpm shaking speed for 7–8 hours. Subsequently, $TB_{amp}$ (50 mL) in a 125-mL Erlenmeyer flask was inoculated with an aliquot (1 mL) of these pre-cultures and allowed to shake for 2–2.5 hours at 37 ºC, and 220 rpm until an optical density at 600 nm ($OD_{600}$) of 0.8–1.0. The culture was then rested on ice for 30–45 min and induced with 0.5 mM IPTG and 1.0 mM ALA (final concentrations). Protein expression was conducted at 22 ºC and 180 rpm for 16–20 hours.

#### Protein Purification

The engineered protoglobin variants were expressed in *Escherichia coli* BL21(DE3) cells following slightly modified procedure in section C. A preculture of engineered protoglobin variants were cultured in an Erlenmeyer flask containing $LB_{amp}$ (50 mL) at 37 ºC with 220 rpm shaking speed for 6 hours. Subsequently, $TB_{amp}$ (1 L) in a 4-L Erlenmeyer flask was inoculated with 50 mL of the preculture and allowed to shake for 2–2.5 hours at 37 ºC, and 180 rpm until an optical density at 600 nm ($OD_{600}$) of 0.8–1.0. The culture was then rested on ice for 30–45 min and induced with 0.5 mM IPTG and 1.0 mM ALA (final concentrations). Protein expression was conducted at 22 ºC and 180 rpm for 16–20 hours and collected by centrifugation at 4,000 r.p.m. at 4 °C. After cell lysis by sonication, and centrifugation (20,000 g, 25 min, and 4 ºC), the proteins were purified by Ni-affinity chromatography. The clarified lysate was transferred to a nickel nitriloacetic acid (Ni-NTA) column equilibrated with equilibration buffer (50 mM KPi buffer, 200 mM NaCl, pH 8.0). The resin was washed with 200 ml Ni-NTA wash buffer (50 mM KPi buffer, 200 mM NaCl, 20 mM imidazole, pH 8.0). The proteins were eluted with Ni-NTA elution buffer (50 mM KPi buffer, 200 mM NaCl, 200 mM imidazole, pH 8.0). After elution, the proteins were dialysed in 50 mM KPi buffer (pH 8.0) using 3 KDa dialysis memberane and concentrated in 3kDa centrifugal filters. The haem-loaded protein concentration was determined using hemochrome assay.

#### Hemochrome Assay for the Determination of Hemoprotein Concentration

*E. coli* cells after expression were pelleted by centrifugation (4,000 g, 8 min, and 4 ºC). $TB_{amp}$ medium was removed. Harvested cell pellets were resuspended to M9–N medium (pH 8.0) supplemented with D-glucose (25 mM final concentration) to $OD_{600}$ = 50. The same cell suspension was used for reaction set-up and determination of protein concentration. Protein concentration in the cell was determined using the hemochrome assay on cell lysate (*4*). Lysate was obtained by sonication using a Qsonica Q500 sonicator (6 minutes, 1 second on, 2 seconds off, 35% amplitude, on wet ice). The cell debris was removed by centrifugation (14,000 g, 10 minutes, 4 °C). To a cuvette, 500 µL of the lysate and 500 µL of solution I [0.2 M NaOH, 40% (v/v) pyridine, 0.5 mM $K_3Fe(CN)_6$] were added. The UV-Vis spectrum (380–620 nm) of the oxidized state Fe(III) was recorded immediately. Sodium dithionite (10 µL of 0.5 M solution in water) was added, and the UV-Vis spectrum of the reduced state Fe(II) was recorded immediately. The protein concentration was calculated using the extinction coefficient and dilution factor (2× dilution in volume): $\varepsilon [557_{reduced} - 540_{oxidized}] = 23.98\ mM^{-1}cm^{-1}$. The hemochrome assay detects total heme level, which is a good approximation of over-expressed heme enzyme.

#### Analytical Reaction Setup and Product Quantification

*Reaction set-up in 96-well plate*

*E. coli* cells after expression in 96-well plates were pelleted by centrifugation (4,000 g, 8 min, and 4 ºC). $TB_{amp}$ medium was removed. Harvested cell pellets were resuspended with 380 µL M9–N medium (pH 7.2). The resuspended cell in 96-well plates were introduced to an anaerobic chamber (coy chamber, oxygen level: <60 ppm). Alkyl azides **1** (20 µL, 0.1 M stock in ethanol) were added using a multi-channel pipette. The reactions were shaken at room temperature overnight at 700 rpm.

*Reaction set-up in 2-mL vials with adjusted $OD_{600}$ (whole-cell)*

*E. coli* cells after expression in 125-mL Erlenmeyer flasks were pelleted by centrifugation (4,000 g, 8 min, and 4 ºC). $TB_{amp}$ medium was removed. Harvested cell pellets were resuspended to M9–N medium (pH 7.2) to $OD_{600}$ = 30. Resuspended cells (380 µL) were aliquoted to 2-mL screw cap vials and introduced to an anaerobic chamber (coy chamber, oxygen level: <60 ppm). Alkyl azides **1** (20 µL, 0.1 M stock in ethanol) were added using a multi-channel pipette. The reactions were shaken at room temperature overnight at 700 rpm.

*Reaction set-up in 2-mL vials with adjusted haem loading (purified protein)*

An aliquot of purified ***Tm*Pgb-PIP-G7** was diluted with M9–N buffer (50 mM, pH 7.2) to a final volume of 380 µL in a 2-mL vial, giving a haem concentration of 20 µM. Solution of purified protein was introduced to an anaerobic chamber (coy chamber, oxygen level: <60 ppm). Alkyl azide **1** (20 µL of a 0.1 M stock solution in ethanol) was added in a single portion under anaerobic conditions. The vial was sealed with a screw-top cap and shaken at 700 rpm overnight at room temperature or 37 °C.

An aliquot of purified **GDM-NT-401** was diluted with M9–N buffer (50 mM, pH 7.2) to a final volume of 370 µL in a 2-mL vial, giving a haem concentration of 10 µM. Solution of purified protein was introduced to an anaerobic chamber (coy chamber, oxygen level: <60 ppm). $Na_2S_2O_4$ (10 µL of a 0.1 M stock solution in M9–N buffer, pH 7.2), alkyl azide **1** (20 µL of a 0.1 M stock solution in ethanol) was added sequentially under anaerobic conditions. The vial was sealed with a screw-top cap and shaken at 700 rpm overnight at room temperature.

*Reaction work-up on 96-well plate*

After the reaction was completed, the vials were removed from the coy chamber. Acetonitrile (780 µL) was used to quench the reaction. 1,2,3-Trimethoxybenzene (5 mM final concentration) was used as internal standard (20 µL, 0.1 M stock in acetonitrile). The deep-well plates were shaken at room temperature for 20 min at 700 rpm and centrifuged (5,000 × g, 20 min, 4 °C). A sample of the supernatant (0.2 mL) was transferred to a 96-well plate and sealed with a heat plate sealer for reverse-phase HPLC-MS analysis. Products were identified and quantified by analyzing the absolute ion count on the single ion channel or comparing to a calibration curve generated with chemically synthesized standard products and 1,2,3-trimethoxybenzene as internal standard. Enantiomeric excess (*ee*) of the reactions were determined by analyzing an internal-standard-free sample using polar ionic HPLC-MS.

*Reaction work-up for analytical reactions in 2-mL vials*

After the reaction was completed, the vials were removed from the coy chamber. Acetonitrile (780 µL) was used to quench the reaction. 1,2,3-Trimethoxybenzene (5 mM final concentration) was used as internal standard (20 µL, 0.1 M stock in acetonitrile). The mixture was shaken at room temperature for 20 min at 700 rpm, transferred to an Eppendorf tube and centrifuged (5,000 × g, 20 min, 4 °C). A sample of the supernatant (0.2 mL) was transferred to a vial equipped with a glass insert for reverse-phase HPLC-MS analysis. Products were identified and quantified by comparing to a calibration curve generated with chemically synthesized standard products and 1,2,3-trimethoxybenzene as internal standard. Enantiomeric excess (*ee*) of the reactions were determined by analyzing an internal-standard-free sample using polar ionic HPLC-MS.

#### Preparative-Scale Reaction Setup and Product Quantification

*Preparative reaction set-up*

An aliquot of purified ***Tm*Pgb-PIP-G7** was diluted with M9–N buffer (50 mM, pH 7.2) to a final haem concentration of 20 µM. Alternatively, *e. coli* expressing **GDM-NT-401** were resuspended to M9–N medium (pH 7.2) to $OD_{600}$ = 30. Solution of purified protein or resuspended cells (190 mL) were introduced to an anaerobic chamber (coy chamber, oxygen level: <60 ppm). Alkyl azide **1** (10 mL of a 0.1 M stock solution in ethanol) was added in a single portion under aerobic conditions. The flask was sealed with a screw-top lid and shaken at 220 rpm for 2 days at 37 °C.

*Preparative reaction work-up*

Upon completion of the reaction, acetonitrile (400 mL) was added to quench the reaction and precipitate the protein. The precipitant was removed by filtration through a pad of Celite. Excess acetonitrile was removed under reduced pressure. The remaining aqueous solution was acidified to pH 3–4 with 1 M HCl and washed with ethyl acetate (150 mL × 3). The aqueous phase was then basified to pH 10 with 2 M NaOH and extracted with ethyl acetate (150 mL × 3). The combined organic layers were dried over $Na_2SO_4$ and concentrated under reduced pressure. The resulting residue was characterized by $^1H$ NMR spectroscopy and was found to be of sufficient purity without further purification.

#### General Procedures for the Preparation of Starting Material

##### General procedure A for the synthesis of alkyl azide substrates

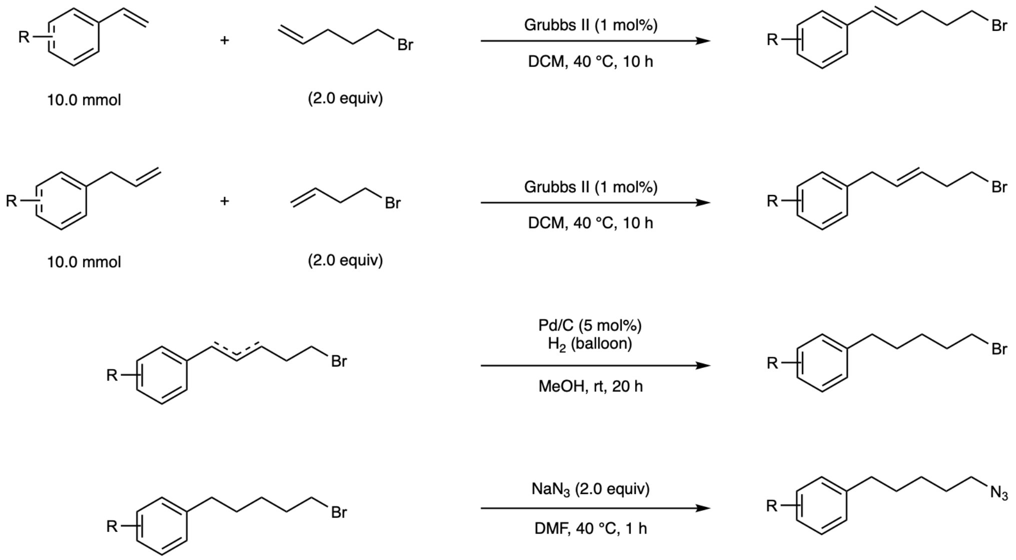

**Scheme S1**. General procedure A for the synthesis of alkyl azides through metathesis, hydrogenation, and azidation.

*Olefin metathesis*

Following a procedure slightly modified from literature (*5*), to a solution of styrene (10.0 mmol) and 5-bromo-1-pentene (20.0 mmol) in DCM (0.3 M) was added Grubbs II (1 mol%). The mixture was stirred at 40 °C for 10 h or until full conversion of the styrene monitored by thin-layer chromatography (TLC). Upon completion, the organic solvent was removed under reduced pressure, and the residue was subjected to next step without further purification. Alternatively, the same procedure could be applied for the combination of allylbenzene (10.0 mmol) and 4-bromo-1-butene (20 mmol).

*Hydrogenation*

To a solution of alkenyl bromide in MeOH (0.5 M) was added palladium on carbon (5 mol%). The reaction mixture was equipped with a hydrogen balloon and stirred for 20 h at room temperature or until full conversion of the styrene monitored by thin-layer chromatography (TLC). Upon completion, the reaction mixture was filtered, and the organic solvent was removed under reduced pressure. The residue was subjected to next step without further purification.

*Azidation*

To a solution of alkyl bromide in DMF (0.5 M) was added $NaN_3$ (2.0 equiv). The reaction mixture was stirred at 40 °C for 10 h or until full conversion of the starting material monitored by thin-layer chromatography (TLC). Upon completion, the reaction mixture was quenched with brine. filtered and the organic solvent was removed under reduced pressure. After this time, the reaction was quenched with water. The aqueous solution was extracted with ethyl acetate (50 mL ×3). The combined organic layers were dried over $Na_2SO_4$. The organic solvent was removed under reduced pressure, and the residue was subjected to flash column chromatography on silica gel with hexane as eluent to afford the 1-(5-azidopentyl)-benzene product.

**General procedure B for the synthesis of 1m**

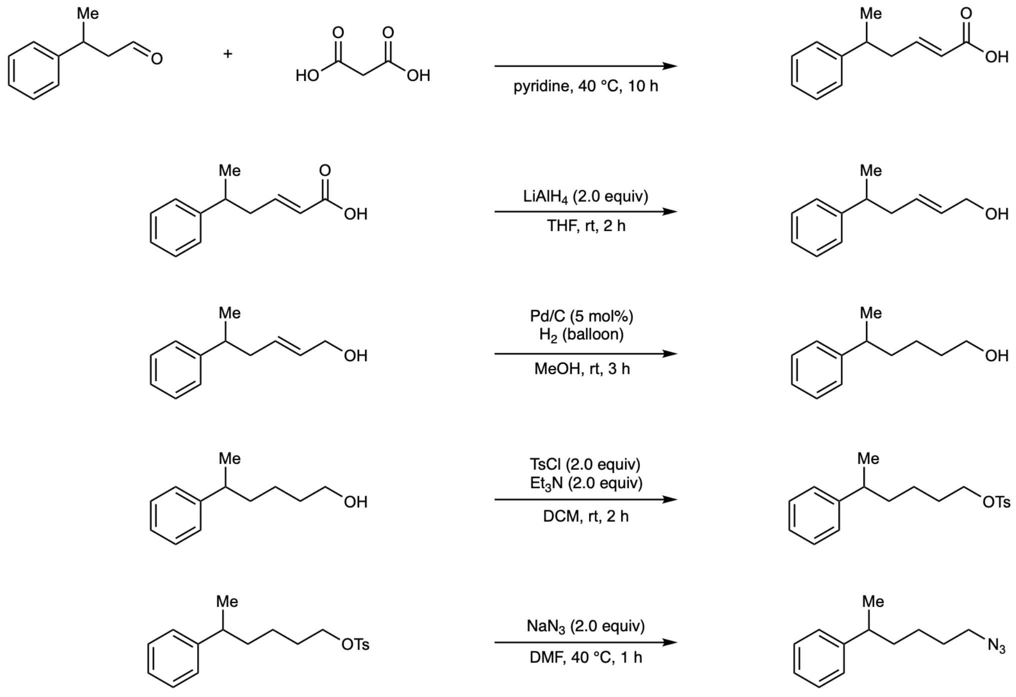

**Scheme S2**. General procedure B for the synthesis of substrate **1m** through Knoevenagel condensation, reduction, hydrogenation, tosylation, and azidation.

*Knoevenagel condensation Doebner modification*

The reaction was performed following a modified procedure from literature. A round-bottom flask containing a mixture of 3-phenylbutanal (12.0 mmol) and malonic acid (10.0 mmol) in pyridine (0.5 M) was equipped with a balloon. The mixture was stirred at 40 °C for 10 h or until full conversion of the substrate monitored by thin-layer chromatography (TLC). Upon completion, the organic solvent was removed under reduced pressure, and the residue was subjected to next step without further purification.

*Reduction of carboxylic acid*

To a solution of the α,β-unsaturated acid in THF, $LiAlH_4$ (2.0 equiv) was added at 0 °C. The reaction was gradually warmed to room temperature and stirred for 2 h or until full conversion of the substrate monitored by thin-layer chromatography (TLC). Upon completion, the reaction was quenched with cold saturated $NH_4Cl$ solution. The aqueous solution was extracted with ethyl acetate (200 mL ×3). The combined organic layers were dried over $Na_2SO_4$. The organic solvent was removed under reduced pressure, and the residue was subjected to next step without further purification.

*Hydrogenation*

To a solution of alkenyl alcohol in MeOH (0.5 M) was added palladium on carbon (5 mol%). The reaction mixture was equipped with a hydrogen balloon and stirred for 20 h at room temperature or until full conversion of the styrene monitored by thin-layer chromatography (TLC). Upon completion, the reaction mixture was filtered, and the organic solvent was removed under reduced pressure. The residue was subjected to next step without further purification.

*Tosylation*

To a solution of alkyl alcohol in DCM (0.5 M) was added TsCl (2.0 equiv) and triethylamine (2.0 equiv). The reaction mixture was stirred at room temperature for 2 h or until full conversion of the styrene monitored by thin-layer chromatography (TLC). Upon completion, the reaction mixture was quenched with brine. The aqueous solution was extracted with DCM (200 mL ×3). The combined organic layers were dried over $Na_2SO_4$. The organic solvent was removed under reduced pressure, and the residue was subjected to next step without further purification.

*Azidation*

To a solution of tosyl protected alkyl alcohol in DMF (0.5 M) was added $NaN_3$ (2.0 equiv). The reaction mixture was stirred at 40 °C for 10 h or until full conversion of the starting material monitored by thin-layer chromatography (TLC). Upon completion, the reaction mixture was quenched with brine. filtered and the organic solvent was removed under reduced pressure. After this time, the reaction was quenched with water. The aqueous solution was extracted with ethyl acetate (50 mL ×3). The combined organic layers were dried over $Na_2SO_4$. The organic solvent was removed under reduced pressure, and the residue was subjected to flash column chromatography on silica gel with hexane as eluent to afford the 1-(5-azidopentyl)-benzene product.

**Selected Small-molecule Catalysts for Intramolecular C–H Amination**

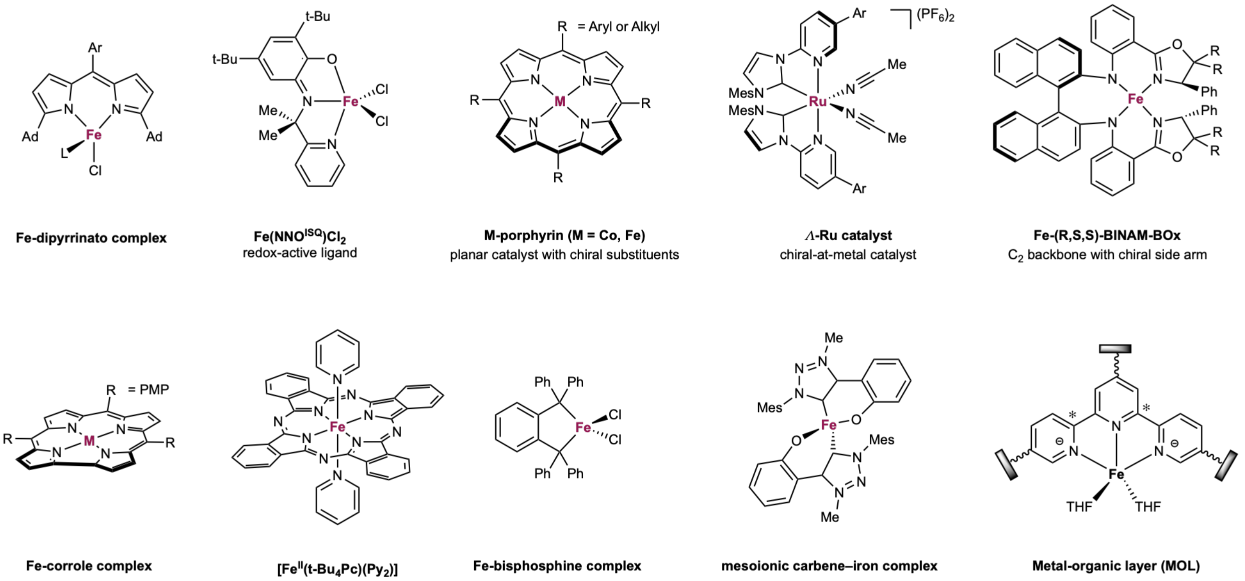

**Scheme S3**. Selected examples of small-molecule catalysts for Intramolecular C–H Amination of Alkyl azides.

**DNA Sequences and Protein Sequences**

**Part 1: *Tm*Pgb-PIP** variants

*DNA sequence of* ***Tm*Pgb-wt**:

ATGCCGCAAATTCCCGGTTACACTTATGGAGATCCCGCCTTACCTCCCAGTCCCGTTTCTTTA
GAGGAATTGGAGCGTCTGAAGGCTTGCTTGCTGTGGACGGAGGAGGATGATAAAGCATTAG
AGCAGGCCGGTAAAGTATTAGAGGACCAAGTAGAAGAAGTGTTAGATTTATGGTACGGCTT
TGTCGGGAGCCATCCTCATTTACTTCACTATTTCACTGATCCTCAGGGGAACCCGATCCCCGA
CTATCTTGAGCGTGTTCGCCGCCGTTTCGGACAGTGGATTCTGGATACCTGCTTTCGTCCCAA
AGACGAAACCTGGCTTCGTTATCAGCATGAGATTGGCTTACGTCATCATCGCACAAAAAAAA
ACCAAACTGACGGCGTGACCTCCGTCCCACATATTCCGTTGCGCTATTTGATCAGTTTTATTT
ATCCCATTACAGCCACCATCAAACCCTTCCTGACTAAGAAGGGCCACAATCCCGAGGAGGTG
GAGCGTATGTATCAGGCATGGTTCAAGGCAGTTGTATTGCAAGTAGCACTTTGGTCCTATCC
TTACACTCAGCCTGGCGACTTTCTCGAG

*Amino acid sequence of* ***Tm*Pgb-wt**:

MPQIPGYTYGDPALPPSPVSLEELERLKACLLWTEEDDKALEQAGKVLEDQVEEVLDLWYGFVG
SHPHLLHYFTDPQGNPIPDYLERVRRRFGQWILDTCFRPKDETWLRYQHEIGLRHHRTKKNQTD
GVTSVPHIPLRYLISFIYPITATIKPFLTKKGHNPEEVERMYQAWFKAVVLQVALWSYPYTQPGDF
LE

*DNA sequence of* ***Tm*Pgb-PIP-G1-5442**:

ATGCCGCAAATTCCCGGTTACACTTATGGAGATCCCGCCTTACCTCCCAGTCCCGTTTCTTTA
GAGGAATTGGAGCGTCTGAAGGCTTGCTTGCTGTGGACGGAGGAGGATGATAAAGCATTAG
AGCAGGCCGGTAAAGTATTAGAGGACCAAGTAGAAGAAGTGTTAGATTTATGGTATGGCTT
TGTCGGGAGCCATCCTCATTTACTTCACTATTTCACTGATCCTCAGGGGAACCCGATCCCCGA
CTATCTTGAGCGTGTTCGCCGCCGTTTCGGACAGTGGATTCTGGATACCTGCTTTCGTCCCAA
AGACGAAACCTGGCTTCGTTATCAGCATGAGATTGGCTTACGTCATCATCGCACAAAAAAAA
ACCAAACTGACGGCGTGACCTCCGTCCCACATATTCCGTTGCGCTATTTGATCAGTTTTATTT
ATCCCACTACAGCCACCATCAAACCCTTCCTGACTAAGAAGGGCCACAATCCCGAGGAGGT
GGAGCGTATGTATCAGGCATGGTTCAAGGCAGTTGTATTGCAAGTAGCACTTTGGTCCTATC
CTTACACTCAGCCTGGCGACTTTCTCGAG

*Amino acid sequence of* ***Tm*Pgb-PIP-G1-5442**:

MPQIPGYTYGDPALPPSPVSLEELERLKACLLWTEEDDKALEQAGKVLEDQVEEVLDLWYGFVG
SHPHLLHYFTDPQGNPIPDYLERVRRRFGQWILDTCFRPKDETWLRYQHEIGLRHHRTKKNQTD
GVTSVPHIPLRYLISFIYPTTATIKPFLTKKGHNPEEVERMYQAWFKAVVLQVALWSYPYTQPGD
FLE

*DNA sequence of* ***Tm*Pgb-PIP-G2-5443**:

ATGCCGCAAATTCCCGGTTACACTTATGGAGATCCCGCCTTACCTCCCGGTCCCGTTTCTTTA
GAGGAATTGGAGCGTCTGAAGGCTTGCTTGCTGTGGACGGAGGAGGATGATAAAGCATTAG
AGCAGGCCGGTAAAGTATTAGAGGACCAAGTAGAAGAAGTGTTAGATTTATGGTATGGCTT
TGTCGGGAGCCATCCTCATCTACTTCACTATTTCACTGATCCTCAGGGGAACCCGATCCCCGA
CTATCTTGAGCGTGTTCGCCGCCGTTTCGGACAGTGGATTCTGGATACCTGCTTTCGTCCCAA
AGACGAAACCTGGCTTCGTTATCAGCATGAGATTGGCTTACGTCATCATCGCACAAAAAAAA
ACCAAACTGACGGCGTGACCTCCGTCCCACATATTCCGTTGCGCTATTCGATCAGTTTTATTT
ATCCCACTACAGCCACCATCAAACCCTTCCTGACTAAGAAGGGCCACAATCCCGAGGAGGT
GGAGCGTATGCATCAGGCATGGTTCAAGGCAGTTGTATTGCAAGTAGCACTTTGGTCCTATC
CTTACACTCAGCCTGGCGACTTTCTCGAG

*Amino acid sequence of* ***Tm*****Pgb-PIP-G2-5443**:

MPQIPGYTYGDPALPPGPVSLEELERLKACLLWTEEDDKALEQAGKVLEDQVEEVLDLWYGFV
GSHPHLLHYFTDPQGNPIPDYLERVRRRFGQWILDTCFRPKDETWLRYQHEIGLRHHRTKKNQT
DGVTSVPHIPLRYSISFIYPTTATIKPFLTKKGHNPEEVERMHQAWFKAVVLQVALWSYPYTQPG
DFLE

*DNA sequence of* ***Tm*****Pgb-PIP-G3-5444**:

ATGCCGCAAATTCCCGGTTACACTTATGGAGATCCCGCCTTACCTCCCGGTCCCGTTTCTTTA
GAGGAATTGGAGCGTCTGAAGGCTAGCTTGCTGTGGACGGAGGAGGATGATAAAGCATTAG
AGCAGGCCGGTAAAGTCTTAGAGGACCAAGTAGAAGAAGTGTTAGATTTATGGTATGGCTTT
GTCGGGAGCCATCCTCATCTACTTCACTATTTCACTGATCCTCAGGGGAACCCGATCCCCGA
CTATCTTGAGCGTGTTCGCCGCCGTTTCGGACAGTGGATTCTGGATACCTGCTTTCGTCCCAA
AGACGAAACCTGGCTTCGTTATCAGCATGAGATTGGCTTACGTCATCATCGCACAAAAAAAA
ACCAAACTGACGGTGTGACCTCCGTCCCACATATTCCGTTGCGCTATTCGATCAGTTTTATTT
ATCCCACTACAGCCACCATCAAACCCTTCCTGACTAAGAAGGGCCACAATCCCGAGGAGGT
GGAGCGTATGCATCAGGCATGGTTCAAGGCAGTTGTATTGCAAGTAGCACTTTGGTCCTATC
CTTACACTCAGCCTGGCGACTTTCTCGAG

*Amino acid sequence of* ***Tm*****Pgb-PIP-G3-5444**:

MPQIPGYTYGDPALPPGPVSLEELERLKASLLWTEEDDKALEQAGKVLEDQVEEVLDLWYGFV
GSHPHLLHYFTDPQGNPIPDYLERVRRRFGQWILDTCFRPKDETWLRYQHEIGLRHHRTKKNQT
DGVTSVPHIPLRYSISFIYPTTATIKPFLTKKGHNPEEVERMHQAWFKAVVLQVALWSYPYTQPG
DFLE

*DNA sequence of* ***Tm*****Pgb-PIP-G4-5445**:

ATGCCGCAAATTCCCGGTTACACTTATGGAGATCCCGCCTTACCTCCCGGTCCCGTTTCTTTA
GAGGAATTGGAGCGTCTGAAGGCTAGCTTGCTGTGGACGGAGGAGGATGATAAAGCATTAG
AGCAGGCCGGTAAAGTCTTAGAGGACCAAGTAGAAGAAGTGTTAGATTTATGGTATGGCTG
GGTCGGGAGCCATCCTCATCTACTTCACTATTTCACTGATCCTCAGGGGAACCCGATCCCCG
ACTATCTTGAGCGTGTTCGCCGCCGTTTCGGACAGTGGATTCTGGATACCTGCTTTCGTCCCA
AAGACGAAACCTGGCTTCGTTATCAGCATGAGATTGGCTTACGTCATCATCGCACAAAAAAA
AACCAAACTGACGGTGTGACCTCCGTCCCACATATTCCGTTGCGCTATTCGATCAGTTTTATT
TATCCCACTACAGCCACCATCAAACCCTTCCTGACTAAGAAGGGCCACAATCCCGAGGAGGT
GGAGCGTATGCATCAGGCATGGTTCAAGGCAGTTGTATTGCAAGTAGCACTTTGGTCCTATC
CTTACACTCAGCCTGGCGACTTTCTCGAG

*Amino acid sequence of* ***Tm*****Pgb-PIP-G4-5445**:

MPQIPGYTYGDPALPPGPVSLEELERLKASLLWTEEDDKALEQAGKVLEDQVEEVLDLWYGWV
GSHPHLLHYFTDPQGNPIPDYLERVRRRFGQWILDTCFRPKDETWLRYQHEIGLRHHRTKKNQT
DGVTSVPHIPLRYSISFIYPTTATIKPFLTKKGHNPEEVERMHQAWFKAVVLQVALWSYPYTQPG
DFLE

*DNA sequence of* ***Tm*****Pgb-PIP-G5-5446**:

ATGCCGCAAATTCCCGGTTACACTTATGGAGATCCCGCCTTACCTCCCGGTCCCGTTTCTTTA
GAGGAATTGGAGCGTCTGAAGGCTAGCTTGCTGTGGACGGAGGAGGATGATAAAGCATTAG
AGCAGGCCGGTAAAGTCTTAGAGGACCAAGTAGAAGAAGTGTTAGATTTATGGTATGGCTG
GGTCGGGAGCCATCCTCATCTACTTCACTATTTCACTGATCCTCAGGGGAACCCGATCCCCG
ACTATCTTGAGCGTGTTCGCCGCCGTTTCGGACAGTGGATTCTGGATACCTGCTTTCGTCCCA
AAGACGAAACCTGGCTTCGTTATCAGCATGAGATTGGCTTACGTCATCATCGCACAAAAAAA
AACCAAACTGACGGTGTGACCTCCGTCCCACATATTCCGTTGCGCTATAATATCAGTTTTATT
TATCCCACTACAGCCACCATCAAACCCTTCCTGACTAAGAAGGGCCACAATCCCGAGGAGGT

GGAGCGTATGCATCAGGCATGGTTCAAGGCAGTTGTATTGCAAGTAGCACTTTGGTCCTATC
CTTACACTCAGCCTGGCGACTTTCTCGAG

*Amino acid sequence of* ***Tm*****Pgb-PIP-G5-5446**:

MPQIPGYTYGDPALPPGPVSLEELERLKASLLWTEEDDKALEQAGKVLEDQVEEVLDLWYGWV
GSHPHLLHYFTDPQGNPIPDYLERVRRRFGQWILDTCFRPKDETWLRYQHEIGLRHHRTKKNQT
DGVTSVPHIPLRYNISFIYPTTATIKPFLTKKGHNPEEVERMHQAWFKAVVLQVALWSYPYTQPG
DFLE

*DNA sequence of* ***Tm*****Pgb-PIP-G6-5447**:

ATGCCGCAAATTCCCGGTTACACTTATGGAGATCCCGCCTTACCTCCCGGTCCCGTTTCTTTA
GAGGAATTGGAGCGTCTGAAGGCTAGCTTGCTGTGGACGGAGGAGGATGATAAAGCATTAG
AGCAGGCCGGTAAAGTCTTAGAGGACCAAGTAGAAGAAGTGTTAGATTTATGGTATGGCTG
GGTCGGGAGCGAACCTCATCTACTTCACTATTTCACTGATCCTCAGGGGAACCCGATCCCCG
ACTATCTTGAGCGTGTTCGCCGCCGTTTCGGACAGTGGATTCTGGATACCTGCTTTCGTCCCA
AAGACGAAACCTGGCTTCGTTATCAGCATGAGATTGGCTTACGTCATCATCGCACAAAAAAA
AACCAAACTGACGGTGTGACCTCCGTCCCACATATTCCGTTGCGCTATAATATCAGTTTTATT
TATCCCACTACAGCCACCATCAAACCCTTCCTGACTAAGAAGGGCCACAATCCCGAGGAGGT
GGAGCGTATGCATCAGGCATGGTTCAAGGCAGTTGTATTGCAAGTAGCACTTTGGTCCTATC
CTTACACTCAGCCTGGCGACTTTCTCGAG

*Amino acid sequence of* ***Tm*****Pgb-PIP-G6-5447**:

MPQIPGYTYGDPALPPGPVSLEELERLKASLLWTEEDDKALEQAGKVLEDQVEEVLDLWYGWV
GSEPHLLHYFTDPQGNPIPDYLERVRRRFGQWILDTCFRPKDETWLRYQHEIGLRHHRTKKNQT
DGVTSVPHIPLRYNISFIYPTTATIKPFLTKKGHNPEEVERMHQAWFKAVVLQVALWSYPYTQPG
DFLE

*DNA sequence of* ***Tm*****Pgb-PIP-G7-5448**:

ATGCCGCAAATTCCCGGTTACACTTATGGAGATCCCGCCTTACCTCCCGGTCCCGTTTCTTTA
GAGGAATTGGAGCGTCTGAAGGCCAGCTTGCTGTGGACGGAGGAGGATGATAAAGCATTAG
AGCAGGCCGGTAAAGTCTTAGAGGACCAAGTAGAAGAAGTGTTAGATTTATGGTATGGCTG
GGTCGGGGGTGAACCTCATCTACTTCACTATTTCACTGATCCTCAGGCGAACCCGATCCCCG
ACTATCTTGAGCGTGTTCGCCGCCGTTTCGGACAGTGGATTCTGGATACCTGCTTTCGTCCCA
AAGACGAAACCTGGCTTCGTTATCAGCATGAGATTGGCTTACGTCATCATCGCACAAAAAAA
AACCAAACTGACGGTGTGACCTCCGTCCCACATATTCCGTTGCGCTATAATATCAGTTTTATT
TATCCCACTACAGCCACCATCAAACCCTTCCTGACTAAGAAGGGCCACAATCCCGAGGAGGT
GGAGCGTATGCATCAGGCATGGTTCAAGGCAGTTGTATTGAGTGTAGCACTTTGGTCCTATC
CTTACACTCAGCCTAGCGACTTTCTCGAG

*Amino acid sequence of* ***Tm*****Pgb-PIP-G7-5448**:

MPQIPGYTYGDPALPPGPVSLEELERLKASLLWTEEDDKALEQAGKVLEDQVEEVLDLWYGWV
GGEPHLLHYFTDPQANPIPDYLERVRRRFGQWILDTCFRPKDETWLRYQHEIGLRHHRTKKNQT
DGVTSVPHIPLRYNISFIYPTTATIKPFLTKKGHNPEEVERMHQAWFKAVVLSVALWSYPYTQPS
DFLE

**Part 2: GDM-NT** variants from recycled scaffold design

*DNA acid sequence of* **GDM-NT-121-5449**

CCTCCGCTTGTGGAGCCAATTGAAGATCCCGCAGAACGTGCAGCAATCTTGGCGGAATTGAA
AGCTTTCTTGCGGTTCCTGGCTGCTCTGAGTCGTGAGGAACTGGAACAGGTCACGAAGCGGC

GCGCTGGGGGTACATTGGATGAACTTCACGCACTTCTCCGTGAATTGGCAGCCGCCGACTTG
GAACTCCTTCTTGCTTTACGGGTGAGAGCAACCCGTGCCGATATAAACGCGTTTGCTAATGC
TTCGGACGAAGAGTTAGCGAAACTTGAAGCAATCCTGGCGGAAGAAATTGACAGATTAGTC
GCTCGTATGGAAGAAGCCCTTCCAGGCCTGGATTATGACGCAATTAGAGCCGAAGTCCTGGC
GGCTGTGAAGGAAGCGCTGGCTGCGCTGAAAGAAGGTGGTCTGGATGCATTACTGGATTAC
GTTTATCGAAGACTGGGAGAATTAGGGGCCAGAGTTTTGAGCTTGGATGTGAAAAAACTGA
GAGAACTGCATCCGGATGCGCCGCTGGGTCCATTAGTGGCCGCTGCCAAGGGCATTACGATA
GTCGAGATGGAAATTCCCGAGGCACTTCTGCCACTTCTCGTACGTGAGTTTCGAGAAGGGTT
GCCGGCACTTGACCGTCGCTATGCAGATGAATTAACTCGCTCCCAGCTGGCTACTTTAGTGG
CACTCTATGCCTTTAGCCTGATTCTTATGGCCTTACTTGCGGCGAAACAAAAACCCGATGCCC
CCAAAGTGCGCGTGTTCATCGTGCACCTTACCTATGATGACGGGACACTCACTCTGTTGCTC
ACTCCGGCA

*Amino acid sequence of* **GDM-NT-121-5449:**

PPLVEPIEDPAERAAILAELKAFLRFLAALSREELEQVTKRRAGGTLDELHALLRELAAADLELLL
ALRVRATRADINAFANASDEELAKLEAILAEEIDRLVARMEEALPGLDYDAIRAEVLAAVKEAL
AALKEGGLDALLDYVYRRLGELGARVLSLDVKKLRELHPDAPLGPLVAAAKGITIVEMEIPEAL
LPLLVREFREGLPALDRRYADELTRSQLATLVALYAFSLILMALLAAKQKPDAPKVRVFIVHLTY
DDGTLTLLLTPA

*DNA sequence of* **GDM-NT-226-5450:**

CCACCTGTCCTGCGCCCAGTTACTGACCCGGCCCGTCGCGCCGAGTTGCACGCCGCGCTGCT
TGAATTACTGCGTCAGTTGGCCGAACTCAGTGATGAACAACTTGCGGCGTGCTTAAAACGCC
GTTACGGGCTGACACAAGATGAGGTTAAAGAATTGTTAAAAGAACTGGCAGCAGCGGATCT
TCGCCTGCTGAACGCACTTGAAGTACATGCTACGAGAGCGGATATCGATGCTGCCCGCGATT
GGTCGGATGAAGAGTGGGCACGCCTGCGGGCGGTTTTTCGCGCGTTCCTGCGCGAACTGGTC
GCCCGGCTTGCTGCCACAGTTCCGGGTTTAGACGGAGAAGCGCTCTTGGCACGTCTGGAAGC
GGCGCTGGATGAGGCATTAGCCGCTCTGCGGGCAGGAGGTTTTGATGCTCTGCTGGATACAG
TAGCGCGTCAGCTCGAAGAGGCAGGTATTCCCTTGGTGGTATTTGATCTCCGCCGACTTGCC
GCTTTGCCAGGAGTCCCACCCCTTCCGGCTGTTCTGGCCGGCGCTCGAGGCCTGGCGATCGC
TGTTCTGGATGTCCCCCCTGCGGTGGCAGATTTATTCGCAGAGGTTTTATCCCGCGACATTCC
AGCATTCCAAGCACGTGTAGCCCCTGAACTTACCGCTTCTCAGATGGCTGTCTTGGCGGCAC
TTGCAACTGCCGCGTTGCTGCTTGCTGCTGCGTTCGTACGTGAAGCGCGACCGGATGCGCCC
ACAGTTCGCTTGGTCGTCGCGCACTTGACCTTTCCGGATGGCAACCTGGTTTTAGAGTTTCGT
GTAGAG

*Pairwise identity* (compared to **GDM-NT-121-5450**): 45.41%

*Amino acid sequence of* **GDM-NT-226-5450:**

PPVLRPVTDPARRAELHAALLELLRQLAELSDEQLAACLKRRYGLTQDEVKELLKELAAADLRL
LNALEVHATRADIDAARDWSDEEWARLRAVFRAFLRELVARLAATVPGLDGEALLARLEAALD
EALAALRAGGFDALLDTVARQLEEAGIPLVVFDLRRLAALPGVPPLPAVLAGARGLAIAVLDVPP
AVADLFAEVLSRDIPAFQARVAPELTASQMAVLAALATAALLLAAAFVREARPDAPTVRLVVA
HLTFPDGNLVLEFRVE

*DNA acid sequence of* **GDM-NT-151-5451**

AAGACTATAACACTGGAATCGGAATCCAGCCTGCCTTTTTCAGTACTCGAGTTGCTGTTGTC
ACCTGAGAATCTGGCCCGTCTTCTGGCGACATGGAAAAGTGAACACATAAGAATAGAAAGT
ATCGAAATACTGGAGTTTGTACCTGGTGAGCGCGTCACGGTCAAAGTTAAAGTGACTTTTAA
AGTGGCAGGTGAAGAAGTTCATCTTGAAAGCGTGGTAACGTTCGAACGCGTCGAGGGCGAT
CGTGCTCGTATCACCGCCGTCATTACCTACGATGAAGACAAGTTTCGTGAGGCACTCGAAAA

ACTTAGCGATGAAGAAATCAAAGACTGGATGTTAGCCGTTGCGAAGGAAGTTGCGGCAGCT
GTCAGAGAGTTGCTTGAGGAATTACGCGAACGAGGAGAAATTGATCTTAGTGACGAAGAAA
TCGATCGCCTCGTCGTTGCGGTCGGTCTGAAACATATGGGTGTGTTAGCTAAGTTAATCGAA
GCGGTTACCGGGAAAGACTACTCTGAACTGCTGGAGGAAGTTAAAAAACGTTTCCTGGAGG
AATGGTTG

*Amino acid sequence of* **GDM-NT-151-5451:**

KTITLESESSLPFSVLELLLSPENLARLLATWKSEHIRIESIEILEFVPGERVTVKVKVTFKVAGEEV
HLESVVTFERVEGDRARITAVITYDEDKFREALEKLSDEEIKDWMLAVAKEVAAAVRELLEELR
ERGEIDLSDEEIDRLVVAVGLKHMGVLAKLIEAVTGKDYSELLEEVKKRFLEEWL

*DNA sequence of* **GDM-NT-362-5452:**

GAAACGCTGGAGTTGGAAAGTGAAGCATCACTGCCCCTCGACGTGCTGGAGTTACTGTTATC
CCCTGAGAATCTTCCTCGTGTGTTGGCAGTGGGCCGTGGAGAACATTTCCGTCTGCTCAGTCT
TGAGGTTCTTGCGAGAGAACCGGGCCGTCGTGTAACGGTCGCTTCTCGCGCTGAAATAAGAC
TGGGTGATCGCTGGATCCCGGTAGATGTCGTGCTGACATTTGAGCGACAGCCGGATGGCAGC
GTGCGCATTAGAGCCGTACTGACATTCGACGCCGACGCCGTGCGAGAGGCATTGGCGGACA
TGACGGATGAAGAAATTGAGGATCTTTTGTTAGGTTTTGCCTTAGAGGTTGCTGCAGCGGTG
AGAGCACTTCTGGAAGAATTGCGCGAAGCTGGACGTATAGATCTGTCTGACGAAGAAATTG
CCGCGCTGGCACTGCGAGTGGGTTTAAAACATATGCGTGTCCTGGCACGGATTTTAGAGAAA
GCGACTGGGCGTGACTTATCTGCGGTATTAGAACGCGCGGAGGAACGCCTCAAAAAAGAAC
TCGCG

*Pairwise identity* (compared to **GDM-NT-151-5451**): 50.6%

*Amino acid sequence of* **GDM-NT-362-5452:**

ETLELESEASLPLDVLELLLSPENLPRVLAVGRGEHFRLLSLEVLAREPGRRVTVASRAEIRLGDR
WIPVDVVLTFERQPDGSVRIRAVLTFDADAVREALADMTDEEIEDLLLGFALEVAAAVRALLEE
LREAGRIDLSDEEIAALALRVGLKHMRVLARILEKATGRDLSAVLERAEERLKKELA

**Part 3: GDM-NT** variants from directed evolution and additional protein engineering thereof.

*DNA acid sequence of* **GDM-NT-399-5453**

GAAACGCTGGAGTTGGAAAGTGAAGCATCACTGCCCCTCGACGTGCTGGAGTTACTGTTATC
CCCTGAGAATCTTCCTCGTGTGTTGGCAGTGGGCCGTGGAGAACATTTCCGTCTGCTCAGTCT
TGAGGTTCTTGCGAGAGAACCGGGCCGTCGTCTAACGGTCGCTTCTCGCGCTGAAATAAGAC
TGGGTGACCGCTGGGTCCCGGTAGATGTCGTGCTGACATTTGAGCGACAGCCGGATGGCAGC
GTGCGCATTAGAGCCGTACTGACATTCGACGCCGACGCCGTGCGAGGGGCATTGGCGGACA
TGACGGATGAAGAAATTGAGGATCTTTTGTTAGGTTTTGCCTTAGAGGTTGCTGCAGCGGTG
AGAGCACTTCTGGAAGAATTGCGCGAAGCTGGACGTATAGATCTGTCTGACGAAGAAATTG
CCGCGCTGGCACTGCGAGTGGGTTTAAAACTTATGCGTGTCCTGGCACGGATTTTGGAGAAA
GCGACTGGGCGTGACTTATCTGCGGTATTAGAACGCGCGGAGGAACGCCTCAAAAAAGAAC
TCGCG

*Amino acid sequence of* **GDM-NT-399-5453:**

ETLELESEASLPLDVLELLLSPENLPRVLAVGRGEHFRLLSLEVLAREPGRRLTVASRAEIRLGDR
WVPVDVVLTFERQPDGSVRIRAVLTFDADAVRGALADMTDEEIEDLLLGFALEVAAAVRALLEE
LREAGRIDLSDEEIAALALRVGLKLMRVLARILEKATGRDLSAVLERAEERLKKELA

*DNA acid sequence of* **GDM-NT-400-5454:**

GAAACGCTGGAGTTGGAAAGTGAAGCATCACTGCCCCTCGACGTGCTGGAGTTACTGTTATC
CCCTGAGAATCTTCCTCGTGTGTTGGCAGTGGGCCGTGGAGAACATTTCCGTCTGCTCAGTCT
TGAGGTTCTTGCGAGAGAACCGGGCCGTCGTGTAACGGTCGCTTCTCGCGCTGAAATAAGAC
TGGGTGATCGCTGGATCCCGGTAGATGTCGTGCTGACATTTGAGCGACAGCCGGATGGCAGC
GTGCGCATTAGAGCCGTACTGACATTCGACGCCGACGCCGTGCGAGAGGCATTGGCGGACA
TGACGGATGAAGAAATTGAGGATCTTTTGTTAGGTTTTGCCTTAGAGGTTGCTGCAGCGGTG
AGAGCACTTCTGGAAGAATTGCGCGAAGCTGGACGTATAGATCTGTCTGACGAAGAAATTG
CCGCGCTGGCACTGCGAGTGGGTTTAAAAATGTTGCGTGTCCTGGCACGGATTTTAGAGAAA
GCGACTGGGCGTGACTTATCTGCGGTATTAGAACGCGCGGAGGAACGCCTCAAAAAAGAAC
TCGCG

*Amino acid sequence of* **GDM-NT-400-5455:**

ETLELESEASLPLDVLELLLSPENLPRVLAVGRGEHFRLLSLEVLAREPGRRVTVASRAEIRLGDR
WIPVDVVLTFERQPDGSVRIRAVLTFDADAVREALADMTDEEIEDLLLGFALEVAAAVRALLEE
LREAGRIDLSDEEIAALALRVGLKMLRVLARILEKATGRDLSAVLERAEERLKKELA

*DNA sequence of* **GDM-NT-401-5455:**

GAAACGCTGGAGTTGGAAAGTGAAGCATCACTGCCCCTCGACGTGCTGGAGTTACTGTTATC
CCCTGAGAATCTTCCTCGTGTGTTGGCAGTGGGCCGTGGAGAACATTTCCGTCTGCTCAGTCT
TGAGGTTCTTGCGAGAGAACCGGGCCGTCGTCTTACGGTCGCTTCTCGCGCTGAAATAAGAC
TGGGTGATCGCTGGGTTCCGGTAGATGTCGTGCTGACATTTGAGCGACAGCCGGATGGCAGC
GTGCGCATTAGAGCCGTACTGACATTCGACGCCGACGCCGTGCGAGGTGCATTGGCGGACAT
GACGGATGAAGAAATTGAGGATCTTTTGTTAGGTTTTGCCTTAGAGGTTGCTGCAGCGGTGA
GAGCACTTCTGGAAGAATTGCGCGAAGCTGGACGTATAGATCTGTCTGACGAAGAAATTGC
CGCGCTGGCACTGCGAGTGGGTTTAAAAATGTTGCGTGTCCTGGCACGGATTTTAGAGAAAG
CGACTGGGCGTGACTTATCTGCGGTATTAGAACGCGCGGAGGAACGCCTCAAAAAAGAACT
CGCG

*Amino acid sequence of* **GDM-NT-401-5455:**

ETLELESEASLPLDVLELLLSPENLPRVLAVGRGEHFRLLSLEVLAREPGRRLTVASRAEIRLGDR
WVPVDVVLTFERQPDGSVRIRAVLTFDADAVRGALADMTDEEIEDLLLGFALEVAAAVRALLEE
LREAGRIDLSDEEIAALALRVGLKMLRVLARILEKATGRDLSAVLERAEERLKKELA

*DNA acid sequence of* **GDM-NT-402-5456:**

GAAACGCTGGAGTTGGAAAGTGAAGCATCACTGCCCCTCGACGTGCTGGAGTTACTGTTATC
CCCTGAGAATCTTCCTCGTGTGTTGGCAGTGGGCCGTGGAGAACATTTCCGTCTGCTCAGTCT
TGAGGTTCTTGCGAGAGAACCGGGCCGTCGTCTTACGGTCGCTTCTCGCGCTGAAATAAGAC
TGGGTGATCGCTGGGTTCCGGTAGATGTCGTGCTGACATTTGAGCGACAGCCGGATGGCAGC
GTGCGCATTAGAGCCGTACTGACATTCGACGCCGACGCCGTGCGAGGTGCATTGGCGGACAT
GACGGATGAAGAAATTGAGGATCTTTTGTTAGGTTTTGCCTTAGAGGTTGCTGCAGCGGTGA
GAGCACTTCTGGAAGAATTGCGCGAAGCTGGACGTATAGATCTGTCTGACGAAGAAATTGC
CGCGCTGGCACTGCGAGTGGGTTTAAAAATGGCACGTGTCCTGGCACGGATTTTAGAGAAA
GCGACTGGGCGTGACTTATCTGCGGTATTAGAACGCGCGGAGGAACGCCTCAAAAAAGAAC
TCGCG

*Amino acid sequence of* **GDM-NT-402-5456:**

ETLELESEASLPLDVLELLLSPENLPRVLAVGRGEHFRLLSLEVLAREPGRRLTVASRAEIRLGDR
WVPVDVVLTFERQPDGSVRIRAVLTFDADAVRGALADMTDEEIEDLLLGFALEVAAAVRALLEE
LREAGRIDLSDEEIAALALRVGLKMARVLARILEKATGRDLSAVLERAEERLKKELA

*DNA acid sequence of* **GDM-NT-403-5457:**

GAAACGCTGGAGTTGGAAAGTGAAGCATCACTGCCCCTCGACGTGCTGGAGTTACTGTTATC
CCCTGAGAATCTTCCTCGTGTGTTGGCAGTGGGCCGTGGAGAACATTTCCGTCTGCTCAGTCT
TGAGGTTCTTGCGAGAGAACCGGGCCGTCGTCTTACGGTCGCTTCTCGCGCTGAAATAAGAC
TGGGTGATCGCTGGGTTCCGGTAGATGTCGTGCTGACATTTGAGCGACAGCCGGATGGCAGC
GTGCGCATTAGAGCCGTACTGACATTCGACGCCGACGCCGTGCGAGGTGCATTGGCGGACAT
GACGGATGAAGAAATTGAGGATCTTTTGTTAGGTTTTGCCTTAGAGGTTGCTGCAGCGGTGA
GAGCACTTCTGGAAGAATTGCGCGAAGCTGGACGTATAGATCTGTCTGACGAAGAAATTGC
CGCGCTGGCACTGCGAGTGGGTTTAAAATTGGCACGTGTCCTGGCACGGATTTTAGAGAAAG
CGACTGGGCGTGACTTATCTGCGGTATTAGAACGCGCGGAGGAACGCCTCAAAAAAGAACT
CGCG

*Amino acid sequence of* **GDM-NT-403-5457:**

ETLELESEASLPLDVLELLLSPENLPRVLAVGRGEHFRLLSLEVLAREPGRRLTVASRAEIRLGDR
WVPVDVVLTFERQPDGSVRIRAVLTFDADAVRGALADMTDEEIEDLLLGFALEVAAAVRALLEE
LREAGRIDLSDEEIAALALRVGLKLARVLARILEKATGRDLSAVLERAEERLKKELA

*DNA acid sequence of* **GDM-NT-404-5458:**

GAAACGCTGGAGTTGGAAAGTGAAGCATCACTGCCCCTCGACGTGCTGGAGTTACTGTTATC
CCCTGAGAATCTTCCTCGTGTGTTGGCAGTGGGCCGTGGAGAACATTTCCGTCTGCTCAGTCT
TGAGGTTCTTGCGAGAGAACCGGGCCGTCGTCTTACGGTCGCTTCTCGCGCTGAAATAAGAC
TGGGTGATCGCTGGGTTCCGGTAGATGTCGTGCTGACATTTGAGCGACAGCCGGATGGCAGC
GTGCGCATTAGAGCCGTACTGACATTCGACGCCGACGCCGTGCGAGGTGCATTGGCGGACAT
GACGGATGAAGAAATTGAGGATCTTTTGTTAGGTTTTGCCTTAGAGGTTGCTGCAGCGGTGA
GAGCACTTCTGGAAGAATTGCGCGAAGCTGGACGTATAGATCTGTCTGACGAAGAAATTGC
CGCGCTGGCACTGCGAGTGGGTTTAAAACATATGCGTGTCCTGGCACGGATTTTAGAGAAAG
CGACTGGGCGTGACTTATCTGCGGTATTAGAACGCGCGGAGGAACGCCTCAAAAAAGAACT
CGCG

*Amino acid sequence of* **GDM-NT-404-5458:**

ETLELESEASLPLDVLELLLSPENLPRVLAVGRGEHFRLLSLEVLAREPGRRLTVASRAEIRLGDR
WVPVDVVLTFERQPDGSVRIRAVLTFDADAVRGALADMTDEEIEDLLLGFALEVAAAVRALLEE
LREAGRIDLSDEEIAALALRVGLKHMRVLARILEKATGRDLSAVLERAEERLKKELA

*DNA acid sequence of* **GDM-NT-405-5459:**

GAAACGCTGGAGTTGGAAAGTGAAGCATCACTGCCCCTCGACGTGCTGGAGTTACTGTTATC
CCCTGAGAATCTTCCTCGTGTGTTGGCAGTGGGCCGTGGAGAACATTTCCGTCTGCTCAGTCT
TGAGGTTCTTGCGAGAGAACCGGGCCGTCGTCTTACGGTCGCTTCTCGCGCTGAAATAAGAC
TGGGTGATCGCTGGGTTCCGGTAGATGTCGTGCTGACATTTGAGCGACAGCCGGATGGCAGC
GTGCGCATTAGAGCCGTACTGACATTCGACGCCGACGCCGTGCGAGGTGCATTGGCGGACAT
GACGGATGAAGAAATTGAGGATCTTTTGTTAGGTTTTGCCTTAGAGGTTGCTGCAGCGGTGA
GAGCACTTCTGGAAGAATTGCGCGAAGCTGGACGTATAGATCTGTCTGACGAAGAAATTGC
CGCGCTGGCACTGCGAGTGGGTTTAAAACATGCACGTGTCCTGGCACGGATTTTAGAGAAAG
CGACTGGGCGTGACTTATCTGCGGTATTAGAACGCGCGGAGGAACGCCTCAAAAAAGAACT
CGCG

*Amino acid sequence of* **GDM-NT-405-5459:**

ETLELESEASLPLDVLELLLSPENLPRVLAVGRGEHFRLLSLEVLAREPGRRLTVASRAEIRLGDR
WVPVDVVLTFERQPDGSVRIRAVLTFDADAVRGALADMTDEEIEDLLLGFALEVAAAVRALLEE
LREAGRIDLSDEEIAALALRVGLKHARVLARILEKATGRDLSAVLERAEERLKKELA

*DNA acid sequence of* **GDM-NT-406-5460:**

GAAACGCTGGAGTTGGAAAGTGAAGCATCACTGCCCCTCGACGTGCTGGAGTTACTGTTATC
CCCTGAGAATCTTCCTCGTGTGTTGGCAGTGGGCCGTGGAGAACATTTCCGTCTGCTCAGTCT
TGAGGTTCTTGCGAGAGAACCGGGCCGTCGTCTTACGGTCGCTTCTCGCGCTGAAATAAGAC
TGGGTGATCGCTGGGTTCCGGTAGATGTCGTGCTGACATTTGAGCGACAGCCGGATGGCAGC
GTGCGCATTAGAGCCGTACTGACATTCGACGCCGACGCCGTGCGAGGTGCATTGGCGGACAT
GACGGATGAAGAAATTGAGGATCTTTTGTTAGGTTTTGCCTTAGAGGTTGCTGCAGCGGTGA
GAGCACTTCTGGAAGAATTGCGCGAAGCTGGACGTATAGATCTGTCTGACGAAGAAATTGC
CGCGCTGGCACTGCGAGTGGGTTTAAAATTGTTGCGTGTCCTGGCACGGATTTTAGAGAAAG
CGACTGGGCGTGACTTATCTGCGGTATTAGAACGCGCGGAGGAACGCCTCAAAAAAGAACT
CGCG

*Amino acid sequence of* **GDM-NT-406-5460:**

ETLELESEASLPLDVLELLLSPENLPRVLAVGRGEHFRLLSLEVLAREPGRRLTVASRAEIRLGDR
WVPVDVVLTFERQPDGSVRIRAVLTFDADAVRGALADMTDEEIEDLLLGFALEVAAAVRALLEE
LREAGRIDLSDEEIAALALRVGLKLLRVLARILEKATGRDLSAVLERAEERLKKELA

*DNA acid sequence of* **GDM-NT-407-5461:**

GAAACGCTGGAGTTGGAAAGTGAAGCATCACTGCCCCTCGACGTGCTGGAGTTACTGTTATC
CCCTGAGAATCTTCCTCGTGTGTTGGCAGTGGGCCGTGGAGAACATTTCCGTCTGCTCAGTCT
TGAGGTTCTTGCGAGAGAACCGGGCCGTCGTCTTACGGTCGCTTCTCGCGCTGAAATAAGAC
TGGGTGATCGCTGGGTTCCGGTAGATGTCGTGCTGACATTTGAGCGACAGCCGGATGGCAGC
GTGCGCATTAGAGCCGTACTGACATTCGACGCCGACGCCGTGCGAGGTGCATTGGCGGACAT
GACGGATGAAGAAATTGAGGATCTTTTGTTAGGTTTTGCCTTAGAGGTTGCTGCAGCGGTGA
GAGCACTTCTGGAAGAATTGCGCGAAGCTGGACGTATAGATCTGTCTGACGAAGAAATTGC
CGCGCTGGCACTGCGAGTGGGTTTAAAAATGATGCGTGTCCTGGCACGGATTTTAGAGAAAG
CGACTGGGCGTGACTTATCTGCGGTATTAGAACGCGCGGAGGAACGCCTCAAAAAAGAACT
CGCG

*Amino acid sequence of* **GDM-NT-407-5461:**

ETLELESEASLPLDVLELLLSPENLPRVLAVGRGEHFRLLSLEVLAREPGRRLTVASRAEIRLGDR
WVPVDVVLTFERQPDGSVRIRAVLTFDADAVRGALADMTDEEIEDLLLGFALEVAAAVRALLEE
LREAGRIDLSDEEIAALALRVGLKMMRVLARILEKATGRDLSAVLERAEERLKKELA

**Part 4: GDM-NT** variants from the third round of "novel scaffold" design

*DNA acid sequence of* **GDM-NT-507-5509:**

AAAACCACATTTGAATTGCGGTATGAAGGTGGCGAGGTGGTCCTGTCCAGTAAAACCGAAG
TGGATGGGAAGGTGTTAGAAACACGTCGCCGTGGTCGGCCGGAAAACATTGCGTTTATTGCG
AGACGTCTGCTTGCTGCGCCTGATGATGAGCTGCTTGATGTGGCTCTGAGAGCGGCGATTGC
TATATTAATTGCTGATGCTATAGCAGATTACCTCAAACTCACGGAAGGCGAAGATGCAGTGC
CATCTTTGGAATTGCGTCTTAAACTGATCGAAATCCTGGCCGAGAGAATTAAATATGAGGTC
GAAAAAATTGAAGGCGGCTTGCGTGTTACTTTCGAAGTCGATCTGGAAGCACTGGCTAAAG
ATCCGTCCGTACCAGAAGAAGTTCGGGAGATCCTGAAAAAATTCAAAAAAGCTAGTGGCAA
AATCACAATTCACGTCACAGAAGCCGACCTGCGCGCATTATTGGCCGCCATTCGCGCGGCCC
TGGCGGCCCTGCCAGCGGACGCAACACCCGAGGAAATTCAAGCTGTCGTGCGCGCAGTGGC
CGGTGCATTTATTGCGGAACGGGGTCTCACTCGTCTTGCCGAGGGGGAAATCGAATTGGAGG
TCAAA

*Amino acid sequence of* **GDM-NT-507-5509:**

KTTFELRYEGGEVVLSSKTEVDGKVLETRRRGRPENIAFIARRLLAAPDDELLDVALRAAIAILIA
DAIADYLKLTEGEDAVPSLELRLKLIEILAERIKYEVEKIEGGLRVTFEVDLEALAKDPSVPEEVRE
ILKKFKKASGKITIHVTEADLRALLAAIRAALAALPADATPEEIQAVVRAVAGAFIAERGLTRLAE
GEIELEVK

*DNA acid sequence of* **GDM-NT-543-5510:**

TTAACTCCGGAGGAACTGTTAGCCCGCGCTCGAGAATTAGAAGCCAGAGCCCCACCTTTTAC
TGTACCTGAGGGGGTTTTAGCCGCCGCGACGGAAGTGCTGCGTGAAGCAGTCGCGAAGCTC
GCCGAAAAAGTGGGAGAAGAAAAAGCAAAAGAATTGGTAGAGCGTGTGTTATCGGATGAA
TATTATGCTCGTCTGACCAAGAAACGTCTGGAGATCCTGGCAGCCGACCCTTGGATCCGGGC
TGTTATGCAGGAAATTCTGGATCTGCTGGAAGAAGCCGTCGCCAAAGGGAAAAACCCTATG
GAAATCCTGTTAAAAGCAATTCTGGAAACGGTGCGCTACCTCATCGAAAAAGTGGGAGAAG
ATACTGTAATCGCCCTGACCCGAGAACGCGCAAAAGGTAAAGTTGTTGTTCAACGGATCGTT
CGCATTGACGGTAAAACTTATGTCATAGTCGCAAGCTTACCTTTGTCTGAAGAAACACTGGA
CAAGAGATTGCTGGGGCCAATTGTATTTGTCTTTGAAGTCCCTAACGAACTTGAAGAAGAGC
TGCGTGAGCTGCTTAAAGAAGATCCGTTTCGTTGTTTAGTTAAAGCGGTGCTGAACGCGAAA
GAGTTGTATCTGTTAGAGCCAGTCCGTGAAGATGGTGAATTAACCGTCCGCCTGAAACCACT
GCATCGCTTACTGCTCGCGATCGCCGTGATGTTAAAGGACATTCGTGACCGCCTCGGGGAGG
ACATAGCTGAGGAAGCCGCCAAAGTCCTGTTAGACACCTTAAAGCGTGAGGTTGCCGCA

*Amino acid sequence of* **GDM-NT-543-5510:**

LTPEELLARARELEARAPPFTVPEGVLAAATEVLREAVAKLAEKVGEEKAKELVERVLSDEYYA
RLTKKRLEILAADPWIRAVMQEILDLLEEAVAKGKNPMEILLKAILETVRYLIEKVGEDTVIALTR
ERAKGKVVVQRIVRIDGKTYVIVASLPLSEETLDKRLLGPIVFVFEVPNELEEELRELLKEDPFRCL
VKAVLNAKELYLLEPVREDGELTVRLKPLHRLLLAIAVMLKDIRDRLGEDIAEEAAKVLLDTLK
REVAA

*DNA acid sequence of* **GDM-NT-589-5511:**

TCACGAGAAGAGATTCGTGAGGCACTGTTAGCAGCGATTGAGGAAGTTCGCGCAGACGAAG
AAGTTCGTGCAGCGCAAGCGGAGGCGGCAGCCATTGCTGACGAAGCAGCGGAAAAAGTGGC
TAGCCCAGCAGTGGCTGCAACCCTGCGCCAGCTGGCAGAGCTCATCCGTCAGCATCCCTTAG
CCCTCGCTACCCTGGAAGTTGCGCTGGGGAAGACGGAACGCGGCCGACGTCTGCGCGCACT
GTTGGTTGCAGCGGCAGAGGCCGCGGCTGGCTTGACAGAAGAAGAACGTGCGGTCTTACGG
CGCGCGTTGGAGCTTGCGGAGCGCATGCGCGATCTGGTACTGAAAAAAGAAATAAACAGTC
CCGAAGAACTGAAAGCGACTGTCGCCGAAATTCGCCGTCTTCTTGCCGAAGCCGAGCCGCTG
ATCGCCGAAGCCGAGGAAATAGCGCGCCGAAAAGGCATTAAAGTTCCGGAACCAGATGCCG
CAGTGGTTGCTGCTGCAACCCCAGAAGATGCCTTTGTAATAGGAAAAGCCGTAGCCAAGATT
GCAAAAAAATTTGGTTTGAAAACGATGTTCATGAAAATTGCAGAATTCGTTCTGGAAGGCAA
ACTGTCTCGTGAAGAGGCACTGGAAGCCTTGGGCGTAGCAGTTAGAGCGCTGCGGGCCGTG
CTTCGCGTGCGTGCGATAGTTCTGGCCATGGCCGCGCAGCTGGTTCCCGAATTACAGGTGAC
CGTGGAATTTCGTCGCCTGCTGTTGACCCTCGCAGAGATCGTAGCGCGCTTCCCGGAGGCAT
TTCTGGAAGGAATAATCGAAGCTTTAAAGGAAGGT

*Amino acid sequence of* **GDM-NT-589-5511:**

SREEIREALLAAIEEVRADEEVRAAQAEAAAIADEAAEKVASPAVAATLRQLAELIRQHPLALAT
LEVALGKTERGRRLRALLVAAAEAAAGLTEEERAVLRRALELAERMRDLVLKKEINSPEELKAT
VAEIRRLLAEAEPLIAEAEEIARRKGIKVPEPDAAVVAAATPEDAFVIGKAVAKIAKKFGLKTMF
MKIAEFVLEGKLSREEALEALGVAVRALRAVLRVRAIVLAMAAQLVPELQVTVEFRRLLLTLAEI
VARFPEAFLEGIIEALKEG

**Part 5: GDM-NT** variants used as negative control for structural analysis

*DNA sequence of* **GDM-NT-105:**

ACTCCAGAGGAACTTGTTGCTTTTATGTTGGATCCAGATGCTGACGTTGATGAAGCACTGGC
TAGACTCGAAGAATTTCTTGCTACTGCCACGCCTGAAGAGTTGCTTGAGCTTGCACGTGTAG
TGTTTGCTGCGGTTGTCGAACGATTCCCTCGGATAATGGCATTACACGCTCATCTGCGAGAA
CATGGAGGGCGCCCTCCGCCGCCGGCGGGCTTTTATTTTGCCATGGGTGTGATGACGCTGCT
GGGAGATAACGCGCGTCGGCAGTTACAGCGTAAATATGGTCTGTCAGAAGAAGAAGGGGCG
CTGTTGTTTCTTGACGCCGTTGACCGCGTTATACCTGAGCTTGTGAGAGCCCTGTTAGGACGT
GAACCGGATCCTCGCCGTCGCCGGTTAACCATGCTTTTCGTTTTGTATCTTGAATATCCAGAA
CTGTTTAAAAAATTATTCACAGAAGAAGAGCAGGCCTTCGTAAGACGTTTATTTCGCCGCGT
TCTGGCTGAAGGTGTTGAGGCCGCATTGGCCGCCTTGACACCGGAAGAACGCGCAAGATTG
GATGCCATGTACGCTAAATTTCTGGCGGCAGCTGAGGAGCTCAAGAAGAAATATCCGGAAG
CCGCTGCAGAGATGGAAGCCCTCGCAGCCCGGCTGCGAGAGCTGGCGCTTCGGATCAATGC
CCGGGTGGCTTCCCTGTCTGAAGCAGACCAGGCGGCACTGCAGGCAGCGGTTGATGCCGCAT
TCGCCGAAGCGGCTGCTCGTACGCCGCTTGCTGATCTTTTGGCG

*Amino acid sequence of* **GDM-NT-105:**

TPEELVAFMLDPDADVDEALARLEEFLATATPEELLELARVVFAAVVERFPRIMALHAHLREHG
GRPPPPAGFYFAMGVMTLLGDNARRQLQRKYGLSEEEGALLFLDAVDRVIPELVRALLGREPDP
RRRRLTMLFVLYLEYPELFKKLFTEEEQAFVRRLFRRVLAEGVEAALAALTPEERARLDAMYAK
FLAAAEELKKKYPEAAAEMEALAARLRELALRINARVASLSEADQAALQAAVDAAFAEAAART
PLADLLA

*DNA sequence of* **GDM-NT-141:**

GGGGACGAGGTTAAAGCTGCCCTGATCCGTGCTGCCGAAGCAGTATATGCGGGTGATCGTG
AGGGTGCGCGCGCTGCCCTGGCGGAGGCAGTTGACGCCCTGGAGAAGAGATCCCGGGAACA
AGTGAGACTGGCAATCAAAGGTCTGCTCGAAATAGCAAAATTTGTAGCAGAACTGCTCAAG
GAGTTAACCCCAGAACAAAAAGAAGCGCTGCTGGCGCAGGGCGTCCTTGAGGCAATTAAAC
GTTTTTTCCTTGCCCTTGCTGCCATAGCAGGTCTGGTGGGCGGTGACGAAATGGTTGATGAA
GTGCTTGAAATGGGAAAAGAGGTCGCGGAACTTTTAAAGGAGGTTGTTGGGGAGGAAAAAG
CGGACGAATTTAAGGAAGAAGTCGAAAAAGGAGTTGCTGAACTTCGGACACTTACTCCAGA
AGAACGCAAAGAAAGAGCCGTTGATCTCCTGCTGGAAGTGTTAGAAGCGATGGTCGCCCGG
GGTCTTAGCCTTGACCAGATTAAAGAGTTTGCGCGTTTGTTGGCAGAACTGCTGACAGCTGA
AACCGAAGAAGAAGTAAAAGCAGTTGTCGAAAAAGTGAAAGCAATCCACAAAGCGAGTGT
TGCACGTCTCAAGGCA

*Amino acid sequence of* **GDM-NT-141:**

GDEVKAALIRAAEAVYAGDREGARAALAEAVDALEKRSREQVRLAIKGLLEIAKFVAELLKELT
PEQKEALLAQGVLEAIKRFFLALAAIAGLVGGDEMVDEVLEMGKEVAELLKEVVGEEKADEFK
EEVEKGVAELRTLTPEERKERAVDLLLEVLEAMVARGLSLDQIKEFARLLAELLTAETEEEVKAV
VEKVKAIHKASVARLKA

**Table S9**. Summary of mutations introduced in each round of evolution for ***Tm*Pgb-PIP** variants:

All mutations are compared to *Tm*Pgb-wt. New mutations are highlighted in red.

| Entry | Variant | Mutations from ***Pa*Pgb-wt** |
|---|---|---|
| 1 | ***Tm*Pgb-PIP-G1-5442** | I148T |
| 2 | ***Tm*Pgb-PIP-G2-5443** | S17G, L141S, I148T, Y170H |
| 3 | ***Tm*Pgb-PIP-G3-5444** | S17G, C30S, L141S, I148T, Y170H |

| | | |
|---|---|---|
| 4 | ***Tm*Pgb-PIP-G4-5445** | S17G, C30S, F62W, L141S, I148T, Y170H |
| 5 | ***Tm*Pgb-PIP-G5-5446** | S17G, C30S, F62W, S141N, I148T, Y170H |
| 6 | ***Tm*Pgb-PIP-G6-5447** | S17G, C30S, F62W, H66E, S141N, I148T, Y170H |
| 7 | ***Tm*Pgb-PIP-G7-5448** | S17G, C30S, F62W, S65G, H66E, G78A, S141N, I148T, Y170H, Q180S, G192S |

**Table S10**. Summary of mutations introduced in the final round of engineering for **GDM-NT** variants:

| Entry | Variant | Mutations from **GDM-NT-362-5452** |
|---|---|---|
| 1 | **GDM-NT-399-5453** | V53L, I68V, E99G, H155L |
| 2 | **GDM-NT-400-5454** | H155M-M156L |
| 3 | **GDM-NT-401-5455** | V53L-I68V-E99G-H155M-M156L |
| 4 | **GDM-NT-402-5456** | V53L-I68V-E99G-H155M-M156A |
| 5 | **GDM-NT-403-5457** | V53L-I68V-E99G-H155L-M156A |
| 6 | **GDM-NT-404-5458** | V53L-I68V-E99G |
| 7 | **GDM-NT-405-5459** | V53L-I68V-E99G-M156A |
| 8 | **GDM-NT-406-5460** | V53L-I68V-E99G-H155L-M156L |
| 9 | **GDM-NT-407-5461** | V53L-I68V-E99G-H155M |

#### Design Pipeline Details

##### Initial Design Details (First Round)

The initial designs were made based on AlphaFold 3 (AF3) predictions of ***Tm*Pgb** and ***Ap*Pgb**. For details of this round of design, see descriptions in our report on AlphaProtein Novo (*6*).

##### Partial Diffusion Details (Second Round)

We performed partial diffusion from "parent" designs **GDM-NT-121** and **GDM-NT-151**, which are described in companion work (*6*). For each "parent" design, we generated two groups of designs by applying enough renoising steps to ensure the post-denoising median TM-score distributions centered around 0.95 or 0.99. **GDM-NT-226** and **GDM-NT-362** were created by this process, derived from **GDM-NT-121** and **GDM-NT-151** respectively.

*Metric Definitions*

We used AF3 (*7*) for structure prediction. Metric definitions are as follows:

*Subsets*

- `sampled/` refers to metrics on the sampled structure
- `sequence/` refers to metrics on the sequence (from LigandMPNN)
- `monomer/` refers to metrics on a monomer (apo) fold of the design
- `complex/` refers to metrics on a complex (protein with ligands from the motif) fold of the design
- `complex/expanded_motif/` refers to motif metrics where the entire residue of a motif (in this case the heme-ligating histidine) in included, versus motif metrics which only involve the atoms fixed in the motif

*Sample Metrics*

- percent_ligand_bb_clashes: Percent of ligand atoms within 2.0 Å of a protein backbone atom
- max_consecutive_identical_aatypes: Max consecutive identical residues
- max_single_aatype_fraction: Max fraction of any single amino acid type
- num_blocklisted_motifs: Number of subsequences matching anything from the following sequences.
    - WSHPQFEK (Strep-tag)
    - EQKLISEEDL (c-Myc-tag)
    - ENLYFQ[SGAMCH] (TEV protease)
    - LEVLFQGP (3C protease cleavage site)
    - YPYDVPDYA (HA-tag)
    - HHHHH (His-tag)
    - DYKDDDDK (FLAG-tag)
    - GKPIPNPLLGLDST (V5-tag)
    - LVPRGS (Thrombin cleavage)
- unknown_count: Number of unknown ('X') amino acid type
- sample_his_rmsd: the RMSD of the ligating His sidechain atoms from the ideal pentagonal plane

*Consistency Metrics*

- tm_score: Template Modelling score (*8*) measuring length-normalised global structural similarity.
- interface_lddt: LDDT (*9*) computed between residues or atoms not on the same chain.
- mean_pocket_bb_aligned_ligand_rmsd: Mean RMSD of ligand heavy atoms after superimposing the predicted structure onto the reference on backbone atoms within 10 Å of the ligand.
- mean_pocket_bb_rmsd: Mean RMSD of pocket backbone atoms as defined above.

- motif_allatom_rmsd: All-atom RMSD between AF3 prediction and input motif structure
- motif_with_ligand_allatom_rmsd: same as above, with inclusion of the ligand motif atoms

*Confidence Metrics*

- predicted_distance_error: AF3 predicted distance error over all atoms
- chain_pair_pde_mean_sym: Symmetrised mean predicted distance error between chains.
- ligand_plddt_mean: Mean ligand pLDDT.

*Structure Prediction Metrics*

- pocket_C9_min_atom_dist: the minimum distance between any atom of the designed protein to the tail carbon atom ('C9') of the substrate's phenyl group, based on structure prediction of the sample with the substrate and heme cofactor
- pocket_C9_allatom_neighbors_mean: the number of design atoms within 4, 6, or 8 Å, averaged across the different distance thresholds, of the substrate's 'C9' atom, based on structure prediction with the substrate
- propionate_hbond_num_atoms: the number of potential H-bond partner atoms to any of the heme propionate's oxygens (O1A, O2A, O1D, O2D), based on the structure prediction of the protein in complex with the heme and substrate
- propionate_hbond_num_res: the number of unique residues corresponding to the above

We also included metrics that quantify the variance of two relevant ligand atoms across structure prediction with five different representations of the small molecules and states involved: heme and substrate, heme and piperidine, heme and pyrrolidine, and the heme covalently bound to the substrate with two different SMILES representations. The SMILES representations are:

- CC1=C(CCC(O)=O)C2=NC1=Cc1c(C=C)c(C)c3\C=C4/N=C(/C=C5\N([Fe](N(CCCCCC6=CC=CC=C6)N=N)n13)\C(=C/2)C(CCC(O)=O)=C5C)C(C)=C4C=C
- [Fe]N(CCCCCC1=CC=CC=C1)N=N.CC1=C(CCC([O-])=O)/C2=C/C3=N/C(=C\C4=C(C)C(C=C)=C([N-]4)/C=C4\N=C(\C=C\1/[N-]\2)C(C=C)=C4C)/C(C)=C3CCC([O-])=O.

And the resulting metrics are:

- fe_self_rmsd: the RMSD of the iron atom in the porphyrin across the five different structure predictions, after global alignment to the sampled structure
- n_self_rmsd: same as above, but for the ligating nitrogen of the substrate

Finally, we specified a metric for regioselectivity based on the Hydrogen Atom Transfer step of the reaction, where the near-attack conformation to the hydrogen of the fourth vs fifth carbon in the alkyl chain suggests regioselectivity for pyrrolidine and piperidine, respectively. Because AF3 does not generate hydrogens, we instead simply use the distance to the corresponding carbon as a proxy metric.

- piperidine_propensity: the difference in distance between nitrogen and the fifth carbon away from the substrate from structure prediction, and the distance between the nitrogen and the fourth carbon. We take the minimum value over 7 structure prediction seeds of the design in complex with "heme and substrate" (as described above) for filtering. Note that AF3 Leaving Atom (AF3-LA) (*6*) was used for computing this metric. All other metrics were computed with AF3.

*Filtering Process*

After denoising and resequencing with LigandMPNN (*10*), the samples were first filtered by:
(*also applied in multi-resequence filtering)*

- sampled/percent_ligand_bb_clashes <= 5.0
- sequence/max_consecutive_identical_aatypes = 5
- sequence/max_single_aatype_fraction <= 0.25

- sequence/num_blocklisted_motifs <= 0
- sequence/unknown_count <= 0
- monomer/predicted_distance_error <= 2.0
- monomer/tm_score >= 80
- complex/chain_pair_pde_mean_sym <= 2.0
- complex/interface_lddt >= 30
- complex/mean_pocket_bb_aligned_ligand_rmsd <= 2.5
- complex/mean_pocket_bb_rmsd <= 2.0
- complex/expanded_motif/motif_allatom_rmsd <= 1.5
- complex/expanded_motif/motif_with_ligand_allatom_rmsd <= 2.5
- complex/ligand_plddt_mean >= 75

(*not applied in multi-resequence filtering)*

- pocket_C9_min_atom_dist between 2.8 and 3.8
- pocket_C9_allatom_neighbors_mean between 38 and 62, inclusive
- propionate_hbond_num_atoms between 3 and 6, inclusive
- propionate_hbond_num_res between 3 and 5, inclusive
- sampled_his_rmsd <= 0.08
- ligating_n_self_rmsd <= 0.8
- his_heme_distance <= 3.0

Additional criteria for a subset of samples:

- The piperidine_propensity filter as predicted by AF3-LA was applied at <= 0.35.
- Samples were subjected to a "multi-resequence" filter. For each candidate (comprising a backbone and its "original" LigandMPNN sequence passing the full set of filters described above), we generated 7 "additional" LigandMPNN sequences. A candidate passed the multi-resequence filter if all of these additional 7 re-sequences met the criteria labeled "Also applied in multi-resequence filtering." These 7 additional re-sequences were only used for filtering: we use the original LigandMPNN sequence for experimental characterisation.

*Selection Process*

In total, 96 total partial diffusion designs were tested (48 derived from **GDM-NT-121** and 48 derived from **GDM-NT-151**). Within each 48-design set, half (24) had a post-denoising median TM-score of 0.95, and the other half had a median TM-score of 0.99. These subgroups were further divided evenly based on previously described selection criteria: 12 designs passed only the base filters, while the remaining 12 passed both the base and additional filters.

Designs in the 0.99 TM-score group were chosen at random. Conversely, designs in the 0.95 TM-score group were selected *via* K-medoids clustering by TM-score to identify representative centroids for each design facet. To minimize the contribution of the termini during clustering, we trimmed the sequences before determining the TM-score by removing the first 4 and last 2 residues from **GDM-NT-121** designs, and the first 3 and last 1 residues from **GDM-NT-151** designs.

#### Structural Diversity of Designs

##### *Design Similarity to PDB Chains*

Heme-containing structures were retrieved from a March 2026 PDB (*11*) snapshot. Heme cofactors were restricted to the following CCD codes representing iron-porphyrin variants: HEM, HEC, HEA, HEB, HEO, HDD, SRM, HCO, 1FH, 2FH. We pre-filtered to PDB entries containing at least one non-polymer entity, then identified heme-containing entries by parsing ligand residue names, yielding 7,543 heme-containing structures. We further restricted this analysis to PDB chains that are ligated to a heme, using proximity to the Fe atom of each heme to disambiguate multiple chains. Sequences were then clustered with MMseqs2 (*12*) (easy-cluster, 70% identity, 80% bidirectional coverage, greedy set-cover mode), yielding 1,016 non-redundant chains.

We computed all all-vs-all structural alignments using TM-align on the representative single-chain structures plus the two partial diffusion designs (**GDM-NT-226** and **GDM-NT-362**) to obtain TM-scores. The pairwise distance between structures was defined as 1 minus the mean TM-score normalised to both chain lengths.

Separately, each design was aligned against all heme-proximal PDB chains (before clustering) using TM-align to identify the closest structural match without sequence-identity reduction. The resulting TM-align superposition for this top-scoring hit is shown in Figure 3b.

##### *Visualizations*

Each heme-binding chain was assigned to a functional family by keyword matching against the concatenated PDB entry description and title fields, applied in the following priority order: P450 ("p450" or CYP followed by a digit or word boundary); Globin ("globin", "hemoglobin", or "myoglobin"); Peroxidase ("catalase-peroxidase", "katg", or "peroxidase"); Peroxygenase ("peroxygenase" or "UPO"); Catalase ("catalase", excluding any catalase-peroxidases as they are classified under Peroxidase); Dioxygenase ("dioxygenase", "TDO", or "IDO"/"IDO1"/"IDO2"); Heme oxygenase ("heme oxygenase" or "haem oxygenase"); Nitric oxide synthase ("nitric oxide synthase"); Cytochrome c ("cytochrome c" without co-occurring "oxidase", "p450", or "peroxidase"; note this results in a heterogeneous group which includes single heme cytochrome c's as well as multi heme proteins); Siroheme (SRM CCD code or "siroheme"). Remaining entries were labelled "Other heme proteins".

Metric MDS was applied to the precomputed distance matrix using scikit-learn (*13*) (n_components=2, dissimilarity='precomputed', metric=True, init='classical_mds', n_init=1, max_iter=10000, random_state=42). Kruskal's stress-1 was 0.35, indicating moderate distortion in the two-dimensional projection. The most structurally similar PDB chains for each design are highlighted in Fig. 3b.

UMAP (*14*) was applied to the same distance matrix (n_components=2, metric=precomputed, n_neighbors=30, min_dist=0.3, random_state=42). Parameters were selected from a sweep spanning local (n_neighbors=5, min_dist=0.05) to global (n_neighbors=50, min_dist=0.5) regimes; the chosen values best preserved global cluster topology while maintaining interpretable inter-family separations.

In both plots, PDB representatives are coloured by the protein family, and de novo designs are overlaid as gold stars.

*UMAP projection based on all-pairs TM-align scores across non-redundant heme-binding chains*

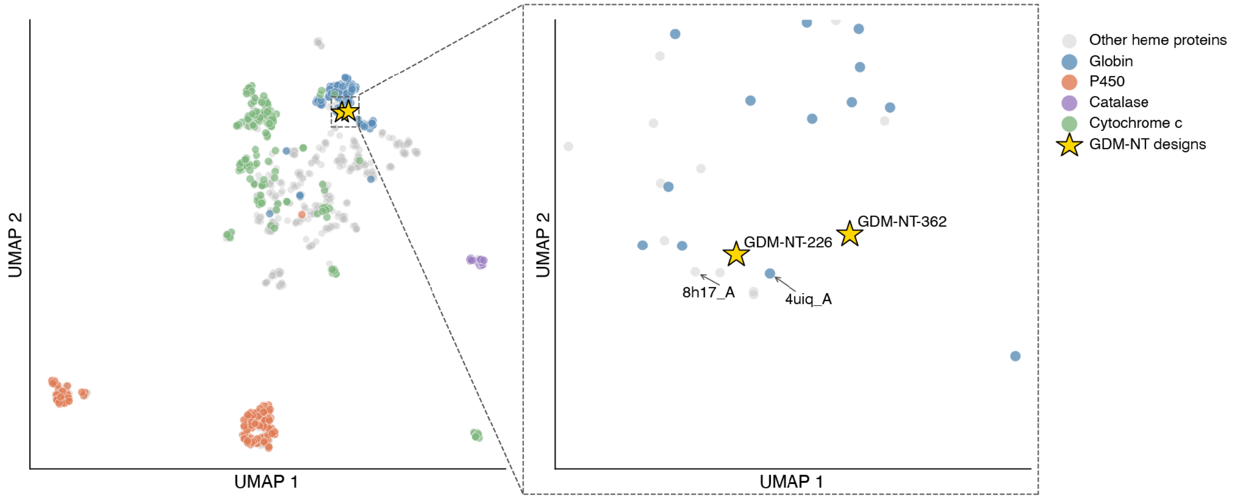

**Figure. S1** UMAP projection based on all-pairs TM-align scores across non-redundant heme-binding chains from the PDB (March 2026), clustered at 70% sequence identity (n=1016). Each point represents a cluster representative, colored by family, with GDM-NT designs projected into the same space. Note: the cytochrome c label is a heterogeneous grouping defined by PDB annotation. The inset (right) zooms into the region surrounding the two designs, with the most structurally similar PDB chain for each design labelled (8h17_A for **GDM-NT-226**; 4uiq_A for **GDM-NT-362**). Structural superpositions of these closest matches are shown in Figure 3b. Designs occupy a sparsely populated region of the landscape

**Motif-Informed Design Details (Third Round)**

With the successful designs obtained from partial diffusion and post-design evolution, we sought to test whether these successful designs could in return inform the design pipeline. Specifically, we anticipated extracting motifs from successful designs after the second round could lead to higher success rate and better scaffold diversity. Guided by this principle, the third design round explored alternative noncanonical residue options at the proximal axial position while otherwise keeping the design approach consistent with the second design round, with exceptions in motif generation and metrics, as described below.

*Motif Generation*
We extracted haem-bound piperidine poses and their corresponding proximal axial residues from AF3-LA predictions of **GDM-NT-362**, **GDM-NT-401**, **GDM-NT-406**. In contrast to earlier rounds, we used AF3-LA (instead of AF3), and we did not enforce a covalent bond between the product and the iron on the HEM ligand. Previously, this extra covalent bond was used to ensure coordination between HEM and the product; we found it unnecessary for this round.

*Metrics*
Because this design round incorporates axial ligands other than histidine, we generalized several metrics to accommodate the geometries introduced by these alternative axial ligands.

1) Axial Heme Distance:
This metric measures the distance between the "tip atom" of the residue at the proximal axial position and the iron on the HEM ligand. We defined the following tip atoms for each axial ligand:

- **NE2** for Histidine (**GDM-NT-362**)
- **SD** for Methionine (**GDM-NT-401**)
- **CD1** for Leucine (**GDM-NT-406**)

For **GDM-NT-362**, AF3-LA predicts an axial heme distance of 2.1 Å. The corresponding threshold used in the second round of nitrene transfer design was 3.0 Å. For motifs derived from **GDM-NT-401** and **GDM-NT-406**, we established thresholds that maintain the same ratio between the threshold and the mean axial heme distance across 10 folding seeds using AF3-LA.

The maximum thresholds used per motif were:

- **GDM-NT-362**: ≤ 3.0 Å
- **GDM-NT-401**: ≤ 4.0 Å
- **GDM-NT-406**: ≤ 4.8 Å

2) Sampled Axial Residue RMSD:
In previous rounds, the axial residue RMSD metric took the residue corresponding to the axial ligand from the generator sample, aligned it to an idealized geometry of that same side-chain, and computed the RMSD.

The goal of this metric is to evaluate the validity of the terminal side-chain frame of the sampled residue. For histidine, the entire side-chain is rigid and can be represented by a single frame, but this implementation had to be adapted for the more flexible leucine and methionine side-chains.

To generalize this metric, we recalibrated the RMSD for histidine into a threshold based on the maximum absolute bond length error (compared to CCD idealised coordinates). An RMSD threshold of 0.08 was previously used for the axial residue RMSD. By regressing the maximum absolute bond length error onto the RMSD for histidine ($R^2 = 0.97$), we derived a maximum absolute bond length error cutoff of 0.14 Å.

Additionally, we introduced a maximum absolute bond angle error cutoff of 20°. We observed that the generator would sometimes produce samples, particularly for leucine, with acceptable bond lengths but severely distorted bond angles. This loose 20° cutoff was applied across all motifs to filter out such anomalies.

3) Multi-resequencing:
In the third round of design, we exclusively selected designs passing 7/7 multi-resequences. This is in contrast to the novel-scaffold facet of the second round of design described in AP Novo (*6*), which selected designs passing at least 4/7 multi-resequences.

*Scaffold Novelty of the Third Round of Design*

The three top-performing designs **GDM-NT-507**, **GDM-NT-543**, **GDM-NT-589** all feature low TM-scores compared to known natural haemproteins. Due to limited space, we show the TM-align superposition of **GDM-NT-543** (blue) with its closest structural match (gray) in the following figure.

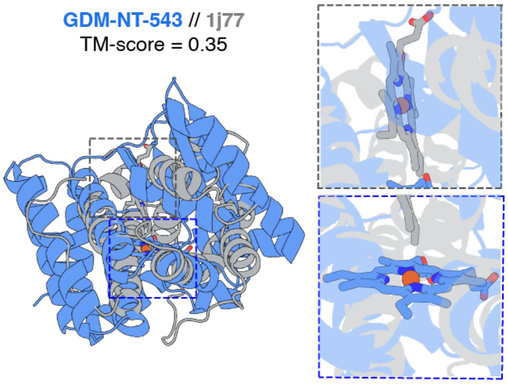

**Figure S2. Structural diversity of additional active designs.** TM-align superposition of **GDM-NT-543** (blue) with its closest structural match (gray) from the PDB (March 2026), showing folds distinct from known haem proteins. Insets show the haem binding pocket of designs and natural proteins

*Beta-Sheet Content Analysis Across Natural and Designed Haem Proteins*

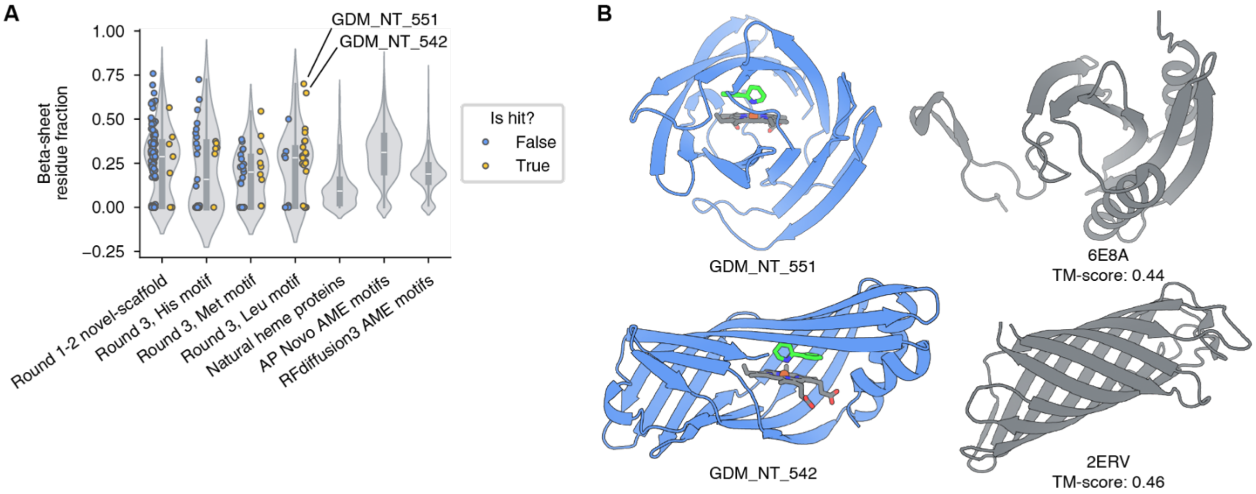

**Figure. S3. Beta-sheet content of designs. (A)** Fraction of residues annotated as beta sheet by DSSP, for different design rounds, natural haem-containing proteins in the PDB, AP Novo designs generated using the input motifs from the AME benchmark, and RFdiffusion3 designs generated for the same AME motifs. PDB proteins were filtered to 20,402 entries containing HEM and then clustered by mmseqs2 at 70% identity to 1,019 entries. Round 1-2 designs and generation settings for the AME benchmark are as described by the AP Novo paper (*6*). **(B)** Design structures (blue) and nearest Foldseek hit in the PDB (grey) for the two active designs with highest beta-sheet content. **GDM-NT-551** has a ***Tm*Pgb-wt** normalized activity of 3.9 and **GDM-NT-542** has an activity of 0.74. Both designs have 99:1 rr for piperidine.

**Design with No-Protein Motif**
We evaluated the distribution of axial ligands when sampling without an explicitly defined axial ligand in the motif.

*Distribution of Residues at the Presumed Axial Position Across Heme-binding Protein Sequences in the PDB and "no-protein motif" designs*

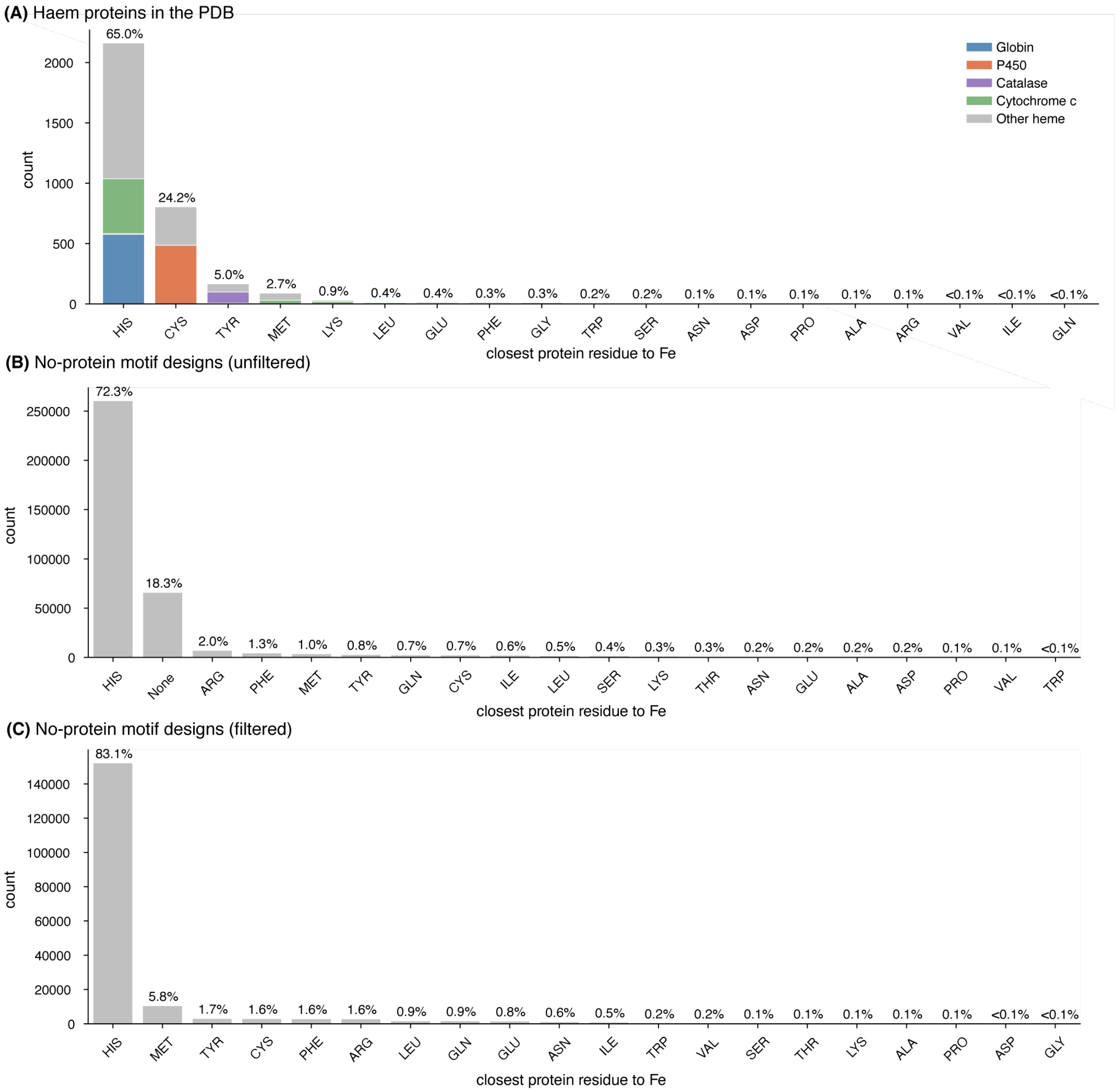

**Figure S4 Distribution of the closest protein residue to Fe in natural haemoproteins and no-protein motif designs. (A)** Haem-binding protein sequences in the PDB. For each haem iron, the nearest protein residue was identified from unique haem-binding sequences in the March 2026 PDB. Entries were deduplicated by collapsing those sharing the same protein sequence and nearest-residue identity; non-standard amino acid codes (e.g., methylhistidine, selenocysteine) were excluded (n = 3,326 unique entries). Bars are coloured by protein family. Histidine is the most frequently observed nearest residue, notably serving as the axial ligand in globins. Cysteine is the second most common characteristic of P450 thiolate ligation. **(B)** Unfiltered no-protein motif designs. Designs were generated using input motifs consisting only of a haem cofactor and N-heterocyclic product, without specifying a presumed axial ligand (n = 360,344 designs). The closest protein residue to Fe was assigned as the amino acid with the nearest side-chain atom to the haem iron, provided the distance was ≤3.5 Å; designs with no residue within this cutoff are labelled "None" and are not included in the next panel. **(C)** No-protein motif designs after basic quality filtering. From the initial no-protein motifdesigns in (B), filters on sequence composition and ligand clashes were applied (retaining n = 183,122, 50.8%). The closest residue to Fe was assigned by majority vote across 7 AF3 seeds, designating the residue with the nearest side-chain atom to the haem iron (≤3.5 Å).

*Generator*

We identified the side-chain atom closest to the HEM iron in the sampled design. If this distance was 3.5 Å or closer, that atom's residue was designated as the axial ligand for that seed.

**Table S11.** Distribution of residues at proximal axial position using no-protein motif designs (unfiltered)

| Residue | Count | Percent |
|---|---|---|
| HIS | 260580 | 72.3% |
| None | 65968 | 18.3% |
| ARG | 7089 | 2.0% |
| PHE | 4515 | 1.3% |
| MET | 3706 | 1.0% |
| TYR | 2865 | 0.8% |
| GLN | 2462 | 0.7% |
| CYS | 2387 | 0.7% |
| ILE | 2192 | 0.6% |
| LEU | 1671 | 0.5% |
| SER | 1480 | 0.4% |
| LYS | 1174 | 0.3% |
| THR | 928 | 0.3% |
| ASN | 853 | 0.2% |
| GLU | 688 | 0.2% |
| ALA | 603 | 0.2% |
| ASP | 549 | 0.2% |
| PRO | 338 | 0.1% |
| VAL | 224 | 0.1% |
| TRP | 72 | <0.1% |

*Generator + Light Filtering (AF3 Agreement)*

We evaluated designs with the heme and substrate across 7 structure prediction seeds. For each seed, we identified the side-chain atom closest to the HEM iron. If this distance was 3.5 Å or closer, that atom's residue was designated as the axial ligand for that seed.

We then took a majority vote across the 7 seeds to determine the final axial ligand, filtering out any designs where no single residue held a majority.

We removed any samples where the maximum axial heme distance exceeded 5.5 Å and applied the following basic filters on the generated sequence and structure:

sample/percent_ligand_bb_clashes: ≤ 5.0
sequence/max_consecutive_identical_aatypes: ≤ 5
sequence/max_single_aatype_frac: ≤ 0.25
sequence/num_blocklisted_motifs: ≤ 0
sequence/unknown_count: ≤ 0
sequence/seq_len: ≥ 90

These basic filters removed roughly half of the initial samples:

- **Passed:** 183,122 / 360,344 (50.8%)
- **Removed:** 177,222 (49.2%)

Of the remaining 183,122 samples, the distribution of generated axial ligands was as follows:

**Table S12.** Distribution of residues at proximal axial position using no-protein motif designs (filtered)

| Residue | Count | Percent |
|---|---|---|
| HIS | 152,200 | 83.1% |
| MET | 10,550 | 5.8% |
| TYR | 3,043 | 1.7% |
| CYS | 2,999 | 1.6% |
| PHE | 2,881 | 1.6% |
| ARG | 2,849 | 1.6% |
| LEU | 1,715 | 0.9% |
| GLN | 1,691 | 0.9% |
| GLU | 1,552 | 0.8% |
| ASN | 1,164 | 0.6% |
| ILE | 895 | 0.5% |
| TRP | 328 | 0.2% |
| VAL | 280 | 0.2% |
| SER | 251 | 0.1% |
| THR | 213 | 0.1% |
| LYS | 179 | 0.1% |

| Residue | Count | Percent |
|---|---|---|
| ALA | 143 | 0.1% |
| PRO | 111 | 0.1% |
| ASP | 75 | <0.1% |
| GLY | 3 | <0.1% |

As these data demonstrate, without explicitly enforcing the axial ligand in the motif, the generator predominantly defaults to histidine and does not generate samples with alternative axial ligands at meaningful frequencies.

It is also worth noting that this distribution reflects only light filtering. Once the full critic is applied–where pass rates are approximately 10 times lower for methionine compared to histidine–this distribution tilts even further toward histidine.

##### Comparison between GDM-NT-401 and *Tm*Pgb-PIP-G7

*Predicted Structures from AF3-LA of Different Protein Scaffolds in Complex with Alkyl Nitrenoid Intermediates*

Predicted structures from AF3-LA of the ***Tm*Pgb-PIP-G7** and **GDM-NT-401** scaffolds in complex with various substrates from the substrate scope analysis. For each complex, 8 structural predictions were generated using AF3-LA with distinct random seeds. The final displayed structure was chosen by taking the top two predictions based on substrate-atom pLDDT, and subsequently selecting the conformation with the minimum N1–C5 distance (measured between the substrate nitrogen and the fifth carbon of the alkyl chain), or the N1-C6 distance for substrate **2n**. Substrates **2a** and **2b** are shown together as AF3-LA cannot model different isotopes. Using this filtering pipeline, AF3-LA consistently predicts a catalytically favourable near-attack conformation for substrates within the **GDM-NT-401** scaffold - a geometry notably absent in the ***Tm*Pgb-PIP-G7** scaffold. The ***Tm*Pgb-PIP-G7** sequence was folded incorporating multiple sequence alignment (MSA) data (using databases MGnify 2022-05, UniRef90 2022-05, SmallBFD).

*Predicted structure of* **GDM-NT-401** *in complex with alkyl nitrenoid intermediates derived from substrate scope study in Fig. 4*

All predicted structures feature a near-attack conformation with the alkyl chain bending toward the heme co-factor.

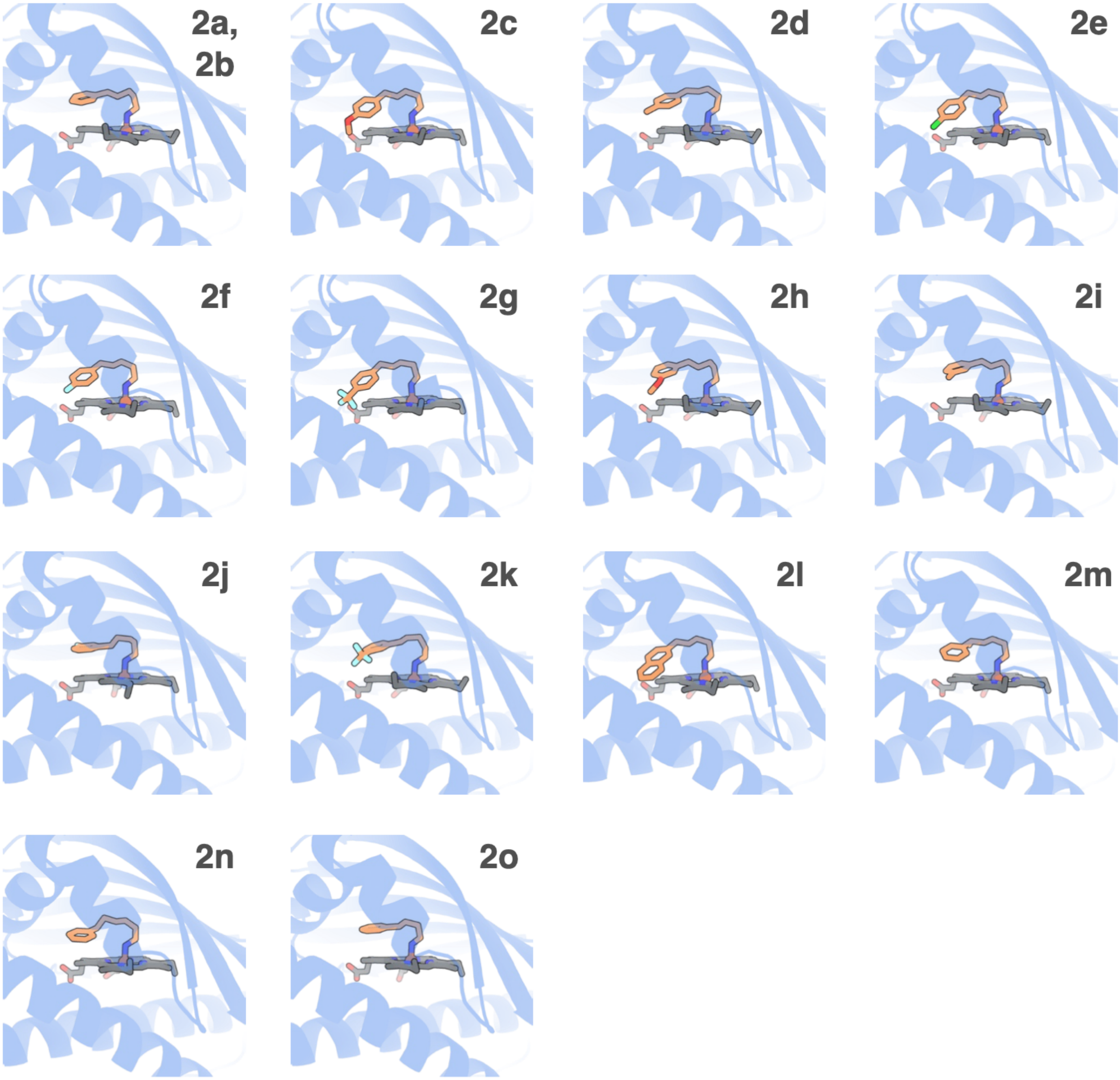

**Figure S5.** Predicted structures from AF3-LA of **GDM-NT-401** in complex with various alkyl nitrenoid intermediates from the substrate scope study.

*Predicted structure of* ***Tm*****Pgb-PIP-G7** *in complex with alkyl nitrenoid intermediates derived from substrate scope study in Fig. 4*

All predicted structures feature a linear conformation

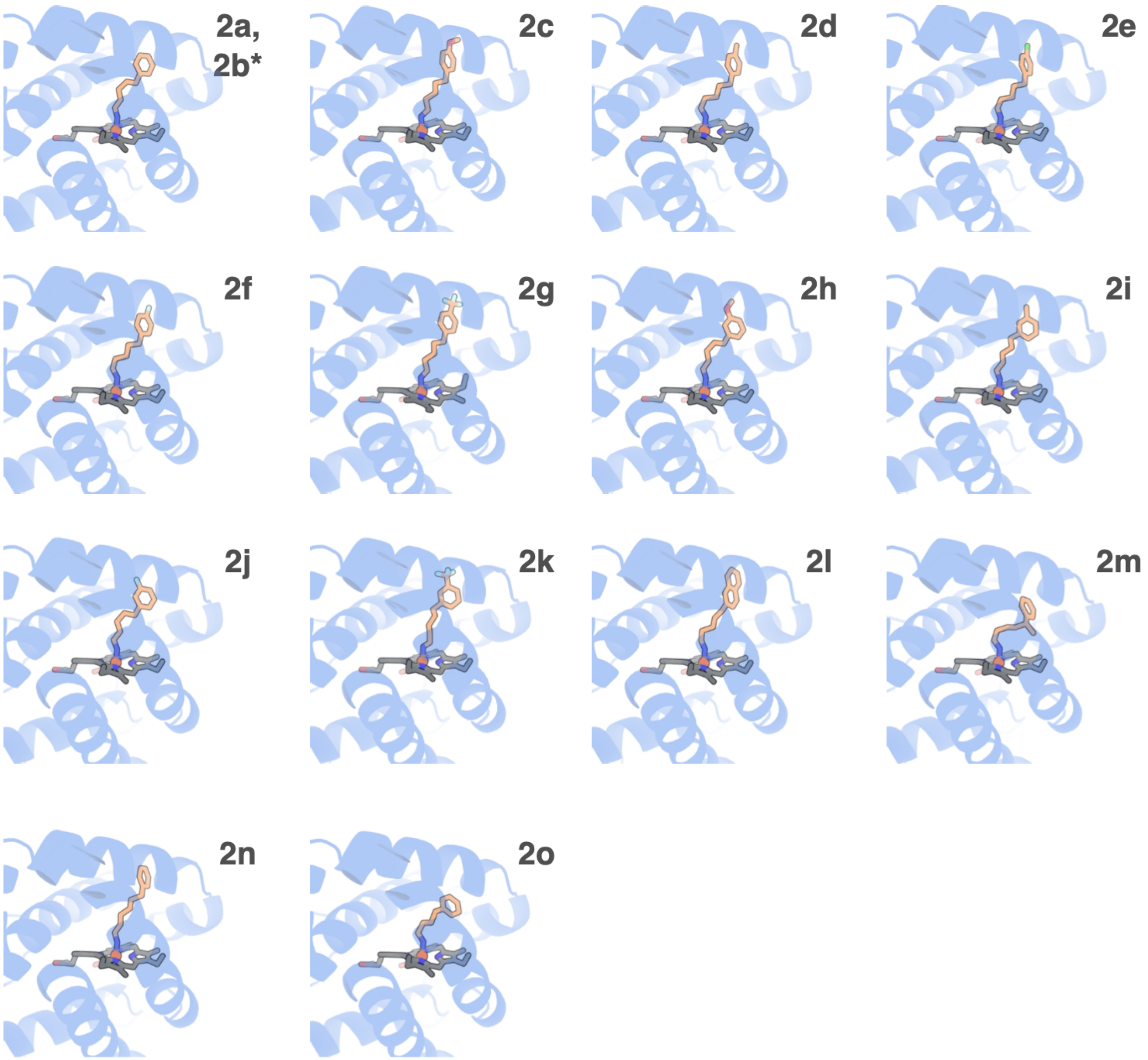

**Figure S6** Predicted structures from AF3-LA of ***Tm*****Pgb-PIP-G7** in complex with various alkyl nitrenoid intermediates from the substrate scope study.

*Summary of N1–C5 distances in predicted structures from AF3-LA of* **GDM-NT-401** and ***TmPgb-PIP-G7*** *in complex with various substrates from the substrate scope analysis.*

In general, the predicted structures of **GDM-NT-401** in complex with different alkyl nitrenoid intermediates have a shorter N1–C5 distances compared to ***Tm*Pgb-PIP-G7**. This result corresponds well with the overall higher yields obtained with **GDM-NT-401**.

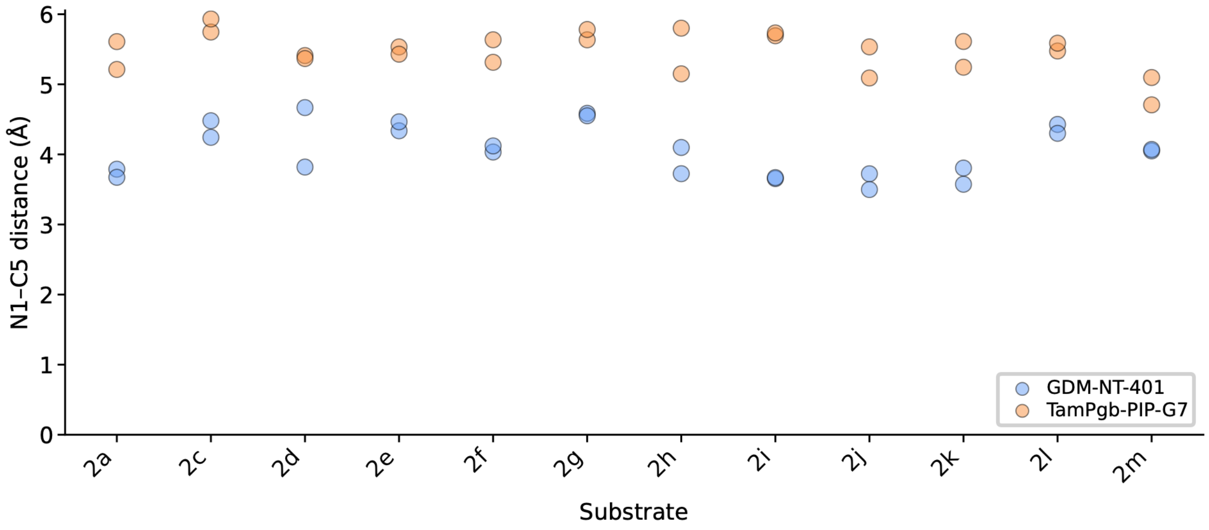

**Figure S7.** N1–C5 distances in predicted structures from AF3-LA of ***Tm*Pgb-PIP-G7** in complex with various alkyl nitrenoid intermediates from the substrate scope study.

*Comparison between* ***Tm*Pgb-PIP-G7** and **GDM-NT-401** *for the formation of* **2n**

A comparison between ***Tm*Pgb-PIP-G7** and **GDM-NT-401** on their predicted structure from AF3-LA in complex with corresponding alkyl nitrenoid intermediate for the formation of **2n** is shown below in Fig. S6. 1) **GDM-NT-401** demonstrated a close-to-attack conformation, while ***Tm*Pgb-PIP-G7** demonstrated a relatively linear conformation. The corresponding N1–C6 distances also demonstrated that **GDM-NT-401** has a shorter average distance than ***Tm*Pgb-PIP-G7**. This result corresponds well to the experimental data where 13% yield of 2n was observed using **GDM-NT-401** and no detectable activity was observed using ***Tm*Pgb-PIP-G7**. 2) Despite the average N1–C6 and N1–C5 distances (Fig. S5) for **GDM-NT-401** are similar (both around 4 Å), the N1–H–C6 versus N1–H–C5 distance and angle clearly favours the formation of a seven-membered ring.

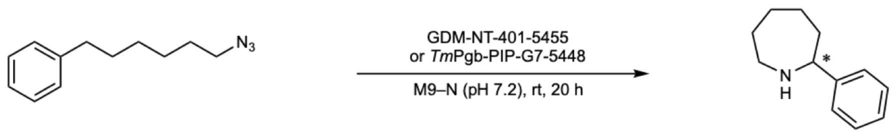

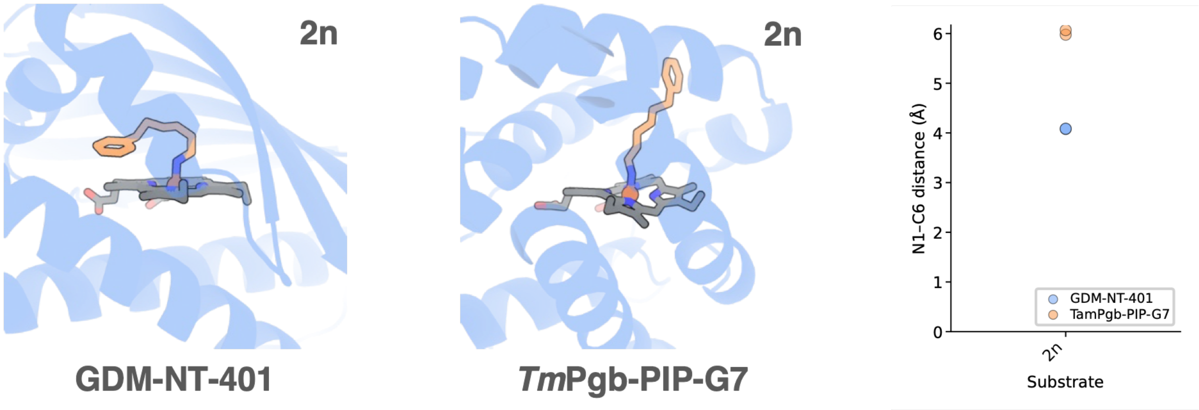

**Figure S8.** Predicted structures from AF3-LA of **GDM-NT-401** in complex with alkyl nitrenoid intermediate to form **2n**.

*UV-Vis Analysis of GDM-NT-401*

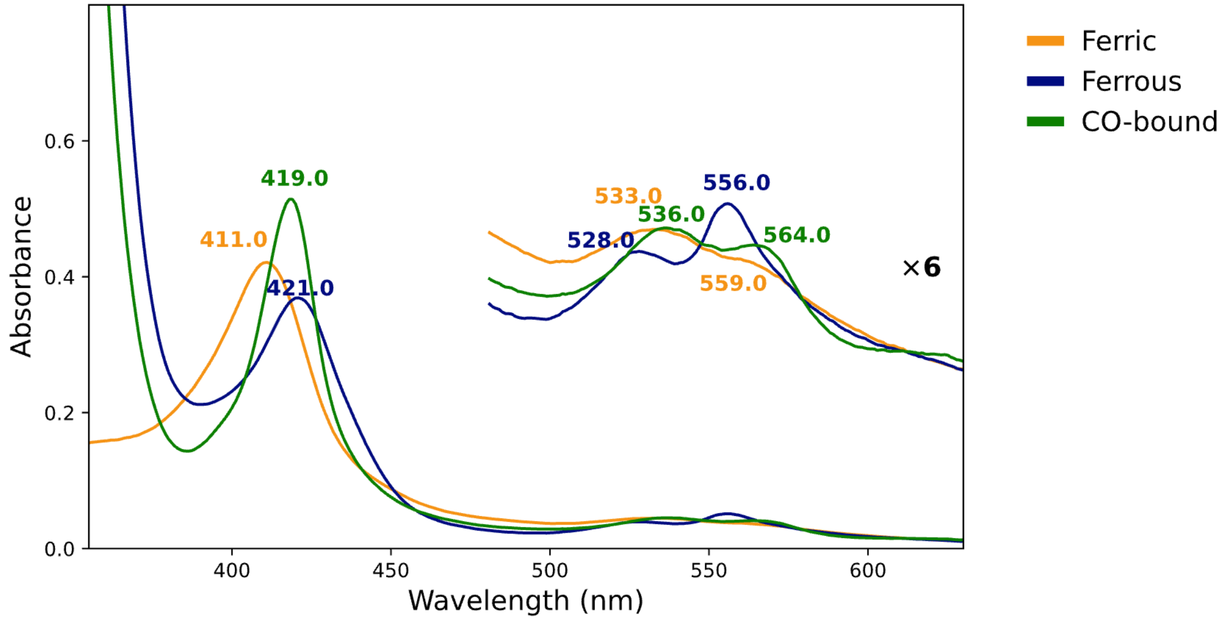

**Figure S9.** UV-vis spectrum of **GDM-NT-401**. Yellow: ferric state. Blue: ferrous state. Green: CO-bound ferrous state.

**Analytical-Scale Enzymatic Reaction Yield Analysis and Calibration Curves for Products**

All enzymatic reactions on analytical scale were conducted following the general procedure described in **General Information**. Reactions for each substrate were set up in triplicate. Product formation was quantified by LC-MS based on the calibration curve of the corresponding racemic standard products.

Calibration curves of reference compounds were conducted for the determination of yield. For each substrate, at least five different concentrations of product (0.125, 0.25, 0.5, 1.0, 1.5, 2, 2.5, 3, 4, and 5 mM) were prepared in 400 µL M9–N (pH = 7.2) buffer. This solution was quenched using the reaction work-up procedure described in **General Information** using 1,2,3-trimethylbenzene as internal standard. The samples were analyzed by HPLC based on UV absorbance at 254, 235, or 210 nm. The calibration curves depict the ratio of concentration in mM (y-axis) against product area to internal standard area (x-axis). *Notes*: Pdt = integration of product on UV spectroscopy, IS = integration of internal standard on UV spectroscopy. In cases where a set of reactions are all having <10% yields, a calibration curve using single ion channel ion count was used without referring to an internal standard.

Chiral protic HPLC-MS was used to determine the enantioselectivity of the reactions. Single ion channel MS spectroscopy was included to support the clean separation of the two enantiomers. For the analysis of regioselectivity, all measurements were made without considering the different response of regio isomers on HPLC-MS. Considering the ratio of the minor regioisomers is generally below 20%, this adjustment won't make large differences in the final regioselectivity.

**Part 1: *Tm*Pgb-PIP lineage analysis.**

All yields were calculated using a single ion channel calibration curve measured on the same day, using the same HPLC-MS, and with the same bottle of eluent. All yields within the ***Tm*Pgb-PIP** lineage using whole *E. coli* cell as catalysts are within the linear range of the calibration curve. All yields represent *only* the yields of **2a**.

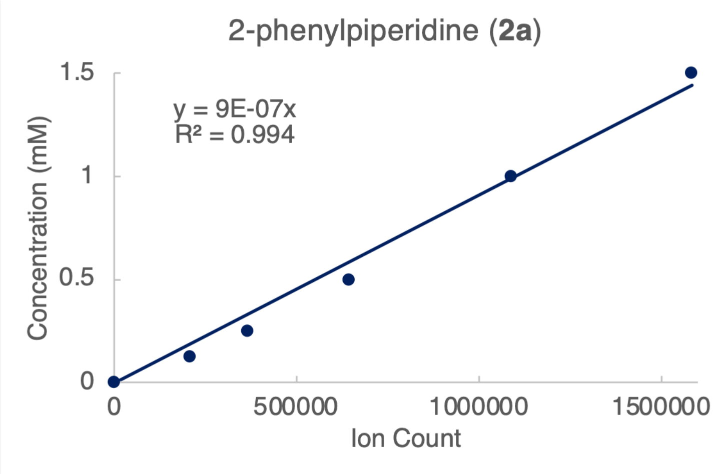

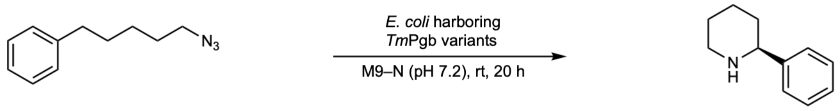

**1a** **2a**

***Tm*Pgb-wt** (whole *E.coli* cell with 11.5 µM heme loading)

| Entry | Product | Yield (%) | Average (%) | STDEV | Average TTN |
|---|---|---|---|---|---|
| 1 | 4156.1 | 0.08 | | | |
| 2 | 3717.6 | 0.07 | 0.07 | 0.01 | 0.3 |
| 3 | 4340.9 | 0.08 | | | |

***Tm*****Pgb-PIP-G1-5442** (whole *E.coli* cell with 11.9 μM heme loading)

| Entry | Product | Yield (%) | Average (%) | STDEV | Average TTN |
|---|---|---|---|---|---|
| 1 | 28315.9 | 0.52 | | | |
| 2 | 28773.1 | 0.52 | 0.54 | 0.03 | 2 |
| 3 | 31091.8 | 0.57 | | | |

***Tm*****Pgb-PIP-G2-5443** (whole *E.coli* cell with 11.4 μM heme loading)

| Entry | Product | Yield (%) | Average (%) | STDEV | Average TTN |
|---|---|---|---|---|---|
| 1 | 43809.0 | 0.80 | | | |
| 2 | 44714.0 | 0.81 | 0.80 | 0.02 | 3 |
| 3 | 42667.4 | 0.78 | | | |

***Tm*****Pgb-PIP-G3-5444** (whole *E.coli* cell with 9.3 μM heme loading)

| Entry | Product | Yield (%) | Average (%) | STDEV | Average TTN |
|---|---|---|---|---|---|
| 1 | 47379.5 | 0.86 | | | |
| 2 | 50912.8 | 0.93 | 0.91 | 0.04 | 5 |
| 3 | 51474.2 | 0.94 | | | |

***Tm*****Pgb-PIP-G4-5445** (whole *E.coli* cell with 11.7 μM heme loading)

| Entry | Product | Yield (%) | Average (%) | STDEV | Average TTN |
|---|---|---|---|---|---|
| 1 | 97787.6 | 1.78 | | | |
| 2 | 106253.1 | 1.94 | 1.84 | 0.08 | 8 |
| 3 | 99661.0 | 1.82 | | | |

***Tm*****Pgb-PIP-G5-5446** (whole *E.coli* cell with 10.9 μM heme loading)

| Entry | Product | Yield (%) | Average (%) | STDEV | Average TTN |
|---|---|---|---|---|---|
| 1 | 183923.9 | 3.35 | | | |
| 2 | 182699.1 | 3.33 | 3.38 | 0.06 | 15 |
| 3 | 189104.1 | 3.45 | | | |

***Tm*****Pgb-PIP-G6-5447** (whole *E.coli* cell with 10.7 μM heme loading)

| Entry | Product | Yield (%) | Average (%) | STDEV | Average TTN |
|---|---|---|---|---|---|
| 1 | 142822.3 | 2.60 | | | |
| 2 | 131523.9 | 2.40 | 2.48 | 0.11 | 12 |
| 3 | 134397.7 | 2.45 | | | |

***Tm*****Pgb-PIP-G7-5448** (whole *E.coli* cell with 10.2 μM heme loading)

| Entry | Product | Yield (%) | Average (%) | STDEV | Average TTN |
|---|---|---|---|---|---|
| 1 | 302443.6 | 5.51 | | | |
| 2 | 333834.6 | 6.08 | 5.97 | 0.41 | 29 |
| 3 | 346630.9 | 6.32 | | | |

**Part 2:** ***De novo*** **design with recycled scaffold data analysis.**

All yields were calculated using a single ion channel calibration curve measured on the same day, using the same HPLC-MS, and with the same bottle of eluent. All yields listed in this section were obtained using whole *E. coli* cell as catalysts and are within the linear range of the calibration curve. All yields represent *only* the yields of **2a**.

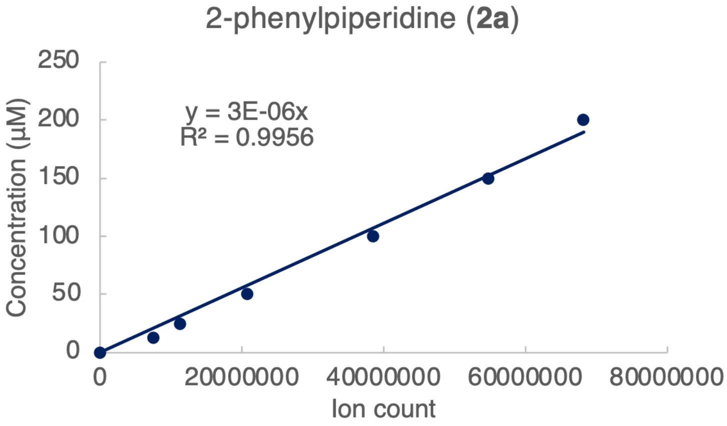

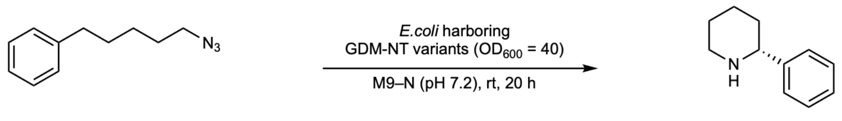

**Table S13**. Results using selected variants in recycled-scaffold design based on **GDM-NT-121**

| | Rxn 1 | Rxn 2 | Rxn 3 | Hemeloading (μM) | Average Yield (%) | Average TTN |
|---|---|---|---|---|---|---|
| ***Tm*****Pgb-wt** | 321052 | 203194 | 238715 | 6.5 | 0.01 | 0.1 |
| **GDM-NT-121** | 1616377 | 1595036 | 1623099 | 2.6 | 0.09 | 1.7 |
| **GDM-NT-215** | 3137075 | 3193042 | 3175384 | 1.7 | 0.18 | 5.2 |
| **GDM-NT-217** | 2094515 | 2021621 | 1987190 | 1.8 | 0.11 | 3.2 |
| **GDM-NT-226** | 28209976 | 27907585 | 28518283 | 2.8 | 1.58 | 28.2 |
| **GDM-NT-227** | 1647956 | 1518064 | 1659073 | 3.1 | 0.09 | 1.5 |
| **GDM-NT-228** | 480373 | 476117 | 483799 | 1.6 | 0.03 | 0.9 |

**Table S14**. Results using selected variants in recycled-scaffold design based on **GDM-NT-151**

| | Rxn 1 | Rxn 2 | Rxn 3 | Hemeloading (μM) | Average Yield (%) | Average TTN |
|---|---|---|---|---|---|---|
| ***Tm*****Pgb-wt** | 321052 | 203194 | 238715 | 6.5 | 0.01 | 0.1 |
| **GDM-NT-151** | 578534 | 578840 | 542094 | 1.7 | 0.03 | 0.9 |
| **GDM-NT-350** | 1795490 | 1722655 | 1735434 | 2.3 | 0.10 | 2.1 |
| **GDM-NT-356** | 5384664 | 5580567 | 6000699 | 2.7 | 0.32 | 5.8 |
| **GDM-NT-359** | 3940367 | 4047304 | 4345490 | 3.6 | 0.23 | 3.2 |
| **GDM-NT-361** | 1321241 | 1475285 | 1588278 | 1.7 | 0.08 | 2.5 |
| **GDM-NT-362** | 8486036 | 8421601 | 8587747 | 13.8 | 0.47 | 1.7 |

**Part 3:** Evolution from **GDM-NT-226** and **GDM-NT-362.**

After identifying GDM-NT-226 **and GDM-NT-362** as the starting point for further evolution, we performed one round of random mutagenesis using error-prone PCR and identified multiple variants with beneficial mutations.

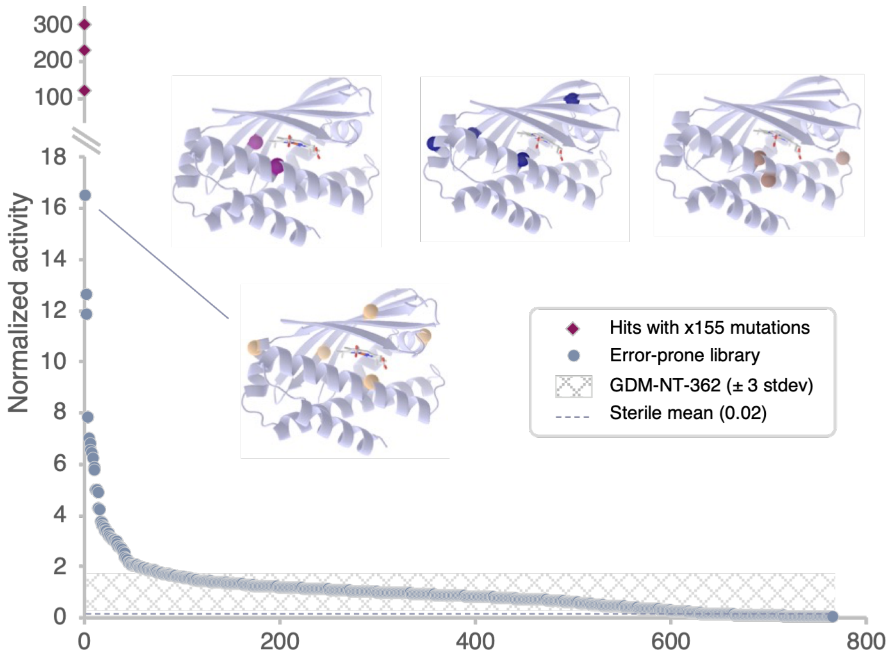

**Figure S10**. Retention of function curve for directed evolution using **GDM-NT-362** as parent. Three variants demonstrated higher than 100-fold improvement compared to **GDM-NT-362**. The illustration of these three variants with mutations highlighted in purple, blue, and brown were generated by AlphaFold 3. Note that all three variants have a mutation at a shared site 155. Three additional variants demonstrated 10- to 20-fold improvement compared to **GDM-NT-362**. The top hit with mutations highlighted in yellow was generated by AlphaFold 3. The area highlighted in cross pattern represents results of **GDM-NT-362** within ±3×standard deviation. The sterile mean of the reaction with respect to **GDM-NT-362** is high in dash line at the bottom of the figure.

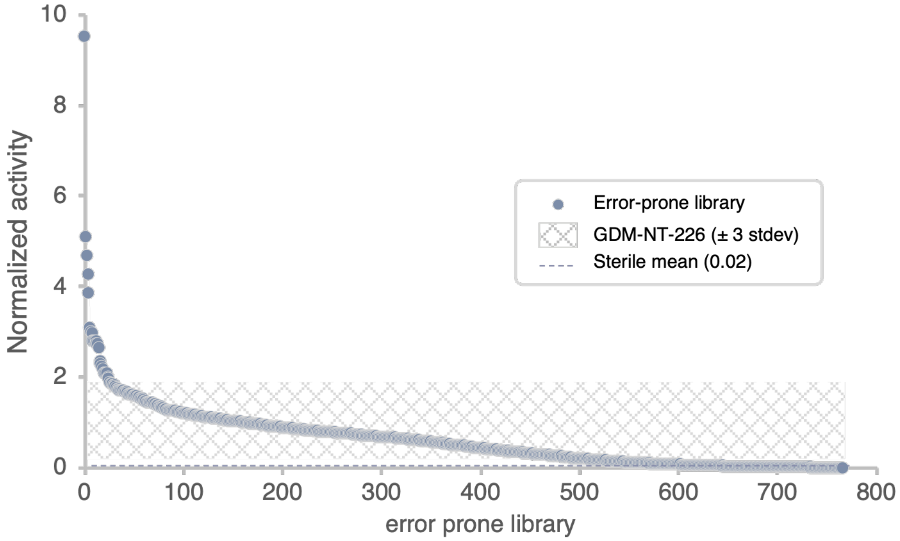

**Figure S11**. Retention of function curve for directed evolution using **GDM-NT-226** as parent. Four variants demonstrated >4-fold improvement compared to **GDM-NT-226**. The area highlighted in cross pattern represents results of **GDM-NT-226** within ±3×standard deviation. The sterile mean of the reaction with respect to **GDM-NT-226** is high in dash line at the bottom of the figure.

**Part 4:** Additional engineering from top hit in error-prone library

One variant with four mutations V53L, 68V, E99G, H155L, later designated as **GDM-NT-399**, was used as the parent for further protein engineering. We identified site 155 as the designed site for axial ligand of the heme co-factor. As a result, we focused on mutations at site 155 and adjacent site 156, while keeping the remaining three mutations at site 53, 68, and 99. We later determined that these three mutations (V53L, 68V, E99G) introduced through error-prone PCR were beneficial to the reaction yield (comparing **GDM-NT-401** with **GDM-NT-400**). All yields represent only the yields of **2a**.

After this final round of protein engineering, the top-performing variant **GDM-NT-401** was selected as final variant for substrate scope study.

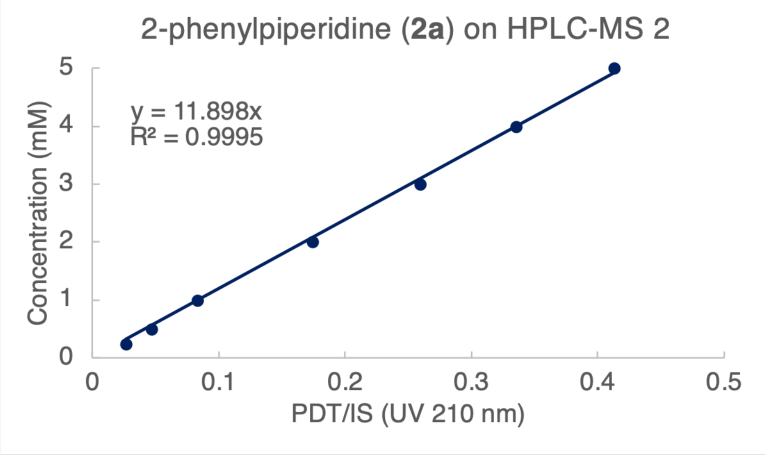

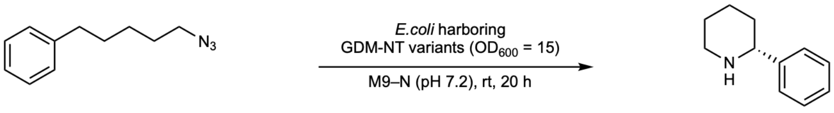

**1a** **2a**

**Table S15**. Results of final round of protein engineering. Analytical-scale reactions were performed on a 96-well plate, with an averaged $OD_{600}$ of 15.

| | x155 | Mutations from GDM-NT-362 | PDT | IS | Yield (%) |
|---|---|---|---|---|---|
| **GDM-NT-362** | H | - | NA | NA | <1 |
| **GDM-NT-404** | H | V53L-I68V-E99G | NA | NA | <1 |
| **GDM-NT-405** | H | V53L-I68V-E99G-M156A | NA | NA | <1 |
| **GDM-NT-399** | L | V53L-I68V-E99G-H155L | 444.1 | 5706.2 | 18.5 |
| **GDM-NT-406** | L | V53L-I68V-E99G-H155L-M156L | 744.6 | 5673.7 | 31.2 |
| **GDM-NT-403** | L | V53L-I68V-E99G-H155L-M156A | 781.8 | 5781.5 | 32.2 |
| **GDM-NT-407** | M | V53L-I68V-E99G-H155M | 127.7 | 5632.3 | 5.4 |
| **GDM-NT-401** | M | V53L-I68V-E99G-H155M-M156L | 872.1 | 5744.3 | 36.1 |
| **GDM-NT-402** | M | V53L-I68V-E99G-H155M-M156A | 565.6 | 5810.3 | 23.2 |
| **GDM-NT-400** | M | H155M-M156L | 630.5 | 5761.5 | 26.0 |

**Part 5: Reaction condition optimization using purified protein.**

Despite reactions performed using *e.coli* harboring ***Tm*Pgb-PIP** variants and **GDM-NT** variants do not require the addition of external reductant; reactions performed with purified protein exhibited diverging behavior between final variants ***Tm*Pgb-PIP-G7** and **GDM-NT-401**. While no external reductant is required for reactions performed with purified ***Tm*Pgb-PIP-G7**, a catalytic amount of $Na_2S_2O_4$ was required for reactions performed with **GDM-NT-401**. Additionally, this trend was consistent with a previously reported variant ***Tm*Pgb-PYR-G2-5441**, where no external reductant was required.

An aliquot of ***Tm*Pgb-PYR-G2-5441**, ***Tm*Pgb-PIP-G7**, or **GDM-NT-401** were added to a 2-mL reaction vial and adjusted to 10 µM *final* heme concentration with M9–N (pH 7.2) buffer. The reaction vials were transferred to a coy chamber. Subsequently, different amount of a freshly prepared stock solution of $Na_2S_2O_4$ (100 mM in M9–N buffer) was added to the vial and gently mixed by shaking the vial holder for 10 seconds. Final concentrations of $Na_2S_2O_4$ added to the reactions vary from 0 mM, 1 mM, 2.5 mM, to 5 mM. Then, 20 µL of a stock solution of **1a** (100 mM in ethanol) was added to the reaction mixture. The reaction vial was capped and shaken for 20 h at 700 rpm and room temperature.

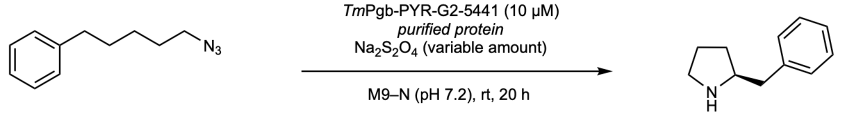

**1a** (5.0 mM) **3a**

**Table S16**. Effect of external reductant for ***Tm*Pgb-PYR-G2-5441** (purified enzyme) on UV channel.

| $Na_2S_2O_4$ (mM) | PDT | IS | Yield (%) | Average Yield (%) | STDEV |
|---|---|---|---|---|---|
| 0 | 742.2 | 5481.3 | 26.7 | | |
| 0 | 721.2 | 5322.4 | 26.7 | 27 | 1.09 |
| 0 | 782.7 | 5397.4 | 28.6 | | |
| 1 | 251.8 | 5279.5 | 9.4 | | |
| 1 | 263.6 | 5286.9 | 9.8 | 10 | 0.79 |
| 1 | 301.3 | 5427.2 | 10.9 | | |
| 2.5 | 32.0 | 5230.0 | 1.2 | | |
| 2.5 | 40.3 | 5259.3 | 1.5 | 1.2 | 0.37 |
| 2.5 | 20.8 | 5340.2 | 0.8 | | |
| 5 | 17.4 | 5310.3 | 0.6 | | |
| 5 | 18.5 | 5345.3 | 0.7 | 0.6 | 0.19 |
| 5 | 8.7 | 5133.0 | 0.3 | | |

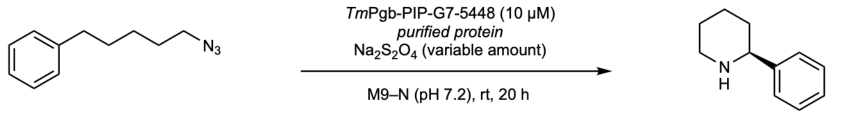

**1a** (5.0 mM) **2a**

**Table S17**. Effect of external reductant for ***Tm*Pgb-PIP-G7-5448** (purified enzyme) on single ion mass channel.

| $Na_2S_2O_4$ (mM) | PDT ion count Rxn 1 | PDT ion count Rxn 2 | PDT ion count Rxn 3 | PDT ion count average | Normalized Yield | STDEV |
|---|---|---|---|---|---|---|
| 0 | 240283.2 | 276580.7 | 252683.0 | 256515.6 | 1.00 | 18449.8 |

| | | | | | | |
|---|---|---|---|---|---|---|
| 1 | 163707.1 | 167370.2 | 205207.6 | 178761.6 | 0.70 | 22976.0 |
| 2.5 | 32425.2 | 26376.1 | 36414.4 | 31738.6 | 0.12 | 5054.3 |
| 5 | 20368.7 | 21099.1 | 20030.4 | 20499.4 | 0.08 | 546.2 |

**Table S18**. Effect of external reductant for ***Tm*Pgb-PIP-G7-5448** (purified enzyme) on UV channel using 0 mM and 1 mM $Na_2S_2O_4$ as external reductant. This result agrees with **Table S15**. Due to low UV intensity, results using 2.5 mM and 5 mM $Na_2S_2O_4$ as external reductant are not available.

| $Na_2S_2O_4$ (mM) | PDT | IS | Yield (%) | Average Yield (%) | STDEV |
|---|---|---|---|---|---|
| 0 | 128.8 | 5367.0 | 4.7 | | |
| 0 | 150.3 | 5385.1 | 5.5 | 5.1 | 0.39 |
| 0 | 144.5 | 5523.5 | 5.2 | | |
| 1 | 83.6 | 5379.3 | 3.1 | | |
| 1 | 82.8 | 5368.0 | 3.0 | 3.2 | 0.26 |
| 1 | 96.4 | 5422.0 | 3.5 | | |

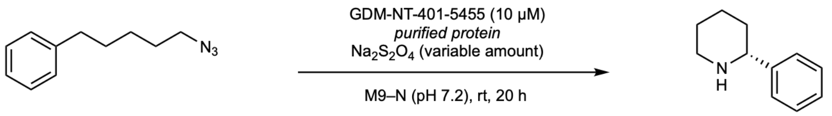

**1a** (5.0 mM) **2a**

**Table S19**. Effect of external reductant for **GDM-NT-401-5455** (purified enzyme) on UV channel.

| $Na_2S_2O_4$ (mM) | PDT | IS | Yield (%) | Average Yield (%) | STDEV |
|---|---|---|---|---|---|
| 0 | 139.4 | 4539.0 | 6.0 | | |
| 0 | 137.1 | 4701.8 | 5.7 | 6 | 0.21 |
| 0 | 136.0 | 4749.3 | 5.6 | | |
| 1 | 1537.7 | 4594.7 | 65.9 | | |
| 1 | 1384.1 | 4621.3 | 59.0 | 60 | 6.03 |
| 1 | 1247.8 | 4558.5 | 53.9 | | |
| 2.5 | 1840.4 | 4517.9 | 80.2 | | |
| 2.5 | 1843.8 | 4619.3 | 78.6 | 78 | 2.32 |
| 2.5 | 1791.3 | 4663.5 | 75.7 | | |
| 5 | 1277.3 | 4548.9 | 55.3 | | |
| 5 | 1318.3 | 4483.2 | 57.9 | 59 | 4.02 |
| 5 | 1492.5 | 4652.5 | 63.2 | | |

**Part 6: Solvent stability test for GDM-NT-401 under high concentration of ethanol.**

Solvent stability test for **GDM-NT-401** was performed by conducting the reaction under different volume percentage of ethanol as co-solvent. An aliquot of **GDM-NT-401** in KPi buffer (pH 8.0) were diluted with corresponding amounts (400μL – 400 μL × ethanol% –17μL enzyme solution – 10 μL $Na_2S_2O_4$ solution) of M9–N buffer (pH 7.2). The solution was brought to a coy chamber. Subsequently, different amounts of ethanol were added (400 μL × ethanol% – 20 μL substrate **1a** solution). We noticed that at 80% or higher ethanol loading, the solution became turbid, suggesting protein denaturation. Then, 10 μL of a freshly prepared stock solution of $Na_2S_2O_4$ (100 mM in M9–N buffer) was added to the vial and gently mixed by shaking the vial holder for 10 seconds. Finally, 20 μL of a stock solution of **1a** (100 mM in ethanol) was added to the reaction mixture. The reaction vial was capped and shaken for 20 h at 700 rpm and room temperature.

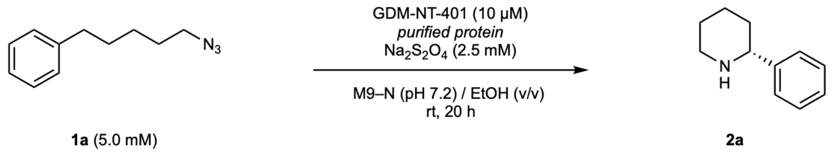

**Table S20**. Results of solvent stability test for **GDM-NT-401** under high concentration of ethanol. Reactions were run in duplicate.

| | PDT Rxn 1 | IS Rxn 1 | PDT Rxn 2 | IS Rxn 2 | Average Yield (%) | STDEV |
|---|---|---|---|---|---|---|
| **5% EtOH** | 2124.8 | 5612.0 | 2081.0 | 5822.8 | 72 | 3.0 |
| **20% EtOH** | 1920.5 | 5466.5 | 1856.6 | 5421.6 | 68 | 1.2 |
| **40% EtOH** | 1107.6 | 5538.9 | 1010.3 | 5676.8 | 37 | 3.1 |
| **60% EtOH** | 1402.8 | 5537.4 | 1136.0 | 5422.0 | 46 | 6.1 |
| **80% EtOH** | 657.9 | 5449.4 | 666.3 | 5572.2 | 24 | 0.2 |
| **90% EtOH** | 109.2 | 5496.9 | 108.4 | 5600.0 | 4 | 0.1 |

**Part 7: $T_{50}$ measurement for GDM-NT-401 and *Tm*Pgb-PIP-G7**

Aliquots of the stock solution of **GDM-NT-401** or ***Tm*Pgb-PIP-G7** were added to PCR tubes and diluted with M9–N (pH 7.2) buffer to 150 μL. The PCR tubes were incubated at different temperatures using a thermocycler for 1 h and cooled down to room temperature for 1 h. The solution was transferred to reaction vials and topped up to 370 μL with M9–N (pH 7.2) buffer. The solution was brought to a coy chamber. Subsequently, 10 μL of a freshly prepared stock solution of $Na_2S_2O_4$ (100 mM in M9–N buffer) was added to the vial and gently mixed by shaking the vial holder for 10 seconds. Then, 20 μL of a stock solution of **1a** (100 mM in ethanol) was added to the reaction mixture. The reaction vial was capped and shaken for 20 h at 700 rpm and room temperature.

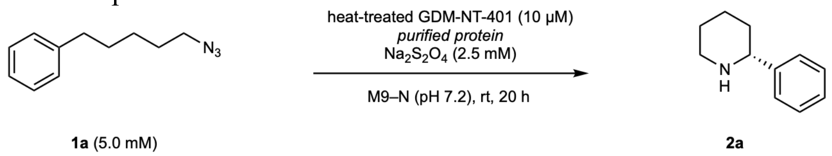

**Table S21**. $T_{50}$ measurement for **GDM-NT-401**

| Temp. (°C) | PDT Rxn 1 | IS Rxn 1 | PDT Rxn 2 | IS Rxn 2 | Average Yield (%) | Normalized Yield | STDEV |
|---|---|---|---|---|---|---|---|
| 4 | 1497.2 | 4937.1 | 1563.6 | 4804.8 | 62 | 1.00 | 0.050 |
| 29.8 | 1628.7 | 4910.7 | 1596.3 | 4824.4 | 65 | 1.05 | 0.002 |
| 35.1 | 1488.5 | 4778.9 | 1550.4 | 4851.3 | 62 | 1.00 | 0.018 |
| 39.2 | 1578.4 | 4892.5 | 1594.7 | 4852.9 | 64 | 1.04 | 0.013 |
| 46.9 | 1601.6 | 4866.7 | 1600.6 | 4854.6 | 65 | 1.05 | 0.001 |
| 54.7 | 1496.0 | 4828.1 | 1541.7 | 4767.6 | 62 | 1.01 | 0.030 |
| 59.6 | 1714.3 | 4804.4 | 1664.3 | 4637.5 | 70 | 1.14 | 0.005 |
| 60.4 | 1620.6 | 4872.4 | 1636.0 | 4884.1 | 66 | 1.06 | 0.005 |
| 65 | 1704.3 | 4865.6 | 1730.8 | 4869.0 | 70 | 1.12 | 0.012 |
| 76.2 | 1693.8 | 4872.7 | 1742.8 | 4936.9 | 69 | 1.11 | 0.012 |

| | | | | | | | |
|---|---|---|---|---|---|---|---|
| 81.4 | 1675.5 | 4850.5 | 1727.1 | 4876.4 | 69 | 1.11 | 0.020 |
| 85 | 1665.7 | 4840.6 | 1690.2 | 4781.4 | 69 | 1.11 | 0.021 |

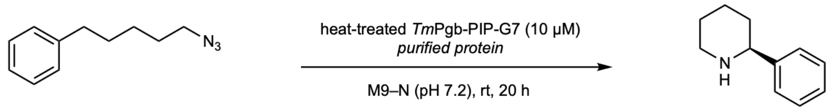

**1a** (5.0 mM) **2a**

**Table S22**. $T_{50}$ measurement for ***Tm*Pgb-PIP-G7**

| Temp. (°C) | PDT ion count Rxn 1 | PDT ion count Rxn 2 | PDT ion count average | Normalized Yield | STDEV |
|---|---|---|---|---|---|
| 4 | 424219.2 | 481448.2 | 452833.7 | 1.00 | 0.089 |
| 29.8 | 380428.5 | 427653.5 | 404041.0 | 0.89 | 0.074 |
| 32.2 | 374280.8 | 445751.5 | 410016.2 | 0.91 | 0.112 |
| 39.2 | 393659.6 | 406688.8 | 400174.2 | 0.88 | 0.020 |
| 46.9 | 389157.2 | 388710.7 | 388933.9 | 0.86 | 0.001 |
| 54.7 | 378291.2 | 390006.4 | 384148.8 | 0.85 | 0.018 |
| 57.7 | 388774.5 | 436880.8 | 412827.6 | 0.91 | 0.075 |
| 59.6 | 383348.1 | 391706.5 | 387527.3 | 0.86 | 0.013 |
| 60.4 | 338108.4 | 390896.8 | 364502.6 | 0.80 | 0.082 |
| 65 | 168664.4 | 184524.0 | 176594.2 | 0.39 | 0.025 |
| 66.7 | 141247.3 | 153473.6 | 147360.5 | 0.33 | 0.019 |
| 68.6 | 51971.4 | 50511.6 | 51241.5 | 0.11 | 0.002 |
| 71.2 | 6431.4 | 7807.4 | 7119.4 | 0.02 | 0.002 |
| 76.2 | trace | trace | trace | trace | NA |
| 81.4 | trace | trace | trace | trace | NA |
| 85 | trace | trace | trace | trace | NA |

**Part 8: Third round of design using refined motifs.**

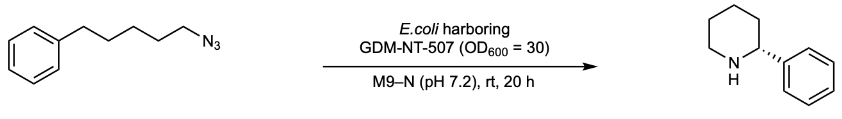

**1a** **2a**

Reaction performed **with GDM-NT-507** (whole *E.coli* cell with 1.8 μM heme loading)

| Entry | Product | IS | P/IS | Yield (%) | Average (%) | STDEV | Average TTN |
|---|---|---|---|---|---|---|---|
| 1 | 483.3 | 5527.7 | 0.087 | 17.2 | | | |
| 2 | 522.0 | 5829.8 | 0.090 | 17.6 | 17.0 | 0.8 | 470 |
| 3 | 471.9 | 5808.5 | 0.081 | 16.0 | | | |

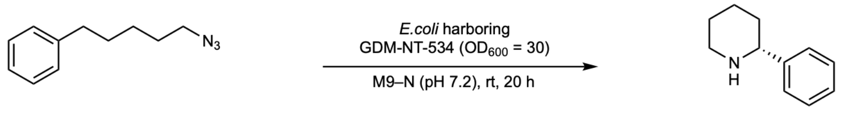

**1a** **2a**

Reaction performed **with GDM-NT-534** (whole *E.coli* cell with 1.8 μM heme loading)

| Entry | Product | IS | P/IS | Yield (%) | Average (%) | STDEV | Average TTN |
|---|---|---|---|---|---|---|---|
| 1 | 392.2 | 5679.2 | 0.069 | 13.6 | | | |
| 2 | 523.6 | 5803.6 | 0.090 | 17.8 | 15.5 | 2.1 | 420 |
| 3 | 448.9 | 5871.5 | 0.076 | 15.1 | | | |

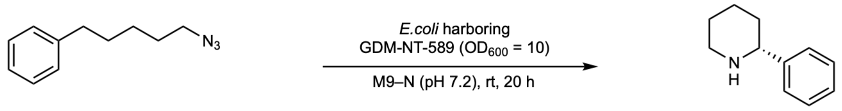

**1a** **2a**

Reaction performed **with GDM-NT-589** (whole *E.coli* cell with $OD_{600}$ = 10)
This variant was only able to express at low cell density likely due to relatively larger protein size.

| Entry | Product | IS | P/IS | Yield (%) | Average (%) | STDEV | Average TTN |
|---|---|---|---|---|---|---|---|
| 1 | 499.2 | 5201.1 | 0.096 | 18.9 | | | |
| 2 | 428.9 | 5267.5 | 0.081 | 16.0 | 17.2 | 1.4 | N.A. |
| 3 | 463.4 | 5141.6 | 0.090 | 17.8 | | | |
| 4 | 442.6 | 5388.1 | 0.082 | 16.2 | | | |

**Part 9: Substrate scope analysis.**
Analytical reaction yield obtained with *E.coli* harboring **GDM-NT-401** ($OD_{600}$ = 30, 2.8 μM heme loading) or purified ***Tm*Pgb-PIP-G7-5448** (20 μM heme loading). All yields were calculated using a calibration curve measured on UV spectrum (210 nm or 254 nm) with 1,2,3-trimethoxybenzene as internal standard. Reactions with <1% yields were omitted.

**Calibration curve and data analysis for 2a and 2b**
**2a** and **2b** were measured using the same calibration curve.

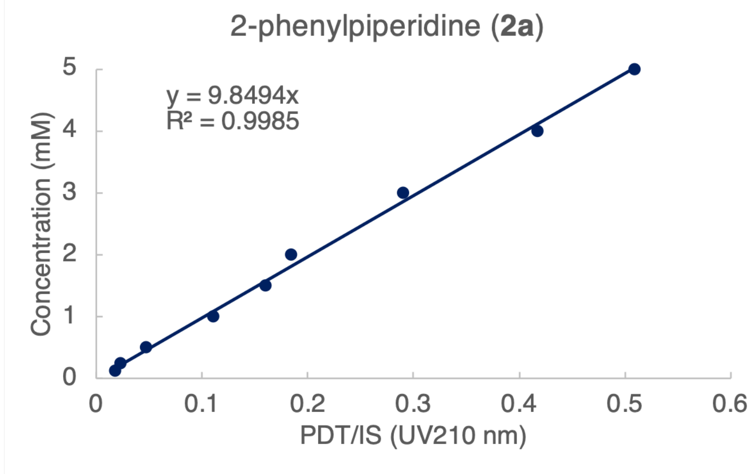

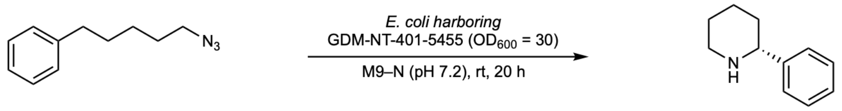

**1a** **2a**

Reaction performed **with GDM-NT-401-5455** (whole *E.coli* cell with 2.8 μM heme loading)

| Entry | Product | IS | P/IS | Yield (%) | Average (%) | STDEV | Average TTN |
|---|---|---|---|---|---|---|---|
| 1 | 1724.8 | 4824.8 | 0.36 | 70.4 | | | |
| 2 | 1564.7 | 4682.5 | 0.33 | 65.8 | 66 | 3.7 | 1200 |
| 3 | 1533.3 | 4781.9 | 0.32 | 63.1 | | | |

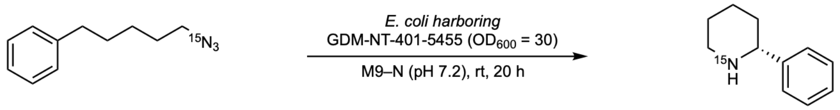

**1a** **2b**

Reaction performed **with GDM-NT-401-5455** (whole *E.coli* cell with 2.8 μM heme loading)

| Entry | Product | IS | P/IS | Yield (%) | Average (%) | STDEV | Average TTN |
|---|---|---|---|---|---|---|---|
| 1 | 1620.5 | 4695.6 | 0.35 | 68.0 | | | |
| 2 | 1348.2 | 4716.3 | 0.29 | 56.3 | 59 | 7.9 | 1100 |
| 3 | 1255.1 | 4651.3 | 0.27 | 53.2 | | | |

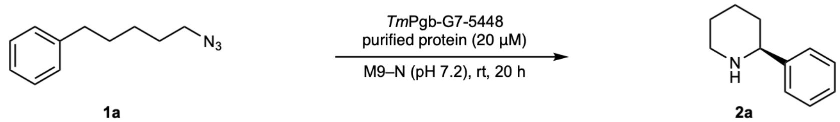

**1a** **2a**

Reaction performed **with *Tm*Pgb-G7-5448** (purified protein with 20 μM heme loading)

| Entry | Product | IS | P/IS | Yield (%) | Average (%) | STDEV | Average TTN |
|---|---|---|---|---|---|---|---|
| 1 | 303.6 | 4736.1 | 0.06 | 12.6 | | | |
| 2 | 276.4 | 4622.1 | 0.06 | 11.8 | 12 | 0.4 | 31 |
| 3 | 288.4 | 4568.7 | 0.06 | 12.4 | | | |

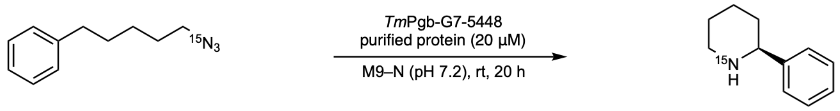

**1a** **2b**

Reaction performed **with *Tm*Pgb-G7-5448** (purified protein with 20 μM heme loading)

| Entry | Product | IS | P/IS | Yield (%) | Average (%) | STDEV | Average TTN |
|---|---|---|---|---|---|---|---|
| 1 | 295.8 | 6145.3 | 0.05 | 9.5 | | | |
| 2 | 337.6 | 6143.7 | 0.05 | 10.8 | 11 | 1.1 | 27 |
| 3 | 368.8 | 6202.4 | 0.06 | 11.7 | | | |

**Calibration curve and data analysis for 2c**

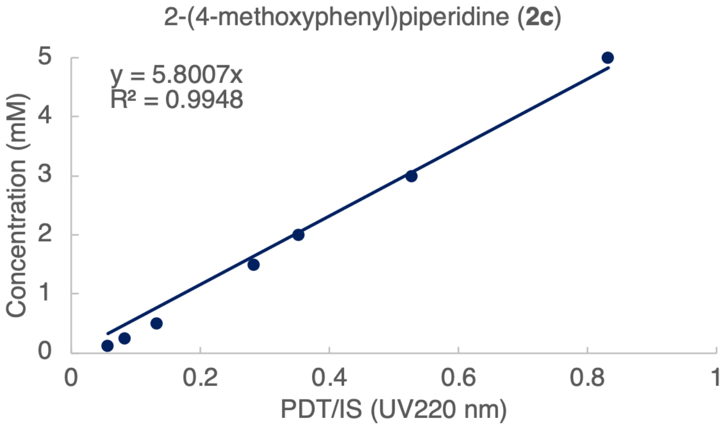

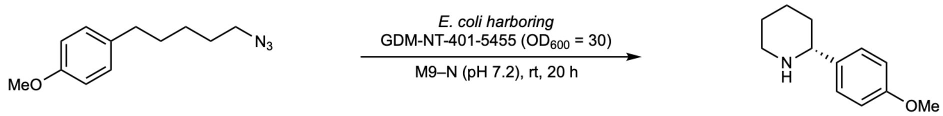

Reaction performed **with GDM-NT-401-5455** (whole *E.coli* cell with 2.8 μM heme loading)

| Entry | Product | IS | P/IS | Yield (%) | Average (%) | STDEV | Average TTN |
|---|---|---|---|---|---|---|---|
| 1 | 201.7 | 2602.4 | 0.08 | 9.0 | | | |
| 2 | 214.0 | 2732.6 | 0.08 | 9.1 | 9 | 0.6 | 168 |
| 3 | 235.1 | 2687.4 | 0.09 | 10.2 | | | |

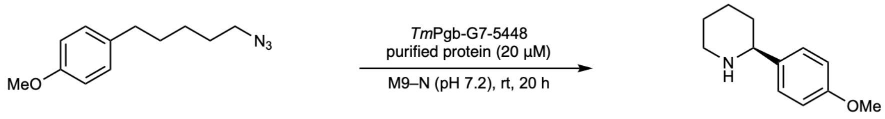

**1c** **2c**, <1% yield

Reaction performed **with *Tm*Pgb-G7-5448** (purified protein with 20 μM heme loading) furnished **2c** in <1% yield.

**Calibration curve and data analysis for 2d**

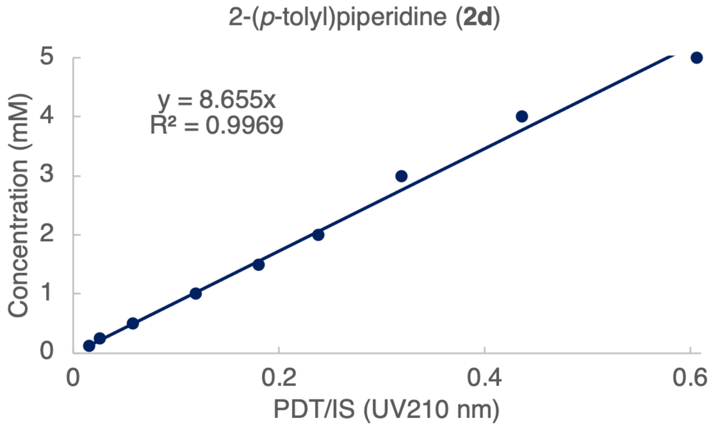

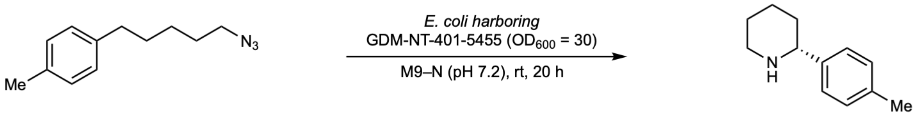

Reaction performed **with GDM-NT-401-5455** (whole *E.coli* cell with 2.8 μM heme loading)

| Entry | Product | IS | P/IS | Yield (%) | Average (%) | STDEV | Average TTN |
|---|---|---|---|---|---|---|---|
| 1 | 1430.3 | 4422.4 | 0.32 | 56.0 | | | |
| 2 | 1291.1 | 4407.5 | 0.29 | 50.7 | 52 | 3.8 | 926 |
| 3 | 1253.6 | 4446.2 | 0.28 | 48.8 | | | |

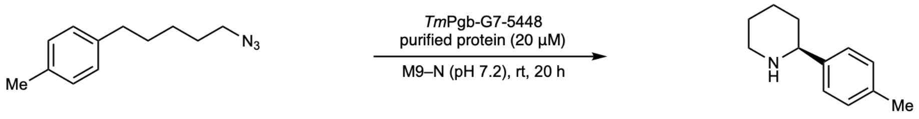

Reaction performed **with *Tm*Pgb-G7-5448** (purified protein)

| Entry | Product | IS | P/IS | Yield (%) | Average (%) | STDEV | Average TTN |
|---|---|---|---|---|---|---|---|
| 1 | 214.6 | 4291.7 | 0.05 | 8.7 | | | |
| 2 | 214.2 | 4459.5 | 0.05 | 8.3 | 9 | 0.2 | 21 |
| 3 | 215.9 | 4307.3 | 0.05 | 8.7 | | | |

**Calibration curve and data analysis for 3e**

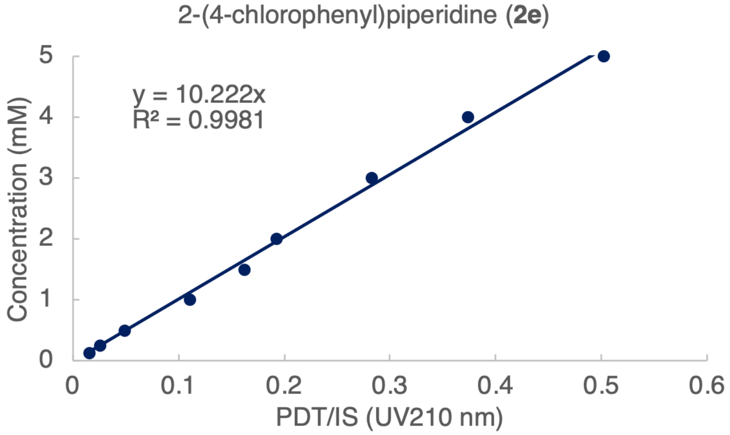

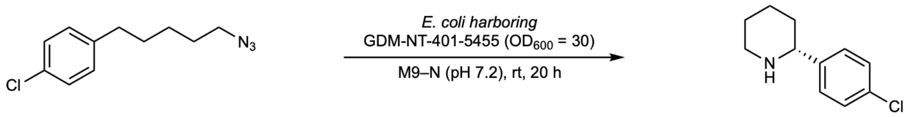

Reaction performed **with GDM-NT-401-5455** (whole *E.coli* cell with 2.8 µM heme loading)

| Entry | Product | IS | P/IS | Yield (%) | Average (%) | STDEV | Average TTN |
|---|---|---|---|---|---|---|---|
| 1 | 128.9 | 4697.0 | 0.03 | 5.6 | | | |
| 2 | 129.2 | 4743.7 | 0.03 | 5.6 | 5 | 0.6 | 94 |
| 3 | 105.5 | 4756.9 | 0.02 | 4.5 | | | |

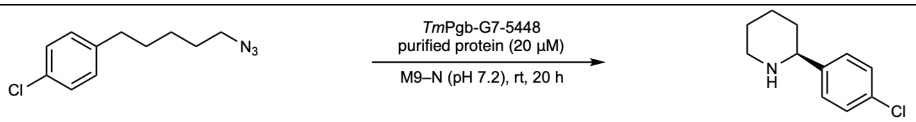

Reaction performed **with *Tm*Pgb-G7-5448** (purified protein)

| Entry | Product | IS | P/IS | Yield (%) | Average (%) | STDEV | Average TTN |
|---|---|---|---|---|---|---|---|
| 1 | 62.8 | 5797.3 | 0.011 | 2.2 | | | |
| 2 | 40.1 | 5726.6 | 0.007 | 1.4 | 2 | 0.5 | 4 |
| 3 | 37.1 | 5924.2 | 0.006 | 1.3 | | | |

**Calibration curve and data analysis for 3f**

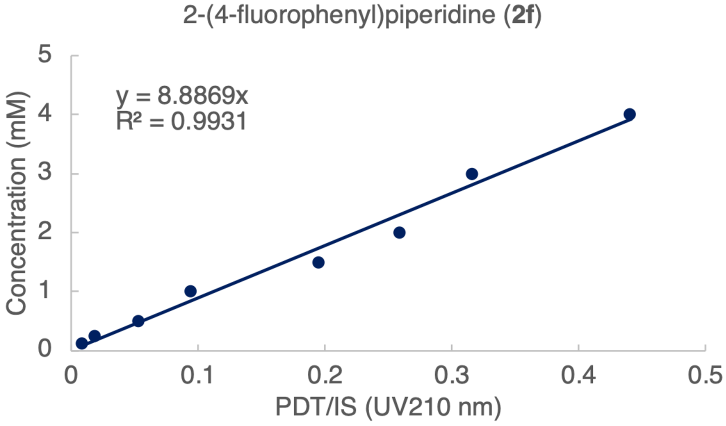

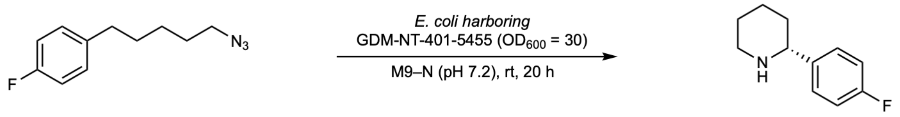

Reaction performed **with GDM-NT-401-5455** (whole *E.coli* cell with 2.8 µM heme loading)

| Entry | Product | IS | P/IS | Yield (%) | Average (%) | STDEV | Average TTN |
|---|---|---|---|---|---|---|---|
| 1 | 956.8 | 4743.6 | 0.20 | 35.9 | | | |
| 2 | 944.7 | 4760.1 | 0.20 | 35.3 | 34 | 2.1 | 600 |
| 3 | 844.2 | 4680.3 | 0.18 | 32.1 | | | |

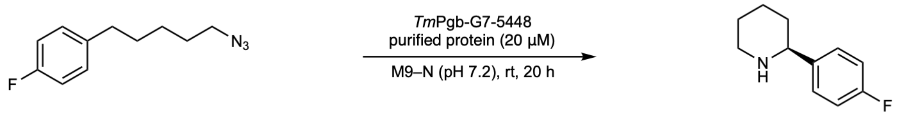

Reaction performed **with *Tm*Pgb-G7-5448** (purified protein)

| Entry | Product | IS | P/IS | Yield (%) | Average (%) | STDEV | Average TTN |
|---|---|---|---|---|---|---|---|
| 1 | 66.5 | 4774.2 | 0.014 | 2.5 | | | |
| 2 | 59.8 | 4471.9 | 0.013 | 2.4 | 3 | 0.4 | 7 |
| 3 | 80.5 | 4583.2 | 0.018 | 3.1 | | | |

**Calibration curve and data analysis for 2g**

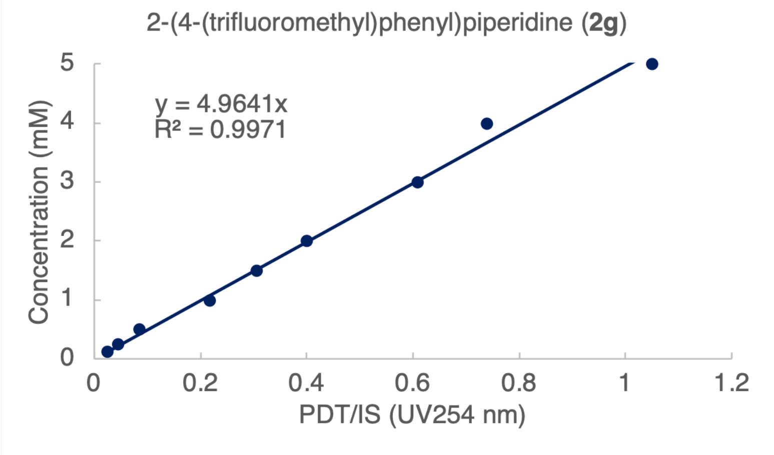

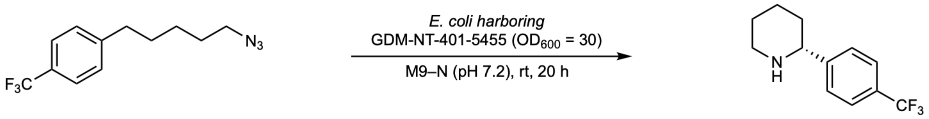

Reaction performed **with GDM-NT-401-5455** (whole *E.coli* cell with 2.8 μM heme loading)

| Entry | Product | IS | P/IS | Yield (%) | Average (%) | STDEV | Average TTN |
|---|---|---|---|---|---|---|---|
| 1 | 36.1 | 111.9 | 0.32 | 32.1 | | | |
| 2 | 34.0 | 118.1 | 0.29 | 28.6 | 30 | 1.7 | 539 |
| 3 | 35.4 | 117.8 | 0.30 | 29.8 | | | |

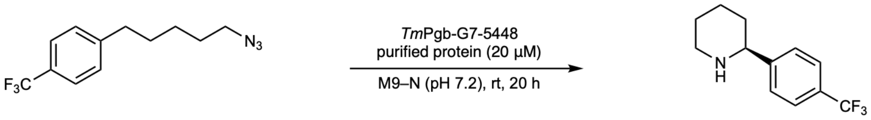

**1g** **2g**, <1% yield

Reaction performed **with *Tm*Pgb-G7-5448** (purified protein with 20 μM heme loading) furnished **2g** in <1% yield.

**Calibration curve and data analysis for 2h**

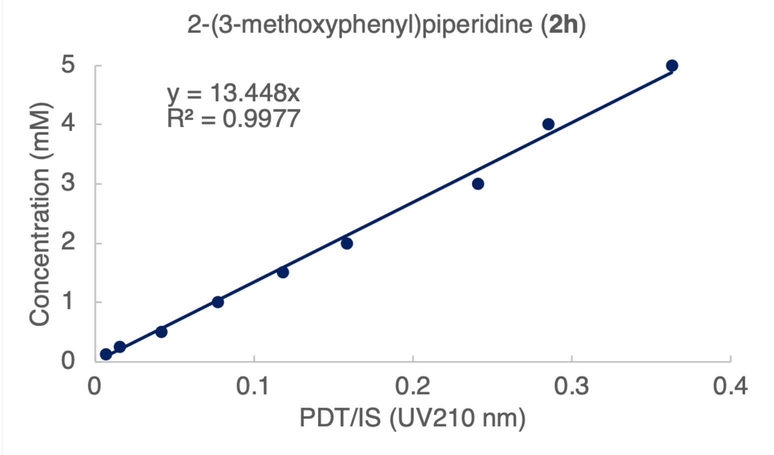

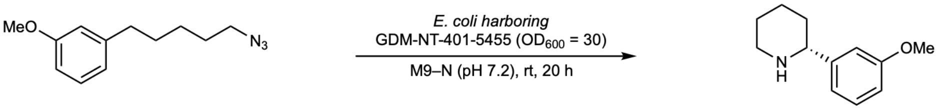

Reaction performed **with GDM-NT-401-5455** (whole *E.coli* cell with 2.8 µM heme loading)

| Entry | Product | IS | P/IS | Yield (%) | Average (%) | STDEV | Average TTN |
|---|---|---|---|---|---|---|---|
| 1 | 786.6 | 4754.6 | 0.17 | 44.5 | | | |
| 2 | 760.0 | 4680.6 | 0.16 | 43.7 | 46 | 3.3 | 800 |
| 3 | 883.0 | 4783.8 | 0.18 | 49.6 | | | |

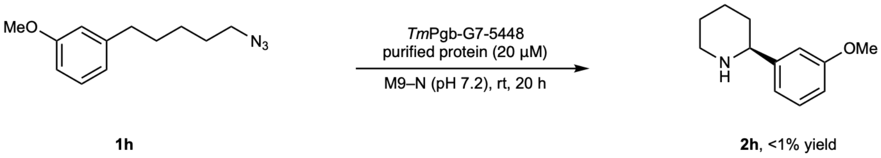

Reaction performed **with *Tm*Pgb-G7-5448** (purified protein with 20 µM heme loading) furnished **2h** in <1% yield.

**Calibration curve and data analysis for 2i**

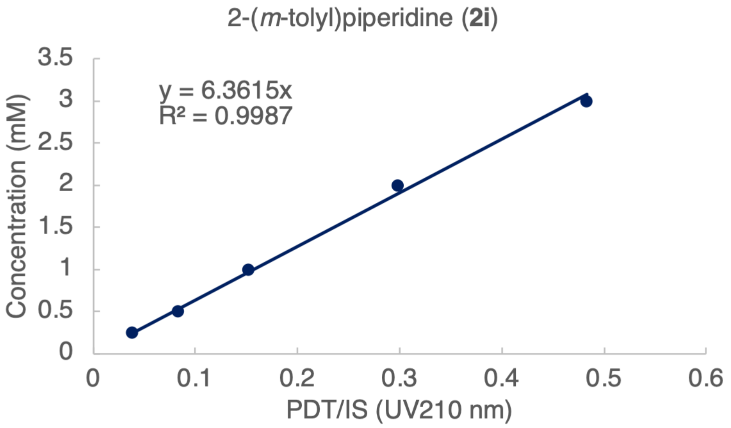

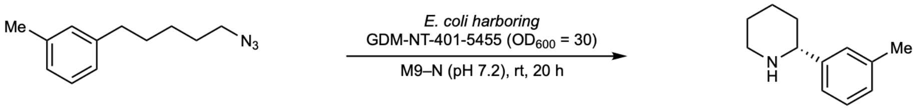

Reaction performed **with GDM-NT-401-5455** (whole *E.coli* cell with 2.8 µM heme loading)

| Entry | Product | IS | P/IS | Yield (%) | Average (%) | STDEV | Average TTN |
|---|---|---|---|---|---|---|---|
| 1 | 1162.5 | 4558.7 | 0.26 | 32.4 | | | |
| 2 | 1185.2 | 4345.1 | 0.27 | 34.7 | 33 | 1.3 | 594 |
| 3 | 1143.0 | 4451.1 | 0.26 | 32.7 | | | |

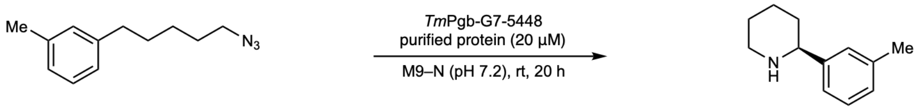

Reaction performed **with *Tm*Pgb-G7-5448** (purified protein)

| Entry | Product | IS | P/IS | Yield (%) | Average (%) | STDEV | Average TTN |
|---|---|---|---|---|---|---|---|
| 1 | 56.9 | 4748.3 | 0.012 | 1.5 | | | |
| 2 | 41.1 | 4396.5 | 0.009 | 1.2 | 1.3 | 0.2 | 3 |
| 3 | 44.0 | 4438.5 | 0.010 | 1.2 | | | |

**Calibration curve and data analysis for 2j**

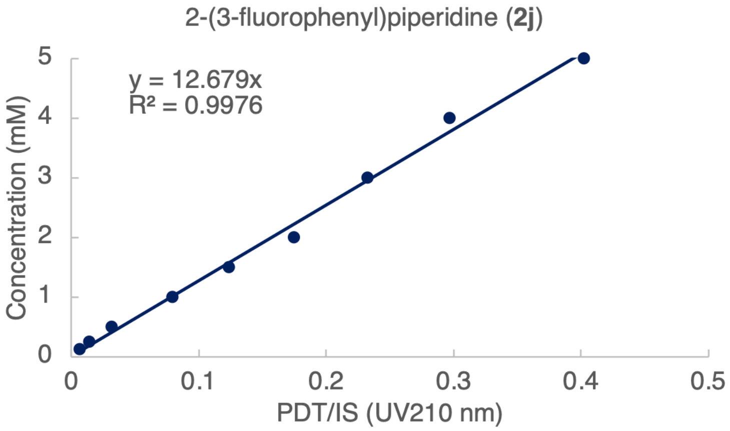

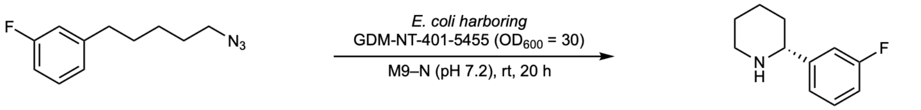

Reaction performed **with GDM-NT-401-5455** (whole *E.coli* cell with 2.8 μM heme loading)

| Entry | Product | IS | P/IS | Yield (%) | Average (%) | STDEV | Average TTN |
|---|---|---|---|---|---|---|---|
| 1 | 476.1 | 4533.5 | 0.11 | 26.6 | | | |
| 2 | 315.2 | 4402.9 | 0.07 | 18.2 | 21 | 4.8 | 376 |
| 3 | 305.3 | 4202.3 | 0.07 | 18.4 | | | |

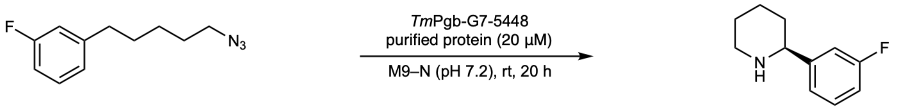

Reaction performed **with *Tm*Pgb-G7-5448** (purified protein)

| Entry | Product | IS | P/IS | Yield (%) | Average (%) | STDEV | Average TTN |
|---|---|---|---|---|---|---|---|
| 1 | 44.0 | 6296.9 | 0.007 | 1.8 | | | |
| 2 | 51.1 | 6303.4 | 0.008 | 2.1 | 2 | 0.1 | 5 |
| 3 | 49.0 | 6279.7 | 0.008 | 2.0 | | | |

**Calibration curve and data analysis for 2k**

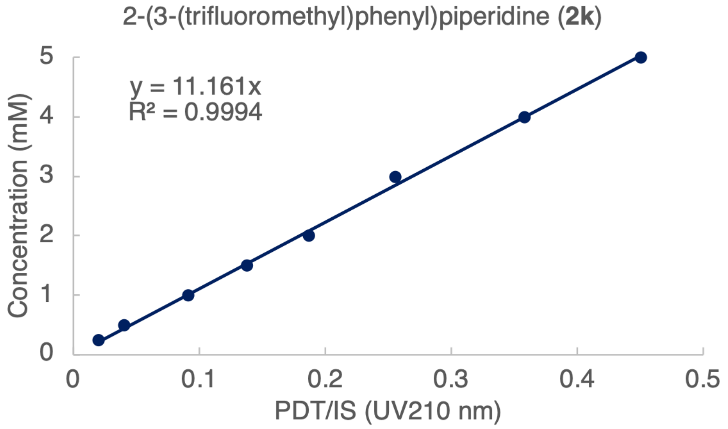

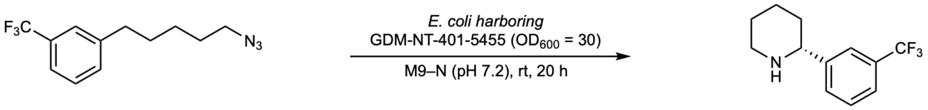

| Reaction performed **with GDM-NT-401-5455** (whole *E.coli* cell with 2.8 μM heme loading) | | | | | | | |
|---|---|---|---|---|---|---|---|
| Entry | Product | IS | P/IS | Yield (%) | Average (%) | STDEV | Average TTN |
| 1 | 396.4 | 4568.3 | 0.09 | 19.4 | | | |
| 2 | 371.1 | 4482.0 | 0.08 | 18.5 | 19 | 0.6 | 334 |
| 3 | 380.5 | 4625.9 | 0.08 | 18.4 | | | |

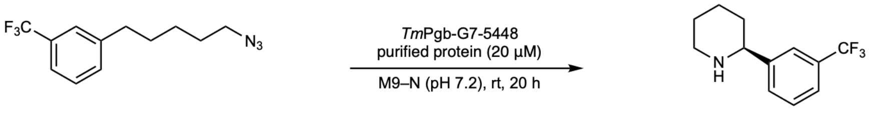

**1k** **2k**, <1% yield

Reaction performed **with *Tm*Pgb-G7-5448** (purified protein with 20 μM heme loading) furnished **2k** in <1% yield.

**Calibration curve and data analysis for 2l**

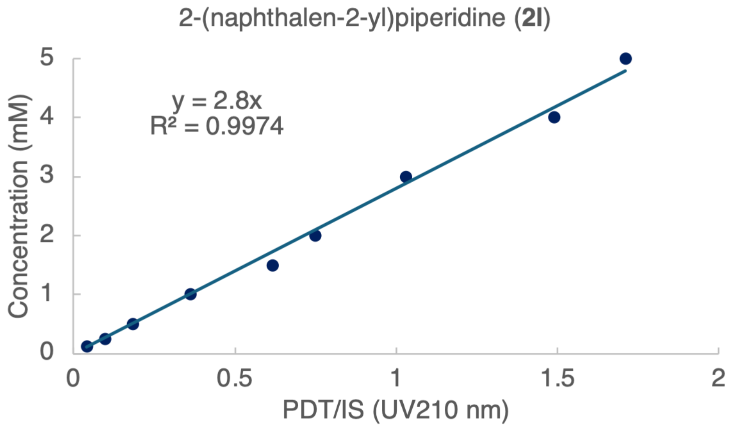

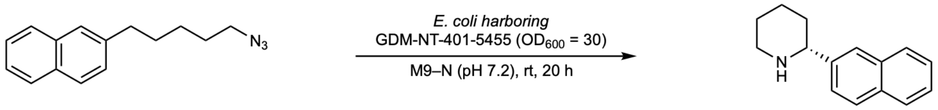

| Reaction performed **with GDM-NT-401-5455** (whole *E.coli* cell with 2.8 μM heme loading) | | | | | | | |
|---|---|---|---|---|---|---|---|
| Entry | Product | IS | P/IS | Yield (%) | Average (%) | STDEV | Average TTN |
| 1 | 306.4 | 4721.5 | 0.06 | 3.6 | | | |
| 2 | 268.8 | 4632.7 | 0.06 | 3.2 | 3 | 0.2 | 60 |
| 3 | 259.9 | 4556.9 | 0.06 | 3.2 | | | |

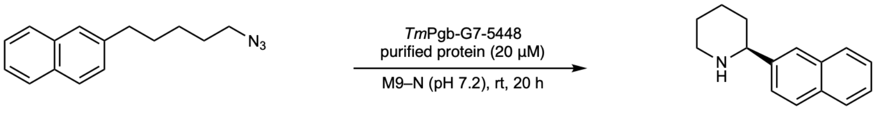

**1l** **2l**, <1% yield

Reaction performed **with *Tm*Pgb-G7-5448** (purified protein with 20 μM heme loading) furnished **2l** in <1% yield.

**Calibration curve and data analysis for 2m**

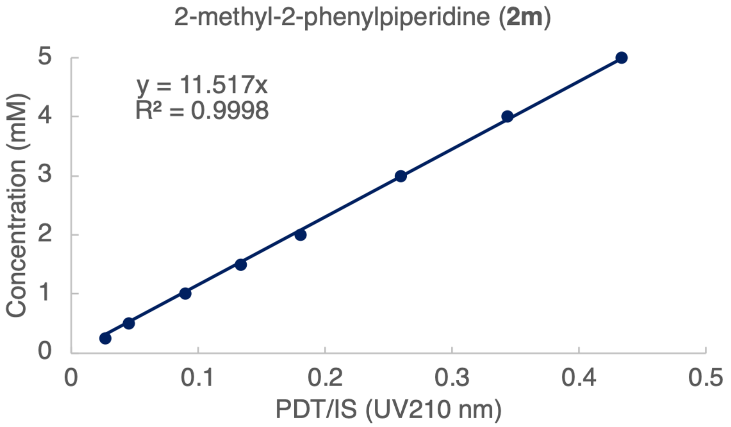

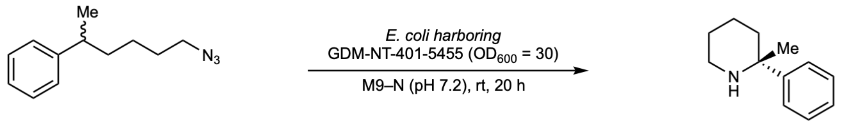

Reaction performed **with GDM-NT-401-5455** (whole *E.coli* cell with 2.8 μM heme loading)

| Entry | Product | IS | P/IS | Yield (%) | Average (%) | STDEV | Average TTN |
|---|---|---|---|---|---|---|---|
| 1 | 425.0 | 4341.0 | 0.10 | 22.6 | | | |
| 2 | 443.0 | 4359.8 | 0.10 | 23.4 | 23 | 0.5 | 407 |
| 3 | 426.9 | 4373.5 | 0.10 | 22.5 | | | |

Reaction performed **with GDM-NT-401-5455** (purified protein with 20 μM heme loading)

| Entry | Product | IS | P/IS | Yield (%) | Average (%) | STDEV | Average TTN |
|---|---|---|---|---|---|---|---|
| 1 | 853.2 | 4295.7 | 0.20 | 45.8 | | | |
| 2 | 851.8 | 4325.7 | 0.20 | 45.4 | 45 | 0.4 | 113 |
| 3 | 847.9 | 4336.6 | 0.20 | 45.0 | | | |

**Calibration curve and data analysis for 2n**

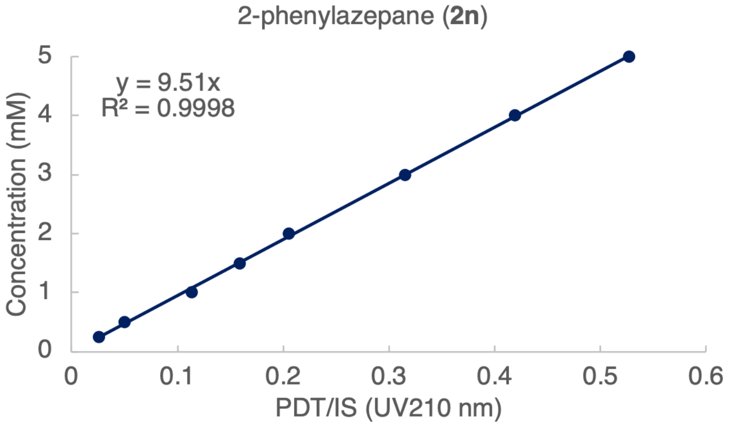

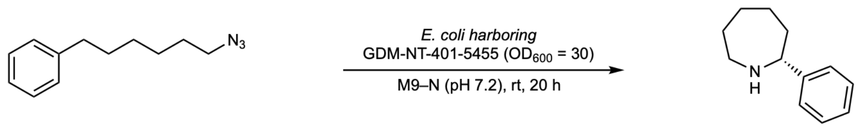

Reaction performed **with GDM-NT-401-5455** (whole *E.coli* cell)

| Entry | Product | IS | P/IS | Yield (%) | Average (%) | STDEV | Average TTN |
|---|---|---|---|---|---|---|---|
| 1 | 282.3 | 4323.3 | 0.07 | 12.4 | | | |
| 2 | 292.8 | 4457.5 | 0.07 | 12.5 | 13 | 0.9 | 232 |
| 3 | 326.9 | 4438.7 | 0.07 | 14.0 | | | |

**Calibration curve and data analysis for 2o**

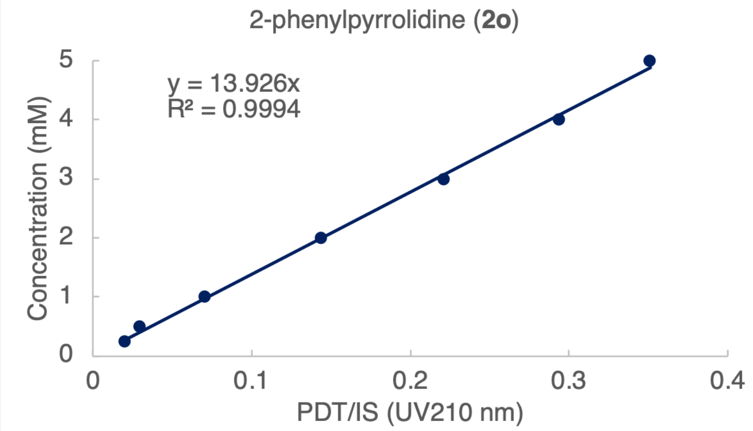

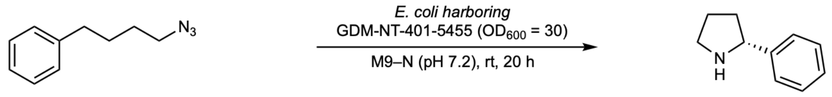

Reaction performed **with GDM-NT-401-5455** (whole *E.coli* cell)

| Entry | Product | IS | P/IS | Yield (%) | Average (%) | STDEV | Average TTN |
|---|---|---|---|---|---|---|---|
| 1 | 754.2 | 4485.0 | 0.17 | 46.8 | | | |
| 2 | 784.2 | 4434.7 | 0.18 | 49.3 | 48 | 1.4 | 851 |
| 3 | 753.1 | 4477.3 | 0.17 | 46.9 | | | |

**Calibration curve and data analysis for 2p**

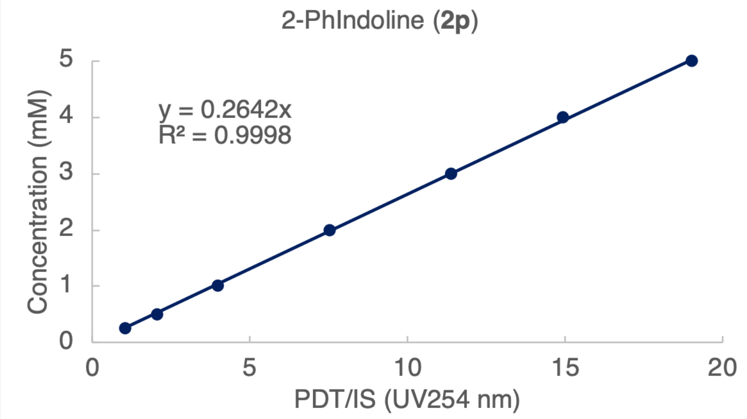

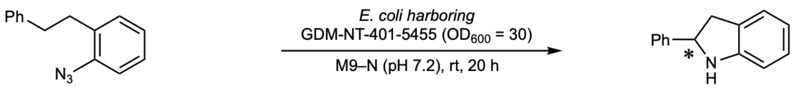

Reaction performed **with GDM-NT-401-5455** (whole *E.coli* cell)

| Entry | Product | IS | P/IS | Yield (%) | Average (%) | STDEV | Average TTN |
|---|---|---|---|---|---|---|---|
| 1 | 1144.5 | 94.7 | 12.09 | 63.9 | 64 | 1.1 | 1144 |
| 2 | 1138.4 | 92.2 | 12.35 | 65.2 | | | |

| | | | | |
|---|---|---|---|---|
| 3 | 1114.6 | 93.4 | 11.93 | 63.1 |

**Calibration curve and data analysis for 2q**

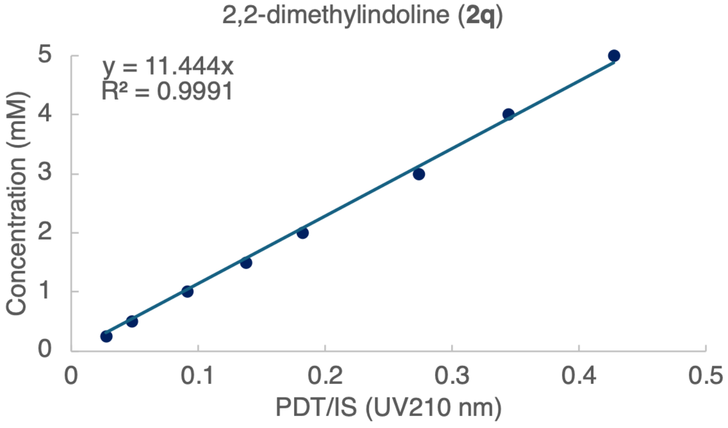

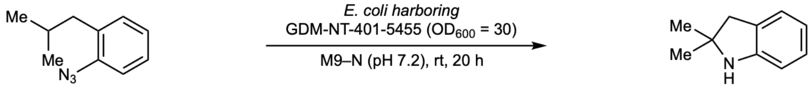

Reaction performed **with GDM-NT-401-5455** (whole *E.coli* cell)

| Entry | Product | IS | P/IS | Yield (%) | Average (%) | STDEV | Average TTN |
|---|---|---|---|---|---|---|---|
| 1 | 2268.5 | 5164.6 | 0.44 | 100.5 | | | |
| 2 | 2152.5 | 5250.2 | 0.41 | 93.8 | 99 | 4.7 | 1769 |
| 3 | 2272.8 | 5057.5 | 0.45 | 102.9 | | | |

**Analytical-Scale Enzymatic Reaction Regioselectivity Analysis**

As all reactions performed with **GDM-NT-401-5455** have >99:1 rr, we omitted this section of original data for simplicity (refer to enantioselectivity analysis section for detailed evidence). In this section, we only demonstrate regioselectivity of reactions performed with ***Tm*Pgb-G7-5448**. All regioselectivities were determined by analyzing the integration ratio on single ion channel mass spectrometry. There are two sources of original data: 1) reverse-phase HPLC-MS analysis using C18 column. 2) chiral polar ionic HPLC-MS analysis using chiral V column. The best source of data for analyzing regioselectivity was chosen based on resolution factor ($R_s$). When HPLC-MS analysis using C18 column was used, we show the screenshots of the single ion channel spectrums below. When HPLC-MS analysis using polar ionic chiral V column was used, please refer to the enantioselectivity analysis section for screenshots of the spectrums.

**Part 1: Regioselectivity analysis of *Tm*Pgb-PIP lineage.**

**Table S23**. Regioselectivity analysis of ***Tm*Pgb-PIP** lineage. Results analyzed on polar ionic HPLC-MS.

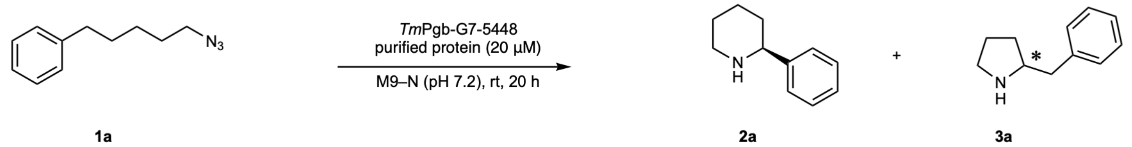

| | **2a**-f1 | **2a**-f2 | **3a**-f1 | **3a**-f2 | **2a** | **3a** | rr (**2a:3a**) |
|---|---|---|---|---|---|---|---|
| ***Tm*Pgb-wt** | 4578 | 230690 | 455684 | 8001453 | 235267 | 8457137 | 3:97 |
| **PIP-G1** | 0 | 1434458 | 1486397 | 270189 | 1434458 | 1756586 | 45:55 |
| **PIP-G2** | 0 | 1996270 | 1270715 | 290422 | 1996270 | 1561137 | 56:44 |
| **PIP-G3** | 0 | 2377421 | 708547 | 267999 | 2377421 | 976546 | 71:29 |
| **PIP-G4** | 0 | 5318512 | 471101 | 219056 | 5318512 | 690157 | 89:11 |
| **PIP-G5** | 0 | 10183343 | 1133631 | 514638 | 10183343 | 1648270 | 86:14 |
| **PIP-G6** | 0 | 8326293 | 1115724 | 356353 | 8326293 | 1472076 | 85:15 |
| **PIP-G7** | 0 | 16744088 | 2771103 | 467014 | 16744088 | 3238117 | 84:16 |

**Part 2: Regioselectivity analysis of GDM-NT variants**

**Table S24**. Regioselectivity analysis of Selected results using selected variants in recycled-scaffold design based on **GDM-NT-121**. Results analyzed on polar ionic HPLC-MS.

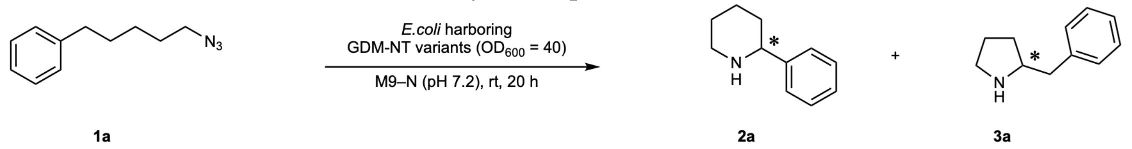

| | **2a**-f1 | **2a**-f2 | **3a**-f1 | **3a**-f2 | **2a** | **3a** | rr (**2a:3a**) |
|---|---|---|---|---|---|---|---|
| ***Tm*Pgb-wt** | 1010 | 64158 | 114644 | 1715495 | 65168 | 1830139 | 3:97 |
| **GDM-NT-121** | 105591 | 244460 | 5714 | 7718 | 350051 | 13432 | 96:4 |
| **GDM-NT-215** | 266139 | 430943 | 8949 | 8810 | 697081 | 17758 | 98:2 |
| **GDM-NT-217** | 285303 | 151843 | 4884 | 12284 | 437147 | 17168 | 96:4 |
| **GDM-NT-226** | 4647607 | 2520857 | 1871 | 3029 | 7168464 | 4901 | >99:1 |
| **GDM-NT-227** | 189184 | 175706 | 11246 | 12413 | 364890 | 23659 | 94:6 |
| **GDM-NT-228** | 85353 | 28646 | 7995 | 5448 | 113999 | 13444 | 89:11 |

**Table S25**. Regioselectivity analysis of Selected results using selected variants in recycled-scaffold design based on **GDM-NT-151**. Results analyzed on polar ionic HPLC-MS.

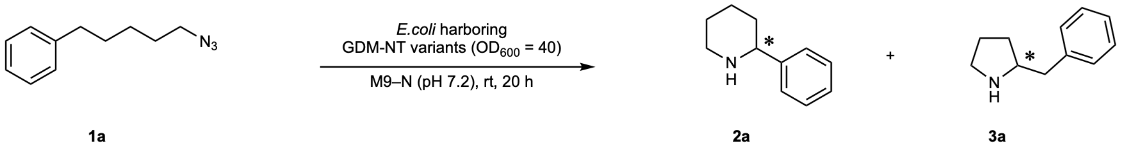

| | **2a**-f1 | **2a**-f2 | **3a**-f1 | **3a**-f2 | **2a** | **3a** | rr (**2a:3a**) |
|---|---|---|---|---|---|---|---|
| ***Tm*Pgb-wt** | 1010 | 64158 | 114644 | 1715495 | 65168 | 1830139 | 3:97 |
| **GDM-NT-151** | 18301 | 98460 | 2733 | 3969 | 116761 | 6702 | 95:5 |
| **GDM-NT-350** | 113820 | 297942 | 0 | 25090 | 411761 | 25090 | 94:6 |
| **GDM-NT-356** | 171827 | 1100276 | 6839 | 6149 | 1272102 | 12988 | 99:1 |
| **GDM-NT-359** | 168656 | 767392 | 3480 | 12155 | 936048 | 15635 | 98:2 |
| **GDM-NT-361** | 124639 | 197673 | 2243 | 16327 | 322312 | 18570 | 95:5 |
| **GDM-NT-362** | 1770315 | 102717 | 2701 | 1195 | 1873032 | 3896 | >99:1 |

**Part 3: Regioselectivity analysis of substrate scope study.**

Reactions performed with ***Tm*Pgb-PIP-G7-5448** on polar ionic chiral HPLC-MS or reverse-phase HPLC-MS.

Regioselectivity analysis of **2a** using ***Tm*Pgb-PIP-G7-5448** (polar ionic HPLC-MS):

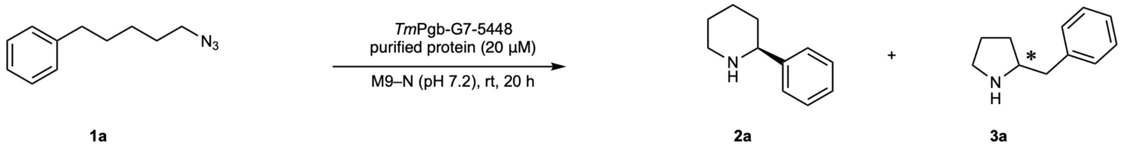

| | **2a**-f1 | **2a**-f2 | **3a**-f1 | **3a**-f2 | **2a** | **3a** | rr (**2a:3a**) |
|---|---|---|---|---|---|---|---|
| **Area (ion counts)** | 0 | 9426293 | 1667976 | 495131 | 9426293 | 2163110 | 81:19 |

Regioselectivity analysis of **2b** using ***Tm*Pgb-PIP-G7-5448** (polar ionic HPLC-MS):

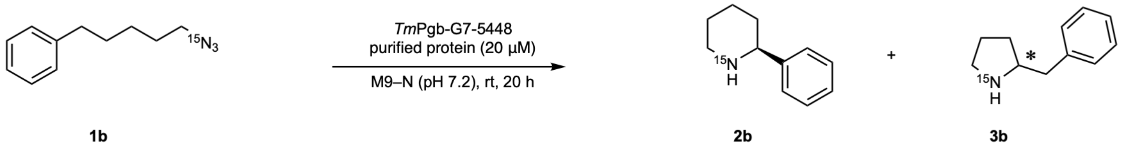

| | **2a**-f1 | **2a**-f2 | **3a**-f1 | **3a**-f2 | **2a** | **3a** | rr (**2a:3a**) |
|---|---|---|---|---|---|---|---|
| **Area (ion counts)** | 0 | 2834641 | 634790 | 184361 | 2834641 | 819151 | 78:22 |

Regioselectivity analysis of **2c** using ***Tm*Pgb-PIP-G7-5448** (polar ionic HPLC-MS):

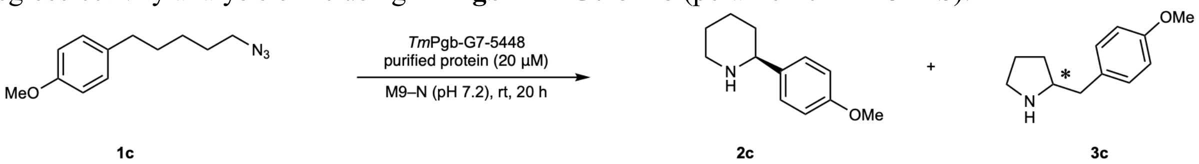

| | **2c**-f1 | **2c**-f2 | **3c**-f1 | **3c**-f2 | **2c** | **3c** | rr (**2c:3c**) |
|---|---|---|---|---|---|---|---|
| **Area (ion counts)** | 0 | 203817.2 | 1315.3 | 37518.9 | 203817.2 | 38834.2 | 84:16 |

Regioselectivity analysis of **2f** using ***Tm*Pgb-PIP-G7-5448** (reverse phase HPLC-MS on C18 column):

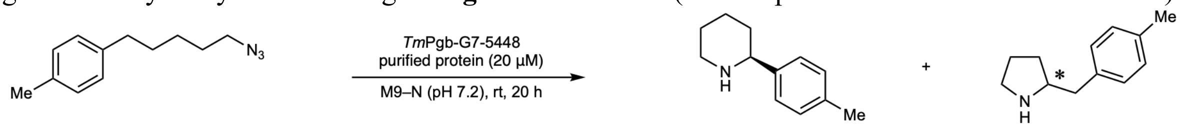

**1d** **2d** **3d**

| | 2e | 3e | rr (2e:3e) | Average rr (2e:3e) |
|---|---|---|---|---|
| **1** | 377340.2 | 30129.2 | 92.6:7.4 | |
| **2** | 391193.4 | 27640.4 | 93.4:6.6 | 93:7 |
| **3** | 387767.1 | 30128.6 | 92.8:7.2 | |

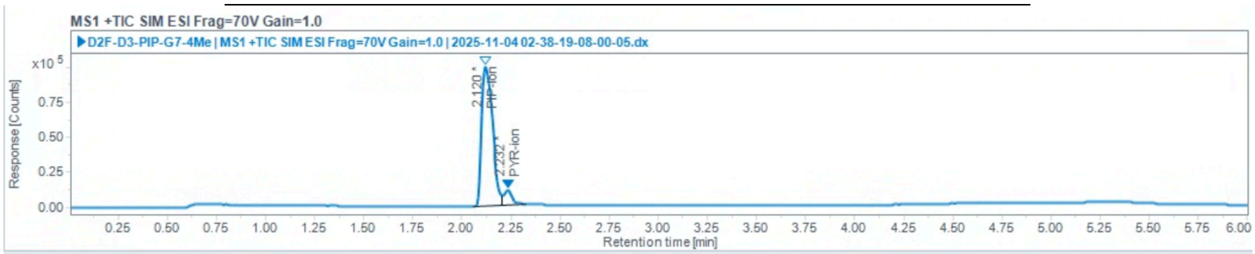

Injection Results

Peaks Summary

| Name | Signal description | RT (min) | Area (Counts·s) | Area% | Height (Counts) | Height% | Start time (min) | End time (min) |
|---|---|---|---|---|---|---|---|---|
| PIP-ion | MS1 +TIC SIM ESI Frag=70V Gain=1.0 | 2.120 | 377340.223 | 92.606 | 99584.696 | 90.24 | 2.059 | 2.205 |
| | MS1 +TIC SIM ESI Frag=70V Gain=1.0 | 2.232 | 30129.180 | 7.394 | 10768.210 | 9.76 | 2.205 | 2.326 |

lcurve-90A.pmx

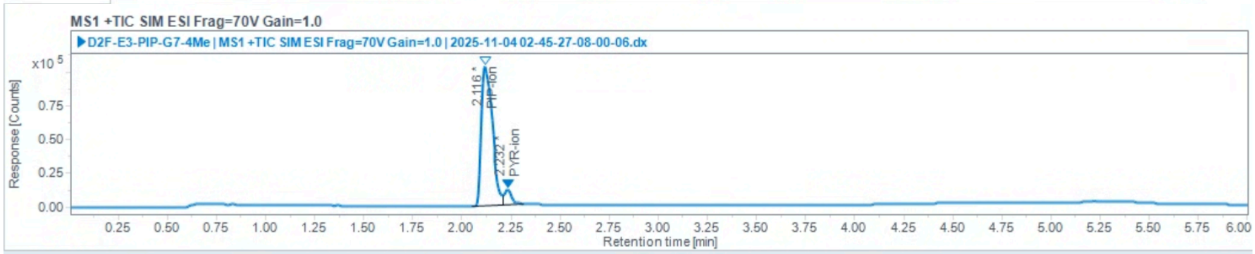

Injection Results

Peaks Summary

| Name | Signal description | RT (min) | Area (Counts·s) | Area% | Height (Counts) | Height% | Start time (min) | End time (min) |
|---|---|---|---|---|---|---|---|---|
| PIP-ion | MS1 +TIC SIM ESI Frag=70V Gain=1.0 | 2.116 | 391193.422 | 93.401 | 103046.399 | 90.28 | 2.049 | 2.209 |
| PYR-ion | MS1 +TIC SIM ESI Frag=70V Gain=1.0 | 2.232 | 27640.372 | 6.599 | 11091.129 | 9.72 | 2.209 | 2.316 |

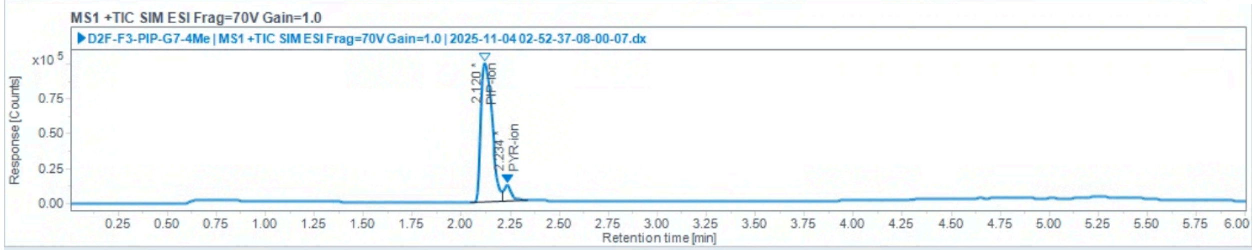

Injection Results

Peaks Summary

| Name | Signal description | RT (min) | Area (Counts·s) | Area% | Height (Counts) | Height% | Start time (min) | End time (min) |
|---|---|---|---|---|---|---|---|---|
| PIP-ion | MS1 +TIC SIM ESI Frag=70V Gain=1.0 | 2.120 | 387767.074 | 92.790 | 99736.682 | 89.84 | 2.048 | 2.209 |
| PYR-ion | MS1 +TIC SIM ESI Frag=70V Gain=1.0 | 2.234 | 30128.574 | 7.210 | 11283.322 | 10.16 | 2.209 | 2.337 |

Regioselectivity analysis of **2f** using ***Tm*Pgb-PIP-G7-5448** (reverse phase HPLC-MS on C18 column):

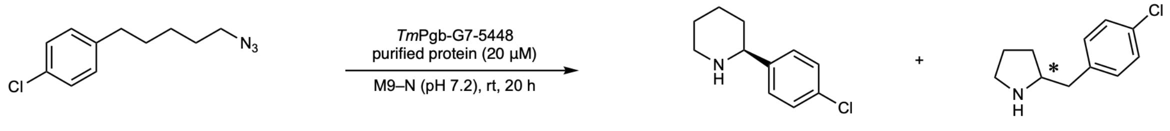

**1e** **2e** **3e**

| | **2e** | **3e** | rr (**2e**:**3e**) | Average rr (**2e**:**3e**) |
|---|---|---|---|---|
| **1** | 170626.9 | 14325.8 | 92.3:7.7 | |
| **2** | 169557.4 | 13730.7 | 92.5:7.5 | 92:8 |
| **3** | 150887.4 | 12134.5 | 92.6:7.4 | |

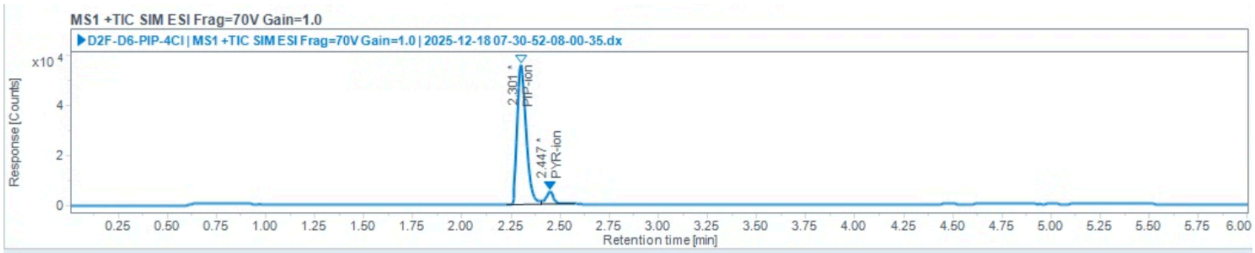

Injection Results

Peaks Summary

| Name | Signal description | RT (min) | Area (Counts·s) | Area% | Height (Counts) | Height% | Start time (min) | End time (min) |
|---|---|---|---|---|---|---|---|---|
| PIP-ion | MS1 +TIC SIM ESI Frag=70V Gain=1.0 | 2.301 | 170626.918 | 92.254 | 56006.345 | 91.55 | 2.226 | 2.404 |
| PYR-ion | MS1 +TIC SIM ESI Frag=70V Gain=1.0 | 2.447 | 14325.824 | 7.746 | 5171.816 | 8.45 | 2.404 | 2.576 |

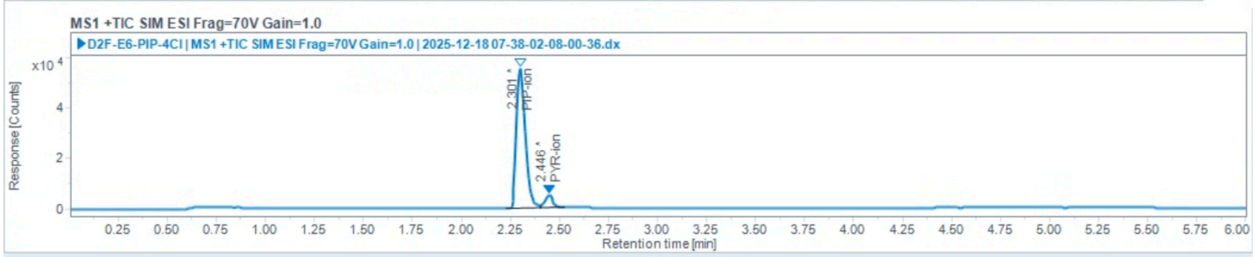

Injection Results

Peaks Summary

| Name | Signal description | RT (min) | Area (Counts·s) | Area% | Height (Counts) | Height% | Start time (min) | End time (min) |
|---|---|---|---|---|---|---|---|---|
| PIP-ion | MS1 +TIC SIM ESI Frag=70V Gain=1.0 | 2.301 | 169557.362 | 92.509 | 55239.240 | 91.86 | 2.228 | 2.404 |
| PYR-ion | MS1 +TIC SIM ESI Frag=70V Gain=1.0 | 2.446 | 13730.691 | 7.491 | 4892.416 | 8.14 | 2.404 | 2.528 |

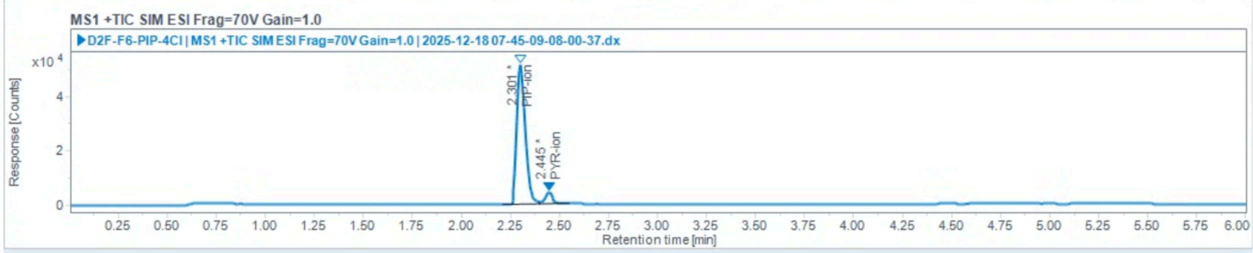

Injection Results

Peaks Summary

| Name | Signal description | RT (min) | Area (Counts·s) | Area% | Height (Counts) | Height% | Start time (min) | End time (min) |
|---|---|---|---|---|---|---|---|---|
| PIP-ion | MS1 +TIC SIM ESI Frag=70V Gain=1.0 | 2.301 | 150887.444 | 92.557 | 51420.110 | 92.35 | 2.210 | 2.401 |
| PYR-ion | MS1 +TIC SIM ESI Frag=70V Gain=1.0 | 2.445 | 12134.513 | 7.443 | 4257.567 | 7.65 | 2.401 | 2.547 |

Regioselectivity analysis of **2f** using ***Tm*Pgb-PIP-G7-5448** (reverse phase HPLC-MS on C18 column):

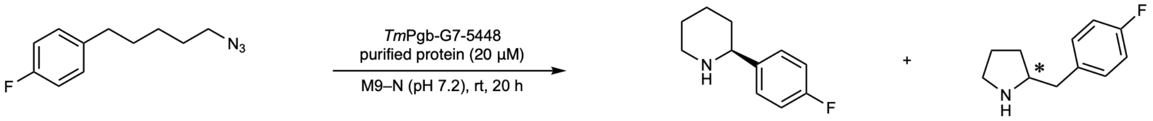

**1f** **2f** **3f**

| | **2f** | **3f** | rr (**2f**:**3f**) | Average rr (**2f**:**3f**) |
|---|---|---|---|---|
| **1** | 154974.7 | 27128.5 | 85.1:14.9 | |
| **2** | 127690.2 | 24034.0 | 84.2:15.8 | 84:16 |
| **3** | 161052.5 | 31395.7 | 83.7:16.3 | |

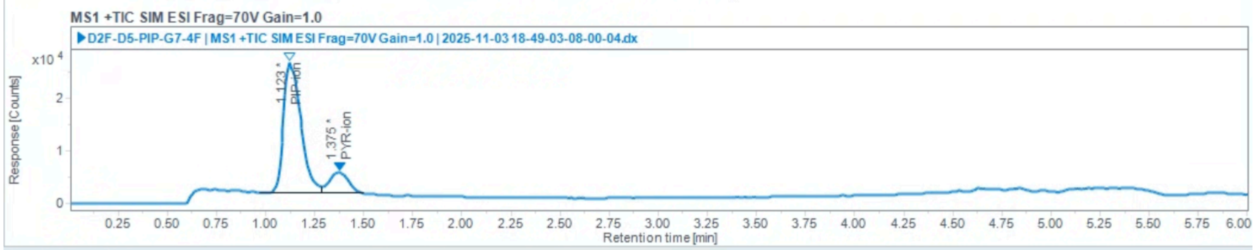

Injection Results

Peaks Summary

| Name | Signal description | RT (min) | Area (Counts·s) | Area% | Height (Counts) | Height% | Start time (min) | End time (min) |
|---|---|---|---|---|---|---|---|---|
| PIP-ion | MS1 +TIC SIM ESI Frag=70V Gain=1.0 | 1.123 | 154974.680 | 85.103 | 24939.911 | 86.01 | 0.972 | 1.289 |
| PYR-ion | MS1 +TIC SIM ESI Frag=70V Gain=1.0 | 1.375 | 27128.520 | 14.897 | 4055.868 | 13.99 | 1.289 | 1.498 |

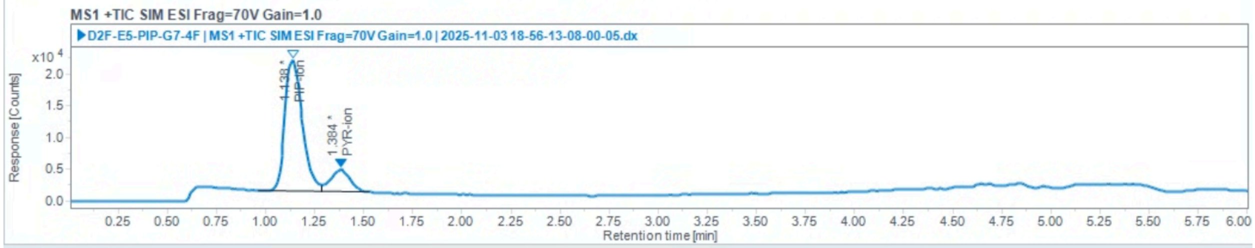

Injection Results

Peaks Summary

| Name | Signal description | RT (min) | Area (Counts·s) | Area% | Height (Counts) | Height% | Start time (min) | End time (min) |
|---|---|---|---|---|---|---|---|---|
| PIP-ion | MS1 +TIC SIM ESI Frag=70V Gain=1.0 | 1.138 | 127690.195 | 84.159 | 20574.759 | 85.15 | 0.961 | 1.289 |
| PYR-ion | MS1 +TIC SIM ESI Frag=70V Gain=1.0 | 1.384 | 24034.029 | 15.841 | 3587.499 | 14.85 | 1.289 | 1.531 |

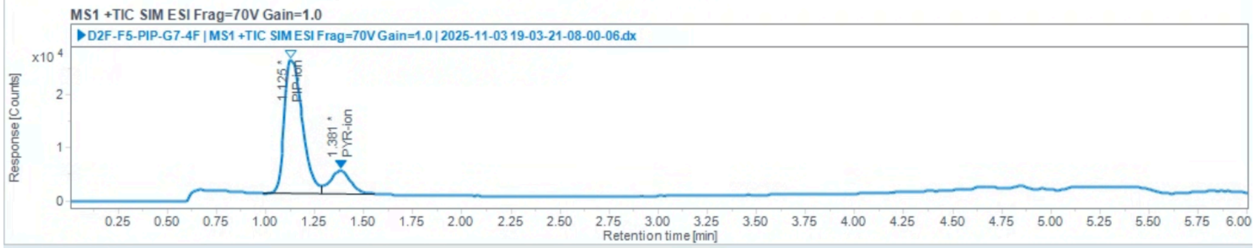

Injection Results

Peaks Summary

| Name | Signal description | RT (min) | Area (Counts·s) | Area% | Height (Counts) | Height% | Start time (min) | End time (min) |
|---|---|---|---|---|---|---|---|---|
| PIP-ion | MS1 +TIC SIM ESI Frag=70V Gain=1.0 | 1.125 | 161052.532 | 83.686 | 25143.111 | 84.87 | 0.989 | 1.289 |
| PYR-ion | MS1 +TIC SIM ESI Frag=70V Gain=1.0 | 1.381 | 31395.699 | 16.314 | 4481.209 | 15.13 | 1.289 | 1.554 |

Regioselectivity analysis of **2g** using ***Tm*Pgb-PIP-G7-5448** (reverse phase HPLC-MS on C18 column):

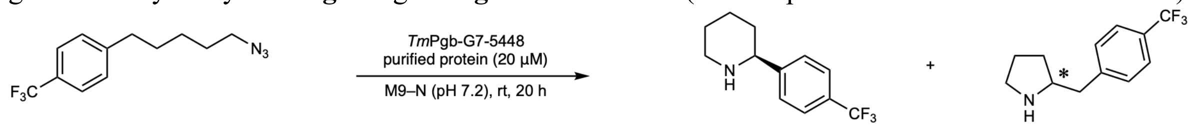

**1g** **2g** **3g**

| | **2g** | **3g** | rr (**2g**:**3g**) | Average rr (**2g**:**3g**) |
|---|---|---|---|---|
| **1** | 33822.5 | 2900.2 | 92.1:7.9 | |
| **2** | 32734.5 | 2659.8 | 92.5:7.5 | 92:8 |
| **3** | 34723.2 | 2842.1 | 92.4:92:6 | |

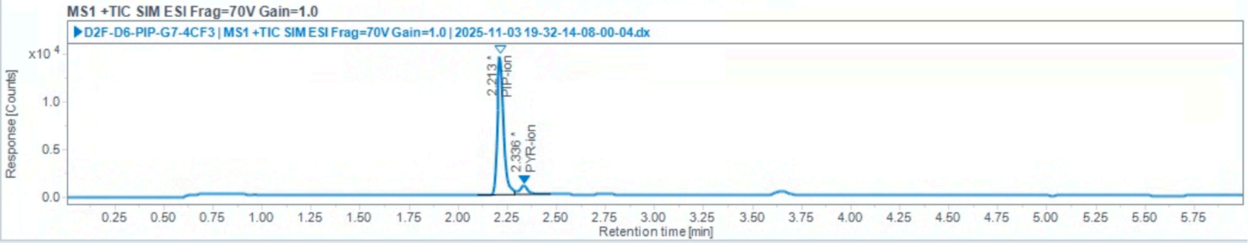

Injection Results

Peaks Summary

| Name | Signal description | RT (min) | Area (Counts·s) | Area% | Height (Counts) | Height% | Start time (min) | End time (min) |
|---|---|---|---|---|---|---|---|---|
| PIP-ion | MS1 +TIC SIM ESI Frag=70V Gain=1.0 | 2.213 | 33822.499 | 92.102 | 14647.982 | 93.77 | 2.101 | 2.287 |
| PYR-ion | MS1 +TIC SIM ESI Frag=70V Gain=1.0 | 2.336 | 2900.200 | 7.898 | 972.908 | 6.23 | 2.287 | 2.468 |

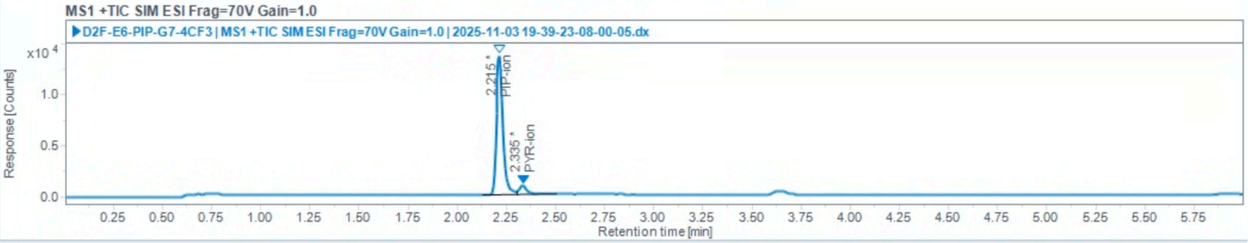

Injection Results

Peaks Summary

| Name | Signal description | RT (min) | Area (Counts·s) | Area% | Height (Counts) | Height% | Start time (min) | End time (min) |
|---|---|---|---|---|---|---|---|---|
| PIP-ion | MS1 +TIC SIM ESI Frag=70V Gain=1.0 | 2.215 | 32734.469 | 92.485 | 13677.457 | 93.62 | 2.129 | 2.304 |
| PYR-ion | MS1 +TIC SIM ESI Frag=70V Gain=1.0 | 2.335 | 2659.771 | 7.515 | 932.196 | 6.38 | 2.304 | 2.507 |

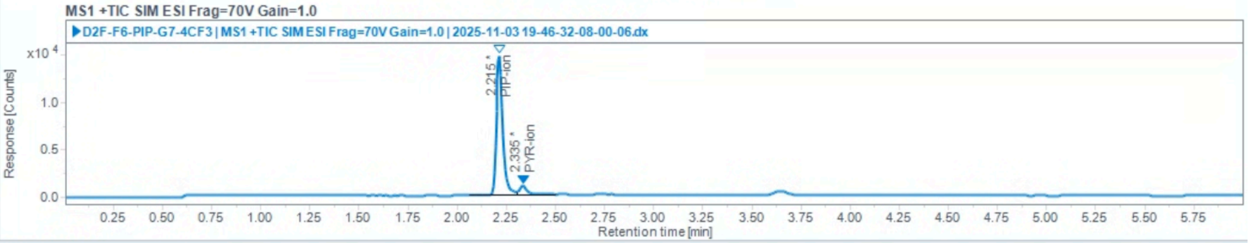

Injection Results

Peaks Summary

| Name | Signal description | RT (min) | Area (Counts·s) | Area% | Height (Counts) | Height% | Start time (min) | End time (min) |
|---|---|---|---|---|---|---|---|---|
| PIP-ion | MS1 +TIC SIM ESI Frag=70V Gain=1.0 | 2.215 | 34723.193 | 92.434 | 14859.534 | 93.80 | 2.061 | 2.304 |
| PYR-ion | MS1 +TIC SIM ESI Frag=70V Gain=1.0 | 2.335 | 2842.050 | 7.566 | 981.927 | 6.20 | 2.304 | 2.502 |

Regioselectivity analysis of **2h** using ***Tm*Pgb-PIP-G7-5448** (polar ionic HPLC-MS):

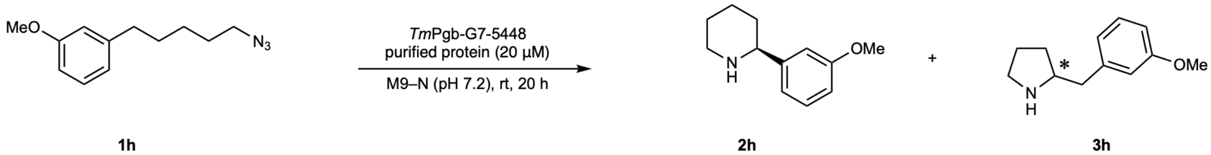

| | **2h**-f1 + **2h**-f2 | **3h**-f1 | **3h**-f2 | **2h** | **3h** | rr (**2h**:**3h**) |
|---|---|---|---|---|---|---|
| **Area (ion counts)** | 2802725.8 | 729527.0 | 89433.7 | 2802725.8 | 818960.7 | 77:23 |

Regioselectivity analysis of **2i** using ***Tm*Pgb-PIP-G7-5448** (reverse phase HPLC-MS on C18 column):

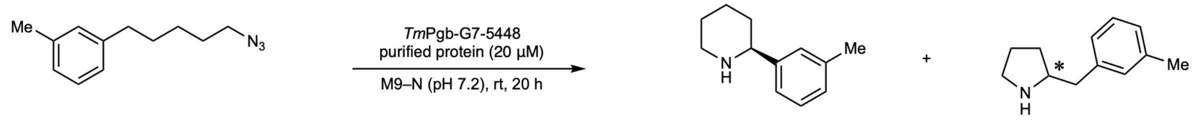

**1i** **2i** **3i**

| | **2i** | **3i** | rr (**2i**:**3i**) | Average rr (**2i**:**3i**) |
|---|---|---|---|---|
| 1 | 143320.0 | 64433.3 | 69.0:31.0 | |
| 2 | 115443.8 | 49819.4 | 69.9:30.1 | 70:30 |
| 3 | 132815.8 | 53755.6 | 71.2:28.8 | |

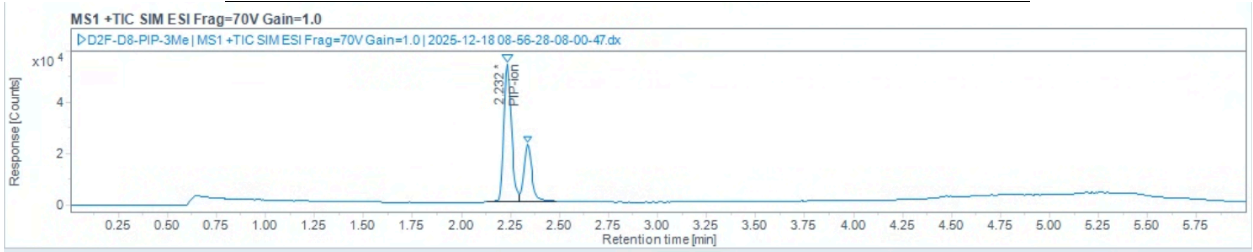

Injection Results

Peaks Summary

| Name | Signal description | RT (min) | Area (Counts·s) | Area% | Height (Counts) | Height% | Start time (min) | End time (min) |
|---|---|---|---|---|---|---|---|---|
| PIP-ion | MS1 +TIC SIM ESI Frag=70V Gain=1.0 | 2.232 | 143319.957 | 68.986 | 53576.910 | 70.43 | 2.131 | 2.296 |
| PYR-ion | MS1 +TIC SIM ESI Frag=70V Gain=1.0 | 2.337 | 64433.288 | 31.014 | 22496.532 | 29.57 | 2.296 | 2.486 |

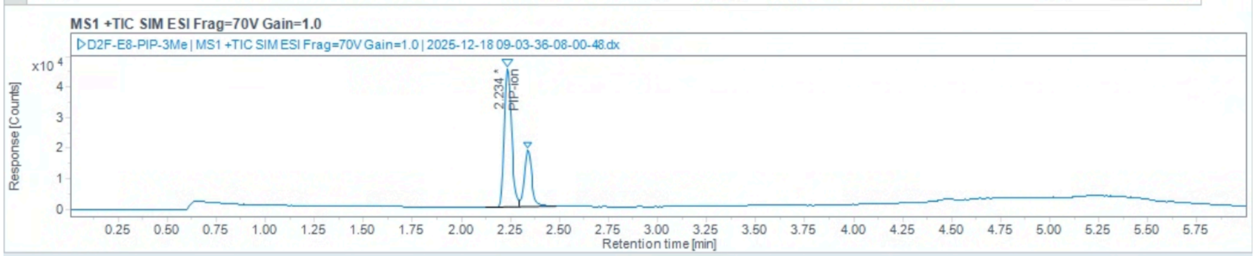

Injection Results

Peaks Summary

| Name | Signal description | RT (min) | Area (Counts·s) | Area% | Height (Counts) | Height% | Start time (min) | End time (min) |
|---|---|---|---|---|---|---|---|---|
| PIP-ion | MS1 +TIC SIM ESI Frag=70V Gain=1.0 | 2.234 | 115443.764 | 69.854 | 45279.191 | 70.95 | 2.123 | 2.296 |
| PYR-ion | MS1 +TIC SIM ESI Frag=70V Gain=1.0 | 2.338 | 49819.424 | 30.146 | 18543.154 | 29.05 | 2.296 | 2.483 |

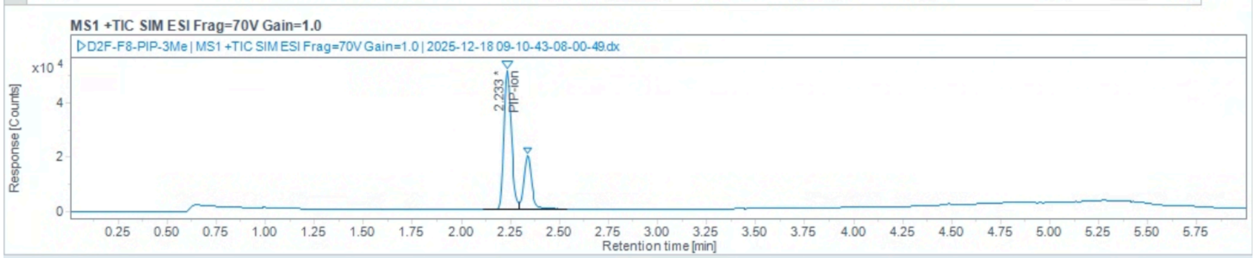

Injection Results

Peaks Summary

| Name | Signal description | RT (min) | Area (Counts·s) | Area% | Height (Counts) | Height% | Start time (min) | End time (min) |
|---|---|---|---|---|---|---|---|---|
| PIP-ion | MS1 +TIC SIM ESI Frag=70V Gain=1.0 | 2.233 | 132815.752 | 71.188 | 50984.159 | 72.07 | 2.114 | 2.294 |
| PYR-ion | MS1 +TIC SIM ESI Frag=70V Gain=1.0 | 2.337 | 53755.629 | 28.812 | 19756.403 | 27.93 | 2.294 | 2.538 |

Regioselectivity analysis of **2j** using ***Tm*Pgb-PIP-G7-5448** (reverse phase HPLC-MS on C18 column):

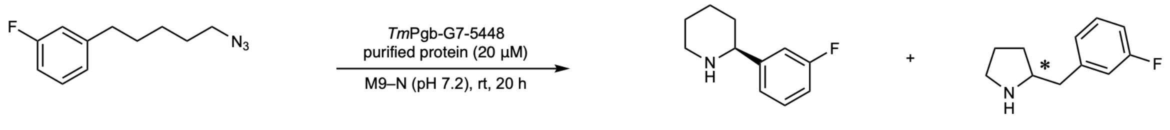

**1j** **2j** **3j**

| | **2j** | **3j** | rr (**2j**:**3j**) | Average rr (**2j**:**3j**) |
|---|---|---|---|---|
| 1 | 131907.4 | 36290.7 | 78.4:21.6 | |
| 2 | 114366.2 | 33668.3 | 77.3:22.7 | 77:23 |
| 3 | 125007.4 | 38649.2 | 76.4:23.6 | |

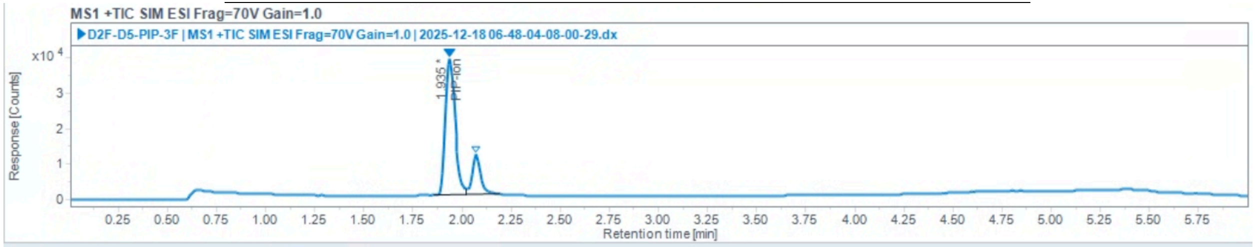

Injection Results

Peaks Summary

| Name | Signal description | RT (min) | Area (Counts·s) | Area% | Height (Counts) | Height% | Start time (min) | End time (min) |
|---|---|---|---|---|---|---|---|---|
| PIP-ion | MS1 +TIC SIM ESI Frag=70V Gain=1.0 | 1.935 | 131907.418 | 78.424 | 38562.151 | 77.34 | 1.854 | 2.018 |
| PYR-ion | MS1 +TIC SIM ESI Frag=70V Gain=1.0 | 2.072 | 36290.663 | 21.576 | 11296.480 | 22.66 | 2.018 | 2.193 |

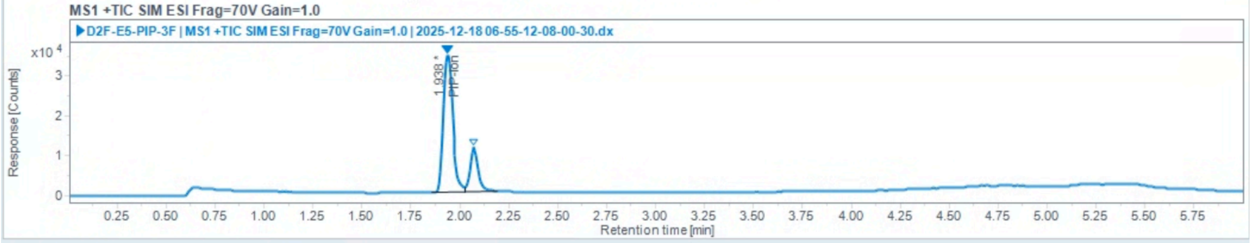

Injection Results

Peaks Summary

| Name | Signal description | RT (min) | Area (Counts·s) | Area% | Height (Counts) | Height% | Start time (min) | End time (min) |
|---|---|---|---|---|---|---|---|---|
| PIP-ion | MS1 +TIC SIM ESI Frag=70V Gain=1.0 | 1.938 | 114366.221 | 77.256 | 34470.986 | 75.50 | 1.857 | 2.025 |
| PYR-ion | MS1 +TIC SIM ESI Frag=70V Gain=1.0 | 2.071 | 33668.268 | 22.744 | 11185.764 | 24.50 | 2.025 | 2.194 |

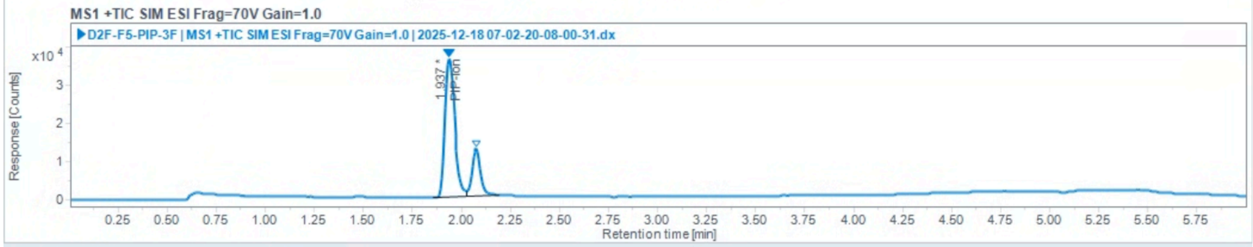

Injection Results

Peaks Summary

| Name | Signal description | RT (min) | Area (Counts·s) | Area% | Height (Counts) | Height% | Start time (min) | End time (min) |
|---|---|---|---|---|---|---|---|---|
| PIP-ion | MS1 +TIC SIM ESI Frag=70V Gain=1.0 | 1.937 | 125007.409 | 76.384 | 36238.843 | 74.33 | 1.854 | 2.027 |
| PYR-ion | MS1 +TIC SIM ESI Frag=70V Gain=1.0 | 2.076 | 38649.198 | 23.616 | 12512.391 | 25.67 | 2.027 | 2.194 |

Regioselectivity analysis of **2k** using ***Tm*Pgb-PIP-G7-5448** (reverse phase HPLC-MS on C18 column):

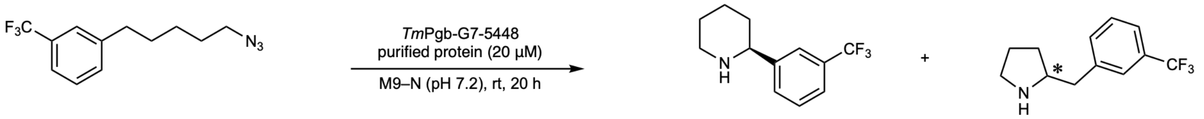

**1k** **2k** **3k**

| | **2k** | **3k** | rr (**2k**:**3k**) | Average rr (**2k**:**3k**) |
|---|---|---|---|---|
| 1 | 640.8 | 1528.1 | 29.5:70.5 | |
| 2 | 566.0 | 1304.5 | 30.3:69.7 | 30:70 |
| 3 | 513.2 | 1229.4 | 29.4:70.6 | |

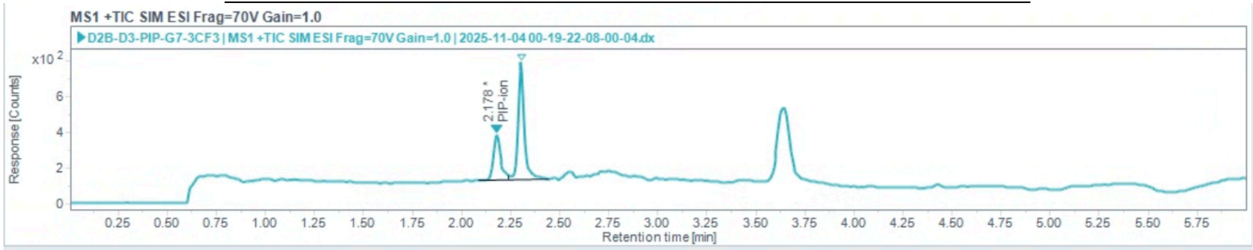

Injection Results

Peaks Summary

| Name | Signal description | RT (min) | Area (Counts·s) | Area% | Height (Counts) | Height% | Start time (min) | End time (min) |
|---|---|---|---|---|---|---|---|---|
| PIP-ion | MS1 +TIC SIM ESI Frag=70V Gain=1.0 | 2.178 | 640.787 | 29.544 | 250.550 | 27.68 | 2.089 | 2.241 |
| PYR-ion | MS1 +TIC SIM ESI Frag=70V Gain=1.0 | 2.303 | 1528.108 | 70.456 | 654.460 | 72.32 | 2.241 | 2.443 |

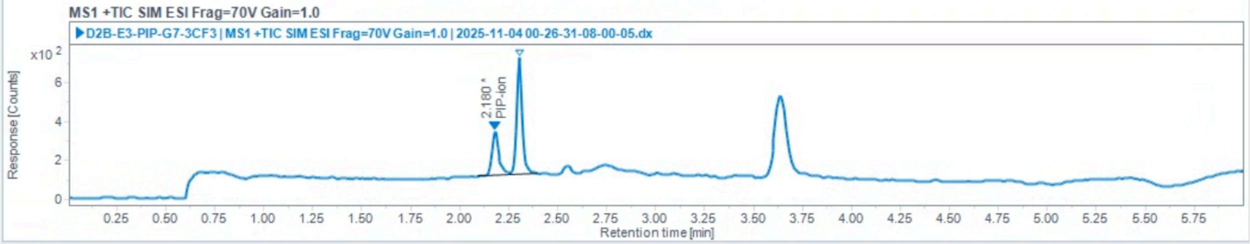

Injection Results

Peaks Summary

| Name | Signal description | RT (min) | Area (Counts·s) | Area% | Height (Counts) | Height% | Start time (min) | End time (min) |
|---|---|---|---|---|---|---|---|---|
| PIP-ion | MS1 +TIC SIM ESI Frag=70V Gain=1.0 | 2.180 | 565.985 | 30.259 | 226.489 | 27.34 | 2.095 | 2.260 |
| PYR-ion | MS1 +TIC SIM ESI Frag=70V Gain=1.0 | 2.304 | 1304.462 | 69.741 | 601.873 | 72.66 | 2.260 | 2.400 |

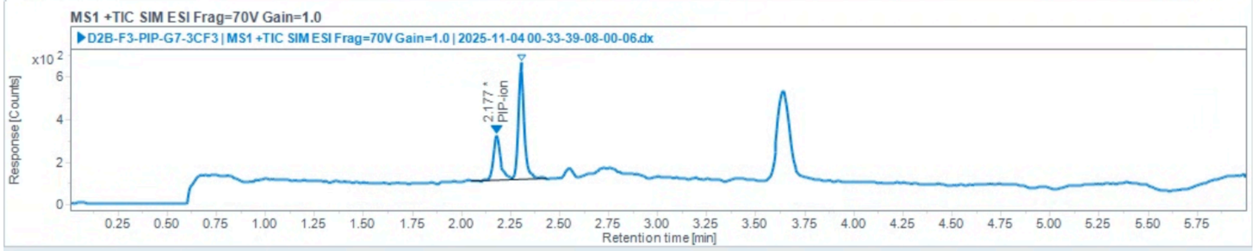

Injection Results

Peaks Summary

| Name | Signal description | RT (min) | Area (Counts·s) | Area% | Height (Counts) | Height% | Start time (min) | End time (min) |
|---|---|---|---|---|---|---|---|---|
| PIP-ion | MS1 +TIC SIM ESI Frag=70V Gain=1.0 | 2.177 | 513.164 | 29.449 | 206.349 | 27.49 | 2.052 | 2.260 |
| PYR-ion | MS1 +TIC SIM ESI Frag=70V Gain=1.0 | 2.303 | 1229.363 | 70.551 | 544.236 | 72.51 | 2.260 | 2.431 |

Regioselectivity analysis of **2l** using ***Tm*Pgb-PIP-G7-5448** (reverse phase HPLC-MS on C18 column):

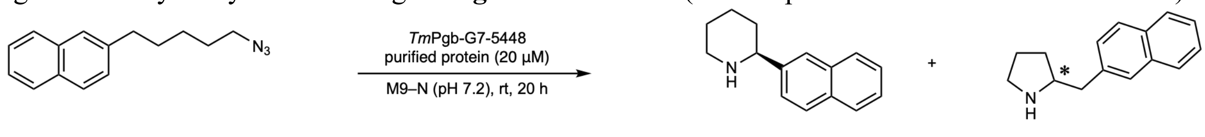

**1l** **2l** **3l**

| | **2l** | **3l** | rr (**2l**:**3l**) | Average rr (**2l**:**3l**) |
|---|---|---|---|---|
| 1 | 14285.0 | 7582.3 | 65.3:34.7 | |
| 2 | 12279.9 | 5893.3 | 67.6:32.4 | 66:34 |
| 3 | 13184.9 | 7277.3 | 64.4:35.6 | |

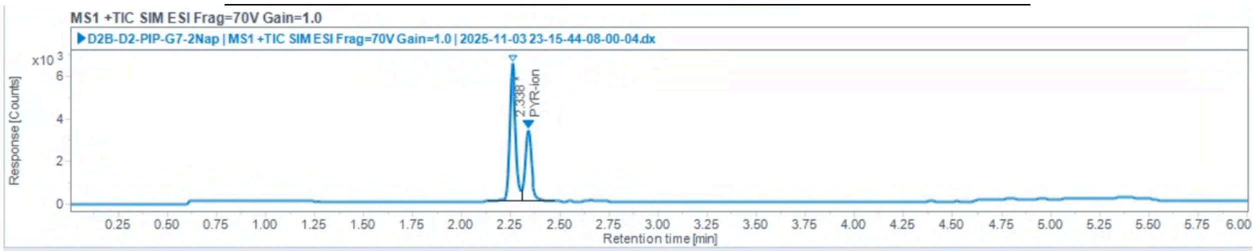

Injection Results

Peaks Summary

| Name | Signal description | RT (min) | Area (Counts·s) | Area% | Height (Counts) | Height% | Start time (min) | End time (min) |
|---|---|---|---|---|---|---|---|---|
| PIP-ion | MS1 +TIC SIM ESI Frag=70V Gain=1.0 | 2.257 | 14285.011 | 65.326 | 6486.467 | 65.84 | 2.131 | 2.306 |
| PYR-ion | MS1 +TIC SIM ESI Frag=70V Gain=1.0 | 2.338 | 7582.329 | 34.674 | 3365.907 | 34.16 | 2.306 | 2.470 |

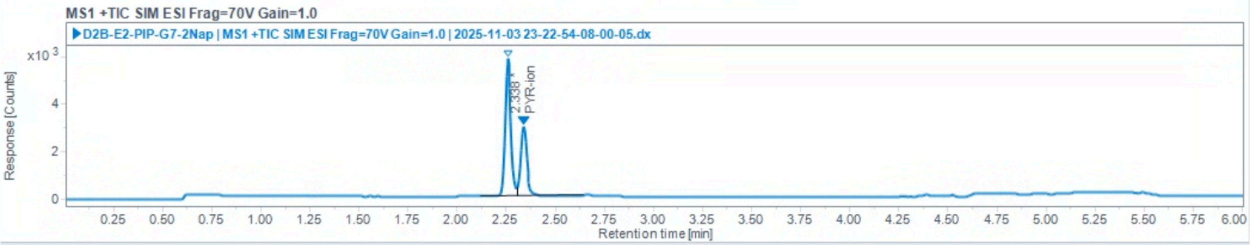

Injection Results

Peaks Summary

| Name | Signal description | RT (min) | Area (Counts·s) | Area% | Height (Counts) | Height% | Start time (min) | End time (min) |
|---|---|---|---|---|---|---|---|---|
| PIP-ion | MS1 +TIC SIM ESI Frag=70V Gain=1.0 | 2.258 | 12279.940 | 67.572 | 5812.029 | 66.53 | 2.120 | 2.306 |
| PYR-ion | MS1 +TIC SIM ESI Frag=70V Gain=1.0 | 2.338 | 5893.267 | 32.428 | 2923.298 | 33.47 | 2.306 | 2.645 |

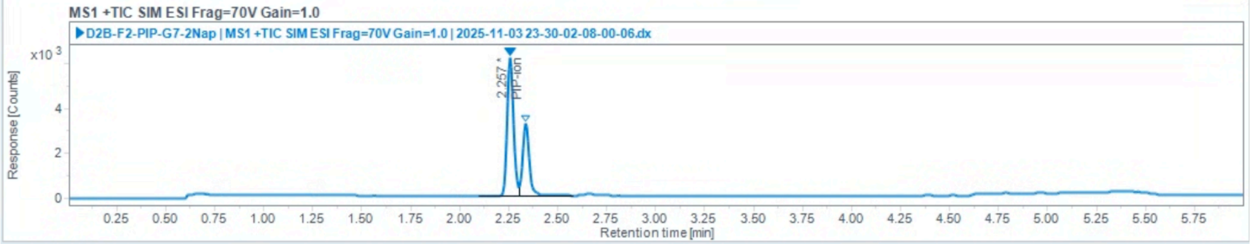

Injection Results

Peaks Summary

| Name | Signal description | RT (min) | Area (Counts·s) | Area% | Height (Counts) | Height% | Start time (min) | End time (min) |
|---|---|---|---|---|---|---|---|---|
| PIP-ion | MS1 +TIC SIM ESI Frag=70V Gain=1.0 | 2.257 | 13184.880 | 64.435 | 6147.658 | 65.56 | 2.101 | 2.304 |
| PYR-ion | MS1 +TIC SIM ESI Frag=70V Gain=1.0 | 2.338 | 7277.280 | 35.565 | 3229.100 | 34.44 | 2.304 | 2.581 |

Regioselectivity analysis of **2n** using **GDM-NT-401** (reverse phase HPLC-MS on C18 column):

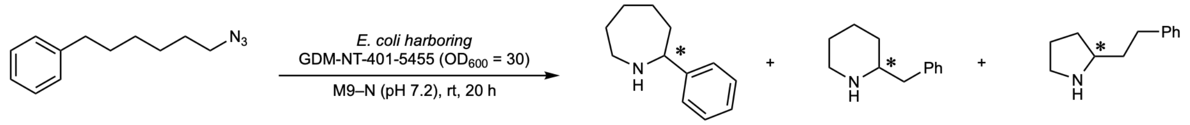

**1n** **2n** **3n** **4n**

89:8:3 rr (**2n**:**3n**:**4n**)

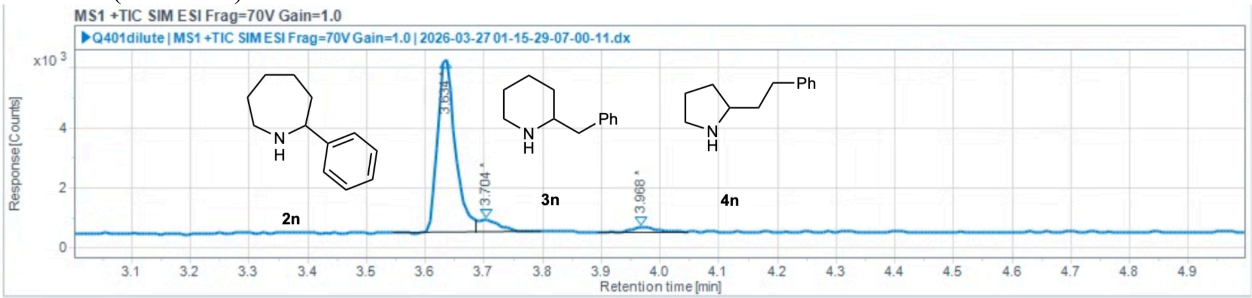

#### Injection Results

Peaks Summary

| # | Name | Signal description | RT (min) | Area (Counts·s) | Area% | Height (Counts) | Height% | Amou | Concentra | Start time ( |
|---|---|---|---|---|---|---|---|---|---|---|
| 1 | | MS1 +TIC SIM ESI Frag=70V Gain | 3.634 | 12083.326 | 88.828 | 5834.231 | 91.13 | | | 3.546 |
| 2 | | MS1 +TIC SIM ESI Frag=70V Gain | 3.704 | 1059.616 | 7.790 | 399.680 | 6.24 | | | 3.687 |
| 3 | | MS1 +TIC SIM ESI Frag=70V Gain | 3.968 | 460.180 | 3.383 | 168.116 | 2.63 | | | 3.895 |

Overlaied view of reaction mixture (orange, obtained using **GDM-NT-401-5455**), 2-phenylazepane (**2n**, purple, commercially available standard), 2-benzylpiperidine (**3n**, cyan, commercially available standard), 2-phenylethylpyrrolidine (**4n**, blue, commercially available standard),

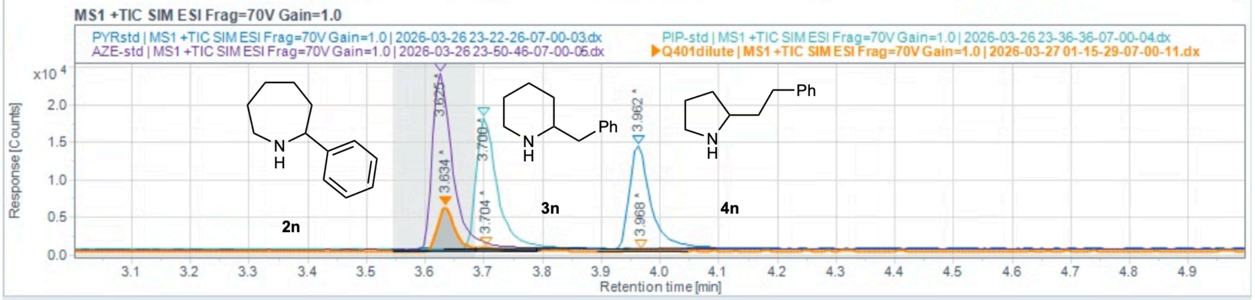

#### Injection Results

Peaks Summary

| # | Name | Signal description | RT (min) | Area (Counts·s) | Area% | Height (Counts) | Height% | Amou | Concentra | Start time ( |
|---|---|---|---|---|---|---|---|---|---|---|
| 1 | | MS1 +TIC SIM ESI Frag=70V Gain | 3.634 | 12083.326 | 88.828 | 5834.231 | 91.13 | | | 3.546 |
| 2 | | MS1 +TIC SIM ESI Frag=70V Gain | 3.704 | 1059.616 | 7.790 | 399.680 | 6.24 | | | 3.687 |
| 3 | | MS1 +TIC SIM ESI Frag=70V Gain | 3.968 | 460.180 | 3.383 | 168.116 | 2.63 | | | 3.895 |

#### Computational Details and Additional Computational Results

##### Computational Details - DFT calculations

All density functional theory (DFT) calculations were carried out using the Gaussian 16 progrm (*15*) . Geometries of intermediates and transition states were optimized in the gas phase using the dispersion-corrected (*16–17*) B3LYP-D3(BJ) functional (*18–19*) with a mixed basis set of SDD (*20*) (*i.e.*, the Stuttgart/Dresden effective core potentials and the associated valence electron basis set) for Fe, and 6-31G(d) for other atoms. Vibrational frequency calculations were performed to verify each optimised structure is a minimum (no imaginary frequencies) or a transition state (exactly one imaginary frequency). Thermal corrections to Gibbs free energies were calculated at 298 K. Single-point energy calculations were carried out using the B3LYP-D3(BJ) functional with the def2-TZVP basis set (*21–22*) for all atoms. Solvation energy corrections were calculated in single point energy calculations using the SMD solvation model (*23*) in diethyl ether to mimic the dielectric constant of enzyme active site (*24*). Wavefunction stability of all structures was confirmed using the "stable=opt" keyword. The bond dissociation enthalpies (BDEs) of δ-C–H and ε-C–H of substrate **1a** were calculated at the M06-2X (*25*)/def2-TZVP level theory in the gas phase, the same level of theory used in the ALFABET BDE database (*26*). CYLview (*27*) was used to visualize 3D structures of DFT-computed complexes and transition states.

##### DFT-computed reaction energy profiles with a 5-methylimidazole axial ligand

Reaction energy profiles of the δ- and ε-C–H amination were calculated using a model iron porphyrin complex with a 5-methylimidazole (Im) axial ligand and azide **1a** as the model substrate. The reaction starts with an iron–azide complex **CP1**, which undergoes nitrogen elimination through transition state **TS1** to generate the iron–nitrene species **INT1** with an energy barrier of 25.4 kcal/mol. Multiple conformers of the iron–nitrene species **INT1** were located, including reactive conformers **RC2a** and **RC2b**, in which the ε- and δ-C–H bonds point towards the nitrene nitrogen, poised for 1,6- and 1,5-hydrogen atom transfer (HAT), respectively. From reactive conformer **RC2b**, 1,5-HAT occurs through **TS2b** with an energy barrier of 17.3 kcal/mol to generate the homobenzylic radical species **INT2b**. The reactive conformer for 1,6-HAT (**RC2a**), which places the ε-C–H in proximity to the nitrene nitrogen, is 0.2 kcal/mol less stable than **RC2b**. From **RC2a**, 1,6-HAT occurs through **TS2a** to generate the benzylic radical species **INT2a**. From the radical species **INT2a** and **INT2b**, C–N radical rebound through **TS3a** and **TS3b** forms the piperidine complex **P2a** and pyrrolidine complex **P2b**, respectively. The C–N radical rebound steps are both highly exergonic and irreversible. In the 1,6- and 1,5-HAT pathways, the Gibbs free energies of the C–N radical rebound transition states (**TS3a** and **TS3b**) is 6.2 and 3.0 kcal/mol lower than those of the HAT transition states (**TS2a** and **TS2b**). These results indicate that the HAT in both pathways are irreversible and thus regioselectivity-determining. Because of the relatively high barriers to the subsequent HAT, the alkyl chain conformational change of the Fe–nitrene intermediate is expected to occur much faster. Under these Curtin–Hammett conditions, the regioselectivity of the HAT is determined by the energy difference between the competing HAT transition states (**TS2a** and **TS2b**, $\Delta\Delta G^{\ddagger}$ = 2.7 kcal/mol). The overall barriers to the 1,6- and 1,5-HAT were calculated with respect to the lowest-energy conformer (**RC2b**): $\Delta G^{\ddagger}$ = 14.6 and 17.3 kcal/mol for 1,6-HAT (**TS2a**) and 1,5-HAT (**TS2b**), respectively.

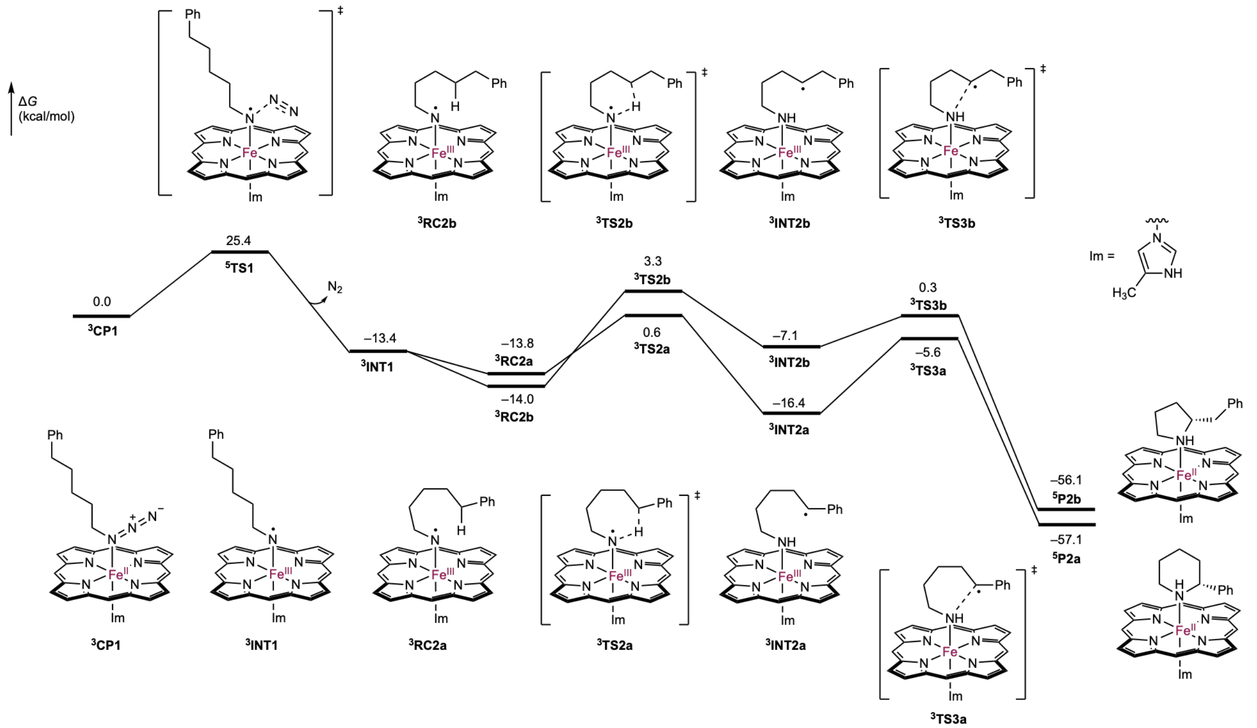

**Figure S12.** DFT-computed reaction energy profiles with a 5-methylimidazole axial ligand.

**Spin state energies of Fe–porphyrin complexes with a 5-methylimidazole axial ligand**

Singlet, triplet, quintet, and septet spin states of intermediates **CP1**, **INT1**, **RC2a**, **RC2b**, **INT2a**, **INT2b**, **P2a**, and **P2b** were located. For the singlet state of each species, both closed-shell singlet and open-shell singlet from broken-symmetry DFT calculations were calculated. For **CP1**, **INT1**, **RC2b**, **P2a**, and **P2b**, the closed-shell singlet wavefunction was stable and no open-shell singlet could be obtained, whereas open-shell singlet was obtained for **RC2a** and **INT2a**. Attempts to locate the singlet of the homobenzylic radical **INT2b** were not successful. Triplet was found to be the most stable spin state for **CP1**, **INT1**, **RC2a**, **RC2b**, **INT2a**, **INT2b**, **TS2a**, **TS2b**, **TS3a**, and **TS3b**. Quintet was found to be the most stable spin state for **TS1**, the piperidine product **P2a** and pyrrolidine product **P2b**. Although no spin crossover was observed for HAT in our study, the relatively small energy differences between triplet and quintet states for both 1,6- and 1,5-HAT transition states (**TS2a** and **TS2b**) are consistent with the previous computational study from Mondal that indicated a small triplet-quintet energy gap at the HAT transition states (*28*).

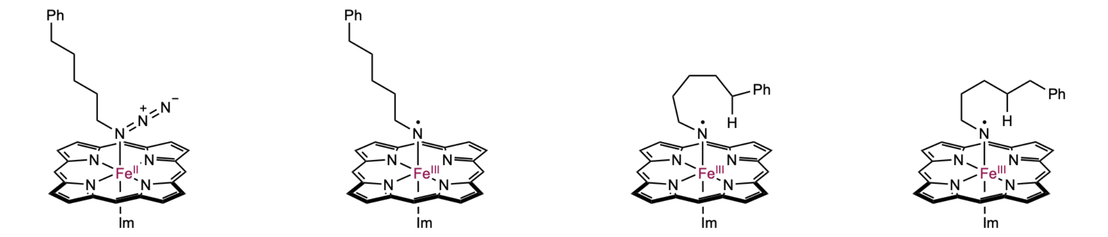

**CP1**

| Spin State | ΔG (kcal/mol) |
|---|---|
| Closed-shell singlet | 6.4 |
| Triplet | 0.0 |
| Quintet | 0.4 |
| Septet | 24.7 |

**INT1**

| Spin State | ΔG (kcal/mol) |
|---|---|
| Closed-shell singlet | 11.5 |
| Triplet | 0.0 |
| Quintet | 6.7 |
| Septet | 10.8 |

**RC2a**

| Spin State | ΔG (kcal/mol) |
|---|---|
| Open-shell singlet | 11.3 |
| Triplet | 0.0 |
| Quintet | 8.7 |
| Septet | 13.5 |

**RC2b**

| Spin State | ΔG (kcal/mol) |
|---|---|
| Closed-shell singlet | 8.0 |
| Triplet | 0.0 |
| Quintet | 7.2 |
| Septet | 13.3 |

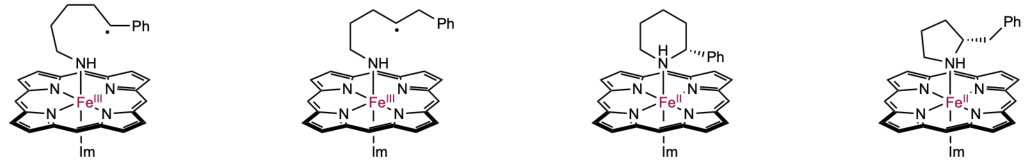

**INT2a**

| Spin State | ΔG (kcal/mol) |
|---|---|
| Open-shell singlet | 23.0 |
| Triplet | 0.0 |
| Quintet | 12.0 |
| Septet | 3.5 |

**INT2b**

| Spin State | ΔG (kcal/mol) |
|---|---|
| Singlet | not located |
| Triplet | 0.0 |
| Quintet | 20.4 |
| Septet | 3.4 |

**P2a**

| Spin State | ΔG (kcal/mol) |
|---|---|
| Closed-shell singlet | 5.4 |
| Triplet | 0.9 |
| Quintet | 0.0 |
| Septet | 23.0 |

**P2b**

| Spin State | ΔG (kcal/mol) |
|---|---|
| Closed-shell singlet | 1.4 |
| Triplet | 1.0 |
| Quintet | 0.0 |
| Septet | 21.7 |

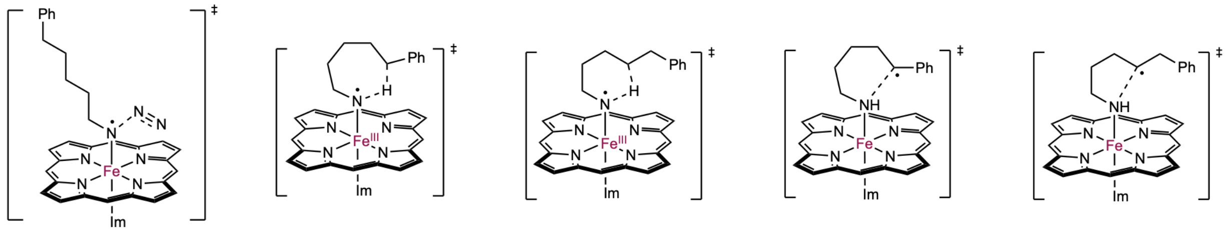

**TS1**

| Spin State | ΔG (kcal/mol) |
|---|---|
| Closed-shell singlet | 5.3 |
| Triplet | 0.4 |
| Quintet | 0.0 |
| Septet | 6.4 |

**TS2a**

| Spin State | ΔG (kcal/mol) |
|---|---|
| Closed-shell singlet | 8.9 |
| Triplet | 0.0 |
| Quintet | 0.3 |
| Septet | 6.8 |

**TS2b**

| Spin State | ΔG (kcal/mol) |
|---|---|
| Closed-shell singlet | 8.5 |
| Triplet | 0.0 |
| Quintet | 0.9 |
| Septet | 6.9 |

**TS3a**

| Spin State | ΔG (kcal/mol) |
|---|---|
| Singlet | not located |
| Triplet | 0.0 |
| Quintet | 2.6 |
| Septet | 8.0 |

**TS3b**

| Spin State | ΔG (kcal/mol) |
|---|---|
| Singlet | not located |
| Triplet | 0.0 |
| Quintet | 10.4 |
| Septet | 10.2 |

**Figure S13.** Spin state energies of intermediates and transition states in the C–H amination catalyzed by a model Fe–porphyrin complex with a 5-methylimidazole (Im) axial ligand. Gibbs free energies are respect to the most stable spin state of each structure.

**DFT-computed reaction energy profiles with a thioether axial ligand**

To model the reaction mechanism with the *de novo* designed **GDM-NT-401**, which has a methionine axial ligand, we computed the reaction energy profiles of the δ- and ε-C–H amination using a model iron porphyrin complex with a ethyl methyl thioether axial ligand and azide **1a** as the model substrate.

The reaction starts with an iron–azide complex **CP1'**, which undergoes nitrogen elimination through transition state **TS1'** to generate the iron–nitrene species **INT1'** with an energy barrier of 24.3 kcal/mol. Multiple conformers of the iron–nitrene species **INT1'** were located, including reactive conformers **RC2a'** and **RC2b'**, in which the ε- and δ-C–H bonds point towards the nitrene nitrogen, poised for 1,6- and 1,5-hydrogen atom transfer (HAT), respectively. From reactive conformer **RC2b'**, 1,5-HAT occurs through **TS2b'** with an energy barrier of 16.5 kcal/mol to generate the homobenzylic radical species **INT2b'**. The reactive conformer for 1,6-HAT (**RC2a'**), which places the ε-C–H in proximity to the nitrene nitrogen, is 0.9 kcal/mol less stable than **RC2b'**. From **RC2a'**, 1,6-HAT occurs through **TS2a'** to generate the benzylic radical species **INT2a'**. From the radical species **INT2a'** and **INT2b'**, C–N radical rebound through **TS3a'** and **TS3b'** forms the piperidine complex **P2a'** and pyrrolidine complex **P2b'**, respectively. The C–N radical rebound steps are both highly exergonic and irreversible. In the 1,6- and 1,5-HAT pathways, the Gibbs free energies of the C–N radical rebound transition states (**TS3a'** and **TS3b'**) is 0.3 and 4.2 kcal/mol lower than those of the HAT transition states (**TS2a'** and **TS2b'**). These results indicate that the HAT in both pathways are irreversible and thus regioselectivity-determining. Because of the relatively high barriers to the subsequent HAT, the alkyl chain conformational change of the Fe–nitrene intermediate is expected to occur much faster. Under these Curtin–Hammett conditions, the regioselectivity of the HAT is determined by the energy difference between the competing HAT transition states (**TS2a'** and **TS2b'**, $\Delta\Delta G^{\ddagger}$ = 3.2 kcal/mol). The overall barriers to the 1,6- and 1,5-HAT were calculated with respect to the lowest-energy conformer (**INT1'**): $\Delta G^{\ddagger}$ = 13.3 and 16.5 kcal/mol for 1,6-HAT (**TS2a'**) and 1,5-HAT (**TS2b'**), respectively.

The DFT calculations indicated that the reaction catalyzed by the Fe–porphyrin catalyst with the thioether axial ligand occurs via the same mechanism as that with the 5-methylimidazole axial ligand, and the computed regioselectivity is not significantly affected by the axial ligand ($\Delta\Delta G^{\ddagger}$ = 3.2 and 2.7 kcal/mol, respectively). These results, along with the classical MD simulations in this study, indicate that the enhanced regioselectivity for the *de novo* designed **GDM-NT-401** compared with the evolved ***Tm*Pgb-PIP-G7** is primarily due to the different active site pocket, rather than the electronic effects of the axial ligand.

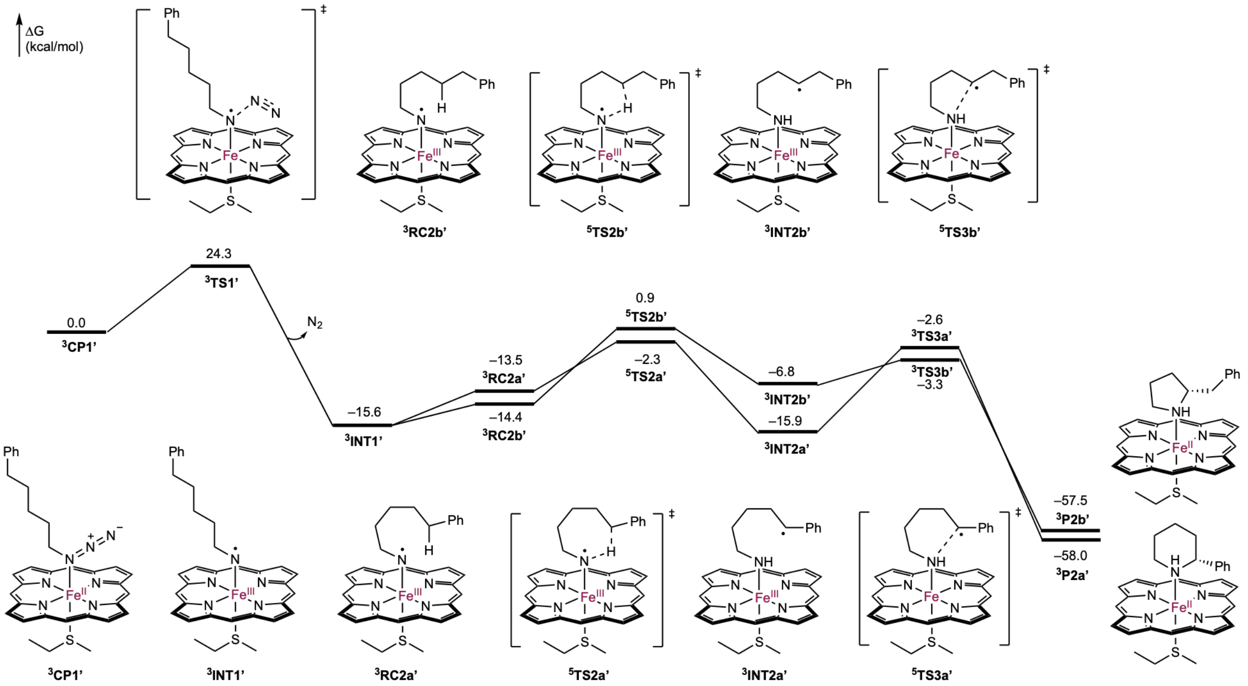

**Figure S14.** DFT-computed reaction energy profiles with a thioether axial ligand.

**Spin state energies of Fe–porphyrin complexes with a thioether axial ligand**

Singlet, triplet, quintet, and septet spin states of intermediates **CP1'**, **INT1'**, **RC2a'**, **RC2b'**, **P2a'**, and **P2b'** were located. For the singlet state of each species, both closed-shell singlet and open-shell singlet from broken-symmetry DFT calculations were calculated. For **CP1'**, **INT1'**, **RC2b'**, **RC2a'**, **P2a'**, and **P2b'**, the closed-shell singlet wavefunction was stable and no open-shell singlet could be obtained. Attempts to locate the singlet of the benzylic radical **INT2a'** and the homobenzylic radical **INT2b'** were not successful. Triplet was found to be the most stable spin state for **CP1'**, **INT1'**, **RC2a'**, **RC2b'**, **INT2a'**, **INT2b'**, **P2a'**, and **P2b'**. Quintet was found to be the most stable spin state for the 1,6- and 1,5-HAT transition states (**TS2a'** and **TS2b'**). The spin crossover prior to HAT is consistent with the previous computational study reported by Mondal with a neutral methanethiol axial ligand (*28*).

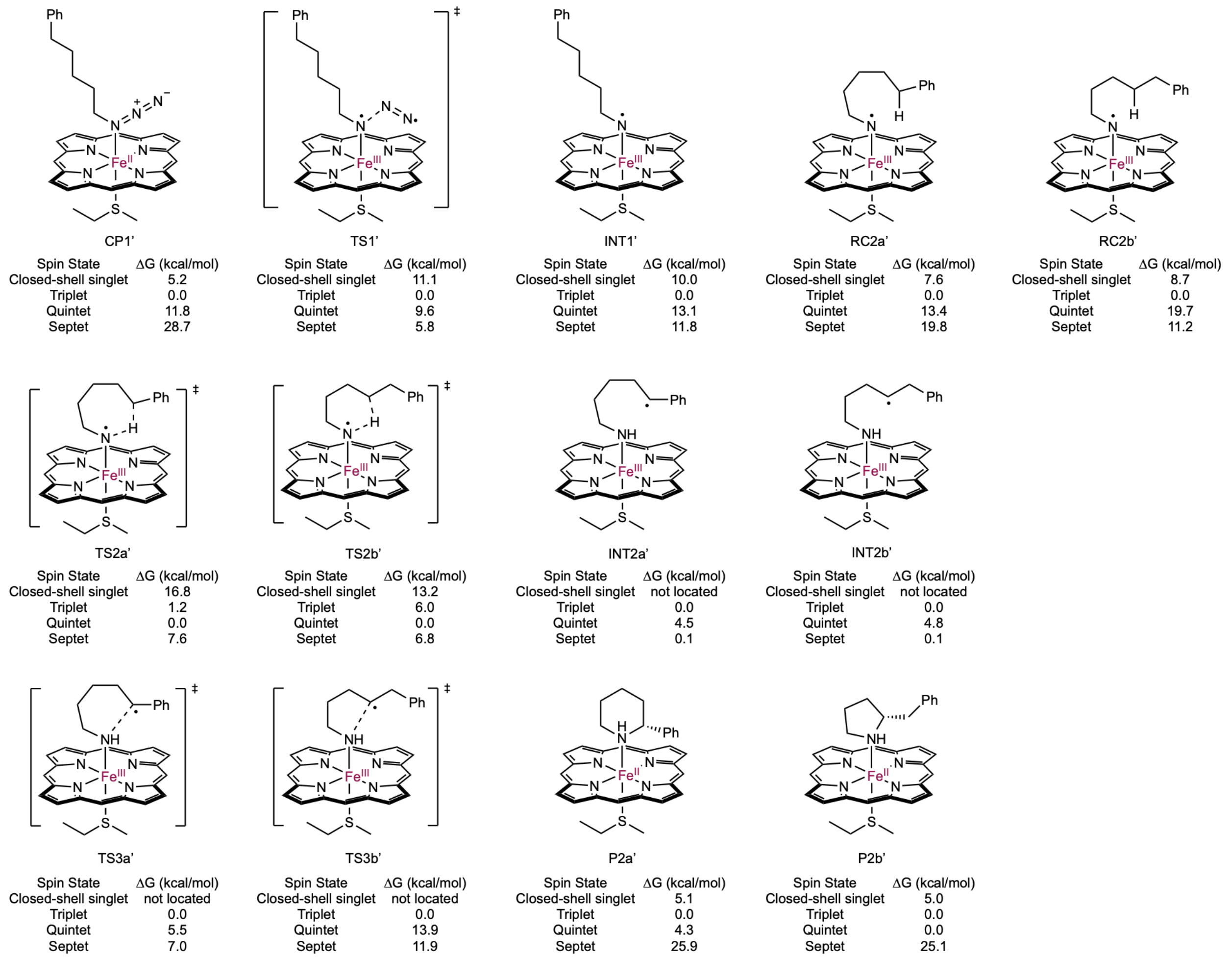

CP1'

| Spin State | ΔG (kcal/mol) |
|---|---|
| Closed-shell singlet | 5.2 |
| Triplet | 0.0 |
| Quintet | 11.8 |
| Septet | 28.7 |

TS1'

| Spin State | ΔG (kcal/mol) |
|---|---|
| Closed-shell singlet | 11.1 |
| Triplet | 0.0 |
| Quintet | 9.6 |
| Septet | 5.8 |

INT1'

| Spin State | ΔG (kcal/mol) |
|---|---|
| Closed-shell singlet | 10.0 |
| Triplet | 0.0 |
| Quintet | 13.1 |
| Septet | 11.8 |

RC2a'

| Spin State | ΔG (kcal/mol) |
|---|---|
| Closed-shell singlet | 7.6 |
| Triplet | 0.0 |
| Quintet | 13.4 |
| Septet | 19.8 |

RC2b'

| Spin State | ΔG (kcal/mol) |
|---|---|
| Closed-shell singlet | 8.7 |
| Triplet | 0.0 |
| Quintet | 19.7 |
| Septet | 11.2 |

TS2a'

| Spin State | ΔG (kcal/mol) |
|---|---|
| Closed-shell singlet | 16.8 |
| Triplet | 1.2 |
| Quintet | 0.0 |
| Septet | 7.6 |

TS2b'

| Spin State | ΔG (kcal/mol) |
|---|---|
| Closed-shell singlet | 13.2 |
| Triplet | 6.0 |
| Quintet | 0.0 |
| Septet | 6.8 |

INT2a'

| Spin State | ΔG (kcal/mol) |
|---|---|
| Closed-shell singlet | not located |
| Triplet | 0.0 |
| Quintet | 4.5 |
| Septet | 0.1 |

INT2b'

| Spin State | ΔG (kcal/mol) |
|---|---|
| Closed-shell singlet | not located |
| Triplet | 0.0 |
| Quintet | 4.8 |
| Septet | 0.1 |

TS3a'

| Spin State | ΔG (kcal/mol) |
|---|---|
| Closed-shell singlet | not located |
| Triplet | 0.0 |
| Quintet | 5.5 |
| Septet | 7.0 |

TS3b'

| Spin State | ΔG (kcal/mol) |
|---|---|
| Closed-shell singlet | not located |
| Triplet | 0.0 |
| Quintet | 13.9 |
| Septet | 11.9 |

P2a'

| Spin State | ΔG (kcal/mol) |
|---|---|
| Closed-shell singlet | 5.1 |
| Triplet | 0.0 |
| Quintet | 4.3 |
| Septet | 25.9 |

P2b'

| Spin State | ΔG (kcal/mol) |
|---|---|
| Closed-shell singlet | 5.0 |
| Triplet | 0.0 |
| Quintet | 0.0 |
| Septet | 25.1 |

**Figure S15.** Spin state energies of intermediates and transition states in the C–H amination catalyzed by a model Fe–porphyrin complex with a thioether axial ligand. Gibbs free energies are respect to the most stable spin state of each structure.

**Computational Details – Classical molecular dynamics simulations**

Classical molecular dynamics (MD) simulations were performed to investigate active site residue–substrate interactions with the Fe–nitrene intermediate prior to the HAT. The Fe–nitrene derived from azide **1a** was used in the classical MD simulations. The starting structures of the alkyl nitrene intermediate in ***Tm*Pgb-PIP-G7** and **GDM-NT-401** were obtained from co-folding models of each protein with the haem–nitrene ligand, generated using Chai-1 (*29*) with the protein sequence given in one-letter code and the haem–nitrene ligand specified by the following SMILES string: [CH2]=[CH][C]1=[C]2[CH]=[C]3[C]([CH3])=[C]([CH]=[CH2])[C]4=[N]3[Fe+4]35([N]6=[C]([C]([CH2][CH2][C]([OH])=[O])=[C]([CH3])[C]6=[CH][C](=[C]1[CH3])[N]23)[CH]=[C]1[C]([CH2][CH2][C]([OH])=[O])=[C]([CH3])[C](=[CH]4)[N]51)=[N][CH2][CH2][CH2][CH2][CH2][C]1=[CH][CH]=[CH][CH]=[CH]1. Because Chai-1 returns heavy-atom coordinates only, hydrogen atoms were added using the H++ server (*30*), and the resulting complex was used as the starting structure for the MD simulations described below.

Classical MD simulations were carried out using the pmemd module (*31*) of the GPU-accelerated Amber 24 software (*32*). The Amber ff19SB force field (*33*) was used in all classical MD simulations. Force field parameters for Fe and surrounding atoms and bonds in the haem–nitrene complex were generated using the MCPB.py module (*34*). The second-generation general Amber force field (gaff2) (*35*) was used for other atoms in the haem–nitrene intermediate. Using the Merz–Singh–Kollman scheme (*36*–*37*), RESP charge fitting (*38*) on electrostatic potential generated at the B3LYP/6-31G(d) level of theory was performed to generate partial charges at triplet spin state for the Fe–nitrene. The enzyme was then put into a solvated cuboid box under periodic boundary conditions using the TIP3P water model (*39*). The minimum distance between the enzyme surface and the edge of the water box was set to 10 Å. Water molecules were treated with the SHAKE algorithm (*40*). The system was neutralized by adding $Na^+$ counterions. Long-range electrostatics were calculated using the particle-mesh-Ewald method (*41*). Lennard-Jones and electrostatic interaction cut-offs were set to 12 Å.

The classical MD simulations were performed in several steps. Three independent replicas of 500 ns MD simulations were performed for the Fe–nitrene intermediate of the ***Tm*Pgb-PIP-G7** and **GDM-NT-401** variants. A 30,000-step energy minimization was first performed with positional restraints for the protein by applying a force constant of 500 kcal·mol$^{-1}$·Å$^{-2}$. Next, the system was gradually heated without any restraint from 0 K to 300 K in 200 ps, which was followed by an equilibration using the isothermal–isobaric ensemble (NPT) in the next 6 ns. Finally, 500 ns production MD simulations were performed using the same conditions as the equilibration with a time step of 2 fs without any restraint. After the MD simulations, clustering analysis was carried out using the CPPTRAJ module (*42*) to identify the most populated structure from the classical MD simulations of all three replicas. Clustering analysis was performed using the k-means algorithm (*43*) (10 clusters, maximum 500 iterations, sieving every 10th frame), with the RMSD of the protein backbone atoms (N, Cα and C) as the distance metric.

To further probe enzyme–substrate interactions at reactive geometries near the HAT transition state, restrained MD simulations of the Fe–nitrene intermediate were performed to sample the near attack conformations (NACs) (*44*) prior to HAT in ***Tm*Pgb-PIP-G7** and **GDM-NT-401**. The restrained MD simulations were performed following the same four-step protocol described above, with a harmonic distance restraint (2.9–3.5 Å, force constant 100 kcal·mol$^{-1}$·Å$^{-2}$) applied between the nitrene nitrogen and the centroid of the δ- and ε-carbon atoms. This distance was chosen according to the corresponding N···C distance in the DFT-optimized transition state structures. Three independent replicas of the restrained MD simulations were performed for each enzyme.

**Unrestrained classical MD simulations of the Fe–nitrene intermediate in *Tm*Pgb-PIP-G7**

To investigate the preferred binding poses of the substrate and the potential interaction modes within the active site, three independent replicas of 500 ns MD simulations were performed for the Fe–nitrene intermediate in ***Tm*Pgb-PIP-G7**. The major binding pose of the alkyl nitrene intermediate in ***Tm*Pgb-PIP-G7** is shown in the main text (Figure 5b, left). The phenyl group of the nitrene alkyl chain is accommodated in a hydrophobic pocket formed by Y60, W59 and F144, engaging in stabilising π···π and C–H···π interactions. These interactions were corroborated by analysing π···π and C–H···π distances along the trajectories. The Cβ of the alkyl chain is 4.7 ± 0.4 Å from the centroid of the aromatic ring of the Y60 side chain, indicating a favorable C–H···π interaction between the nitrene alkyl chain and Y60 (Figure S16). The centroid of the substrate phenyl ring is 5.0 ± 0.7 Å from that of the W59 pyrrole ring (Figure S17) and 4.9 ± 0.8 Å from that of the F144 phenyl ring (Figure S18), indicating stabilising π···π interactions. These persistent interactions hold the nitrene alkyl chain in an extended geometry, with most of the rotatable C($sp^3$)–C($sp^3$) bonds adopting the *trans* conformation (most populated $\theta_2$, $\theta_3$, $\theta_4 \approx 180°$; Figures S19–S22).

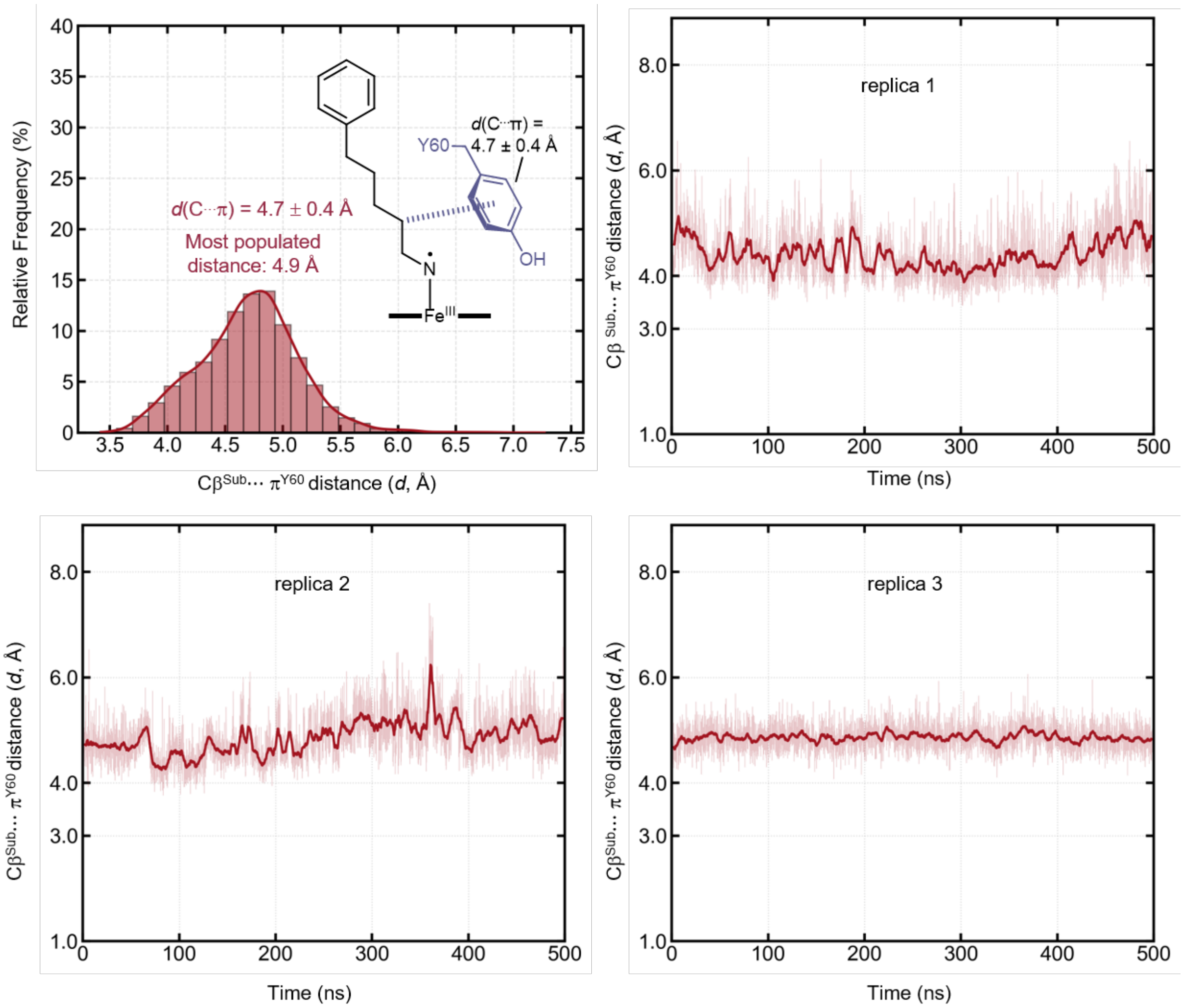

**Figure S16.** Distance between the centroid of the benzene ring of the Y60 side chain ($\pi^{Y60}$) and $C_\beta$ atom of nitrene observed in unrestrained MD simulations with ***Tm*Pgb-PIP-G7**.

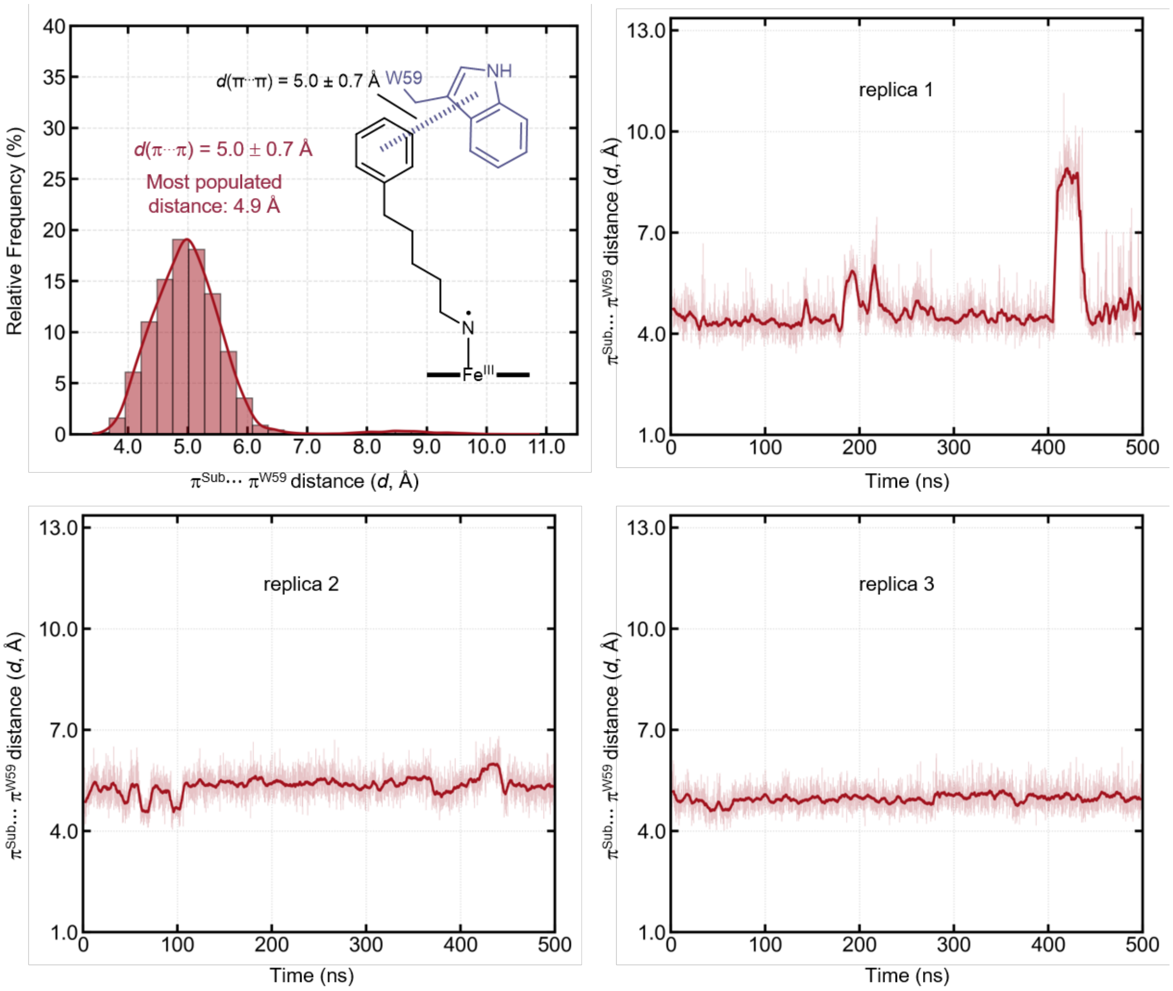

**Figure S17.** Distance between the centroid of the benzene of the substrate ($\pi^{Sub}$) and the centroid of the pyrrole ring of W59 side chain ($\pi^{W59}$) observed in unrestrained MD simulations with ***TmPgb*-PIP-G7**.

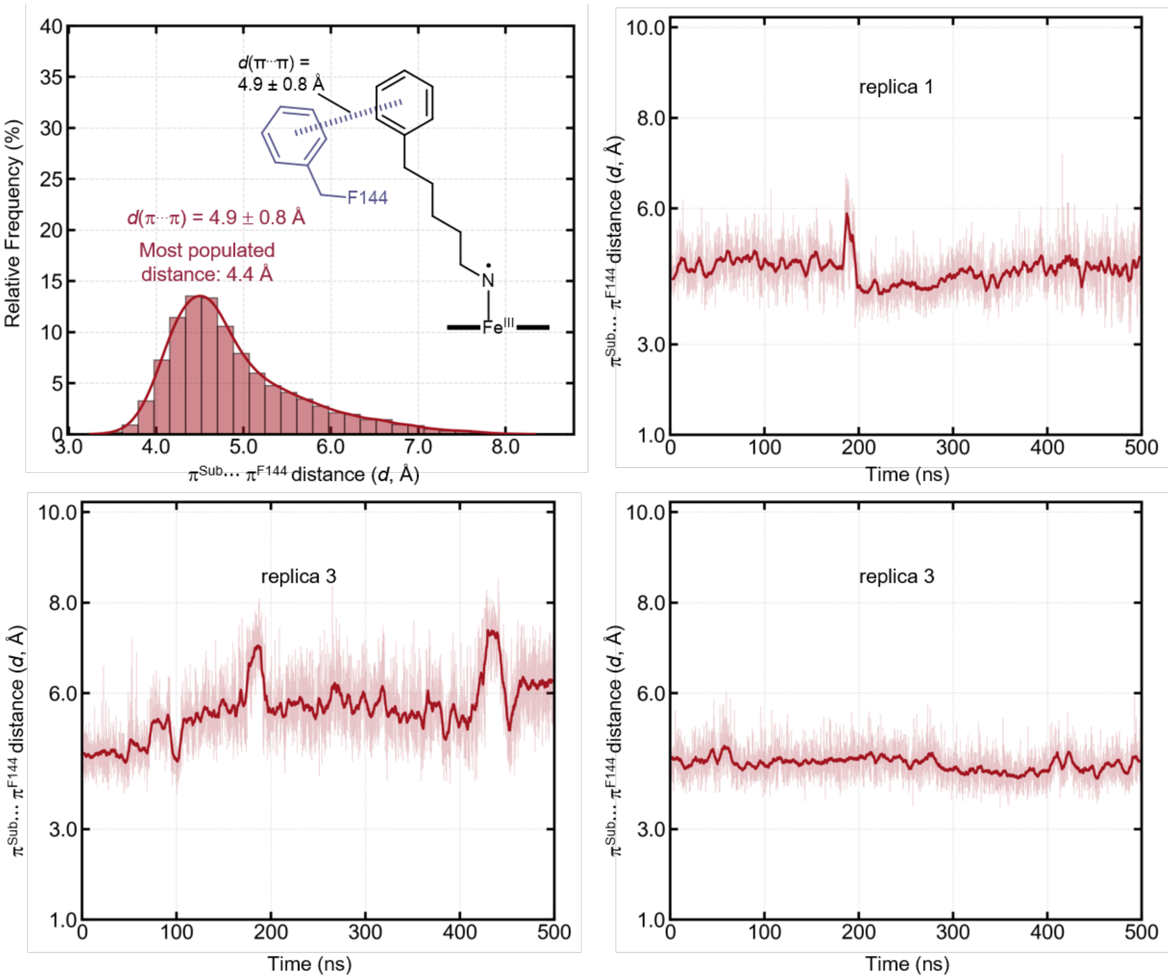

**Figure S18**. Distance between the centroid of the benzene ring of the substrate ($\pi^{Sub}$) and the centroid of the benzene ring of F144 side chain ($\pi^{F144}$) observed in unrestrained MD simulations with ***Tm*Pgb-PIP-G7**.

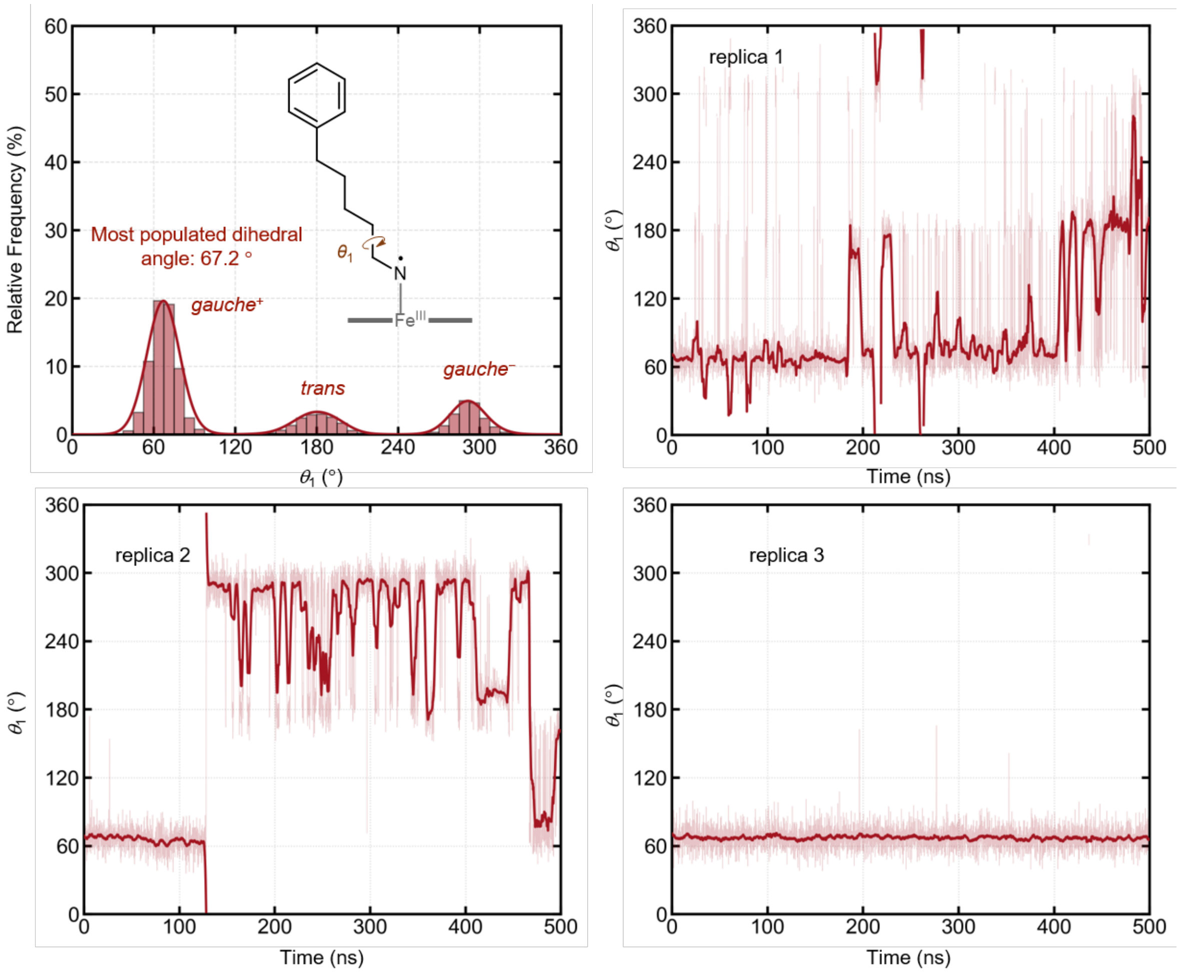

**Figure S19**. Dihedral angle $\theta_1$ (NCαCβCγ) of the alkyl nitrene intermediate observed in unrestrained MD simulations with ***Tm*****Pgb-PIP-G7**.

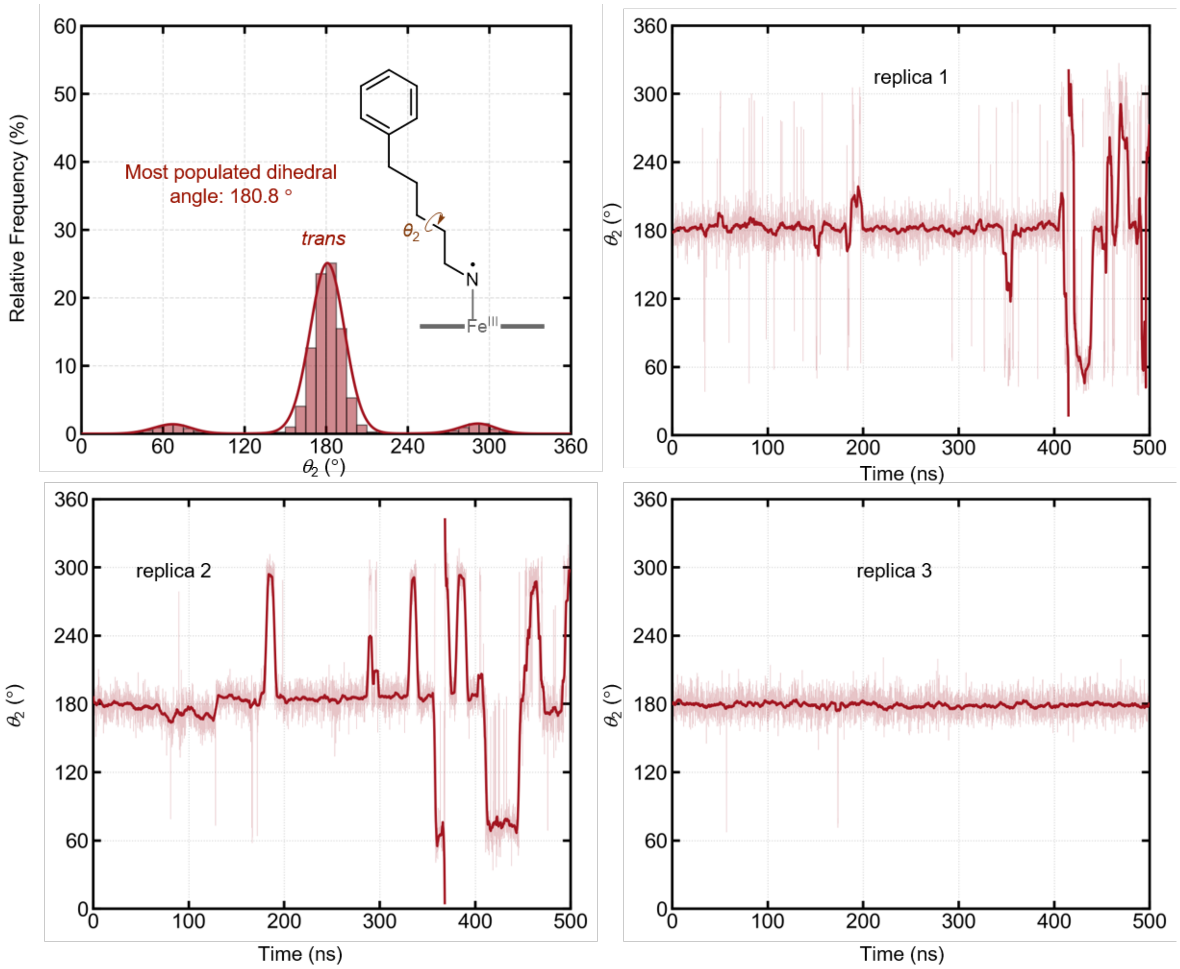

**Figure S20.** Dihedral angle $\theta_2$ (CαCβCγCδ) of the alkyl nitrene intermediate observed in unrestrained MD simulations with ***Tm*****Pgb-PIP-G7**.

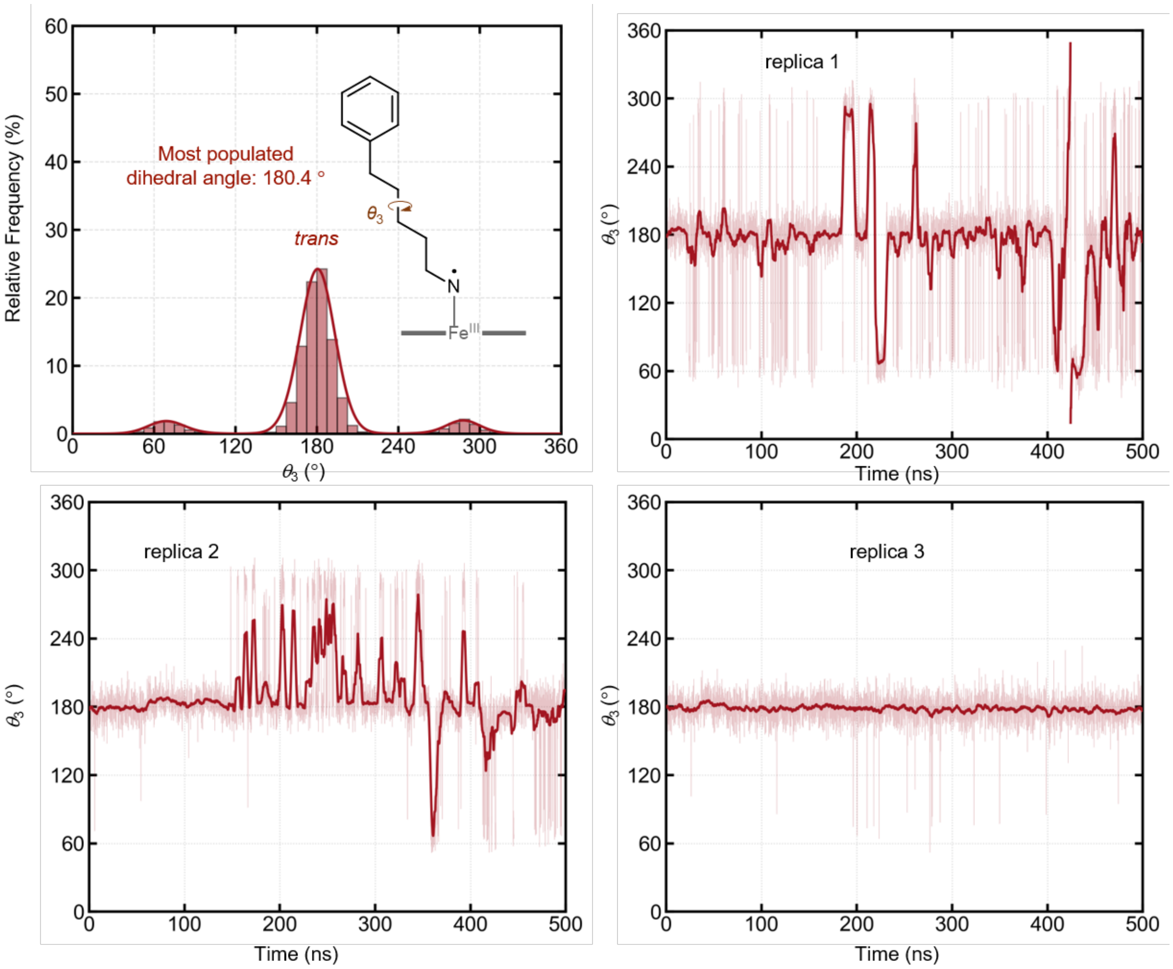

**Figure S21**. Dihedral angle $\theta_3$ (CβCγCδCε) of the alkyl nitrene intermediate observed in unrestrained MD simulations with ***Tm*Pgb-PIP-G7**.

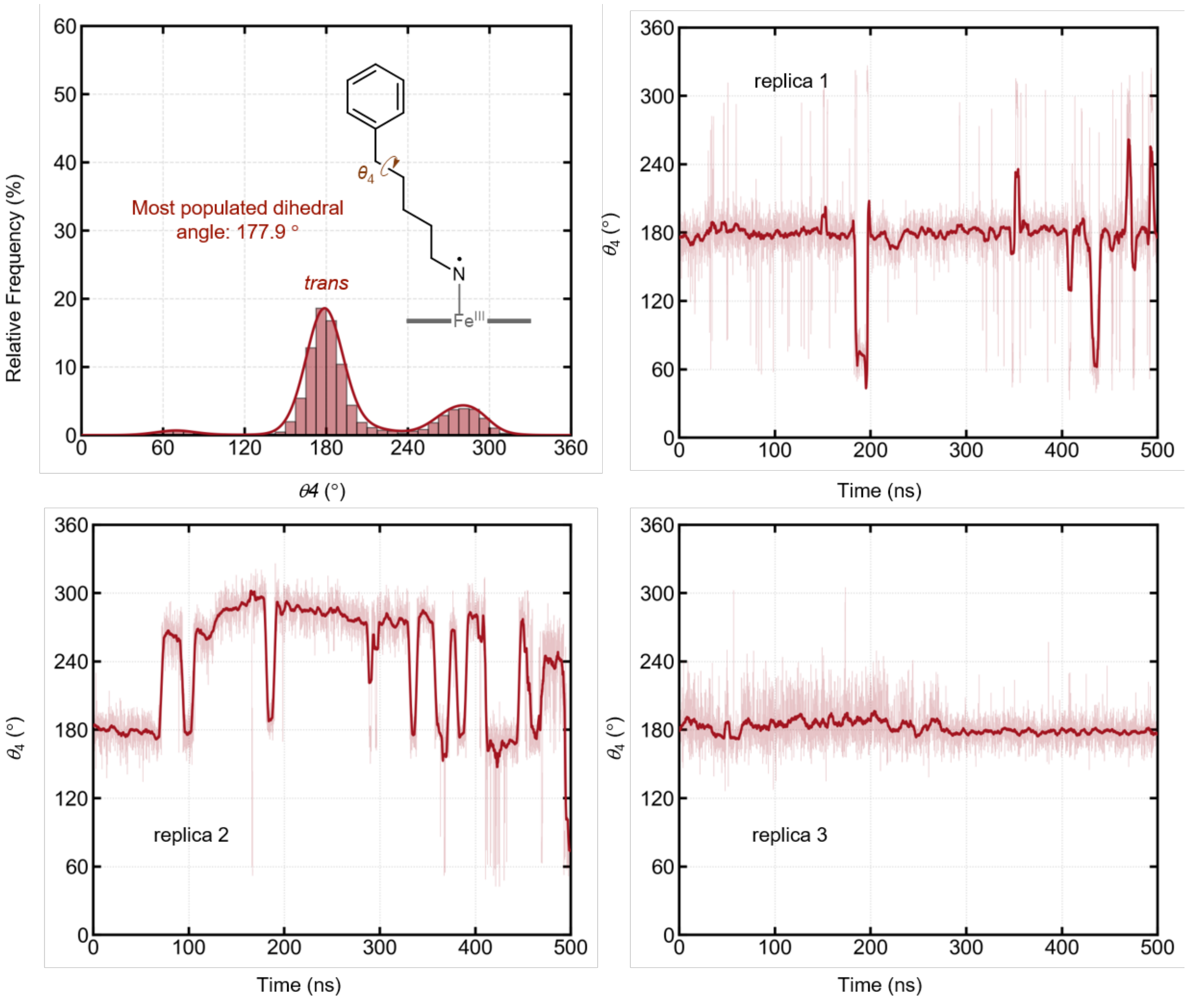

**Figure S22**. Dihedral angle $\theta_4$ (CγCδCεCphenyl) of the alkyl nitrene intermediate observed in unrestrained MD simulations with ***Tm*Pgb-PIP-G7**.

**Unrestrained classical MD simulations of the Fe–nitrene intermediate in GDM-NT-401**

To investigate the preferred binding poses of the substrate and the interaction modes within the active site, three independent replicas of 500 ns MD simulations were performed for the Fe–nitrene intermediate in **GDM-NT-401**. The predominant binding pose of the alkyl nitrene intermediate in **GDM-NT-401** is shown in the main text (Figure 5b, right). The Cδ of L20 forms a persistent C–H···π interaction with the substrate phenyl ring, with a mean Cδ···phenyl distance of 4.0 ± 0.4 Å (Figure S23). The side chains of V28, L19, and L16 create a hydrophobic pocket that enforces a bent, reactive conformation of the alkyl chain (Figures S24–S27).

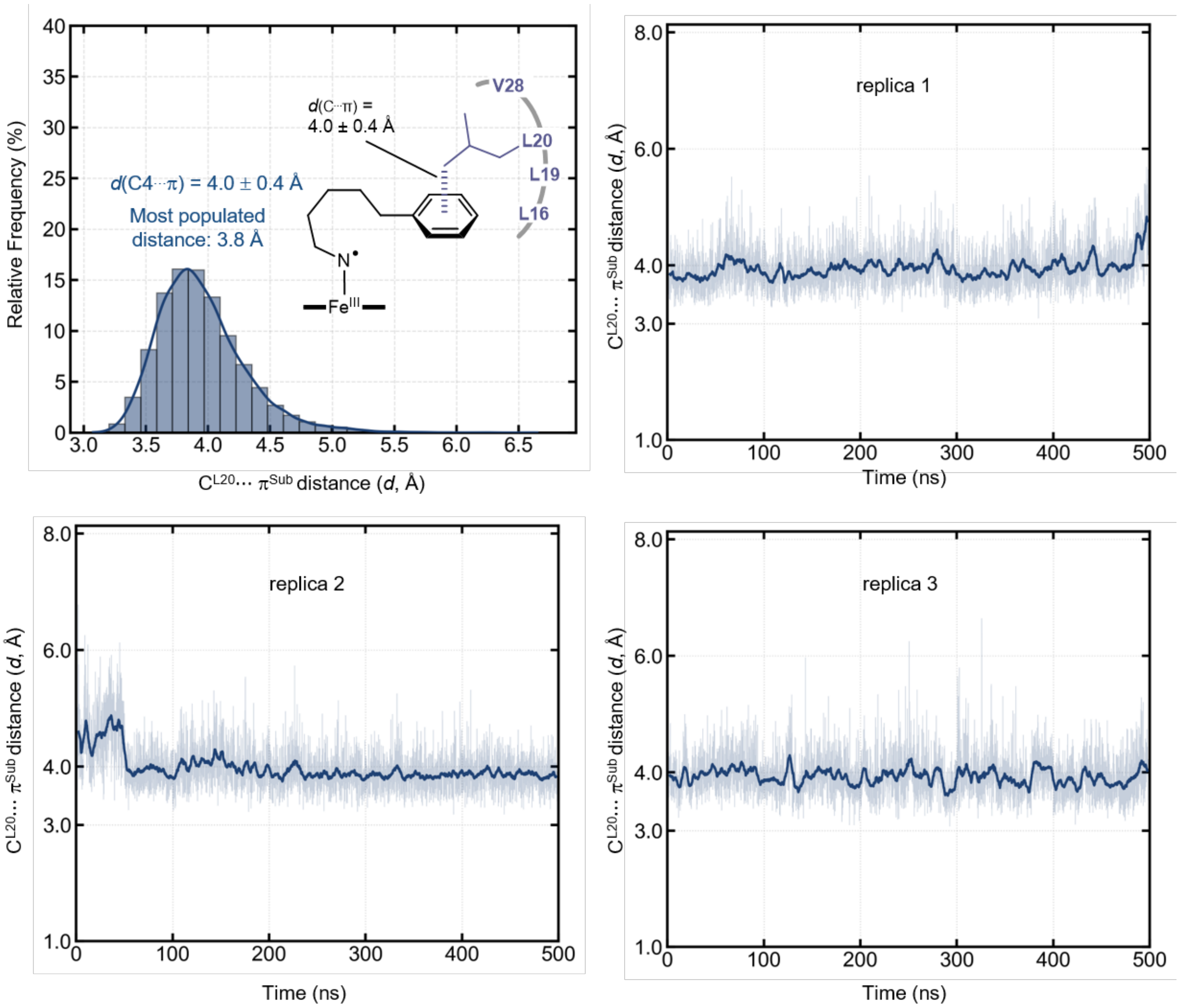

**Figure S23**. Distance between the centroid of the benzene of the substrate ($\pi^{Sub}$) and $C_\delta$ atom of L20 observed in unrestrained MD simulations with **GDM-NT-401**.

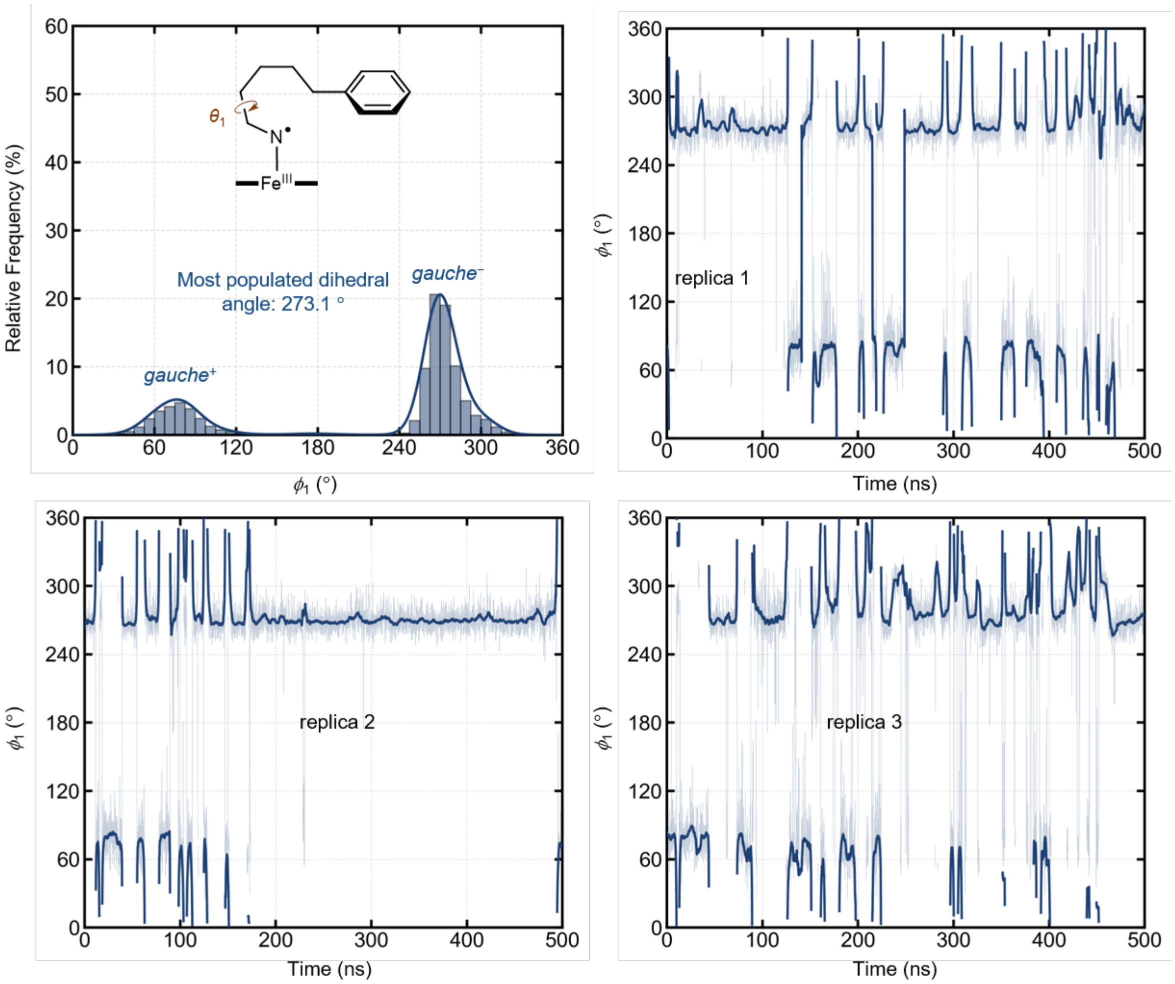

**Figure S24.** Dihedral angle $\theta_1$ (NCαCβCγ) of the alkyl nitrene intermediate observed in unrestrained MD simulations with **GDM-NT-401**.

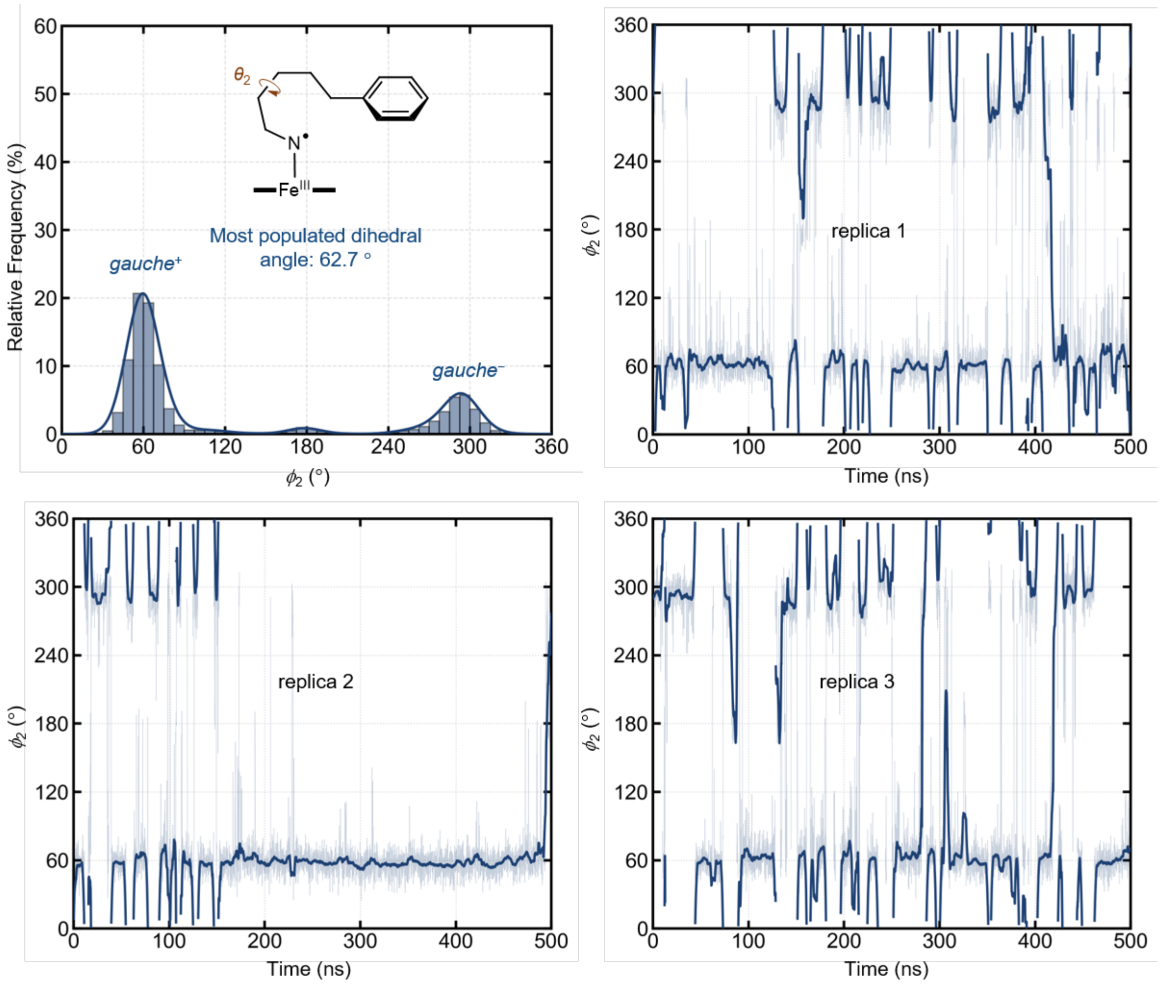

**Figure S25.** Dihedral angle $\theta_2$ (CαCβCγCδ) of the alkyl nitrene intermediate observed in unrestrained MD simulations with **GDM-NT-401**.

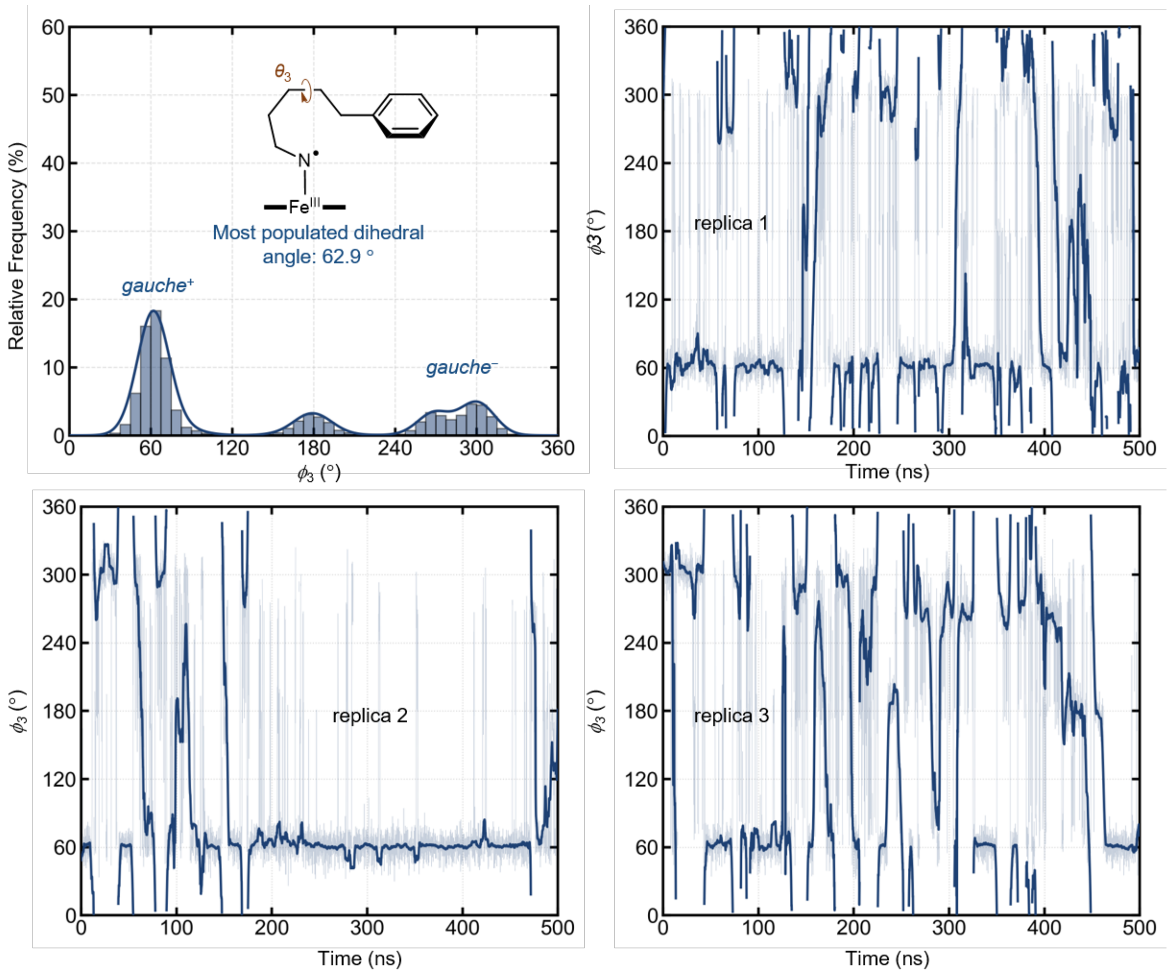

**Figure S26.** Dihedral angle $\theta_3$ (CβCγCδCε) of the alkyl nitrene intermediate observed in unrestrained MD simulations with **GDM-NT-401**.

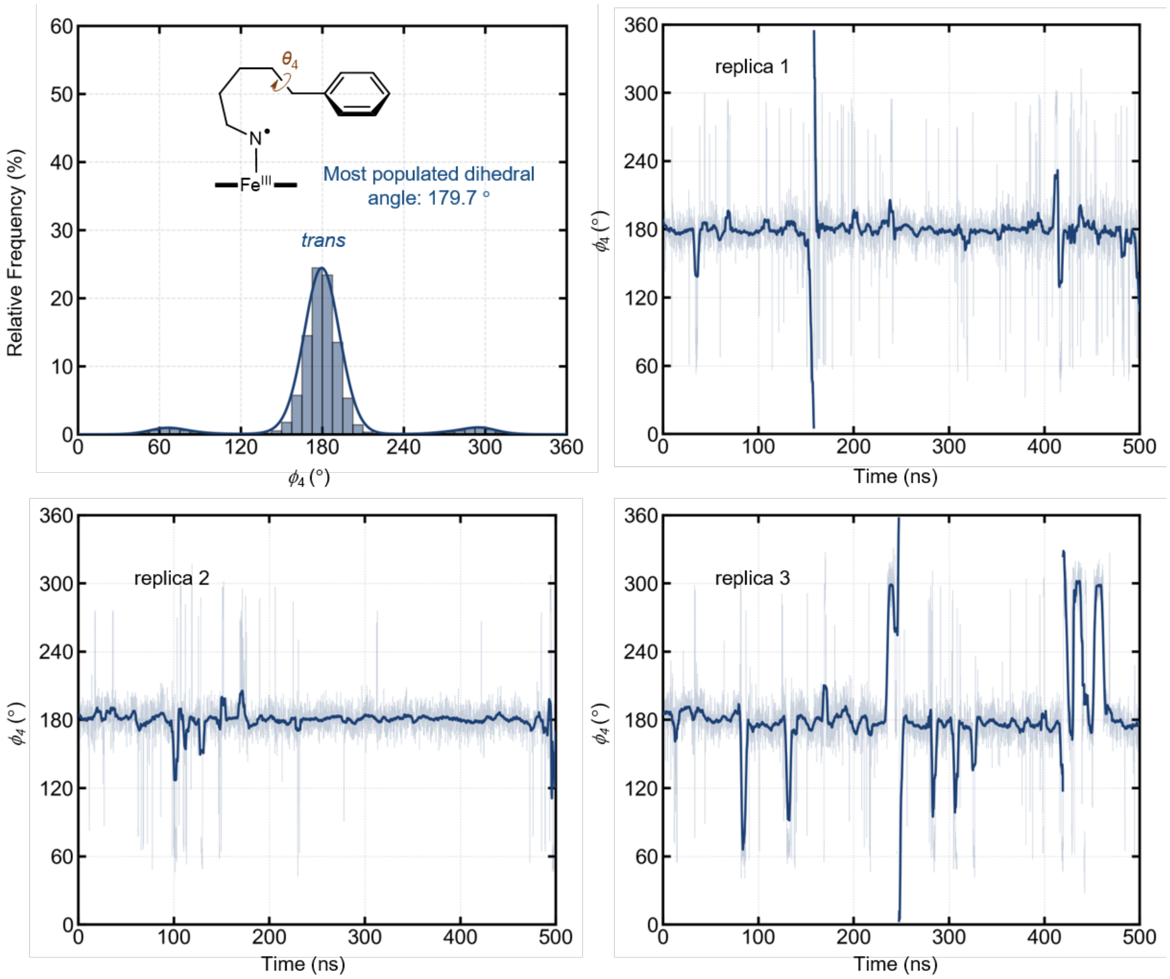

**Figure S27.** Dihedral angle $\theta_4$ (CγCδCεCphenyl) of the alkyl nitrene intermediate observed in unrestrained MD simulations with **GDM-NT-401**.

**Restrained classical MD simulations of the near attack conformations prior to Fe–nitrene HAT in *Tm*Pgb-PIP-G7 and GDM-NT-401**

To further investigate the enzyme–substrate interactions at reactive geometries near the HAT transition state, a distance restraint (2.9–3.5 Å, force constant 100 kcal $mol^{-1}$ $Å^{-2}$) was applied between the nitrene nitrogen and the centroid of the δ- and ε-carbon atoms. In these restrained near-attack conformation (NAC) simulations, ***Tm*Pgb-PIP-G7** populated 1,6-HAT (ε-C–H) geometries in 32.3% and 1,5-HAT (δ-C–H) geometries in 2.8% of the frames (Figure S28a, left), whereas **GDM-NT-401** reached 42.4% and 2.1%, respectively (Figure S28a, right), consistent with the higher regioselectivity of **GDM-NT-401**. The major near-attack conformations in **GDM-NT-401** and ***Tm*Pgb-PIP-G7** are shown in Figure S28b and S28c, respectively. For **GDM-NT-401**, the substrate alkyl chain binding pose is essentially unchanged between the unrestrained and restrained simulations (Figure S28b), because the alkyl chain is already preorganised into the reactive bent geometry in the unrestrained MD. The C–H···π interaction between L20 side chain and the phenyl group of the substrate is maintained under the restraint, with essentially the same centroid–centroid distance as in the unrestrained simulation (4.0 ± 0.4 and 4.0 ± 0.3 Å, respectively; Figure S29). Consistent with this, the per-residue RMSF changes little upon introducing the restraint, and is in fact slightly lower in the restrained than in the unrestrained simulation, possibily induced by the reduced substrate flexibility (Figure S28d, left). By contrast, in ***Tm*Pgb-PIP-G7** the nitrene alkyl chain adopts an extended geometry in the unrestrained simulations, so reaching the HAT transition state-like bent geometry under the restraint requires substantial reorganisation (Figure S28c). The less complementary shape of the ***Tm*Pgb-PIP-G7** active site with the HAT transition state is reflected in the RMSF, which increases upon applying the restraint relative to the unrestrained simulations (Figure S28d, right). Moreover, the conformational change disrupts the favourable interactions of the Y60 and W59 aromatic side chains with the alkyl chain of the nitrene, and weakens the remaining F144–substrate π···π interaction, whose centroid–centroid distance increases from 4.6 ± 0.6 Å in the unrestrained simulations to 5.4 ± 0.7 Å under restraint (Figure S30).

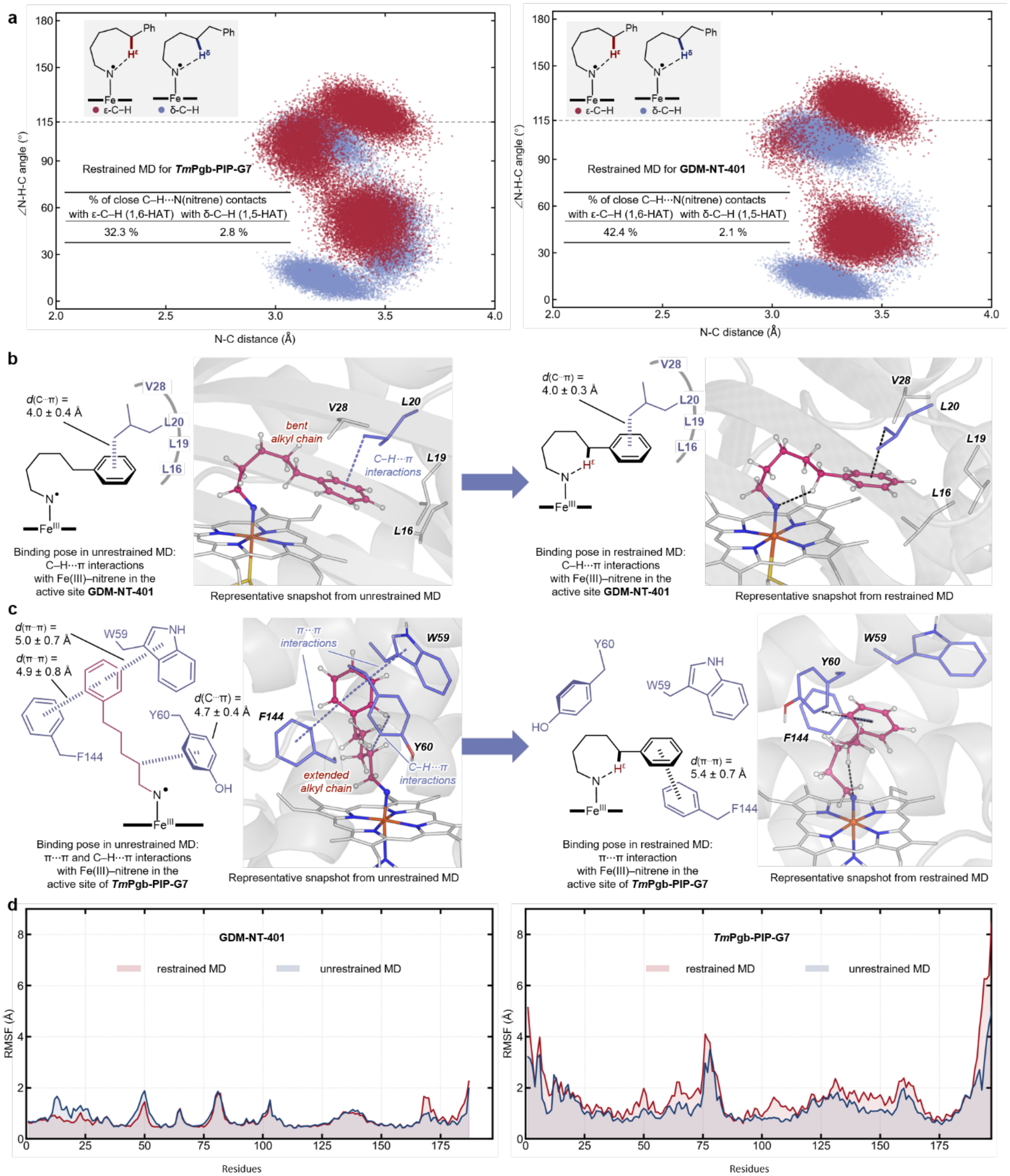

**Figure S28.** **a,** Close contacts between the nitrene N atom and the ε- and δ-C–H bonds observed in the restrained MD simulations of ***Tm*Pgb-PIP-G7** (left) and **GDM-NT-401** (right). **b,** Representative binding pose of the alkyl nitrene intermediate in the active site of **GDM-NT-401** (left, unrestrained MD; right, restrained MD). **c,** Representative binding pose of the alkyl nitrene intermediate in the active site of ***Tm*Pgb-PIP-G7** (left, unrestrained MD; right, restrained MD). **d,** RMSF profiles of the backbone $C_\alpha$ atoms for **GDM-NT-401** (restrained vs unrestrained MD) and ***Tm*Pgb-PIP-G7** (restrained vs unrestrained MD).

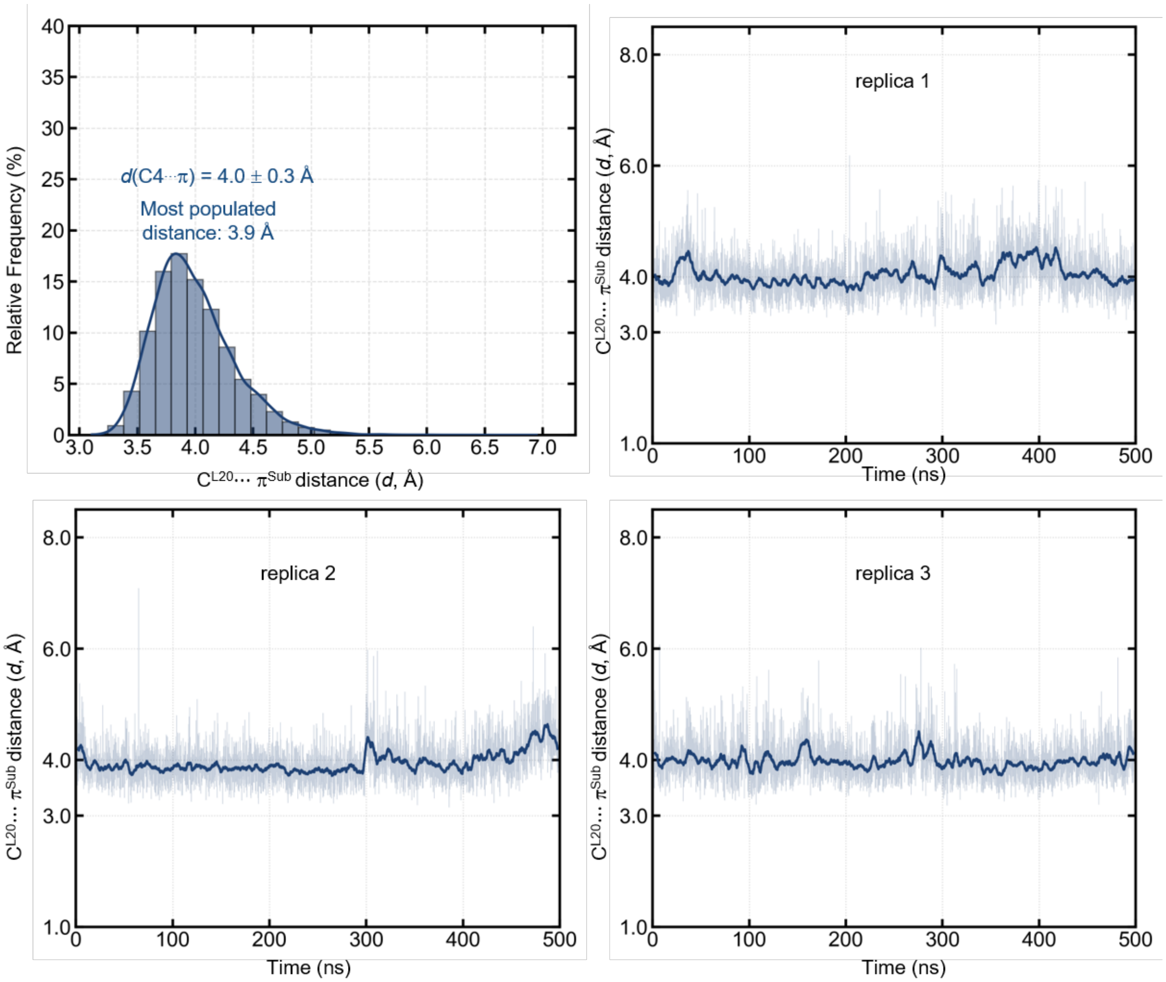

**Figure S29.** Distance between the centroid of the benzene ring of the substrate ($\pi^{Sub}$) and the $C_\delta$ atom of L20 in restrained MD simulations with **GDM-NT-401**.

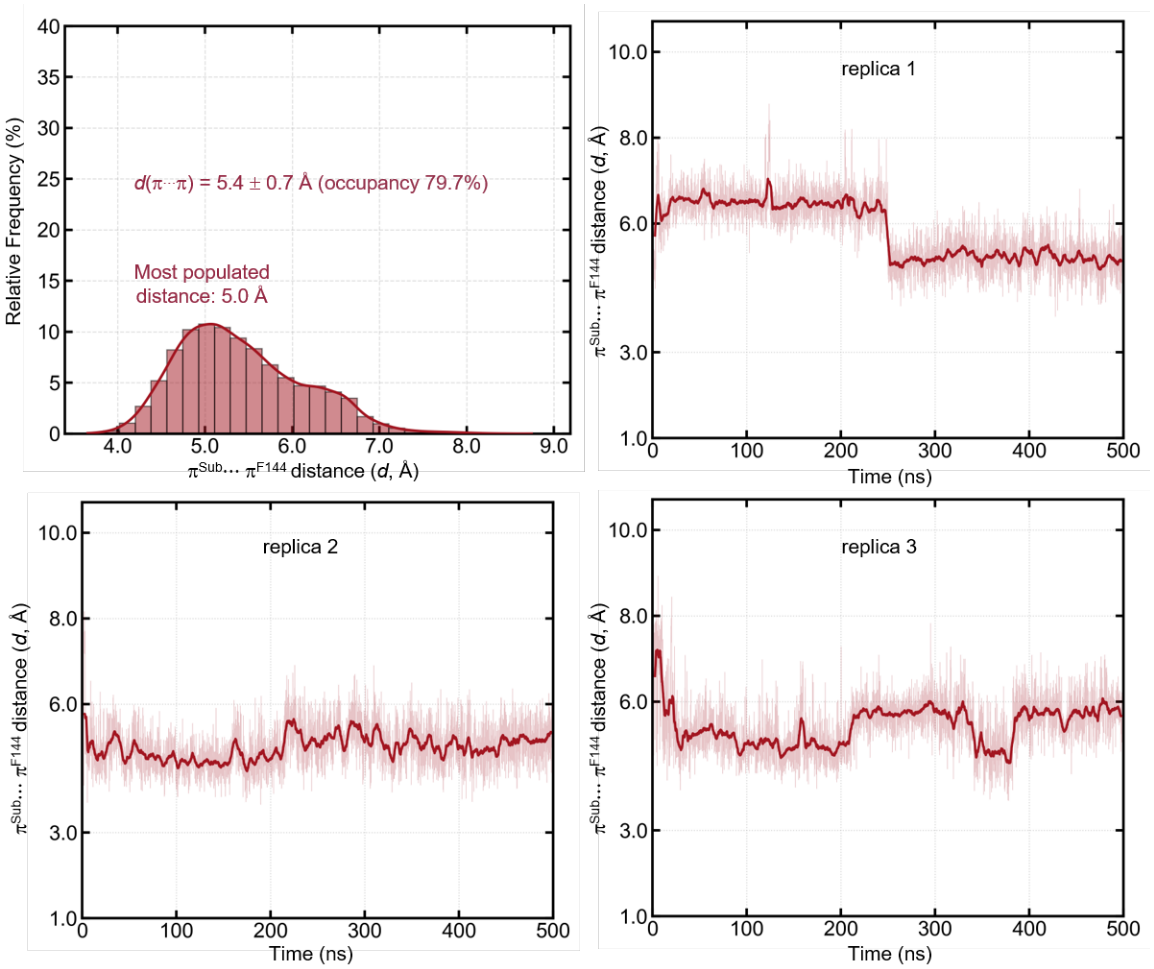

**Figure S30.** Distance between the centroid of the benzene ring of the substrate ($\pi^{Sub}$) and the centroid of the benzene ring of F144 side chain ($\pi^{F144}$) in restrained MD simulations with ***TmPgb*-PIP-G7**.

#### DFT-calculated energies of intermediates and transition states

Cartesian coordinates of all optimized geometries are provided in a separate file in the .xyz format.

**Table S26**. Calculated total (SCF) energies, enthalpies, and Gibbs free energies of intermediates and transition states.

| | Level of theory: B3LYP-D3(BJ)/def2TZVP/SMD($Et_2O$)//B3LYP-D3(BJ)/SDD-6-31G(d) | | |
|---|---|---|---|
| | SCF energy (a.u.) | Enthalpy (a.u.) | Gibbs Free energy (a.u.) |
| **CP1-quintet** | -3111.101532 | -3110.436652 | -3110.558096 |
| **CP1-septet** | -3111.060722 | -3110.397338 | -3110.519353 |
| **CP1-singlet** | -3111.101361 | -3110.434284 | -3110.548611 |
| **CP1-triplet** | -3111.100184 | -3110.434326 | -3110.558762 |
| **INT1-quintet** | -3001.522783 | -3000.871176 | -3000.987808 |
| **INT1-septet** | -3001.511573 | -3000.861297 | -3000.981276 |
| **INT1-singlet** | -3001.525419 | -3000.871093 | -3000.980237 |
| **INT1-triplet** | -3001.537833 | -3000.884628 | -3000.998551 |
| **INT2a-quintet** | -3001.521609 | -3000.871688 | -3000.984300 |
| **INT2a-septet** | -3001.534966 | -3000.884968 | -3000.997768 |

| | | | |
|---|---|---|---|
| **INT2a-singlet** | -3001.513399 | -3000.860142 | -3000.966810 |
| **INT2a-triplet** | -3001.548196 | -3000.894246 | -3001.003424 |
| **INT2b-quintet** | -3001.492097 | -3000.841849 | -3000.955961 |
| **INT2b-septet** | -3001.518630 | -3000.869280 | -3000.983126 |
| **INT2b-triplet** | -3001.531464 | -3000.878079 | -3000.988548 |
| **P2a-quintet** | -3001.613515 | -3000.958087 | -3001.068240 |
| **P2a-septet** | -3001.578180 | -3000.923476 | -3001.031571 |
| **P2a-singlet** | -3001.616195 | -3000.958149 | -3001.059685 |
| **P2a-triplet** | -3001.613808 | -3000.957223 | -3001.066860 |
| **P2b-quintet** | -3001.610559 | -3000.955452 | -3001.066702 |
| **P2b-septet** | -3001.576587 | -3000.922237 | -3001.032105 |
| **P2b-singlet** | -3001.618598 | -3000.959315 | -3001.064489 |
| **P2b-triplet** | -3001.609119 | -3000.953114 | -3001.065095 |
| **RC2a-quintet** | -3001.525372 | -3000.873481 | -3000.985267 |
| **RC2a-septet** | -3001.514601 | -3000.863999 | -3000.977697 |
| **RC2a-singlet** | -3001.525233 | -3000.871133 | -3000.981141 |
| **RC2a-triplet** | -3001.542590 | -3000.887835 | -3000.999173 |
| **RC2b-quintet** | -3001.527742 | -3000.876675 | -3000.988051 |
| **RC2b-septet** | -3001.516608 | -3000.865845 | -3000.978407 |
| **RC2b-singlet** | -3001.534163 | -3000.879040 | -3000.986792 |
| **RC2b-triplet** | -3001.545701 | -3000.890697 | -3000.999582 |
| **TS1-quintet** | -3111.058521 | -3110.397652 | -3110.518332 |
| **TS1-septet** | -3111.047634 | -3110.386905 | -3110.508072 |
| **TS1-singlet** | -3111.057956 | -3110.394846 | -3110.509861 |
| **TS1-triplet** | -3111.065968 | -3110.402709 | -3110.517719 |
| **TS2a-quintet** | -3001.511935 | -3000.866788 | -3000.975892 |
| **TS2a-septet** | -3001.498886 | -3000.854379 | -3000.965513 |
| **TS2a-singlet** | -3001.507251 | -3000.859428 | -3000.962146 |
| **TS2a-triplet** | -3001.519847 | -3000.872377 | -3000.976291 |
| **TS2b-quintet** | -3001.503886 | -3000.859109 | -3000.970649 |
| **TS2b-septet** | -3001.493961 | -3000.849419 | -3000.960991 |
| **TS2b-singlet** | -3001.503454 | -3000.855394 | -3000.958461 |
| **TS2b-triplet** | -3001.510783 | -3000.863689 | -3000.972014 |
| **TS3a-triplet** | -3001.532820 | -3000.881057 | -3000.986225 |
| **TS3a-quintet** | -3001.520082 | -3000.871102 | -3000.982085 |
| **TS3a-septet** | -3001.515090 | -3000.866185 | -3000.973514 |
| **TS3b-triplet** | -3001.520445 | -3000.868722 | -3000.976704 |
| **TS3b-quintet** | -3001.495653 | -3000.847229 | -3000.960153 |
| **TS3b-septet** | -3001.496822 | -3000.848232 | -3000.960370 |
| **CP1′-quintet** | -3362.837249 | -3362.165557 | -3362.285389 |
| **CP1′-septet** | -3362.807778 | -3362.137789 | -3362.258465 |
| **CP1′-singlet** | -3362.852395 | -3362.178136 | -3362.295865 |
| **CP1′-triplet** | -3362.854157 | -3362.181194 | -3362.304147 |
| **TS1′-quintet** | -3362.797100 | -3362.129884 | -3362.250020 |
| **TS1′-septet** | -3362.802062 | -3362.134304 | -3362.256079 |
| **TS1′-singlet** | -3362.804743 | -3362.134438 | -3362.247792 |
| **TS1′-triplet** | -3362.818904 | -3362.148438 | -3362.265385 |
| **INT1′-quintet** | -3253.267817 | -3252.610907 | -3252.726582 |
| **INT1′-septet** | -3253.265969 | -3252.608691 | -3252.728605 |
| **INT1′-singlet** | -3253.282891 | -3252.621204 | -3252.731586 |
| **INT1′-triplet** | -3253.293474 | -3252.632873 | -3252.747467 |

| | | | |
|---|---|---|---|
| **RC2b′-quintet** | -3253.260217 | -3252.603921 | -3252.714158 |
| **RC2b′-septet** | -3253.273134 | -3252.615236 | -3252.727794 |
| **RC2b′-singlet** | -3253.286650 | -3252.624986 | -3252.731748 |
| **RC2b′-triplet** | -3253.298888 | -3252.637812 | -3252.745600 |
| **RC2a′-quintet** | -3253.269783 | -3252.612587 | -3252.722691 |
| **RC2a′-septet** | -3253.255721 | -3252.598110 | -3252.712488 |
| **RC2a′-singlet** | -3253.284012 | -3252.623196 | -3252.732068 |
| **RC2a′-triplet** | -3253.295532 | -3252.634635 | -3252.744115 |
| **TS2b′-quintet** | -3253.262031 | -3252.610014 | -3252.721254 |
| **TS2b′-septet** | -3253.250508 | -3252.598777 | -3252.710375 |
| **TS2b′-singlet** | -3253.252537 | -3252.597577 | -3252.700178 |
| **TS2b′-triplet** | -3253.260502 | -3252.606693 | -3252.711749 |
| **TS2a′-quintet** | -3253.269074 | -3252.616755 | -3252.726220 |
| **TS2a′-septet** | -3253.256325 | -3252.604354 | -3252.714181 |
| **TS2a′-singlet** | -3253.252781 | -3252.597702 | -3252.699501 |
| **TS2a′-triplet** | -3253.274323 | -3252.619358 | -3252.724310 |
| **INT2b′-quintet** | -3253.270632 | -3252.613172 | -3252.725799 |
| **INT2b′-septet** | -3253.275537 | -3252.619071 | -3252.733311 |
| **INT2b′-triplet** | -3253.284049 | -3252.624504 | -3252.733405 |
| **INT2a′-quintet** | -3253.286577 | -3252.628577 | -3252.740790 |
| **INT2a′-septet** | -3253.291645 | -3252.634557 | -3252.747836 |
| **INT2a′-triplet** | -3253.300711 | -3252.640581 | -3252.747936 |
| **P2b′-quintet** | -3253.365346 | -3252.703186 | -3252.814124 |
| **P2b′-septet** | -3253.326245 | -3252.665082 | -3252.774151 |
| **P2b′-singlet** | -3253.367921 | -3252.701681 | -3252.806188 |
| **P2b′-triplet** | -3253.366838 | -3252.703355 | -3252.814184 |
| **P2a′-quintet** | -3253.361646 | -3252.699228 | -3252.808108 |
| **P2a′-septet** | -3253.327806 | -3252.666376 | -3252.773689 |
| **P2a′-singlet** | -3253.370231 | -3252.703652 | -3252.806852 |
| **P2a′-triplet** | -3253.370197 | -3252.706291 | -3252.815040 |
| **TS3a′-septet** | -3253.262661 | -3252.606245 | -3252.715700 |
| **TS3a′-triplet** | -3253.278375 | -3252.619374 | -3252.726820 |
| **TS3a′-quintet** | -3253.265614 | -3252.608514 | -3252.718009 |
| **TS3b′-septet** | -3253.252530 | -3252.596867 | -3252.708882 |
| **TS3b′-triplet** | -3253.274806 | -3252.616748 | -3252.727832 |
| **TS3b′-quintet** | -3253.250854 | -3252.595489 | -3252.705629 |

**Representative Analytical Data of Starting Materials**

All product standards were purchased from commercial sources and used without further purification. Most alkyl azides are not stable under FD/FI/TOF conditions; therefore, we were not able to collect the HRMS data for the following compounds.
For review only: Most analytical data of starting materials are reported in our pyrrolidine synthase paper (*ref. 67 in manuscript*). At the discretion of the reviewers, we can provide a copy for the review process.

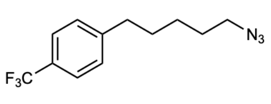

**1-(5-azidopentyl)-4-(trifluoromethyl)benzene (1g):** The reaction was carried out following General Procedure A. **$^{1}$H NMR** (400 MHz, $CDCl_3$) δ 7.81–7.52 (m, 2H), 7.34–7.27 (m, 2H), 3.27 (t, $J$ = 6.8 Hz, 2H), 2.68 (t, $J$ = 7.7 Hz, 2H), 1.94–1.60 (m, 4H), 1.53–1.35 (m, 2H). **$^{13}$C NMR** (101 MHz, $CDCl_3$) $^{13}$C NMR (101 MHz, $CDCl_3$) δ 146.49, 128.80, 128.66–128.44 (m), 125.41 (q, $J_{C–F}$ = 3.7 Hz), 123.15, 51.47, 35.75, 30.84, 28.85, 26.45. **$^{19}$F NMR** (376 MHz, $CDCl_3$) δ –62.30.

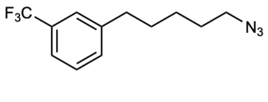

**1-(5-azidopentyl)-3-(trifluoromethyl)benzene (1k):** The reaction was carried out following General Procedure A. $^{1}$H NMR (400 MHz, $CDCl_3$) δ 7.72–7.29 (m, 4H), 3.27 (td, $J$ = 6.9, 4.0 Hz, 2H), 2.80–2.56 (m, 2H), 1.84–1.59 (m, 4H), 1.46–1.36 (m, 2H). **$^{13}$C NMR** (101 MHz, $CDCl_3$) δ 143.26, 131.91 (d, $J_{C–F}$ = 1.6 Hz), 130.78 (d, $J_{C–F}$ = 31.7 Hz), 128.99, 128.87, 125.16 (q, $J_{C–F}$ = 3.8 Hz), 122.85 (q, $J_{C–F}$ = 3.9 Hz), 51.46, 35.73, 30.94, 28.84, 26.46. **$^{19}$F NMR** (376 MHz, $CDCl_3$) δ –62.57.

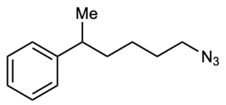

**(6-azidohexan-2-yl)benzene (1m):** The reaction was carried out following General Procedure B. $^{1}$H NMR (400 MHz, $CDCl_3$) δ 7.46–7.26 (m, 2H), 7.23–7.11 (m, 3H), 3.21 (td, $J$ = 7.0, 1.7 Hz, 2H), 2.68 (q, $J$ = 7.1 Hz, 1H), 1.72–1.53 (m, 4H), 1.46–1.14 (m, 5H). **$^{13}$C NMR** (101 MHz, $CDCl_3$) δ 147.46, 128.51, 127.08, 126.10, 51.51, 40.01, 38.03, 29.05, 25.01, 22.45.

### Representative NMR spectrum

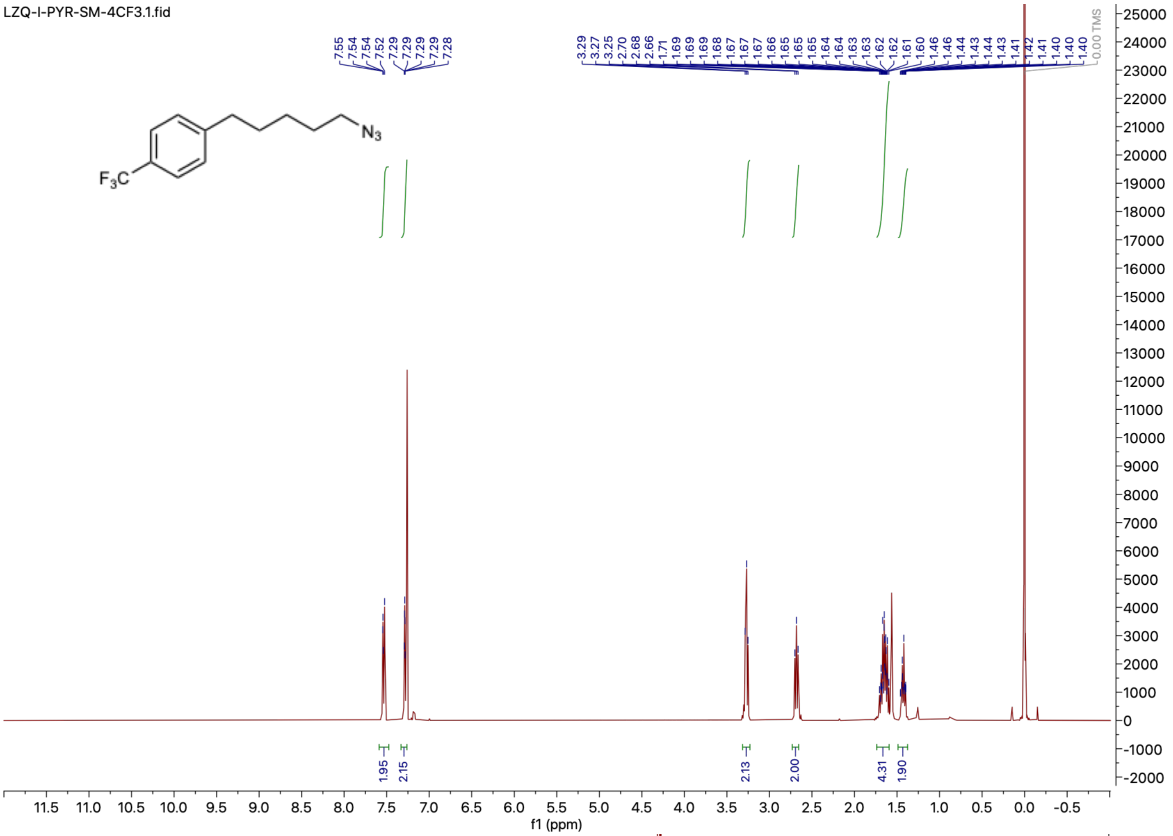

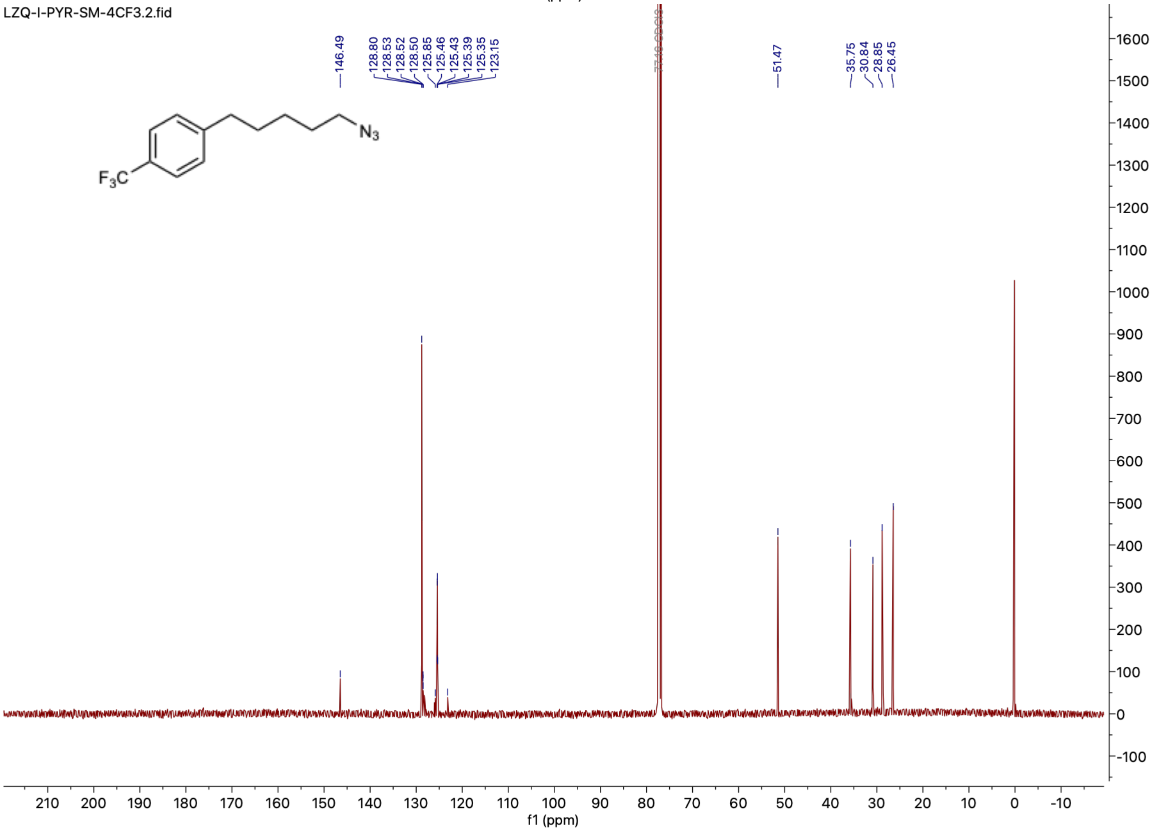

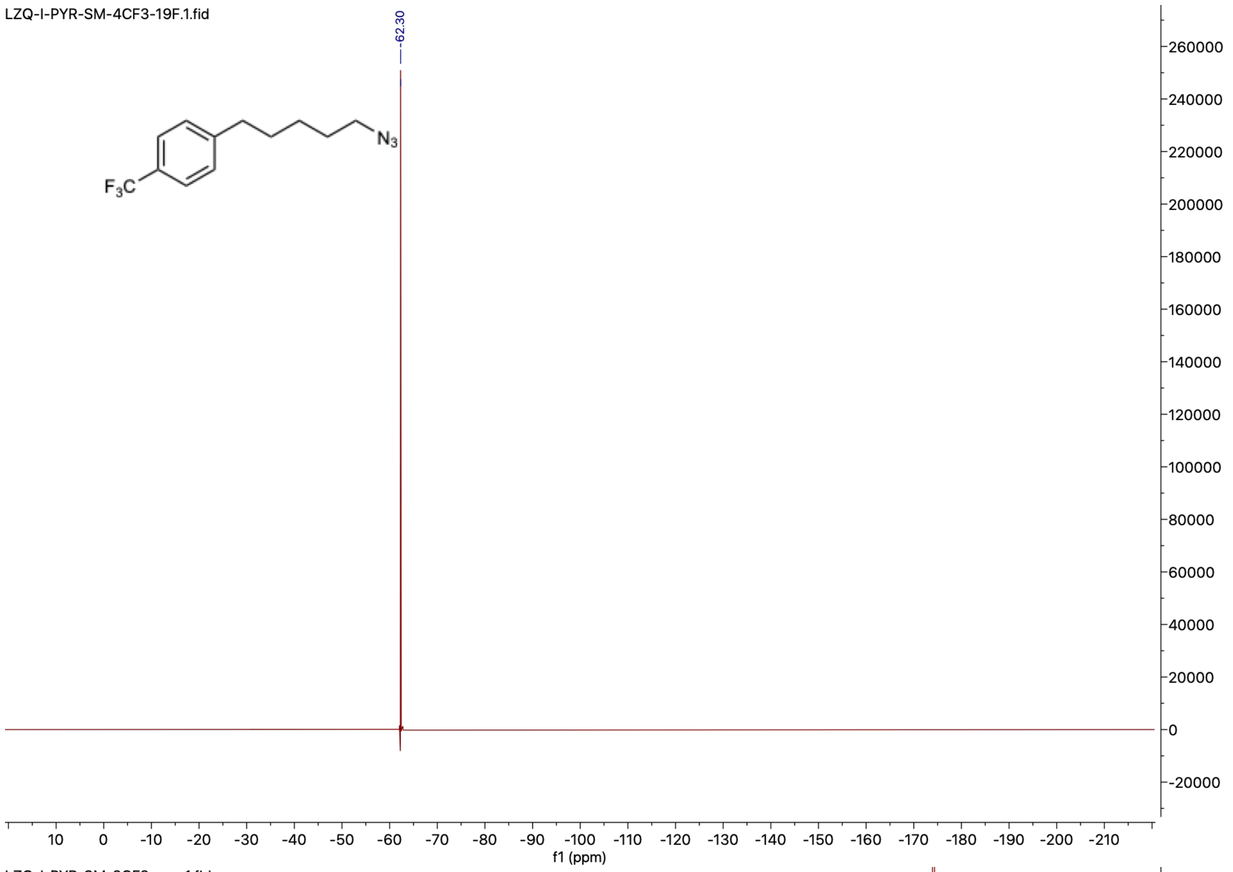
LZQ-I-PYR-SM-4CF3-19F.1.fid
-62.30
N3
F3C
f1 (ppm)

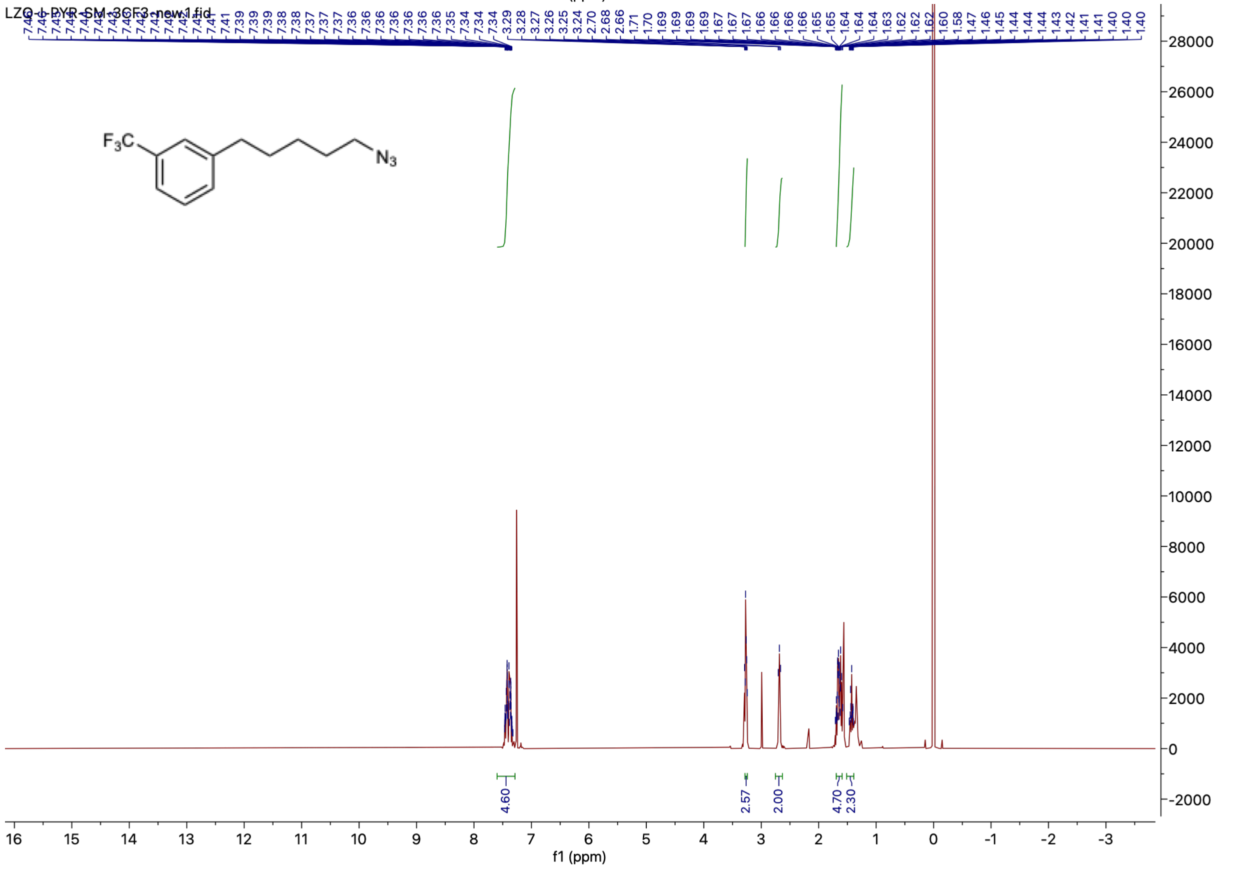
LZQ-I-PYR-SM-3CF3-new.1.fid
F3C
N3
4.60
2.57
2.00
4.70
2.30
f1 (ppm)

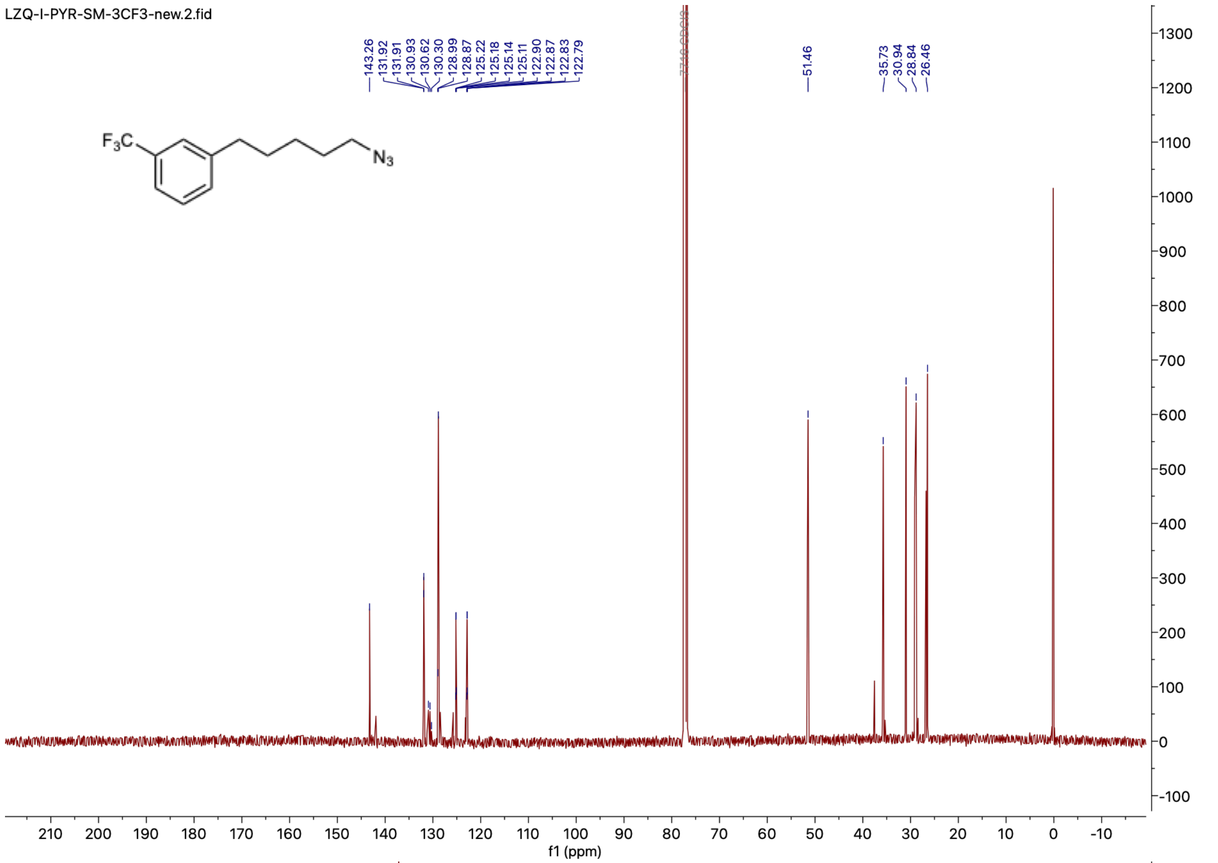
LZQ-I-PYR-SM-3CF3-new.2.fid
143.26
131.92
131.91
130.93
130.62
130.30
128.99
128.87
125.22
125.18
125.14
125.11
122.90
122.87
122.83
122.79
51.46
35.73
30.94
28.84
26.46
$F_3C$
$N_3$
f1 (ppm)

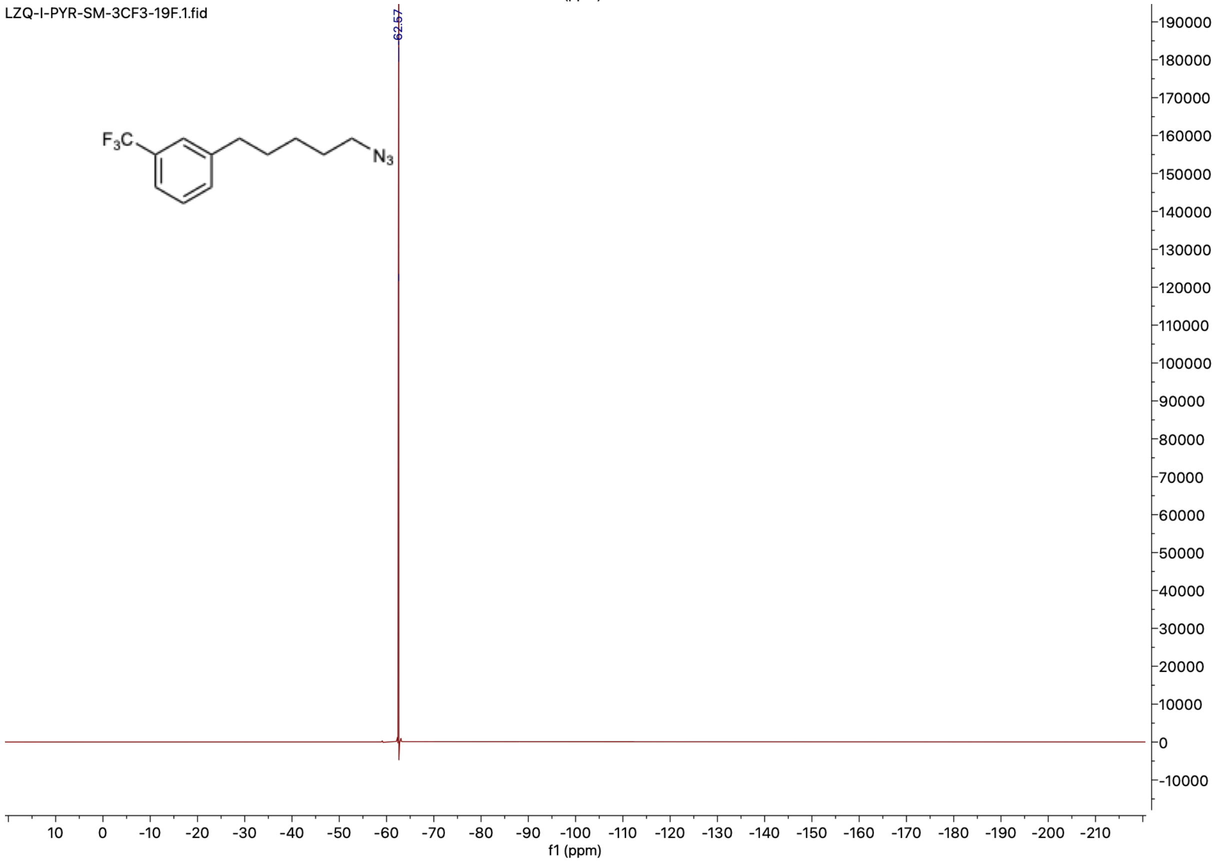
LZQ-I-PYR-SM-3CF3-19F.1.fid
-62.57
$F_3C$
$N_3$
f1 (ppm)

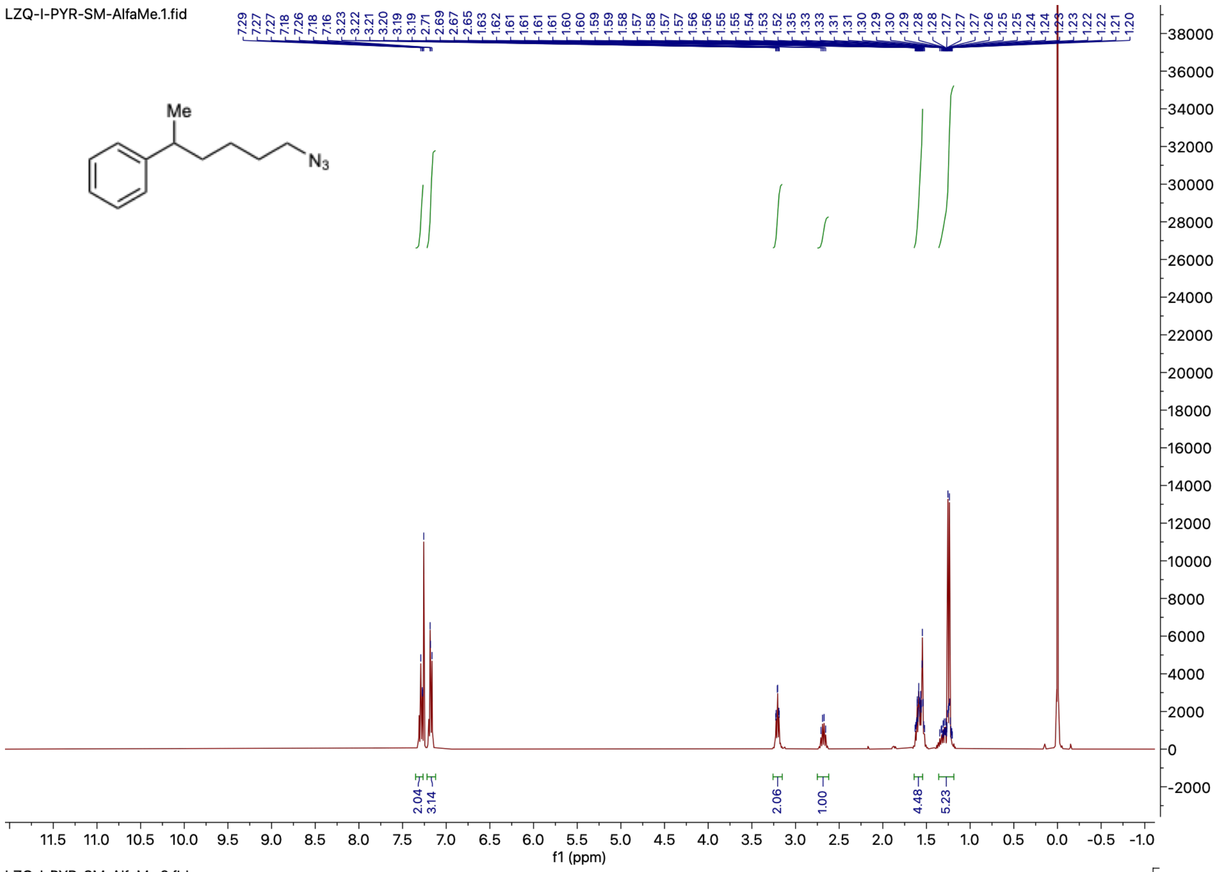
LZQ-I-PYR-SM-AlfaMe.1.fid
Me
N3
2.04
3.14
2.06
1.00
4.48
5.23
f1 (ppm)

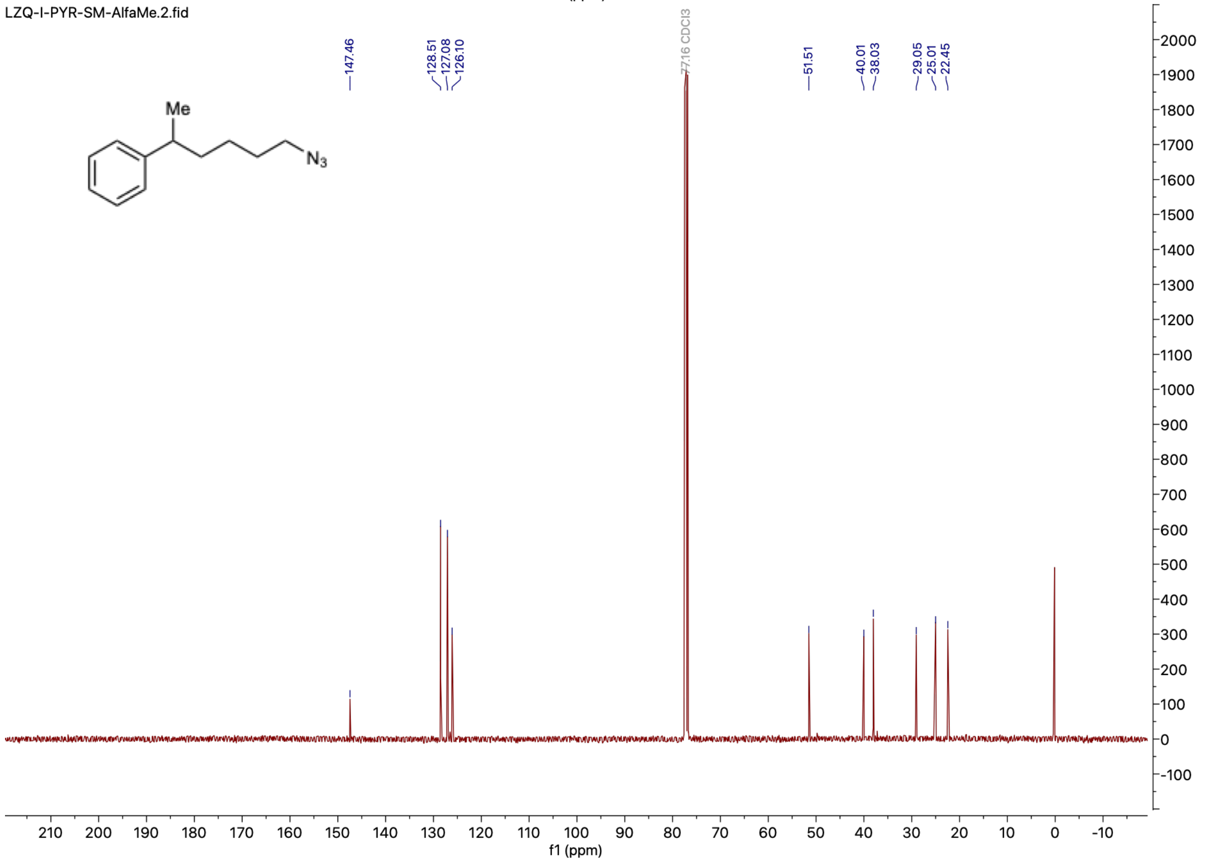
LZQ-I-PYR-SM-AlfaMe.2.fid
147.46
128.51
127.08
126.10
77.16 CDCl3
51.51
40.01
38.03
29.05
25.01
22.45
Me
N3
f1 (ppm)

#### Representative Chiral HPLC-MS data

**Part 1:** Regio- and enantioselectivity analysis of **2a** and **3a** on reactions performed with ***Tm*Pgb-PIP** lineage.
Results analyzed on polar ionic chiral HPLC-MS.

Chiral separation method: polar ionic HPLC-MS using *single* Agilent InfinityLab Poroshell 120 Chiral-V, 2.7 µm, 2.1 × 150 mm; eluent: 40% MeOH (0.2 w% $NH_4CO_2H$), 60% $H_2O$ (20 mM $NH_4CO_2H$, pH 4.0 adjusted with $HCO_2H$); flow rate: 0.2 mL/min.
$t_{R\text{-}2a}$ = 15.439 min, $t_{S\text{-}2a}$ = 16.270 min. $t_{R\text{-}3a}$ = 14.466 min, $t_{S\text{-}3a}$ = 17.430 min.

(*rac*)-**2a** (commercially available standard)

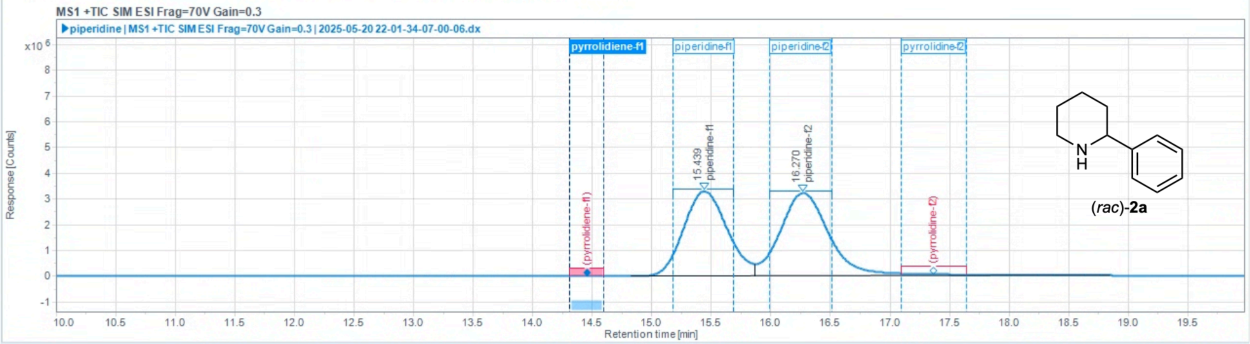

Injection Results

| # | Name | Signal description | RT (min) | Area (Counts·s) | Area% | Height (Counts) | Height% | Amount | Concentration | Start time (min) | End time (min) |
|---|---|---|---|---|---|---|---|---|---|---|---|
| 1 | piperidine-f1 | MS1 +TIC SIM ESI Frag=70V Gain=0.3 | 15.439 | 83873295.146 | 47.132 | 3287426.993 | 50.42 | | | 14.825 | 15.868 |
| 2 | piperidine-f2 | MS1 +TIC SIM ESI Frag=70V Gain=0.3 | 16.270 | 94081538.107 | 52.868 | 3232983.971 | 49.58 | | | 15.868 | 18.851 |

(*rac*)-**3a** (commercially available standard)

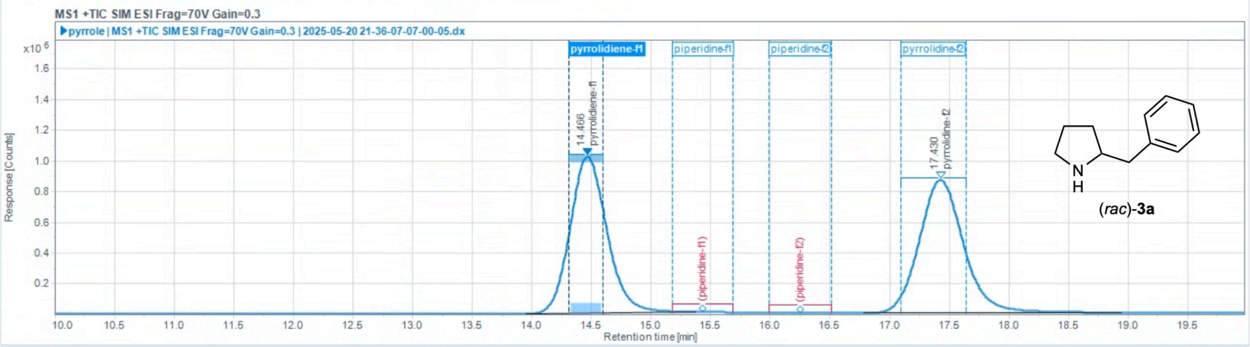

Injection Results

| # | Name | Signal description | RT (min) | Area (Counts·s) | Area% | Height (Counts) | Height% | Amount | Concentration | Start time (min) | End time (min) |
|---|---|---|---|---|---|---|---|---|---|---|---|
| 1 | pyrrolidiene-f1 | MS1 +TIC SIM ESI Frag=70V Gain=0.3 | 14.466 | 21786543.915 | 49.217 | 1020171.145 | 54.01 | | | 13.947 | 15.371 |
| 2 | pyrrolidine-f2 | MS1 +TIC SIM ESI Frag=70V Gain=0.3 | 17.430 | 22480066.557 | 50.783 | 868519.200 | 45.99 | | | 16.780 | 18.942 |

**Table S27**. Regioselectivity and enantioselectivity of TmPgb-PIP lineage.

| | rr (**2a**:**3a**) | **2a** ee (%) |
|---|---|---|
| ***Tm*Pgb-wt** | 3:97 | –96 |
| ***Tm*Pgb-PIP-G1** | 45:55 | >99 |
| ***Tm*Pgb-PIP-G2** | 56:44 | >99 |
| ***Tm*Pgb-PIP-G3** | 71:29 | >99 |
| ***Tm*Pgb-PIP-G4** | 89:11 | >99 |
| ***Tm*Pgb-PIP-G5** | 86:14 | >99 |
| ***Tm*Pgb-PIP-G6** | 85:15 | >99 |
| ***Tm*Pgb-PIP-G7** | 84:16 | >99 |

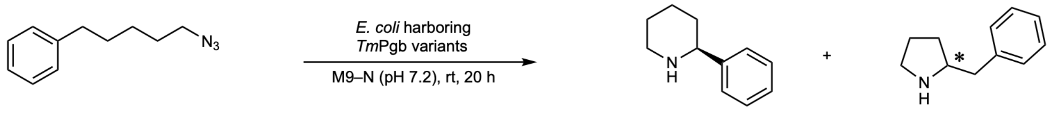

Reaction performed on analytical scale using ***Tm*Pgb-wt**

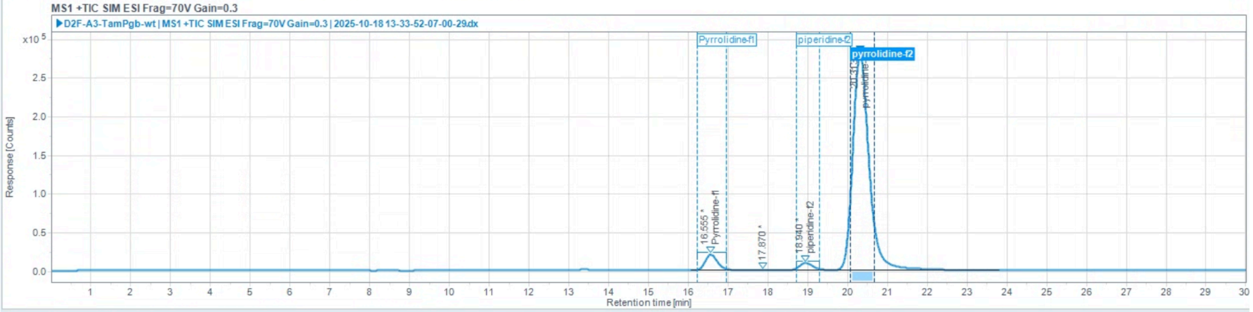

Injection Results

Peaks Summary

| # | Name | Signal description | RT (min) | Area (Counts·s) | Area% | Height (Counts) | Height% | Amount | Concentration | Start time (min) | End time (min) |
|---|---|---|---|---|---|---|---|---|---|---|---|
| 1 | Pyrrolidine-f1 | MS1 +TIC SIM ESI Frag=70V Gain=0.3 | 16.555 | 455684.495 | 5.242 | 20462.948 | 6.59 | | | 16.058 | 17.526 |
| 2 | | MS1 +TIC SIM ESI Frag=70V Gain=0.3 | 17.870 | 4577.763 | 0.053 | 199.969 | 0.06 | | | 17.526 | 18.371 |
| 3 | piperidine-f2 | MS1 +TIC SIM ESI Frag=70V Gain=0.3 | 18.940 | 230689.720 | 2.654 | 9198.722 | 2.96 | | | 18.371 | 19.638 |
| 4 | pyrrolidine-f2 | MS1 +TIC SIM ESI Frag=70V Gain=0.3 | 20.313 | 8001452.668 | 92.051 | 280436.847 | 90.38 | | | 19.638 | 23.819 |

Reaction performed on analytical scale using ***Tm*Pgb-PIP-G1-5442**

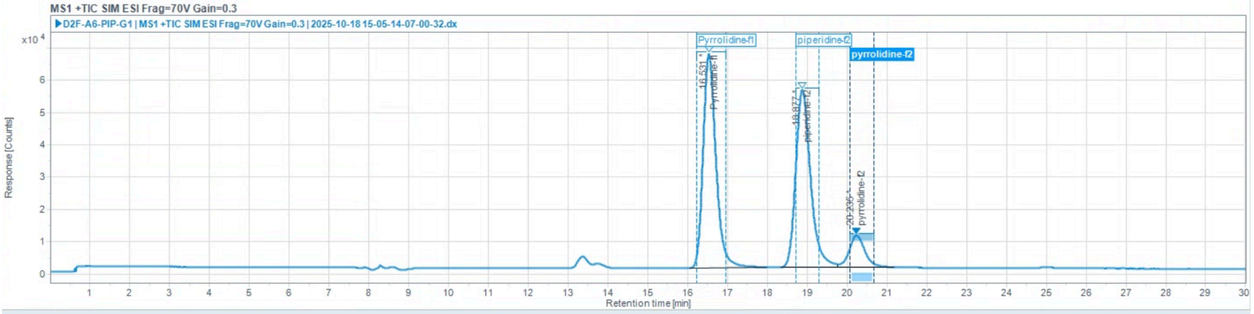

Injection Results

Peaks Summary

| # | Name | Signal description | RT (min) | Area (Counts·s) | Area% | Height (Counts) | Height% | Amount | Concentration | Start time (min) | End time (min) |
|---|---|---|---|---|---|---|---|---|---|---|---|
| 1 | Pyrrolidine-f1 | MS1 +TIC SIM ESI Frag=70V Gain=0.3 | 16.531 | 1486396.975 | 46.741 | 66325.262 | 50.59 | | | 16.038 | 17.970 |
| 2 | piperidine-f2 | MS1 +TIC SIM ESI Frag=70V Gain=0.3 | 18.877 | 1423458.054 | 44.762 | 54927.665 | 41.90 | | | 18.330 | 19.749 |
| 3 | pyrrolidine-f2 | MS1 +TIC SIM ESI Frag=70V Gain=0.3 | 20.235 | 270189.383 | 8.496 | 9839.649 | 7.51 | | | 19.749 | 21.173 |

Reaction performed on analytical scale using ***Tm*****Pgb-PIP-G2-5443**

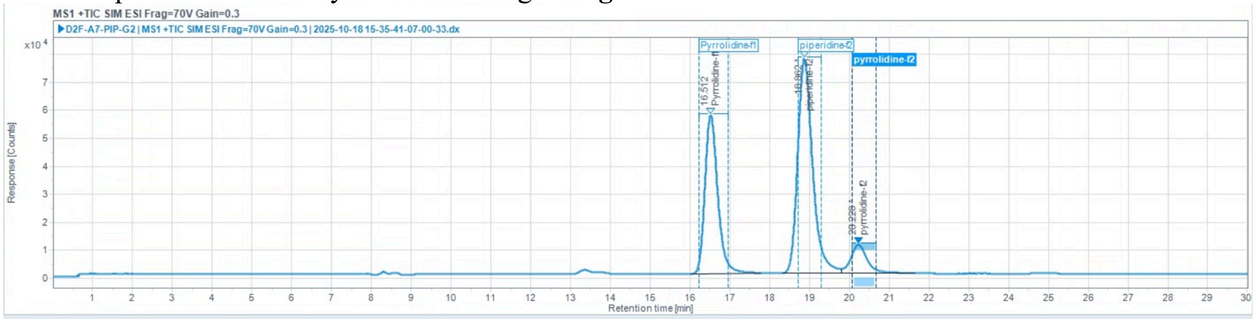

Injection Results

Peaks Summary

| # | Name | Signal description | RT (min) | Area (Counts·s) | Area% | Height (Counts) | Height% | Amount | Concentration | Start time (min) | End time (min) |
|---|---|---|---|---|---|---|---|---|---|---|---|
| 1 | Pyrrolidine-f1 | MS1 +TIC SIM ESI Frag=70V Gain=0.3 | 16.512 | 1270715.320 | 35.720 | 56647.764 | 39.48 | | | 16.014 | 17.782 |
| 2 | piperidine-f2 | MS1 +TIC SIM ESI Frag=70V Gain=0.3 | 18.862 | 1996270.480 | 56.116 | 76677.708 | 53.44 | | | 18.312 | 19.794 |
| 3 | pyrrolidine-f2 | MS1 +TIC SIM ESI Frag=70V Gain=0.3 | 20.228 | 290421.869 | 8.164 | 10165.492 | 7.08 | | | 19.794 | 21.640 |

Reaction performed on analytical scale using ***Tm*****Pgb-PIP-G3-5444**

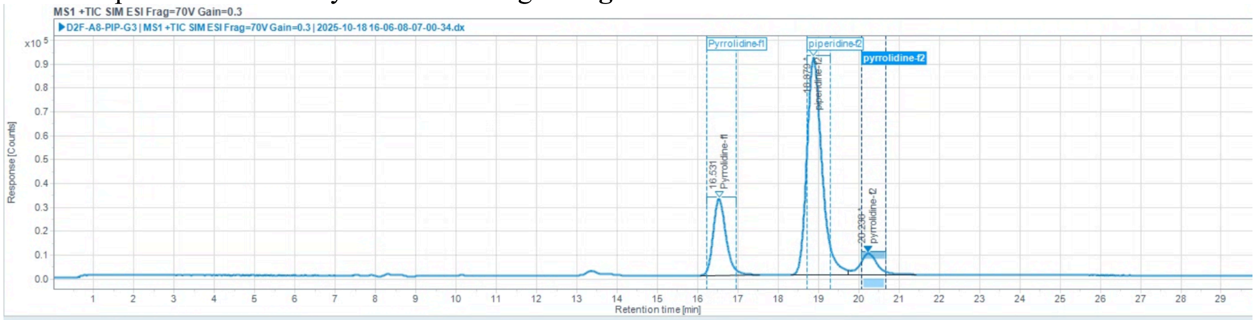

Injection Results

Peaks Summary

| # | Name | Signal description | RT (min) | Area (Counts·s) | Area% | Height (Counts) | Height% | Amount | Concentration | Start time (min) | End time (min) |
|---|---|---|---|---|---|---|---|---|---|---|---|
| 1 | Pyrrolidine-f1 | MS1 +TIC SIM ESI Frag=70V Gain=0.3 | 16.531 | 708546.705 | 21.126 | 31871.509 | 24.13 | | | 16.076 | 17.546 |
| 2 | piperidine-f2 | MS1 +TIC SIM ESI Frag=70V Gain=0.3 | 18.879 | 2377421.122 | 70.884 | 91218.578 | 69.05 | | | 18.312 | 19.749 |
| 3 | pyrrolidine-f2 | MS1 +TIC SIM ESI Frag=70V Gain=0.3 | 20.238 | 267999.182 | 7.991 | 9017.526 | 6.83 | | | 19.749 | 21.417 |

Reaction performed on analytical scale using ***Tm*****Pgb-PIP-G4-5445**

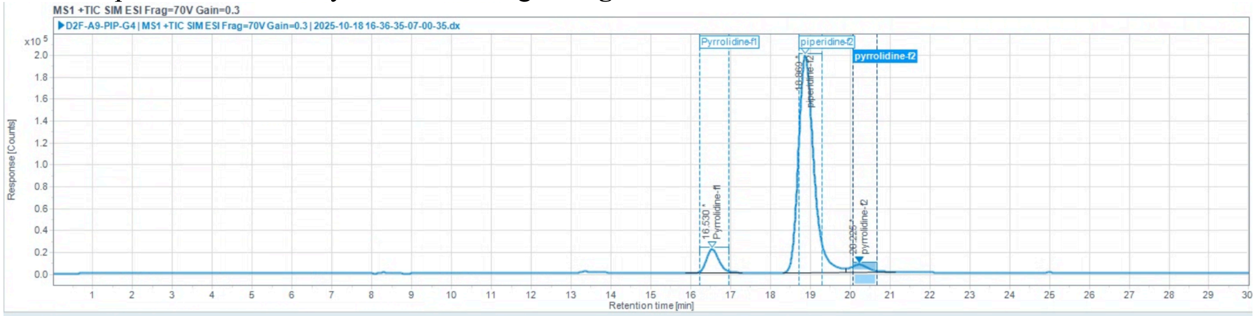

Injection Results

Peaks Summary

| # | Name | Signal description | RT (min) | Area (Counts·s) | Area% | Height (Counts) | Height% | Amount | Concentration | Start time (min) | End time (min) |
|---|---|---|---|---|---|---|---|---|---|---|---|
| 1 | Pyrrolidine-f1 | MS1 +TIC SIM ESI Frag=70V Gain=0.3 | 16.530 | 471100.639 | 7.840 | 21447.295 | 9.45 | | | 15.858 | 17.281 |
| 2 | piperidine-f2 | MS1 +TIC SIM ESI Frag=70V Gain=0.3 | 18.869 | 5318512.123 | 88.514 | 198256.366 | 87.37 | | | 18.305 | 19.883 |
| 3 | pyrrolidine-f2 | MS1 +TIC SIM ESI Frag=70V Gain=0.3 | 20.225 | 219056.070 | 3.646 | 7216.273 | 3.18 | | | 19.883 | 21.128 |

Reaction performed on analytical scale using ***Tm*****Pgb-PIP-G5-5446**

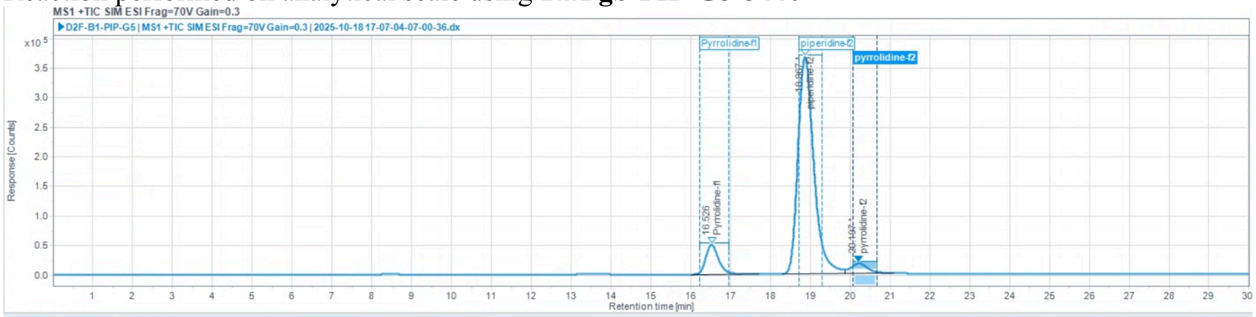

Injection Results

Peaks Summary

| # | Name | Signal description | RT (min) | Area (Counts·s) | Area% | Height (Counts) | Height% | Amount | Concentration | Start time (min) | End time (min) |
|---|---|---|---|---|---|---|---|---|---|---|---|
| 1 | Pyrrolidine-f1 | MS1 +TIC SIM ESI Frag=70V Gain=0.3 | 16.526 | 1133631.338 | 9.581 | 50267.241 | 11.57 | | | 16.032 | 17.708 |
| 2 | piperidine-f2 | MS1 +TIC SIM ESI Frag=70V Gain=0.3 | 18.867 | 10183342.844 | 86.069 | 367395.020 | 84.54 | | | 18.293 | 19.861 |
| 3 | pyrrolidine-f2 | MS1 +TIC SIM ESI Frag=70V Gain=0.3 | 20.197 | 514638.304 | 4.350 | 16941.926 | 3.90 | | | 19.861 | 21.084 |

Reaction performed on analytical scale using ***Tm*****Pgb-PIP-G6-5447**

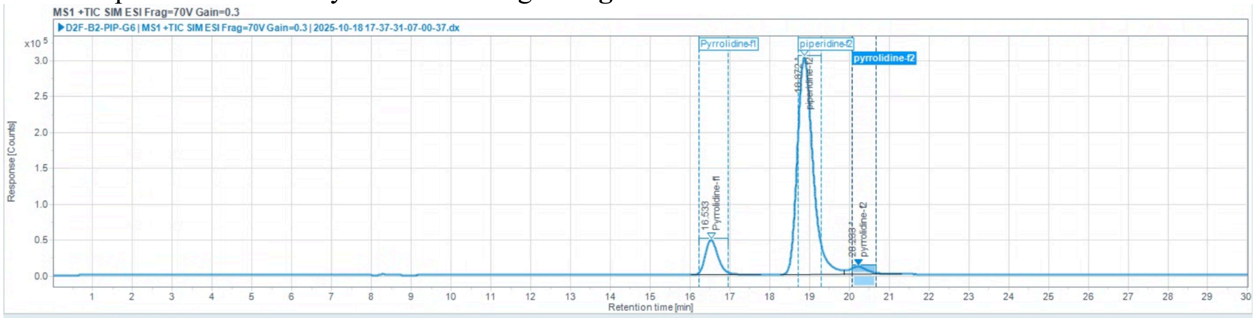

Injection Results

Peaks Summary

| # | Name | Signal description | RT (min) | Area (Counts·s) | Area% | Height (Counts) | Height% | Amount | Concentration | Start time (min) | End time (min) |
|---|---|---|---|---|---|---|---|---|---|---|---|
| 1 | Pyrrolidine-f1 | MS1 +TIC SIM ESI Frag=70V Gain=0.3 | 16.533 | 1115723.894 | 11.387 | 49101.793 | 13.54 | | | 16.026 | 17.782 |
| 2 | piperidine-f2 | MS1 +TIC SIM ESI Frag=70V Gain=0.3 | 18.872 | 8326292.544 | 84.976 | 302740.448 | 83.47 | | | 18.281 | 19.861 |
| 3 | pyrrolidine-f2 | MS1 +TIC SIM ESI Frag=70V Gain=0.3 | 20.233 | 356352.585 | 3.637 | 10841.920 | 2.99 | | | 19.861 | 21.328 |

Reaction performed on analytical scale using ***Tm*****Pgb-PIP-G7-5448**

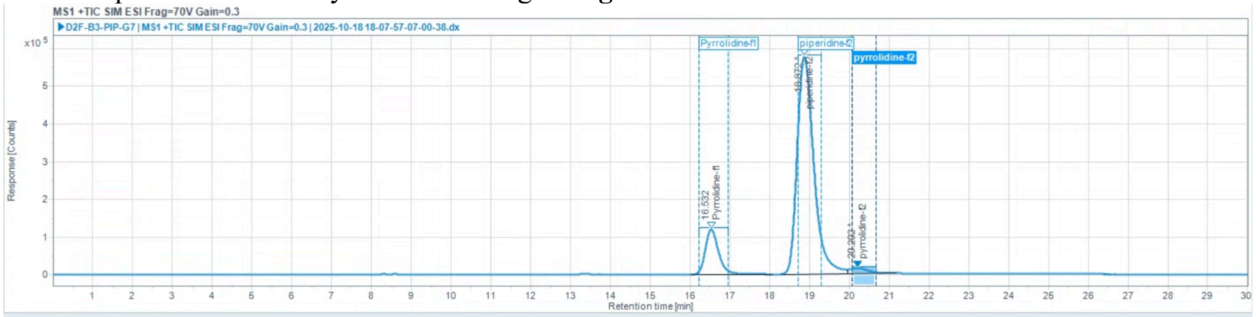

Injection Results

Peaks Summary

| # | Name | Signal description | RT (min) | Area (Counts·s) | Area% | Height (Counts) | Height% | Amount | Concentration | Start time (min) | End time (min) |
|---|---|---|---|---|---|---|---|---|---|---|---|
| 1 | Pyrrolidine-f1 | MS1 +TIC SIM ESI Frag=70V Gain=0.3 | 16.532 | 2771102.938 | 13.868 | 119696.705 | 16.86 | | | 16.020 | 18.075 |
| 2 | piperidine-f2 | MS1 +TIC SIM ESI Frag=70V Gain=0.3 | 18.872 | 16744088.324 | 83.795 | 575424.514 | 81.07 | | | 18.287 | 19.950 |
| 3 | pyrrolidine-f2 | MS1 +TIC SIM ESI Frag=70V Gain=0.3 | 20.202 | 467014.020 | 2.337 | 14659.697 | 2.07 | | | 19.950 | 21.173 |

**Part 2: Regio- and enantioselectivity analysis of 2a and 3a on reactions performed with GDM-NT variants.**

Results analyzed on polar ionic chiral HPLC-MS.

(*rac*)-**2a** (commercially available standard)

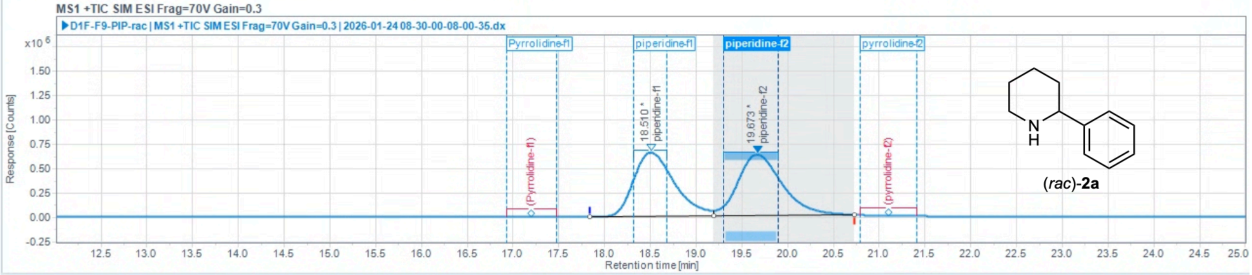

Injection Results

Peaks | Summary

| # | Name | Signal description | RT (min) | Area (Counts·s) | Area% | Height (Counts) | Height% | Amount | Concentration | Start time (min) | End time (min) |
|---|---|---|---|---|---|---|---|---|---|---|---|
| 1 | piperidine-f1 | MS1 +TIC SIM ESI Frag=70V Gain=0.3 | 18.510 | 20366752.140 | 49.171 | 656971.736 | 51.21 | | | 17.838 | 19.188 |
| 2 | piperidine-f2 | MS1 +TIC SIM ESI Frag=70V Gain=0.3 | 19.673 | 21053749.561 | 50.829 | 625903.993 | 48.79 | | | 19.188 | 20.723 |

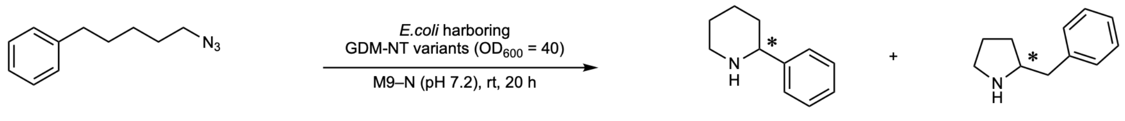

**1a** **2a** **3a**

Reaction performed on analytical scale using ***Tm*Pgb-wt**: 3:97 rr (**2a**:**3a**), –97% ee favoring (*S*)-**2a**.

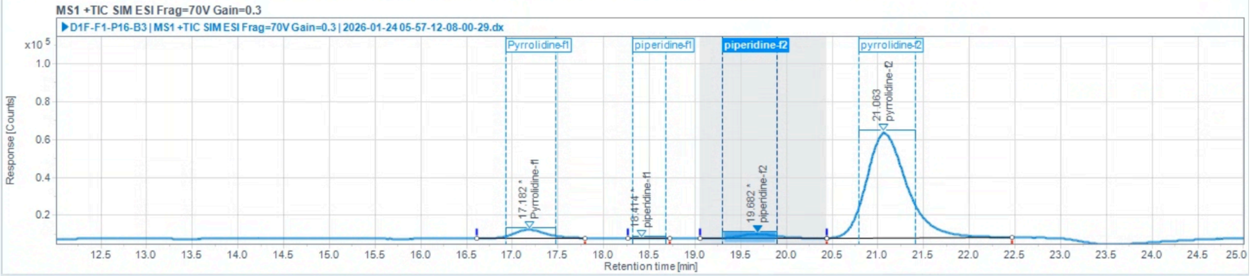

Injection Results

Peaks | Summary

| # | Name | Signal description | RT (min) | Area (Counts·s) | Area% | Height (Counts) | Height% | Amount | Concentration | Start time (min) | End time (min) |
|---|---|---|---|---|---|---|---|---|---|---|---|
| 1 | Pyrrolidine-f1 | MS1 +TIC SIM ESI Frag=70V Gain=0.3 | 17.182 | 114643.960 | 6.049 | 4585.096 | 7.30 | | | 16.606 | 17.796 |
| 2 | piperidine-f1 | MS1 +TIC SIM ESI Frag=70V Gain=0.3 | 18.414 | 1009.813 | 0.053 | 114.362 | 0.18 | | | 18.259 | 18.728 |
| 3 | piperidine-f2 | MS1 +TIC SIM ESI Frag=70V Gain=0.3 | 19.682 | 64158.247 | 3.385 | 2323.900 | 3.70 | | | 19.058 | 20.442 |
| 4 | pyrrolidine-f2 | MS1 +TIC SIM ESI Frag=70V Gain=0.3 | 21.063 | 1715494.866 | 90.513 | 55758.236 | 88.81 | | | 20.442 | 22.465 |

**Table S28**. Regioselectivity and enantioselectivity of selected GDM-NT variants using recycled scaffold design.

| Variants | rr (**2a**:**3a**) | ee (%) | Variants | rr (**2a**:**3a**) | ee (%) |
|---|---|---|---|---|---|
| **GDM-NT-121** | 96:4 | –40 | **GDM-NT-151** | 95:5 | –69 |
| **GDM-NT-215** | 98:2 | –24 | **GDM-NT-350** | 94:6 | –45 |
| **GDM-NT-217** | 96:4 | 31 | **GDM-NT-356** | 99:1 | –73 |
| **GDM-NT-226** | >99:1 | 30 | **GDM-NT-359** | 98:2 | –64 |
| **GDM-NT-227** | 94:6 | 4 | **GDM-NT-361** | 95:5 | –23 |
| **GDM-NT-228** | 89:11 | 50 | **GDM-NT-362** | >99:1 | 89 |

Reaction performed on analytical scale using **GDM-NT-121**: 96:4 rr (**2a**:**3a**), –40% ee favoring (*S*)-**2a**.

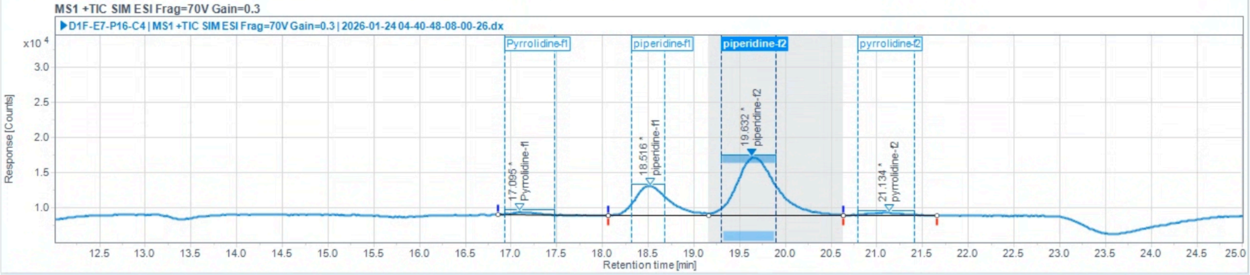

Injection Results

Peaks | Summary

| # | Name | Signal description | RT (min) | Area (Counts·s) | Area% | Height (Counts) | Height% | Amount | Concentration | Start time (min) | End time (min) |
|---|---|---|---|---|---|---|---|---|---|---|---|
| 1 | Pyrrolidine-f1 | MS1 +TIC SIM ESI Frag=70V Gain=0.3 | 17.095 | 5713.548 | 1.572 | 413.341 | 3.16 | | | 16.852 | 18.056 |
| 2 | piperidine-f1 | MS1 +TIC SIM ESI Frag=70V Gain=0.3 | 18.516 | 105591.081 | 29.050 | 4116.973 | 31.46 | | | 18.056 | 19.166 |
| 3 | piperidine-f2 | MS1 +TIC SIM ESI Frag=70V Gain=0.3 | 19.632 | 244460.026 | 67.255 | 8197.223 | 62.64 | | | 19.166 | 20.627 |
| 4 | pyrrolidine-f2 | MS1 +TIC SIM ESI Frag=70V Gain=0.3 | 21.134 | 7718.407 | 2.123 | 359.373 | 2.75 | | | 20.627 | 21.655 |

Reaction performed on analytical scale using **GDM-NT-215**: 98:2 rr (**2a**:**3a**), –24% ee favoring (*S*)-**2a**.

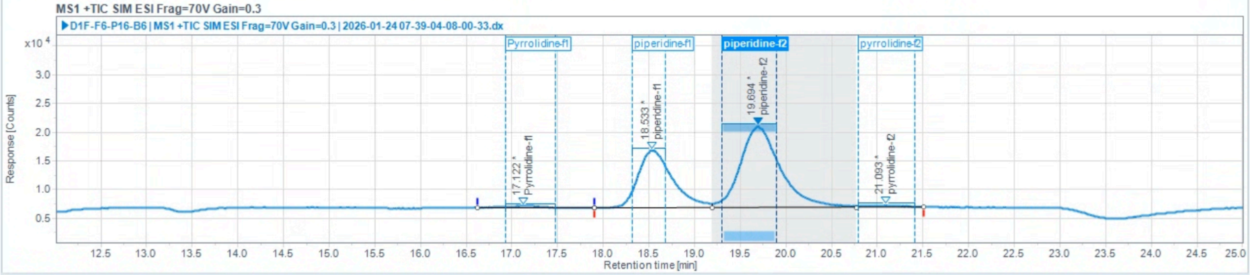

Injection Results

Peaks | Summary

| # | Name | Signal description | RT (min) | Area (Counts·s) | Area% | Height (Counts) | Height% | Amount | Concentration | Start time (min) | End time (min) |
|---|---|---|---|---|---|---|---|---|---|---|---|
| 1 | Pyrrolidine-f1 | MS1 +TIC SIM ESI Frag=70V Gain=0.3 | 17.122 | 8948.585 | 1.252 | 351.331 | 1.41 | | | 16.626 | 17.896 |
| 2 | piperidine-f1 | MS1 +TIC SIM ESI Frag=70V Gain=0.3 | 18.533 | 266138.586 | 37.231 | 10047.514 | 40.27 | | | 17.896 | 19.187 |
| 3 | piperidine-f2 | MS1 +TIC SIM ESI Frag=70V Gain=0.3 | 19.694 | 430942.757 | 60.285 | 14207.662 | 56.95 | | | 19.187 | 20.772 |
| 4 | pyrrolidine-f2 | MS1 +TIC SIM ESI Frag=70V Gain=0.3 | 21.093 | 8809.530 | 1.232 | 341.911 | 1.37 | | | 20.772 | 21.511 |

Reaction performed on analytical scale using **GDM-NT-217**: 96:4 rr (**2a**:**3a**), 31% ee favoring (*R*)-**2a**.

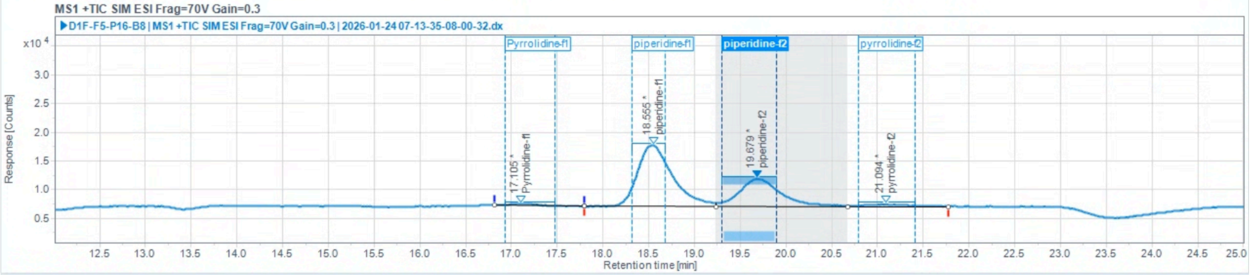

Injection Results

Peaks | Summary

| # | Name | Signal description | RT (min) | Area (Counts·s) | Area% | Height (Counts) | Height% | Amount | Concentration | Start time (min) | End time (min) |
|---|---|---|---|---|---|---|---|---|---|---|---|
| 1 | Pyrrolidine-f1 | MS1 +TIC SIM ESI Frag=70V Gain=0.3 | 17.105 | 4883.638 | 1.075 | 270.723 | 1.68 | | | 16.811 | 17.800 |
| 2 | piperidine-f1 | MS1 +TIC SIM ESI Frag=70V Gain=0.3 | 18.555 | 285303.454 | 62.799 | 10671.048 | 66.10 | | | 17.800 | 19.230 |
| 3 | piperidine-f2 | MS1 +TIC SIM ESI Frag=70V Gain=0.3 | 19.679 | 151843.229 | 33.423 | 4793.964 | 29.70 | | | 19.230 | 20.670 |
| 4 | pyrrolidine-f2 | MS1 +TIC SIM ESI Frag=70V Gain=0.3 | 21.094 | 12283.893 | 2.704 | 406.988 | 2.52 | | | 20.670 | 21.778 |

Reaction performed on analytical scale using **GDM-NT-226**: >99:1 rr (**2a**:**3a**), 30% ee favoring (*R*)-**2a**.

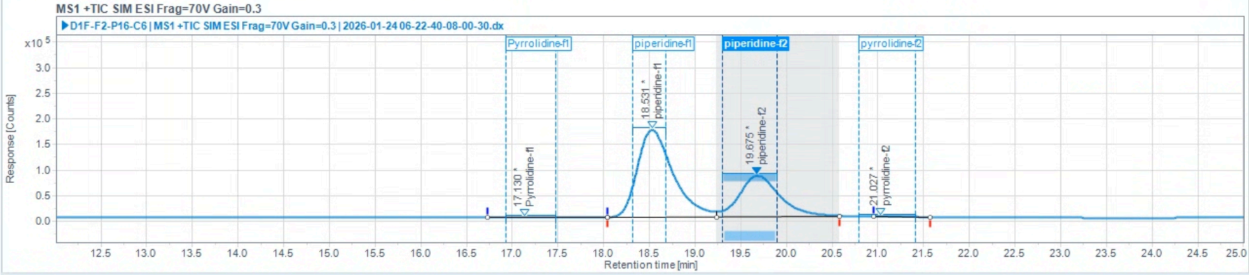

#### Injection Results

Peaks | Summary

| # | Name | Signal description | RT (min) | Area (Counts·s) | Area% | Height (Counts) | Height% | Amount | Concentration | Start time (min) | End time (min) |
|---|---|---|---|---|---|---|---|---|---|---|---|
| 1 | Pyrrolidine-f1 | MS1 +TIC SIM ESI Frag=70V Gain=0.3 | 17.130 | 1871.163 | 0.026 | 565.766 | 0.22 | | | 16.729 | 18.044 |
| 2 | piperidine-f1 | MS1 +TIC SIM ESI Frag=70V Gain=0.3 | 18.531 | 4647606.653 | 64.790 | 170386.701 | 67.68 | | | 18.044 | 19.230 |
| 3 | piperidine-f2 | MS1 +TIC SIM ESI Frag=70V Gain=0.3 | 19.675 | 2520856.918 | 35.142 | 80549.085 | 31.99 | | | 19.230 | 20.579 |
| 4 | pyrrolidine-f2 | MS1 +TIC SIM ESI Frag=70V Gain=0.3 | 21.027 | 3029.448 | 0.042 | 267.841 | 0.11 | | | 20.957 | 21.573 |

Reaction performed on analytical scale using **GDM-NT-227**: 94:6 rr (**2a**:**3a**), 4% ee favoring (*R*)-**2a**.

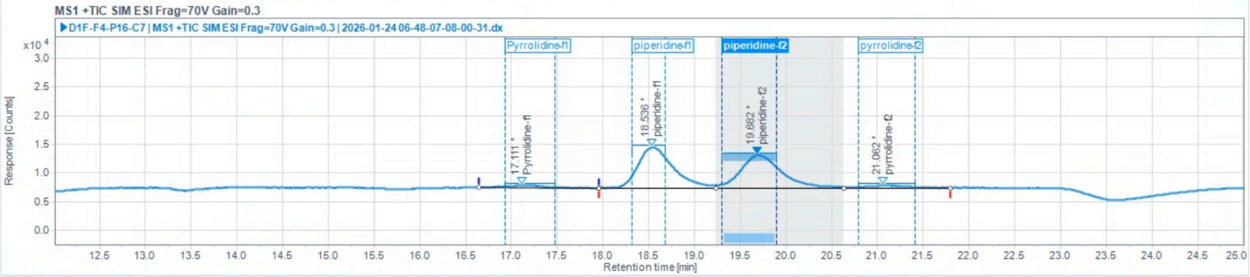

#### Injection Results

Peaks | Summary

| # | Name | Signal description | RT (min) | Area (Counts·s) | Area% | Height (Counts) | Height% | Amount | Concentration | Start time (min) | End time (min) |
|---|---|---|---|---|---|---|---|---|---|---|---|
| 1 | Pyrrolidine-f1 | MS1 +TIC SIM ESI Frag=70V Gain=0.3 | 17.111 | 11245.983 | 2.894 | 464.415 | 3.37 | | | 16.647 | 17.960 |
| 2 | piperidine-f1 | MS1 +TIC SIM ESI Frag=70V Gain=0.3 | 18.536 | 189183.832 | 48.690 | 7150.281 | 51.87 | | | 17.960 | 19.230 |
| 3 | piperidine-f2 | MS1 +TIC SIM ESI Frag=70V Gain=0.3 | 19.682 | 175706.389 | 45.221 | 5766.752 | 41.83 | | | 19.230 | 20.629 |
| 4 | pyrrolidine-f2 | MS1 +TIC SIM ESI Frag=70V Gain=0.3 | 21.062 | 12413.295 | 3.195 | 404.340 | 2.93 | | | 20.629 | 21.799 |

Reaction performed on analytical scale using **GDM-NT-228**: 89:11 rr (**2a**:**3a**), 50% ee favoring (*R*)-**2a**.

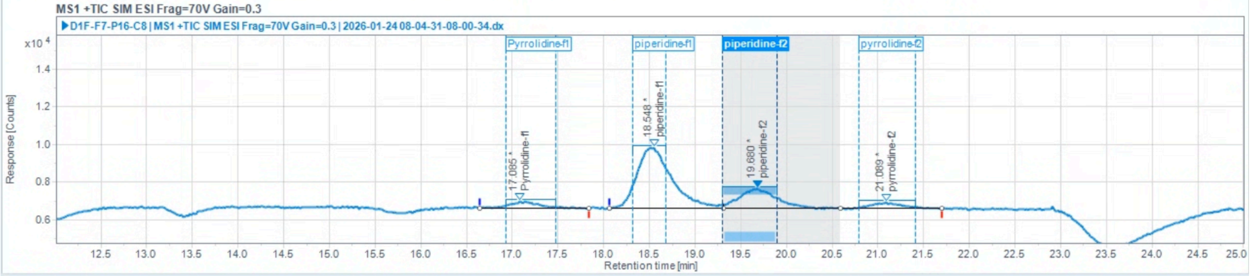

#### Injection Results

Peaks | Summary

| # | Name | Signal description | RT (min) | Area (Counts·s) | Area% | Height (Counts) | Height% | Amount | Concentration | Start time (min) | End time (min) |
|---|---|---|---|---|---|---|---|---|---|---|---|
| 1 | Pyrrolidine-f1 | MS1 +TIC SIM ESI Frag=70V Gain=0.3 | 17.085 | 7995.271 | 6.274 | 362.023 | 7.35 | | | 16.647 | 17.837 |
| 2 | piperidine-f1 | MS1 +TIC SIM ESI Frag=70V Gain=0.3 | 18.548 | 85353.398 | 66.974 | 3217.096 | 65.30 | | | 18.056 | 19.304 |
| 3 | piperidine-f2 | MS1 +TIC SIM ESI Frag=70V Gain=0.3 | 19.680 | 28645.553 | 22.477 | 1042.446 | 21.16 | | | 19.304 | 20.588 |
| 4 | pyrrolidine-f2 | MS1 +TIC SIM ESI Frag=70V Gain=0.3 | 21.089 | 5448.273 | 4.275 | 304.924 | 6.19 | | | 20.588 | 21.696 |

Reaction performed on analytical scale using **GDM-NT-151**: 95:5 rr (**2a**:**3a**), –69% ee favoring (*S*)-**2a**.

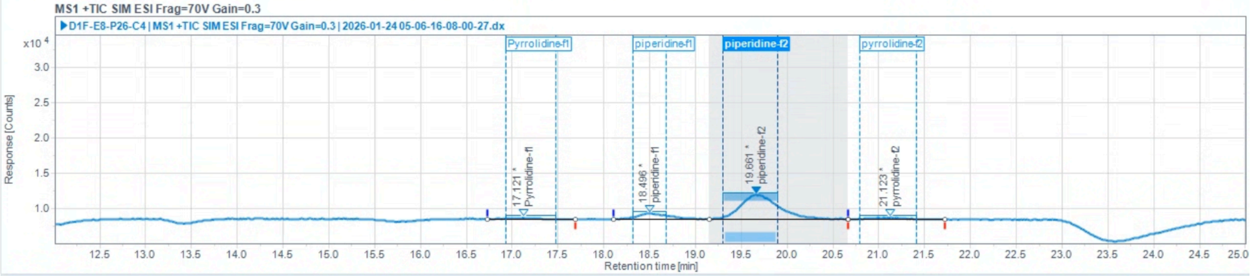

Injection Results

Peaks | Summary

| # | Name | Signal description | RT (min) | Area (Counts·s) | Area% | Height (Counts) | Height% | Amount | Concentration | Start time (min) | End time (min) |
|---|---|---|---|---|---|---|---|---|---|---|---|
| 1 | Pyrrolidine-f1 | MS1 +TIC SIM ESI Frag=70V Gain=0.3 | 17.121 | 2733.013 | 2.214 | 180.961 | 3.88 | | | 16.729 | 17.694 |
| 2 | piperidine-f1 | MS1 +TIC SIM ESI Frag=70V Gain=0.3 | 18.496 | 18300.904 | 14.823 | 797.049 | 17.07 | | | 18.110 | 19.155 |
| 3 | piperidine-f2 | MS1 +TIC SIM ESI Frag=70V Gain=0.3 | 19.661 | 98460.233 | 79.748 | 3461.337 | 74.14 | | | 19.155 | 20.659 |
| 4 | pyrrolidine-f2 | MS1 +TIC SIM ESI Frag=70V Gain=0.3 | 21.123 | 3969.436 | 3.215 | 229.479 | 4.92 | | | 20.659 | 21.716 |

Reaction performed on analytical scale using **GDM-NT-350**: 94:6 rr (**2a**:**3a**), –45% ee favoring (*S*)-**2a**.

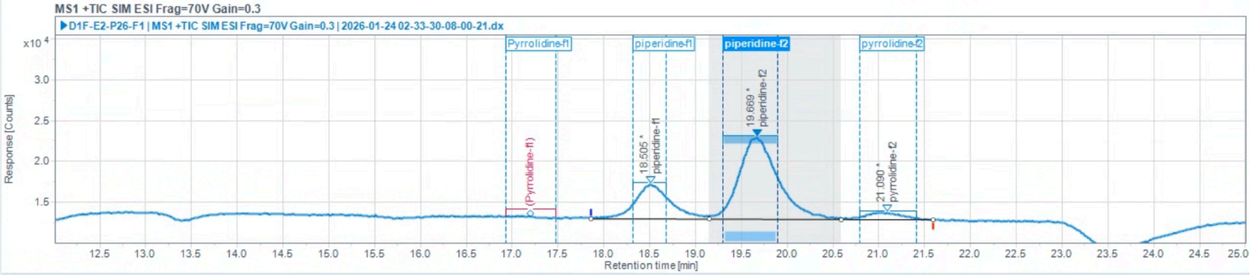

Injection Results

Peaks | Summary

| # | Name | Signal description | RT (min) | Area (Counts·s) | Area% | Height (Counts) | Height% | Amount | Concentration | Start time (min) | End time (min) |
|---|---|---|---|---|---|---|---|---|---|---|---|
| 1 | piperidine-f1 | MS1 +TIC SIM ESI Frag=70V Gain=0.3 | 18.505 | 113819.634 | 26.055 | 4175.578 | 27.63 | | | 17.864 | 19.155 |
| 2 | piperidine-f2 | MS1 +TIC SIM ESI Frag=70V Gain=0.3 | 19.669 | 297941.707 | 68.202 | 10046.695 | 66.48 | | | 19.155 | 20.591 |
| 3 | pyrrolidine-f2 | MS1 +TIC SIM ESI Frag=70V Gain=0.3 | 21.090 | 25089.895 | 5.743 | 890.241 | 5.89 | | | 20.591 | 21.590 |

Reaction performed on analytical scale using **GDM-NT-356**: 99:1 rr (**2a**:**3a**), –73% ee favoring (*S*)-**2a**.

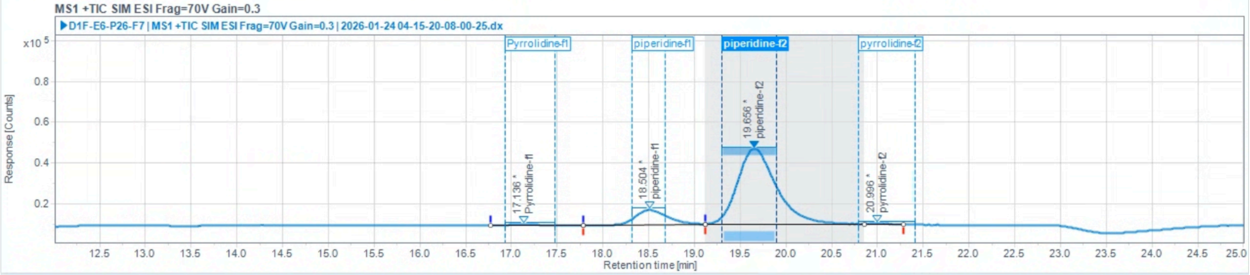

Injection Results

Peaks | Summary

| # | Name | Signal description | RT (min) | Area (Counts·s) | Area% | Height (Counts) | Height% | Amount | Concentration | Start time (min) | End time (min) |
|---|---|---|---|---|---|---|---|---|---|---|---|
| 1 | Pyrrolidine-f1 | MS1 +TIC SIM ESI Frag=70V Gain=0.3 | 17.136 | 6839.246 | 0.532 | 396.744 | 0.88 | | | 16.770 | 17.779 |
| 2 | piperidine-f1 | MS1 +TIC SIM ESI Frag=70V Gain=0.3 | 18.504 | 171826.561 | 13.371 | 7148.996 | 15.91 | | | 17.779 | 19.121 |
| 3 | piperidine-f2 | MS1 +TIC SIM ESI Frag=70V Gain=0.3 | 19.656 | 1100275.660 | 85.619 | 37011.923 | 82.39 | | | 19.121 | 20.854 |
| 4 | pyrrolidine-f2 | MS1 +TIC SIM ESI Frag=70V Gain=0.3 | 20.996 | 6149.236 | 0.479 | 367.435 | 0.82 | | | 20.854 | 21.285 |

Reaction performed on analytical scale using **GDM-NT-359**: 98:2 rr (**2a**:**3a**), –64% ee favoring (*S*)-**2a**.

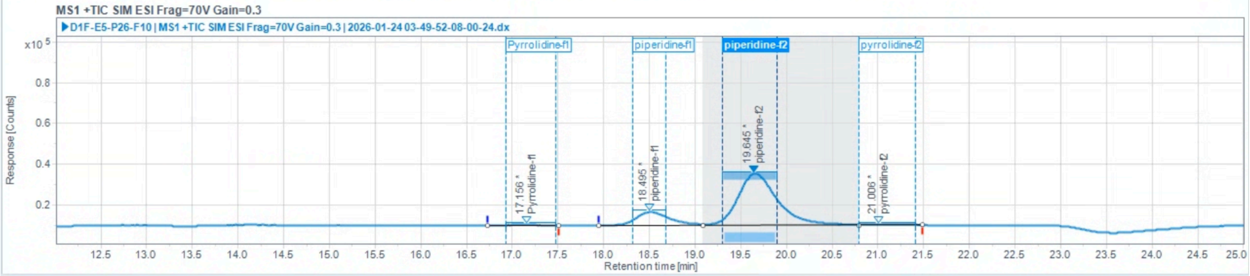

Injection Results

Peaks | Summary

| # | Name | Signal description | RT (min) | Area (Counts·s) | Area% | Height (Counts) | Height% | Amount | Concentration | Start time (min) | End time (min) |
|---|---|---|---|---|---|---|---|---|---|---|---|
| 1 | Pyrrolidine-f1 | MS1 +TIC SIM ESI Frag=70V Gain=0.3 | 17.156 | 3479.897 | 0.366 | 288.288 | 0.88 | | | 16.729 | 17.509 |
| 2 | piperidine-f1 | MS1 +TIC SIM ESI Frag=70V Gain=0.3 | 18.495 | 168656.341 | 17.722 | 6556.707 | 20.02 | | | 17.950 | 19.091 |
| 3 | piperidine-f2 | MS1 +TIC SIM ESI Frag=70V Gain=0.3 | 19.645 | 767392.089 | 80.635 | 25416.737 | 77.61 | | | 19.091 | 20.793 |
| 4 | pyrrolidine-f2 | MS1 +TIC SIM ESI Frag=70V Gain=0.3 | 21.006 | 12155.011 | 1.277 | 487.839 | 1.49 | | | 20.793 | 21.491 |

Reaction performed on analytical scale using **GDM-NT-361**: 95:5 rr (**2a**:**3a**), –23% ee favoring (*S*)-**2a**.

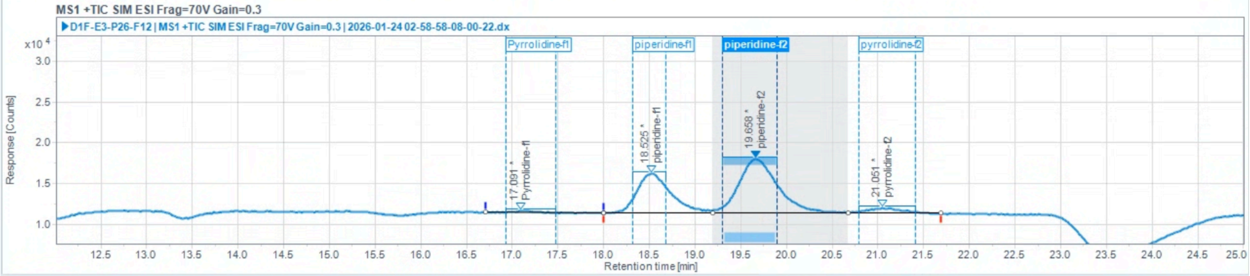

Injection Results

Peaks | Summary

| # | Name | Signal description | RT (min) | Area (Counts·s) | Area% | Height (Counts) | Height% | Amount | Concentration | Start time (min) | End time (min) |
|---|---|---|---|---|---|---|---|---|---|---|---|
| 1 | Pyrrolidine-f1 | MS1 +TIC SIM ESI Frag=70V Gain=0.3 | 17.091 | 2242.808 | 0.658 | 225.259 | 1.82 | | | 16.708 | 17.992 |
| 2 | piperidine-f1 | MS1 +TIC SIM ESI Frag=70V Gain=0.3 | 18.525 | 124639.446 | 36.564 | 4826.447 | 38.92 | | | 17.992 | 19.187 |
| 3 | piperidine-f2 | MS1 +TIC SIM ESI Frag=70V Gain=0.3 | 19.658 | 197672.871 | 57.989 | 6663.560 | 53.73 | | | 19.187 | 20.670 |
| 4 | pyrrolidine-f2 | MS1 +TIC SIM ESI Frag=70V Gain=0.3 | 21.051 | 16326.845 | 4.790 | 686.799 | 5.54 | | | 20.670 | 21.683 |

Reaction performed on analytical scale using **GDM-NT-362**: >99:1 rr (**2a**:**3a**), 89% ee favoring (*R*)-**2a**.

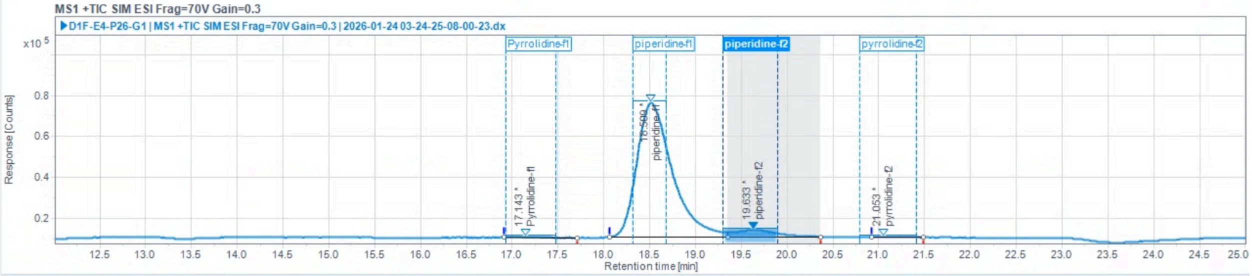

Injection Results

Peaks | Summary

| # | Name | Signal description | RT (min) | Area (Counts·s) | Area% | Height (Counts) | Height% | Amount | Concentration | Start time (min) | End time (min) |
|---|---|---|---|---|---|---|---|---|---|---|---|
| 1 | Pyrrolidine-f1 | MS1 +TIC SIM ESI Frag=70V Gain=0.3 | 17.143 | 2700.514 | 0.144 | 228.557 | 0.33 | | | 16.914 | 17.714 |
| 2 | piperidine-f1 | MS1 +TIC SIM ESI Frag=70V Gain=0.3 | 18.509 | 1770314.949 | 94.320 | 65740.064 | 94.69 | | | 18.062 | 19.358 |
| 3 | piperidine-f2 | MS1 +TIC SIM ESI Frag=70V Gain=0.3 | 19.633 | 102716.631 | 5.473 | 3348.211 | 4.82 | | | 19.358 | 20.360 |
| 4 | pyrrolidine-f2 | MS1 +TIC SIM ESI Frag=70V Gain=0.3 | 21.053 | 1195.317 | 0.064 | 110.671 | 0.16 | | | 20.916 | 21.491 |

**2-phenylpiperidine** (**2a** and **2b**)
Chiral separation method: polar ionic HPLC-MS using *dual* Agilent InfinityLab Poroshell 120 Chiral-V, 2.7 µm, 2.1 × 150 mm; eluent: 40% MeOH (0.2 w% $NH_4CO_2H$), 60% $H_2O$ (20 mM $NH_4CO_2H$, pH 4.0 adjusted with $HCO_2H$); flow rate: 0.2 mL/min. $t_R$ = 33.601 min, $t_S$ = 35.347 min.

Reaction performed on analytical scale using **GDM-NT-507**: 98:2 rr (**2a**:**3a**), 52% ee favoring (*R*)-**2a**.

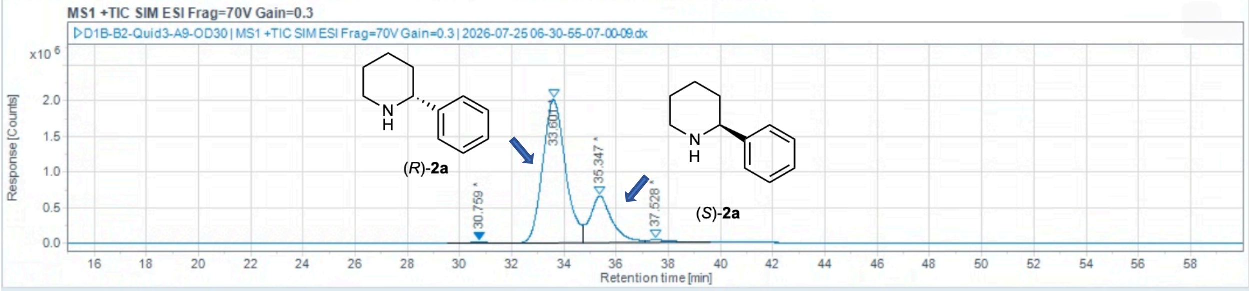

Injection Results

Peaks Summary

| # | Name | Signal description | RT (min) | Area (Counts-s) | Area% | Height (Counts) | Height% | Start time (min) | End time (min) |
|---|---|---|---|---|---|---|---|---|---|
| 1 | | MS1 +TIC SIM ESI Frag=70V Gain=0.3 | 30.759 | 529208.228 | 0.325 | 10346.358 | 0.38 | 29.522 | 31.848 |
| 2 | | MS1 +TIC SIM ESI Frag=70V Gain=0.3 | 33.601 | 121211635.787 | 74.409 | 2018226.112 | 74.29 | 31.848 | 34.710 |
| 3 | | MS1 +TIC SIM ESI Frag=70V Gain=0.3 | 35.347 | 38645343.611 | 23.724 | 651550.876 | 23.98 | 34.710 | 37.096 |
| 4 | | MS1 +TIC SIM ESI Frag=70V Gain=0.3 | 37.528 | 2512253.740 | 1.542 | 36505.739 | 1.34 | 37.096 | 39.601 |

Reaction performed on analytical scale using **GDM-NT-534**: >99:1 rr (**2a**:**3a**), 63% ee favoring (*R*)-**2a**.

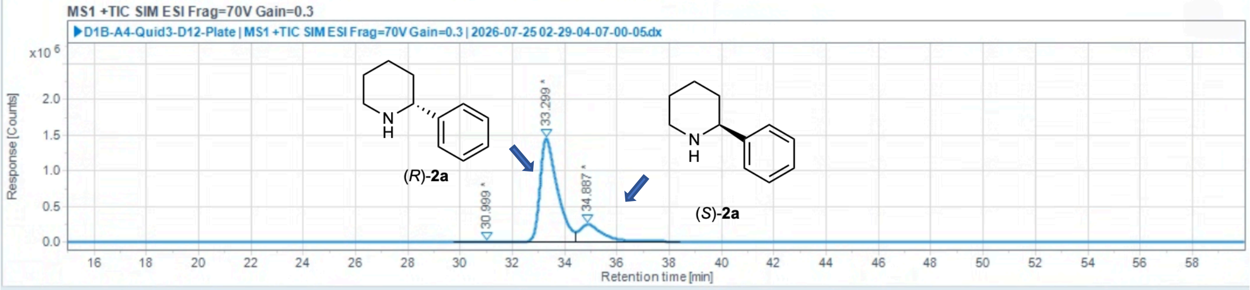

Injection Results

Peaks Summary

| # | Name | Signal description | RT (min) | Area (Counts-s) | Area% | Height (Counts) | Height% | Start time (min) | End time (min) |
|---|---|---|---|---|---|---|---|---|---|
| 1 | | MS1 +TIC SIM ESI Frag=70V Gain=0.3 | 30.999 | 135907.235 | 0.166 | 2556.737 | 0.15 | 29.760 | 32.265 |
| 2 | | MS1 +TIC SIM ESI Frag=70V Gain=0.3 | 33.299 | 66758429.687 | 81.466 | 1449242.570 | 85.51 | 32.265 | 34.436 |
| 3 | | MS1 +TIC SIM ESI Frag=70V Gain=0.3 | 34.887 | 15051558.847 | 18.368 | 242966.759 | 14.34 | 34.436 | 38.408 |

Reaction performed on analytical scale using **GDM-NT-589**: >99:1 rr (**2a**:**3a**), 94% ee favoring (*R*)-**2a**.

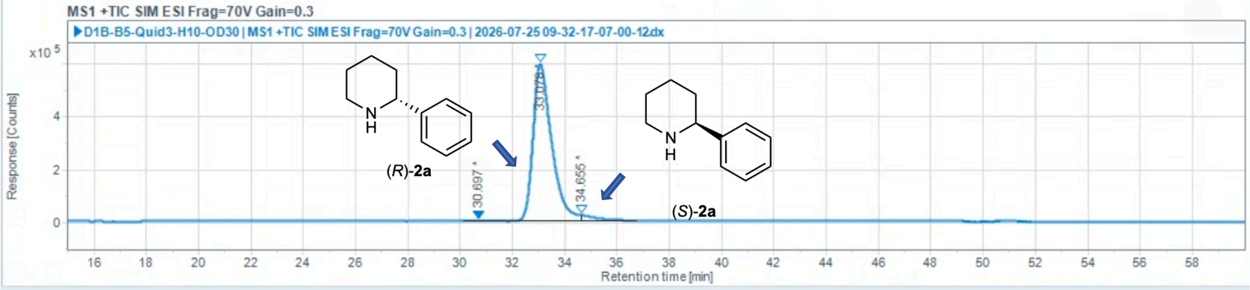

Injection Results

| # | Name | Signal description | RT (min) | Area (Counts·s) | Area% | Height (Counts) | Height% | Start time (min) | End time (min) |
|---|---|---|---|---|---|---|---|---|---|
| 1 | | MS1 +TIC SIM ESI Frag=70V Gain=0.3 | 30.697 | 31343.261 | 0.104 | 826.902 | 0.13 | 30.118 | 32.070 |
| 2 | | MS1 +TIC SIM ESI Frag=70V Gain=0.3 | 33.078 | 29271040.482 | 97.077 | 594655.374 | 96.36 | 32.070 | 34.651 |
| 3 | | MS1 +TIC SIM ESI Frag=70V Gain=0.3 | 34.655 | 849908.043 | 2.819 | 21660.289 | 3.51 | 34.651 | 36.738 |

**Part 3:** Regio- and enantioselectivity analysis of substrate scope study on reactions performed with ***Tm*Pgb-PIP-G7-5455**
Results analyzed on polar ionic chiral HPLC-MS.

**2-phenylpiperidine** (**2a** and **2b**)
Chiral separation method: polar ionic HPLC-MS using *dual* Agilent InfinityLab Poroshell 120 Chiral-V, 2.7 µm, 2.1 × 150 mm; eluent: 40% MeOH (0.2 w% $NH_4CO_2H$), 60% $H_2O$ (20 mM $NH_4CO_2H$, pH 4.0 adjusted with $HCO_2H$); flow rate: 0.2 mL/min. $t_R$ = 31.434 min, $t_S$ = 33.346 min.

(*rac*)-**2a** (commercially available standard):

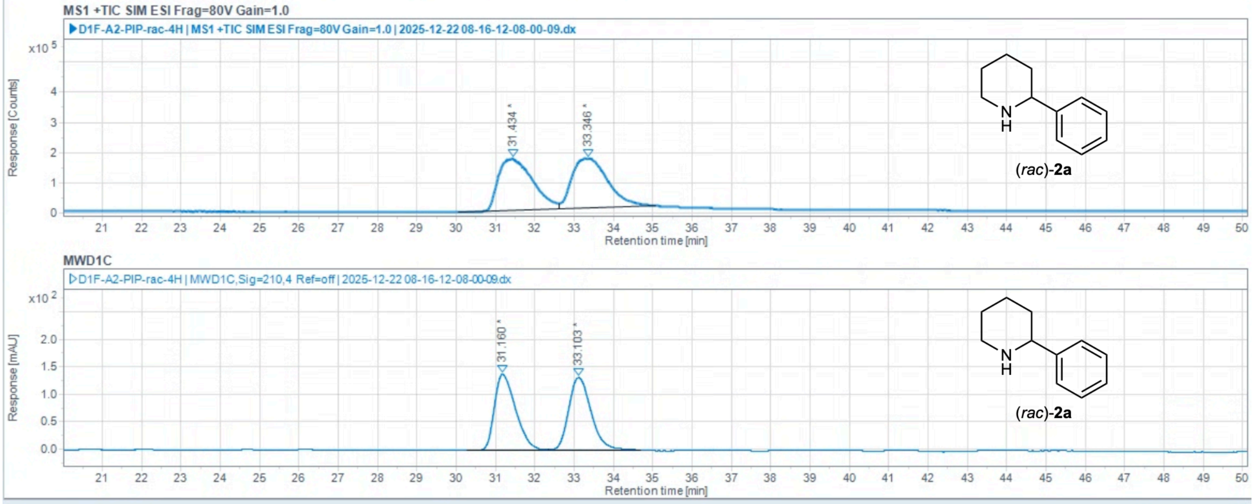

Injection Results

| # | Name | Signal description | RT (min) | Area | Area% | Height | Height% | Amou | Concentra | Start time ( | End t |
|---|---|---|---|---|---|---|---|---|---|---|---|
| 2 | | MS1 +TIC SIM ESI Frag=80V Gain | 31.434 | 10142139.118 | 49.436 | 170479.105 | 50.76 | | | 30.059 | 2.614 |
| 4 | | MS1 +TIC SIM ESI Frag=80V Gain | 33.346 | 10373701.900 | 50.564 | 165350.870 | 49.24 | | | 32.614 | 5.059 |
| 1 | | MWD1C,Sig=210,4 Ref=off | 31.160 | 5211.883 | 49.692 | 139.025 | 51.24 | | | 30.277 | 2.342 |
| 3 | | MWD1C,Sig=210,4 Ref=off | 33.103 | 5276.424 | 50.308 | 132.275 | 48.76 | | | 32.342 | 4.679 |

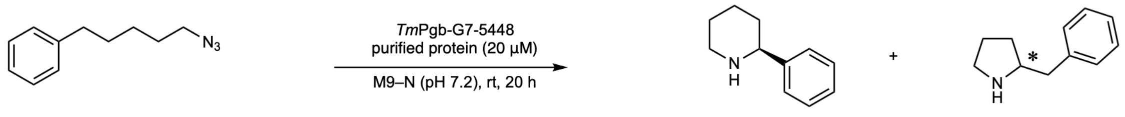

**1a** **2a** **3a**

Reaction performed on analytical scale using ***Tm*Pgb-PIP-G7-5448**: 81:19 rr, >99% ee favoring (*S*)-**2a**.

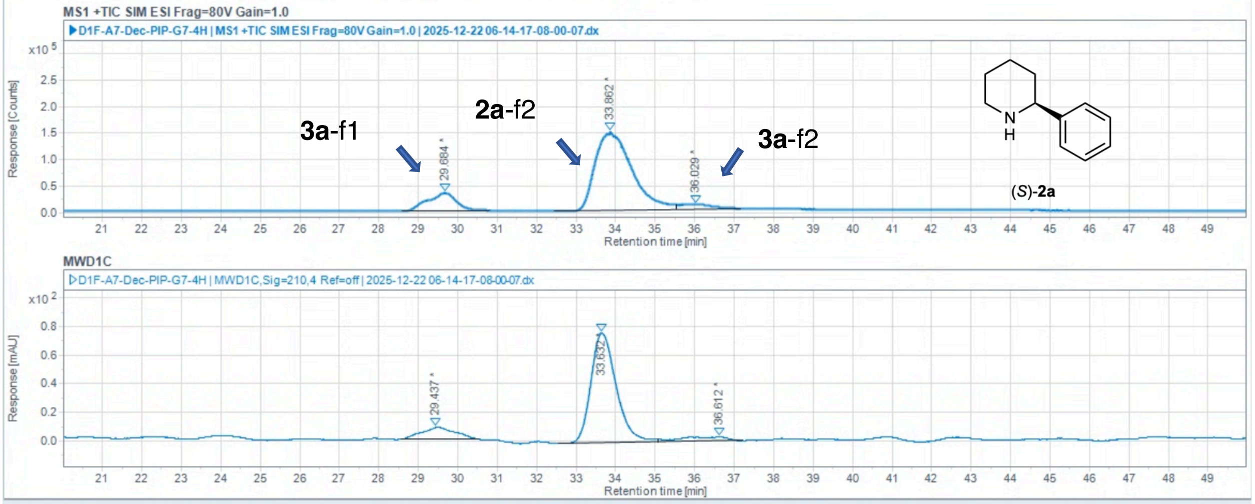

#### Injection Results

Peaks Summary

| # | Name | Signal description | RT (min) | Area | Area% | Height | Height% | Amou | Concentra | Start time ( | End t |
|---|---|---|---|---|---|---|---|---|---|---|---|
| 2 | | MS1 +TIC SIM ESI Frag=80V Gain | 29.684 | 1667976.292 | 14.392 | 33368.448 | 17.60 | | | 28.590 | 0.794 |
| 4 | | MS1 +TIC SIM ESI Frag=80V Gain | 33.862 | 9426293.287 | 81.335 | 147120.682 | 77.61 | | | 32.439 | 5.542 |
| 5 | | MS1 +TIC SIM ESI Frag=80V Gain | 36.029 | 495131.124 | 4.272 | 9086.690 | 4.79 | | | 35.542 | 7.174 |
| 1 | | MWD1C,Sig=210,4 Ref=off | 29.437 | 492.277 | 11.877 | 8.257 | 9.39 | | | 28.550 | 0.545 |
| 3 | | MWD1C,Sig=210,4 Ref=off | 33.632 | 3409.544 | 82.262 | 76.959 | 87.50 | | | 32.539 | 5.078 |
| 6 | | MWD1C,Sig=210,4 Ref=off | 36.612 | 242.942 | 5.861 | 2.733 | 3.11 | | | 35.078 | 7.214 |

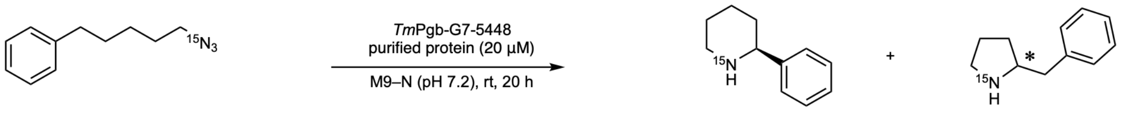

**1b** **2b** **3b**

Reaction performed on analytical scale using ***Tm*Pgb-PIP-G7-5448**: 78:22 rr, >99% ee favoring (*S*)-**2b**.

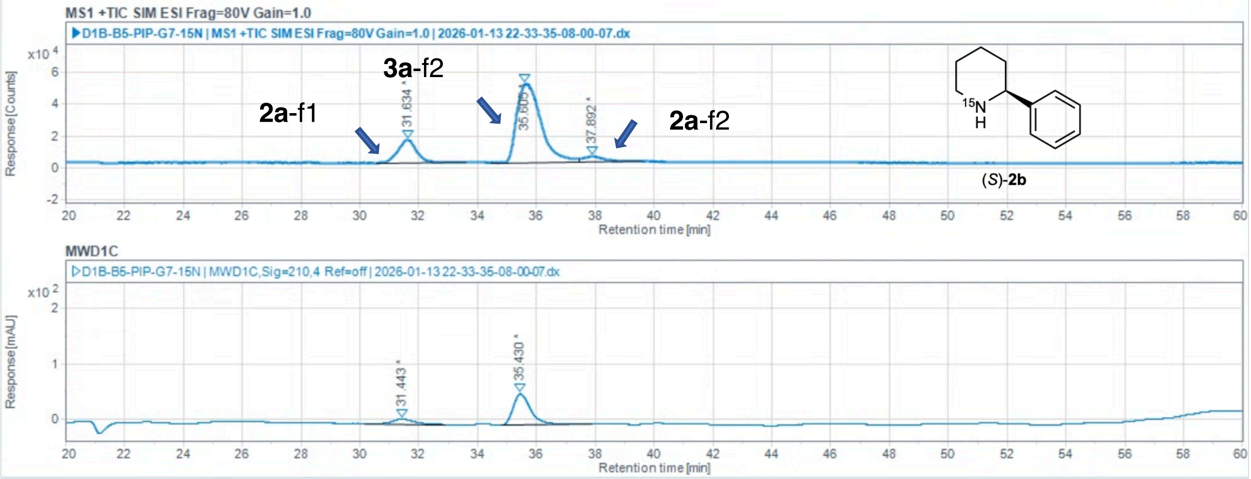

#### Injection Results

Peaks | Summary

| # | Name | Signal description | RT (min) | Area | Area% | Height | Height% | Amou | Concentra | Start time ( | End t |
|---|---|---|---|---|---|---|---|---|---|---|---|
| 2 | | MS1 +TIC SIM ESI Frag=80V Gain | 31.634 | 634789.751 | 17.373 | 14525.135 | 21.39 | | | 30.568 | 3.600 |
| 4 | | MS1 +TIC SIM ESI Frag=80V Gain | 35.605 | 2834640.698 | 77.581 | 49852.346 | 73.42 | | | 34.433 | 7.434 |
| 5 | | MS1 +TIC SIM ESI Frag=80V Gain | 37.892 | 184360.914 | 5.046 | 3521.979 | 5.19 | | | 37.434 | 9.606 |
| 1 | | MWD1C,Sig=210,4 Ref=off | 31.443 | 425.780 | 15.435 | 8.582 | 13.26 | | | 30.152 | 2.887 |
| 3 | | MWD1C,Sig=210,4 Ref=off | 35.430 | 2332.761 | 84.565 | 56.120 | 86.74 | | | 34.789 | 7.881 |

**2-(4-methoxyphenyl)piperidine (2c)**

Chiral separation method: polar ionic HPLC-MS using *dual* Agilent InfinityLab Poroshell 120 Chiral-V, 2.7 µm, 2.1 × 150 mm; eluent: 40% MeOH (0.2 w% $NH_4CO_2H$), 60% $H_2O$ (20 mM $NH_4CO_2H$, pH 4.0 adjusted with $HCO_2H$); flow rate: 0.2 mL/min. $t_R$ = 32.567 min, $t_S$ = 35.979 min.

(*rac*)-**2c** (commercially available standard):

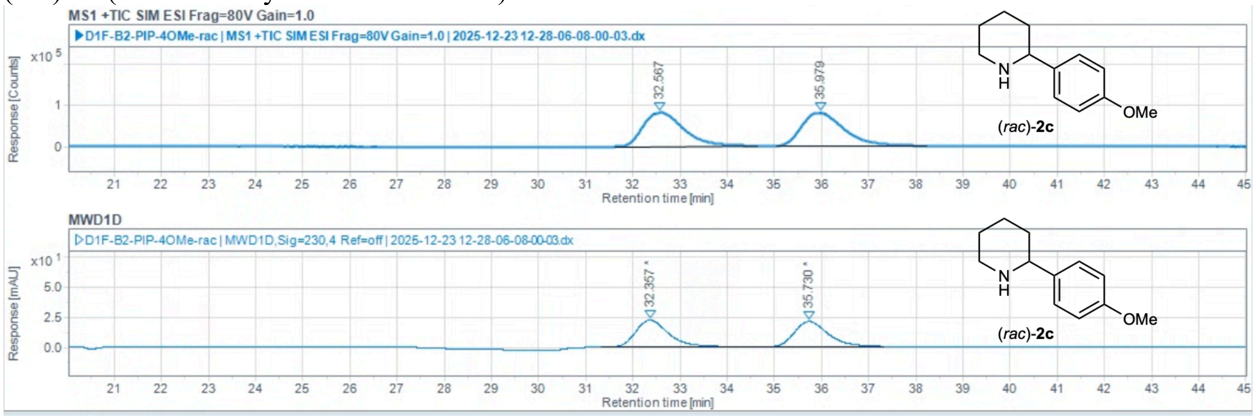

Injection Results

Peaks | Summary

| # | Name | Signal description | RT (min) | Area | Area% | Height | Height% | Amou | Concentra | Start time ( | End t |
|---|---|---|---|---|---|---|---|---|---|---|---|
| 2 | | MS1 +TIC SIM ESI Frag=80V Gain | 32.567 | 4872984.093 | 49.550 | 82639.489 | 50.81 | | | 31.612 | 4.644 |
| 4 | | MS1 +TIC SIM ESI Frag=80V Gain | 35.979 | 4961437.264 | 50.450 | 80006.498 | 49.19 | | | 35.045 | 8.231 |
| 1 | | MWD1D,Sig=230,4 Ref=off | 32.357 | 1073.228 | 51.465 | 22.122 | 51.71 | | | 31.314 | 4.432 |
| 3 | | MWD1D,Sig=230,4 Ref=off | 35.730 | 1012.123 | 48.535 | 20.659 | 48.29 | | | 34.432 | 7.303 |

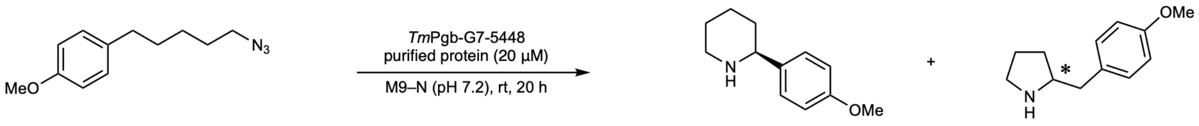

**1c** **2c** **3c**

Reaction performed on analytical scale using ***Tm*Pgb-PIP-G7-5448**: 84:16 rr, >99% ee favoring (*S*)-**2c**.

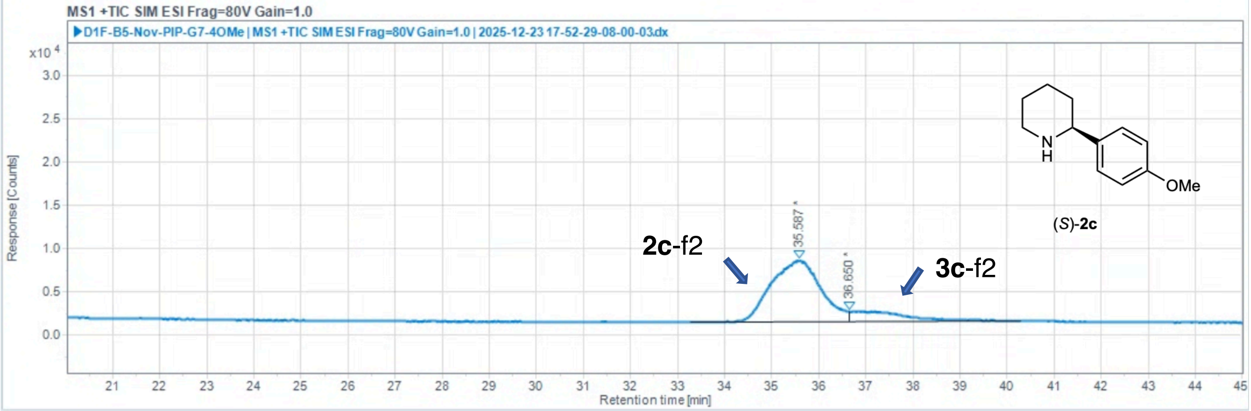

Injection Results

Peaks | Summary

| # | Name | Signal description | RT (min) | Area (Counts·s) | Area% | Height (Counts) | Height% | Amou | Concentra | Start time ( | End t |
|---|---|---|---|---|---|---|---|---|---|---|---|
| 1 | | MS1 +TIC SIM ESI Frag=80V Gain | 35.587 | 512009.489 | 84.336 | 7140.513 | 86.14 | | | 33.275 | 6.634 |
| 2 | | MS1 +TIC SIM ESI Frag=80V Gain | 36.650 | 95097.175 | 15.664 | 1149.014 | 13.86 | | | 36.634 | 0.277 |

co-injection of (*rac*)-**2c** and (*rac*)-**3c**

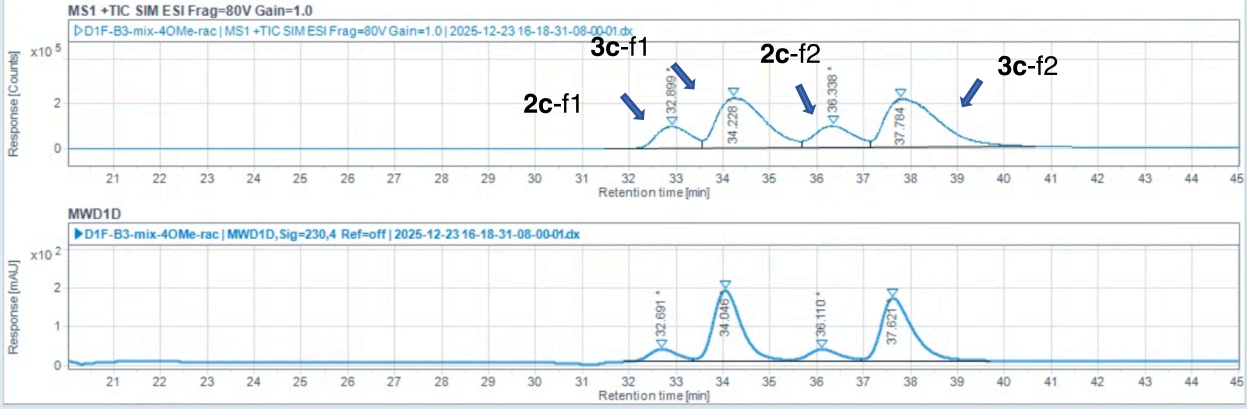

Injection Results

Peaks | Summary

| # | Name | Signal description | RT (min) | Area | Area% | Height | Height% | Amou | Concentra | Start time ( | End t |
|---|---|---|---|---|---|---|---|---|---|---|---|
| 2 | | MS1 +TIC SIM ESI Frag=80V Gain | 32.899 | 4783823.291 | 11.124 | 97694.623 | 15.44 | | | 31.461 | 3.565 |
| 4 | | MS1 +TIC SIM ESI Frag=80V Gain | 34.228 | 15656170.914 | 36.406 | 223658.349 | 35.36 | | | 33.565 | 5.669 |
| 6 | | MS1 +TIC SIM ESI Frag=80V Gain | 36.338 | 5413053.959 | 12.587 | 95594.015 | 15.11 | | | 35.669 | 7.129 |
| 8 | | MS1 +TIC SIM ESI Frag=80V Gain | 37.784 | 17151217.736 | 39.883 | 215641.185 | 34.09 | | | 37.129 | 0.669 |
| 1 | | MWD1D,Sig=230,4 Ref=off | 32.691 | 1356.519 | 7.489 | 32.291 | 7.89 | | | 31.882 | 3.343 |
| 3 | | MWD1D,Sig=230,4 Ref=off | 34.046 | 7714.704 | 42.593 | 183.687 | 44.87 | | | 33.343 | 5.446 |
| 5 | | MWD1D,Sig=230,4 Ref=off | 36.110 | 1431.716 | 7.905 | 30.493 | 7.45 | | | 35.446 | 6.956 |
| 7 | | MWD1D,Sig=230,4 Ref=off | 37.621 | 7609.647 | 42.013 | 162.936 | 39.80 | | | 36.956 | 9.679 |

**2-(p-tolyl)piperidine (2d)**

Chiral separation method: polar ionic HPLC-MS using *dual* Agilent InfinityLab Poroshell 120 Chiral-V, 2.7 µm, 2.1 × 150 mm; eluent: 40% MeOH (0.2 w% $NH_4CO_2H$), 60% $H_2O$ (20 mM $NH_4CO_2H$, pH 4.0 adjusted with $HCO_2H$); flow rate: 0.2 mL/min. $t_R$ = 30.346 min, $t_S$ = 36.371 min.

(*rac*)-**2d** (commercially available standard):

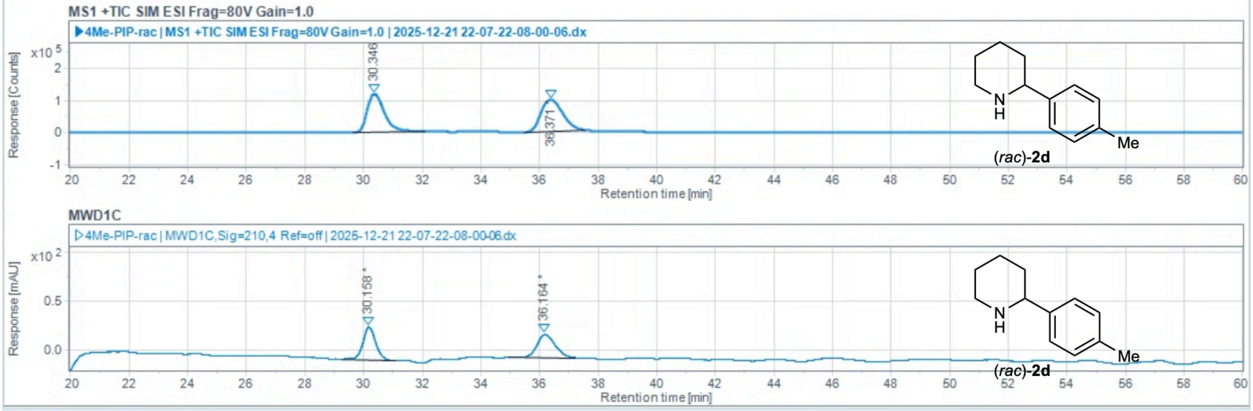

#### Injection Results

Peaks | Summary

| # | Name | Signal description | RT (min) | Area | Area% | Height | Height% | Amou | Concentra | Start time ( | End t |
|---|---|---|---|---|---|---|---|---|---|---|---|
| 2 | | MS1 +TIC SIM ESI Frag=80V Gain | 30.346 | 4844107.781 | 48.892 | 118443.148 | 54.46 | | | 29.653 | 2.094 |
| 4 | | MS1 +TIC SIM ESI Frag=80V Gain | 36.371 | 5063746.785 | 51.108 | 99030.507 | 45.54 | | | 35.493 | 7.595 |
| 1 | | MWD1C,Sig=210,4 Ref=off | 30.158 | 1000.324 | 49.030 | 33.637 | 58.39 | | | 29.227 | 1.091 |
| 3 | | MWD1C,Sig=210,4 Ref=off | 36.164 | 1039.886 | 50.970 | 23.973 | 41.61 | | | 34.938 | 7.238 |

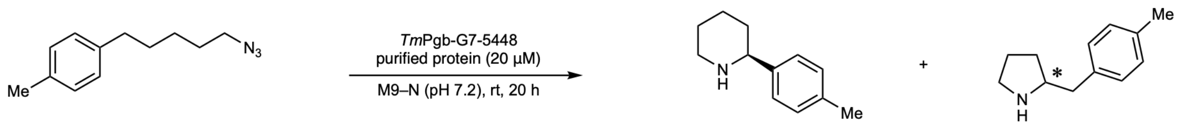

**1d** **2d** **3d**

Reaction performed on analytical scale using ***Tm*Pgb-PIP-G7-5448**: >99% ee favoring (*S*)-**2d**.

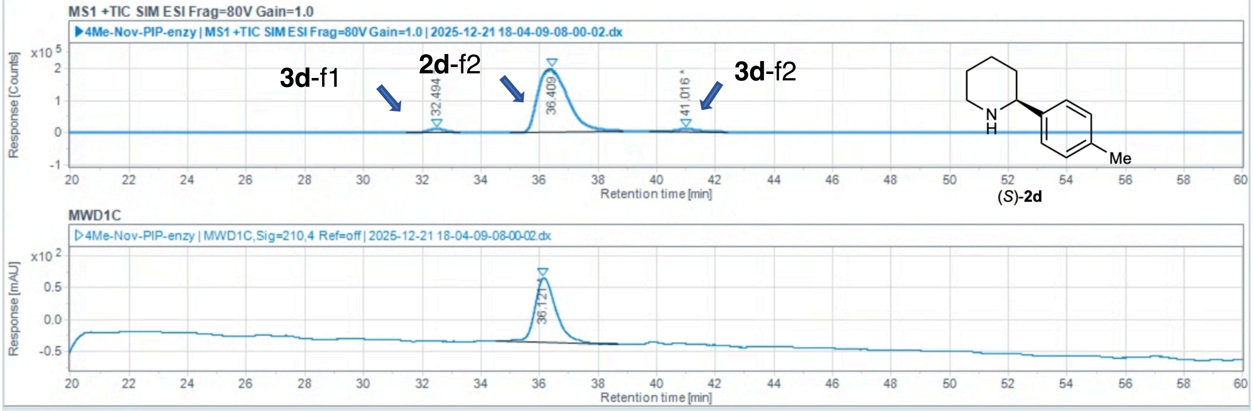

#### Injection Results

Peaks | Summary

| # | Name | Signal description | RT (min) | Area | Area% | Height | Height% | Amou | Concentra | Start time ( | End t |
|---|---|---|---|---|---|---|---|---|---|---|---|
| 1 | | MS1 +TIC SIM ESI Frag=80V Gain | 32.494 | 445134.209 | 3.051 | 10995.479 | 5.08 | | | 31.443 | 3.270 |
| 3 | | MS1 +TIC SIM ESI Frag=80V Gain | 36.409 | 13736715.937 | 94.144 | 196923.774 | 91.06 | | | 35.018 | 8.831 |
| 4 | | MS1 +TIC SIM ESI Frag=80V Gain | 41.016 | 409372.701 | 2.806 | 8336.744 | 3.86 | | | 39.745 | 2.446 |
| 2 | | MWD1C,Sig=210,4 Ref=off | 36.121 | 5029.515 | 100.000 | 100.953 | 100.00 | | | 34.502 | 8.712 |

co-injection of (*rac*)-**2d** and (*rac*)-**3d**

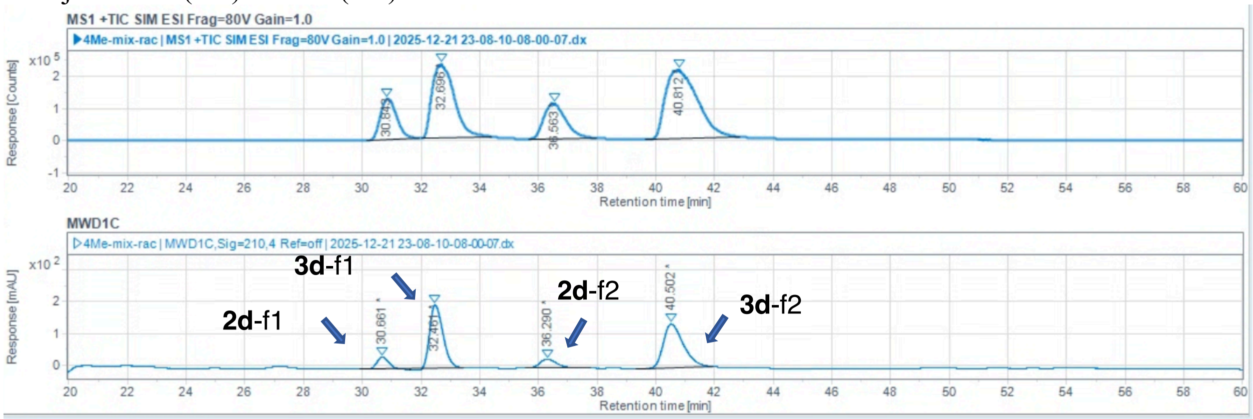

Injection Results

Peaks | Summary

| # | Name | Signal description | RT (min) | Area | Area% | Height | Height% | Amou | Concentra | Start time ( | End t |
|---|---|---|---|---|---|---|---|---|---|---|---|
| 2 | | MS1 +TIC SIM ESI Frag=80V Gain | 30.843 | 4900426.268 | 12.677 | 127078.854 | 18.66 | | | 30.168 | 1.935 |
| 4 | | MS1 +TIC SIM ESI Frag=80V Gain | 32.696 | 12049355.890 | 31.171 | 230068.223 | 33.78 | | | 32.014 | 4.421 |
| 6 | | MS1 +TIC SIM ESI Frag=80V Gain | 36.563 | 5642213.493 | 14.596 | 109617.977 | 16.10 | | | 35.691 | 7.992 |
| 8 | | MS1 +TIC SIM ESI Frag=80V Gain | 40.812 | 16064213.268 | 41.557 | 214266.271 | 31.46 | | | 39.658 | 2.870 |
| 1 | | MWD1C,Sig=210,4 Ref=off | 30.661 | 893.573 | 6.114 | 37.203 | 9.26 | | | 29.901 | 1.606 |
| 3 | | MWD1C,Sig=210,4 Ref=off | 32.461 | 6070.685 | 41.540 | 198.902 | 49.51 | | | 31.606 | 3.431 |
| 5 | | MWD1C,Sig=210,4 Ref=off | 36.290 | 1145.972 | 7.842 | 28.398 | 7.07 | | | 35.572 | 7.794 |
| 7 | | MWD1C,Sig=210,4 Ref=off | 40.502 | 6503.774 | 44.504 | 137.270 | 34.17 | | | 39.340 | 1.958 |

**2-(4-chlorophenyl)piperidine (2e)**

Chiral separation method: polar ionic HPLC-MS using *dual* Agilent InfinityLab Poroshell 120 Chiral-V, 2.7 µm, 2.1 × 150 mm; eluent: 40% MeOH (0.2 w% $NH_4CO_2H$), 60% $H_2O$ (20 mM $NH_4CO_2H$, pH 4.0 adjusted with $HCO_2H$); flow rate: 0.2 mL/min. $t_R$ = 38.233 min, $t_S$ = 41.414 min.

(*rac*)-**2e** (commercially available standard):

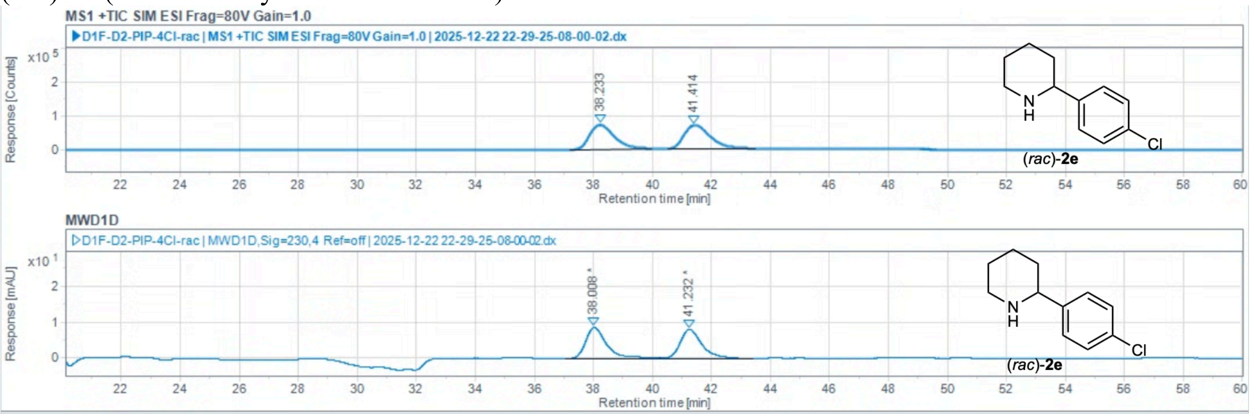

Injection Results

Peaks | Summary

| # | Name | Signal description | RT (min) | Area | Area% | Height | Height% | Amou | Concentra | Start time ( | End t |
|---|---|---|---|---|---|---|---|---|---|---|---|
| 2 | | MS1 +TIC SIM ESI Frag=80V Gain | 38.233 | 4139118.572 | 49.188 | 72203.414 | 50.94 | | | 37.225 | 9.969 |
| 4 | | MS1 +TIC SIM ESI Frag=80V Gain | 41.414 | 4275804.727 | 50.812 | 69550.966 | 49.06 | | | 40.520 | 3.506 |
| 1 | | MWD1D,Sig=230,4 Ref=off | 38.008 | 431.194 | 51.190 | 8.651 | 51.73 | | | 37.027 | 0.230 |
| 3 | | MWD1D,Sig=230,4 Ref=off | 41.232 | 411.153 | 48.810 | 8.073 | 48.27 | | | 40.230 | 3.393 |

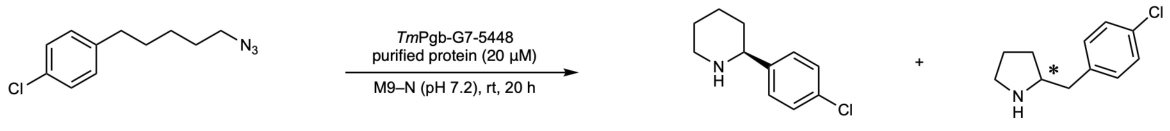

**1e** **2e** **3e**

Reaction performed on analytical scale using ***Tm*Pgb-PIP-G7-5448**: >99% ee favoring (*S*)-**2e**.

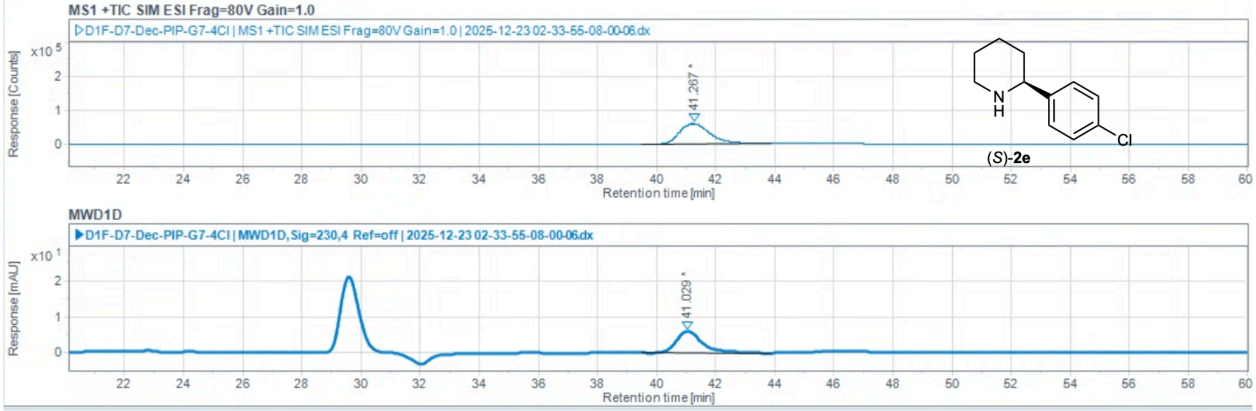

Injection Results

Peaks | Summary

| # | Name | Signal description | RT (min) | Area | Area% | Height | Height% | Amou | Concentra | Start time ( | End t |
|---|---|---|---|---|---|---|---|---|---|---|---|
| 2 | | MS1 +TIC SIM ESI Frag=80V Gain | 41.267 | 4257149.474 | 100.000 | 59748.585 | 100.00 | | | 39.479 | 3.868 |
| 1 | | MWD1D,Sig=230,4 Ref=off | 41.029 | 372.864 | 100.000 | 6.018 | 100.00 | | | 39.479 | 3.868 |

co-injection of (*rac*)-**2e** and (*rac*)-**3e**

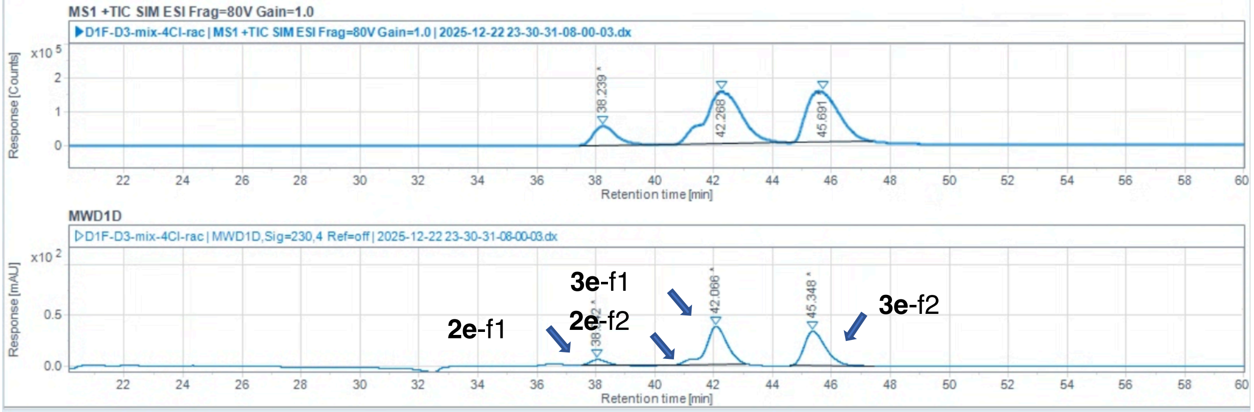

Injection Results

Peaks | Summary

| # | Name | Signal description | RT (min) | Area | Area% | Height | Height% | Amou | Concentra | Start time ( | End t |
|---|---|---|---|---|---|---|---|---|---|---|---|
| 2 | | MS1 +TIC SIM ESI Frag=80V Gain | 38.239 | 3035759.805 | 10.940 | 56526.417 | 15.59 | | | 37.423 | 0.111 |
| 4 | | MS1 +TIC SIM ESI Frag=80V Gain | 42.268 | 13374636.613 | 48.199 | 155494.708 | 42.89 | | | 40.111 | 4.421 |
| 6 | | MS1 +TIC SIM ESI Frag=80V Gain | 45.691 | 11338319.075 | 40.861 | 150516.240 | 41.52 | | | 44.421 | 7.387 |
| 1 | | MWD1D,Sig=230,4 Ref=off | 38.042 | 175.447 | 4.712 | 5.433 | 7.15 | | | 37.502 | 0.507 |
| 3 | | MWD1D,Sig=230,4 Ref=off | 42.066 | 1921.098 | 51.595 | 37.354 | 49.15 | | | 40.507 | 3.117 |
| 5 | | MWD1D,Sig=230,4 Ref=off | 45.348 | 1626.868 | 43.693 | 33.210 | 43.70 | | | 44.580 | 7.427 |

**2-(4-fluorophenyl)piperidine** (**2f**)

Chiral separation method: polar ionic HPLC-MS using *dual* Agilent InfinityLab Poroshell 120 Chiral-V, 2.7 µm, 2.1 × 150 mm; eluent: 40% MeOH (0.2 w% $NH_4CO_2H$), 60% $H_2O$ (20 mM $NH_4CO_2H$, pH 4.0 adjusted with $HCO_2H$); flow rate: 0.2 mL/min. $t_R$ = 27.976 min, $t_S$ = 29.024 min.

(*rac*)-**2f** (commercially available standard):

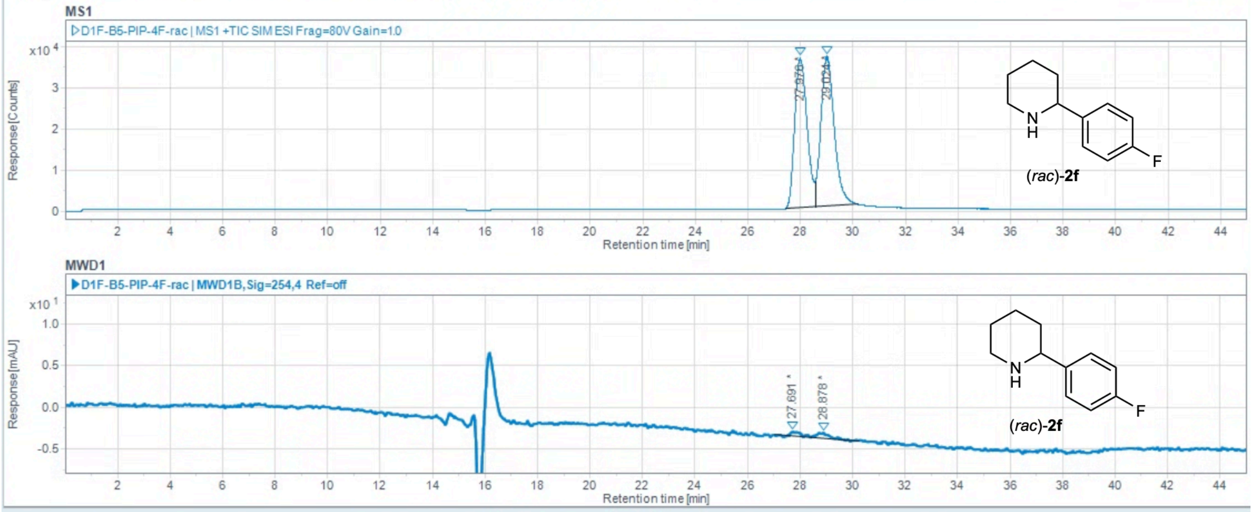

Injection Results

Peaks Summary

| # | Name | Signal description | RT (min) | Area | Area% | Height | Height% | Amou | Concentra | Start time ( | End t |
|---|---|---|---|---|---|---|---|---|---|---|---|
| 2 | | MS1 +TIC SIM ESI Frag=80V Gain | 27.976 | 1191246.065 | 47.993 | 36558.659 | 50.07 | | | 27.446 | 8.565 |
| 4 | | MS1 +TIC SIM ESI Frag=80V Gain | 29.024 | 1290856.144 | 52.007 | 36450.674 | 49.93 | | | 28.565 | 0.176 |
| 1 | | MWD1B,Sig=254,4 Ref=off | 27.691 | 13.202 | 41.550 | 0.592 | 44.77 | | | 27.011 | 8.410 |
| 3 | | MWD1B,Sig=254,4 Ref=off | 28.878 | 18.572 | 58.450 | 0.730 | 55.23 | | | 28.410 | 0.233 |

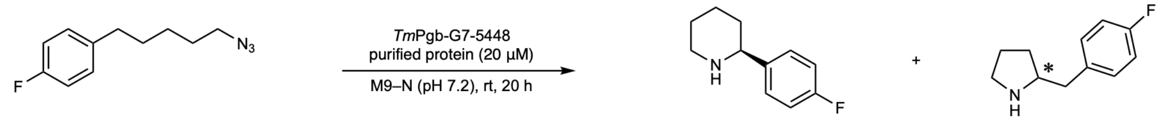

**1f** **2f** **3f**

Reaction performed on analytical scale using ***Tm*Pgb-PIP-G7-5448**: >99% ee favoring (*S*)-**2f**.

By analyzing the following spectrum together with the calculated average regioselectivity based on reverse-phase HPLC-MS using C18 column, the composition of the following spectrum is: major peak (co-elution of 6% **3f**-f1, 84% **2f**-f2) and minor peak (10% **3f**-f2).

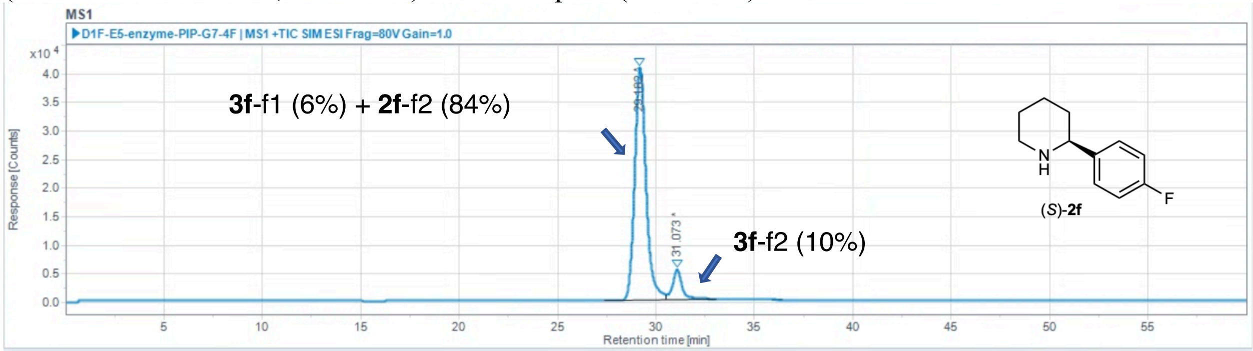

Injection Results

Peaks Summary

| # | Name | Signal description | RT (min) | Area (Counts·s) | Area% | Height (Counts) | Height% | Amou | Concentra | Start time ( | End t |
|---|---|---|---|---|---|---|---|---|---|---|---|
| 1 | | MS1 +TIC SIM ESI Frag=80V Gain | 29.182 | 1643918.847 | 90.225 | 40776.212 | 88.36 | | | 27.430 | 0.540 |
| 2 | | MS1 +TIC SIM ESI Frag=80V Gain | 31.073 | 178110.789 | 9.775 | 5369.829 | 11.64 | | | 30.540 | 3.084 |

co-injection of (*rac*)-**2f** and (*rac*)-**3f**

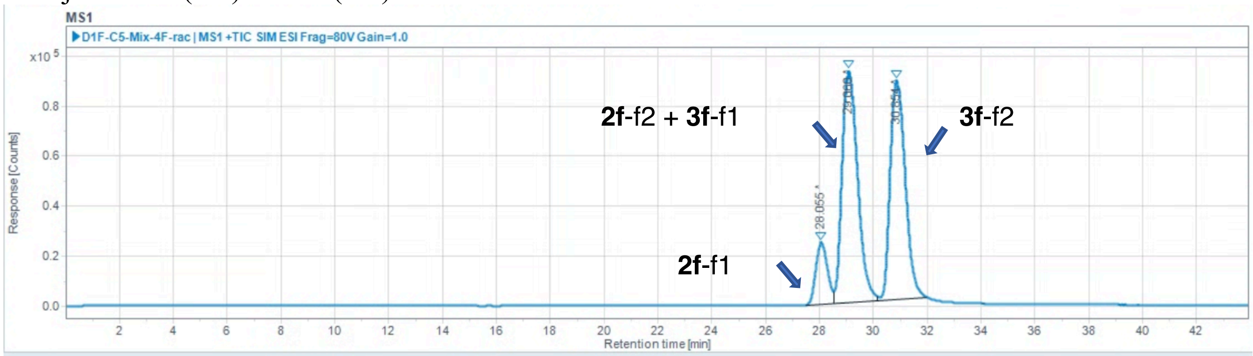

Injection Results

Peaks | Summary

| # | Name | Signal description | RT (min) | Area (Counts·s) | Area% | Height (Counts) | Height% | Amou | Concentra | Start time ( | End t |
|---|---|---|---|---|---|---|---|---|---|---|---|
| 1 | | MS1 +TIC SIM ESI Frag=80V Gain | 28.055 | 736548.867 | 9.378 | 24882.195 | 12.11 | | | 27.478 | 8.513 |
| 2 | | MS1 +TIC SIM ESI Frag=80V Gain | 29.060 | 3723211.113 | 47.407 | 92884.544 | 45.19 | | | 28.513 | 0.168 |
| 3 | | MS1 +TIC SIM ESI Frag=80V Gain | 30.854 | 3394038.109 | 43.215 | 87776.457 | 42.70 | | | 30.168 | 1.988 |

**2-(3-fluorophenyl)piperidine (2j)**

Chiral separation method: polar ionic HPLC-MS using *dual* Agilent InfinityLab Poroshell 120 Chiral-V, 2.7 µm, 2.1 × 150 mm; eluent: 40% MeOH (0.2 w% $NH_4CO_2H$), 60% $H_2O$ (20 mM $NH_4CO_2H$, pH 4.0 adjusted with $HCO_2H$); flow rate: 0.2 mL/min. $t_R$ = 31.205 min, $t_S$ = 33.295 min.

(*rac*)-**2j** (commercially available standard):

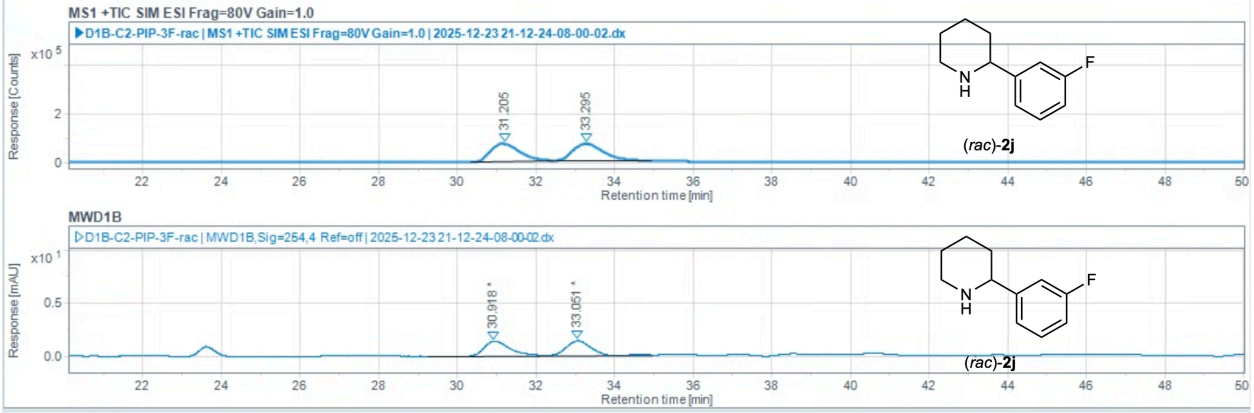

Injection Results

Peaks | Summary

| # | Name | Signal description | RT (min) | Area | Area% | Height | Height% | Amou | Concentra | Start time ( | End t |
|---|---|---|---|---|---|---|---|---|---|---|---|
| 2 | | MS1 +TIC SIM ESI Frag=80V Gain | 31.205 | 3730119.493 | 50.015 | 73781.663 | 51.47 | | | 30.349 | 2.492 |
| 4 | | MS1 +TIC SIM ESI Frag=80V Gain | 33.295 | 3727889.147 | 49.985 | 69565.262 | 48.53 | | | 32.498 | 4.934 |
| 1 | | MWD1B,Sig=254,4 Ref=off | 30.918 | 58.558 | 50.005 | 1.376 | 50.07 | | | 29.262 | 2.175 |
| 3 | | MWD1B,Sig=254,4 Ref=off | 33.051 | 58.546 | 49.995 | 1.373 | 49.93 | | | 32.175 | 4.970 |

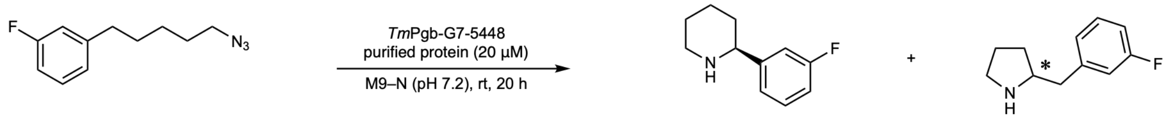

**1j** **2j** **3j**

Reaction performed on analytical scale using ***Tm*Pgb-PIP-G7-5448**: >99% ee favoring (*S*)-**2j**.

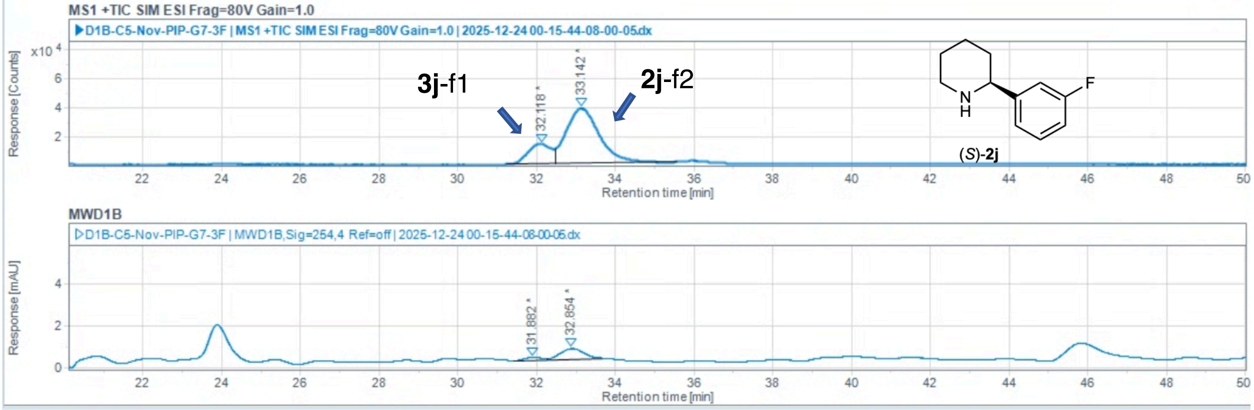

Injection Results

Peaks | Summary

| # | Name | Signal description | RT (min) | Area | Area% | Height | Height% | Amou | Concentra | Start time ( | End t |
|---|---|---|---|---|---|---|---|---|---|---|---|
| 2 | | MS1 +TIC SIM ESI Frag=80V Gain | 32.118 | 559072.859 | 20.184 | 13689.162 | 26.63 | | | 31.220 | 2.465 |
| 4 | | MS1 +TIC SIM ESI Frag=80V Gain | 33.142 | 2210741.958 | 79.816 | 37707.826 | 73.37 | | | 32.465 | 5.548 |
| 1 | | MWD1B,Sig=254,4 Ref=off | 31.882 | 4.201 | 16.747 | 0.142 | 22.07 | | | 31.405 | 2.258 |
| 3 | | MWD1B,Sig=254,4 Ref=off | 32.854 | 20.884 | 83.253 | 0.501 | 77.93 | | | 32.258 | 3.663 |

co-injection of (*rac*)-**2j** and (*rac*)-**3j**

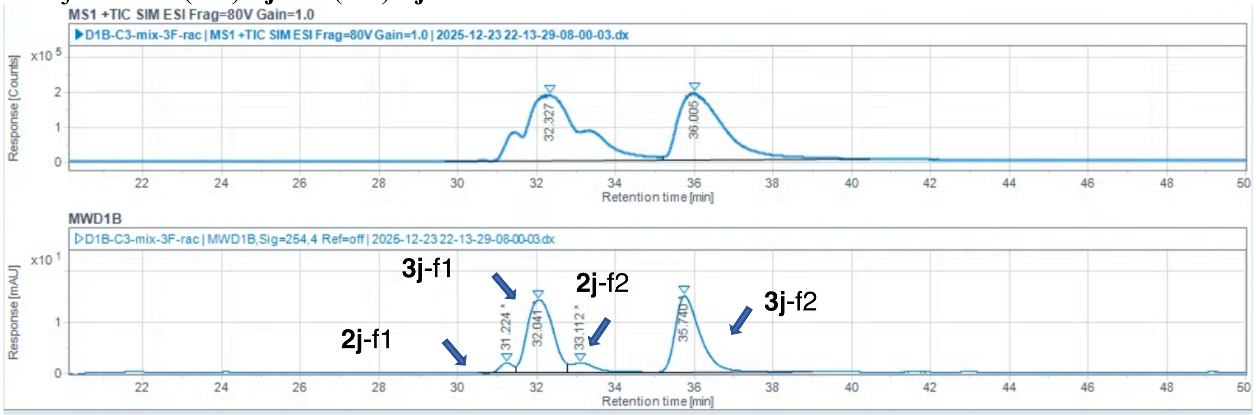

Injection Results

Peaks | Summary

| # | Name | Signal description | RT (min) | Area | Area% | Height | Height% | Amou | Concentra | Start time ( | End t |
|---|---|---|---|---|---|---|---|---|---|---|---|
| 3 | | MS1 +TIC SIM ESI Frag=80V Gain | 32.327 | 19773204.123 | 55.941 | 191414.825 | 49.52 | | | 29.675 | 5.206 |
| 6 | | MS1 +TIC SIM ESI Frag=80V Gain | 36.005 | 15573625.495 | 44.059 | 195126.221 | 50.48 | | | 35.206 | 0.439 |
| 1 | | MWD1B,Sig=254,4 Ref=off | 31.224 | 44.021 | 3.101 | 1.943 | 5.89 | | | 30.510 | 1.461 |
| 2 | | MWD1B,Sig=254,4 Ref=off | 32.041 | 644.785 | 45.416 | 14.237 | 43.17 | | | 31.461 | 2.787 |
| 4 | | MWD1B,Sig=254,4 Ref=off | 33.112 | 86.319 | 6.080 | 1.913 | 5.80 | | | 32.787 | 4.973 |
| 5 | | MWD1B,Sig=254,4 Ref=off | 35.740 | 644.598 | 45.403 | 14.890 | 45.14 | | | 34.973 | 9.013 |

**2-(naphthalen-2-yl)piperidine (2l)**

Chiral separation method: polar ionic HPLC-MS using *dual* Agilent InfinityLab Poroshell 120 Chiral-V, 2.7 µm, 2.1 × 150 mm; eluent: 40% MeOH (0.2 w% $NH_4CO_2H$), 60% $H_2O$ (20 mM $NH_4CO_2H$, pH 4.0 adjusted with $HCO_2H$); flow rate: 0.2 mL/min. $t_R$ = 64.074 min, $t_S$ = 75.475 min.

(*rac*)-**2l** (commercially available standard):

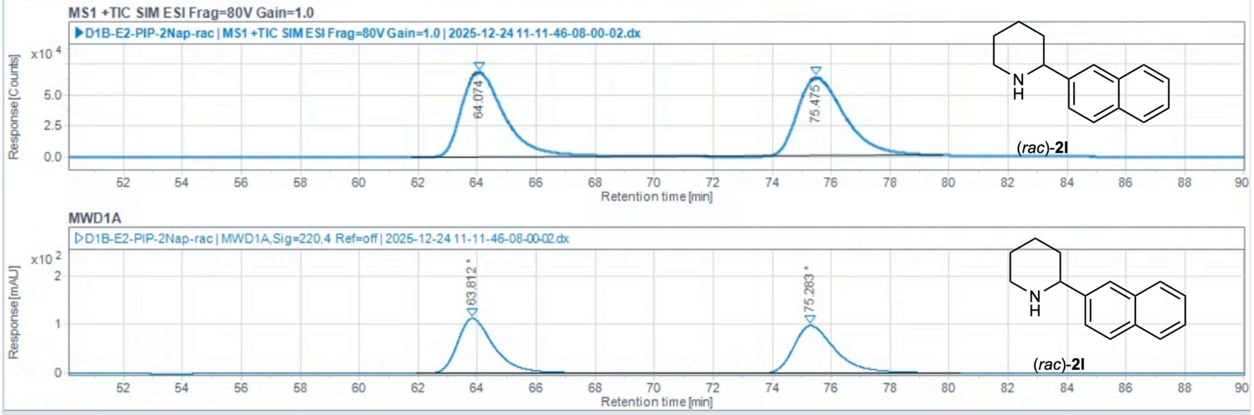

Injection Results

Peaks | Summary

| # | Name | Signal description | RT (min) | Area | Area% | Height | Height% | Amou | Concentra | Start time ( | End t |
|---|---|---|---|---|---|---|---|---|---|---|---|
| 2 | | MS1 +TIC SIM ESI Frag=80V Gain | 64.074 | 6866458.716 | 50.046 | 68128.588 | 51.85 | | | 61.766 | 0.304 |
| 4 | | MS1 +TIC SIM ESI Frag=80V Gain | 75.475 | 6853896.816 | 49.954 | 63261.452 | 48.15 | | | 70.304 | 9.795 |
| 1 | | MWD1A,Sig=220,4 Ref=off | 63.812 | 9781.591 | 50.474 | 113.998 | 53.60 | | | 61.925 | 9.470 |
| 3 | | MWD1A,Sig=220,4 Ref=off | 75.283 | 9598.050 | 49.526 | 98.666 | 46.40 | | | 69.470 | 0.391 |

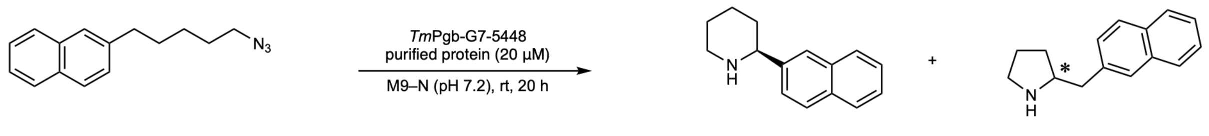

**1l** **2l** **3l**

Reaction performed on analytical scale using ***Tm*Pgb-PIP-G7-5448**: >99% ee favoring (*S*)-**2l**.

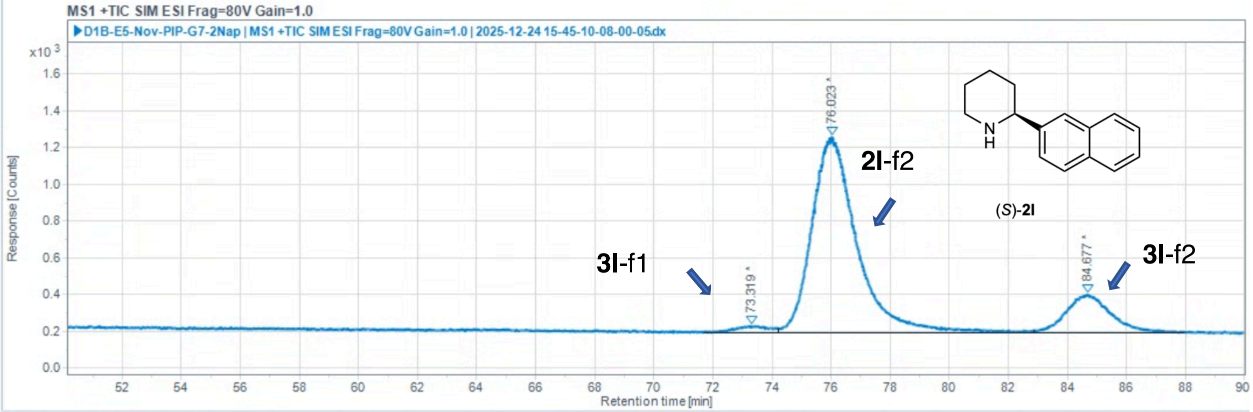

Injection Results

Peaks | Summary

| # | Name | Signal description | RT (min) | Area (Counts·s) | Area% | Height (Counts) | Height% | Amou | Concentra | Start time ( | End t |
|---|---|---|---|---|---|---|---|---|---|---|---|
| 1 | | MS1 +TIC SIM ESI Frag=80V Gain | 73.319 | 2721.216 | 2.015 | 35.363 | 2.70 | | | 71.666 | 4.195 |
| 2 | | MS1 +TIC SIM ESI Frag=80V Gain | 76.023 | 111538.137 | 82.580 | 1067.800 | 81.67 | | | 74.195 | 2.849 |
| 3 | | MS1 +TIC SIM ESI Frag=80V Gain | 84.677 | 20807.920 | 15.406 | 204.256 | 15.62 | | | 82.849 | 7.986 |

co-injection of (*rac*)-**2l** and (*rac*)-**3l**

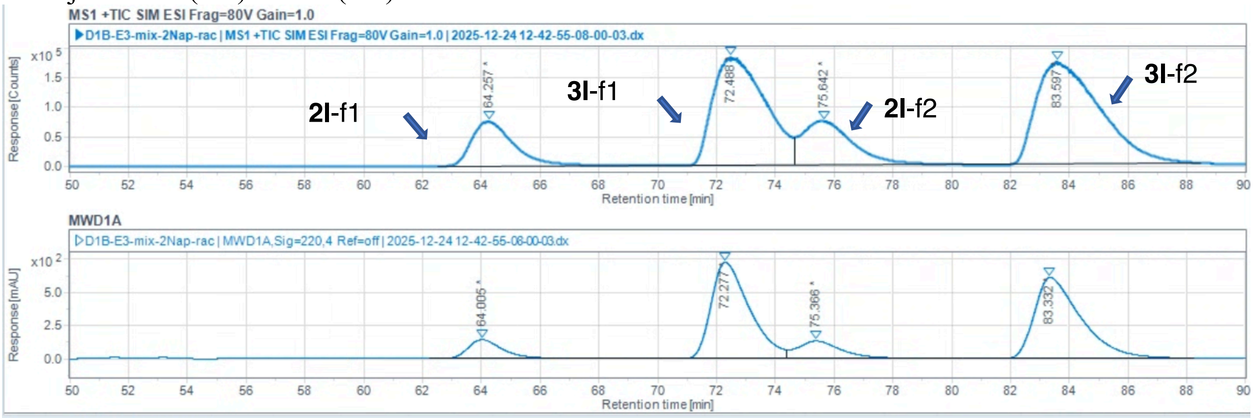

Injection Results

Peaks | Summary

| # | Name | Signal description | RT (min) | Area | Area% | Height | Height% | Amou | Concentra | Start time ( | End t |
|---|---|---|---|---|---|---|---|---|---|---|---|
| 2 | | MS1 +TIC SIM ESI Frag=80V Gain | 64.257 | 7429091.587 | 11.580 | 76304.937 | 15.01 | | | 62.521 | 8.795 |
| 4 | | MS1 +TIC SIM ESI Frag=80V Gain | 72.488 | 22467663.451 | 35.021 | 184149.241 | 36.22 | | | 68.795 | 4.641 |
| 6 | | MS1 +TIC SIM ESI Frag=80V Gain | 75.642 | 8744396.594 | 13.630 | 75069.078 | 14.76 | | | 74.641 | 9.994 |
| 8 | | MS1 +TIC SIM ESI Frag=80V Gain | 83.597 | 25513239.464 | 39.769 | 172923.615 | 34.01 | | | 79.994 | 8.492 |
| 1 | | MWD1A,Sig=220,4 Ref=off | 64.005 | 11920.408 | 7.586 | 142.924 | 8.90 | | | 62.243 | 8.716 |
| 3 | | MWD1A,Sig=220,4 Ref=off | 72.277 | 65518.931 | 41.697 | 724.926 | 45.15 | | | 68.716 | 4.394 |
| 5 | | MWD1A,Sig=220,4 Ref=off | 75.366 | 13444.843 | 8.556 | 130.755 | 8.14 | | | 74.394 | 9.954 |
| 7 | | MWD1A,Sig=220,4 Ref=off | 83.332 | 66246.218 | 42.160 | 607.128 | 37.81 | | | 79.954 | 8.254 |

**Part 4:** Regio- and enantioselectivity analysis of substrate scope study on reactions performed with **GDM-NT-401**.
Results analyzed on polar ionic chiral HPLC-MS, normal phase HPLC, or chiral GC.

**2-phenylpiperidine** (**2a** and **2b**)
Chiral separation method: polar ionic HPLC-MS using *dual* Agilent InfinityLab Poroshell 120 Chiral-V, 2.7 µm, 2.1 × 150 mm; eluent: 40% MeOH (0.2 w% $NH_4CO_2H$), 60% $H_2O$ (20 mM $NH_4CO_2H$, pH 4.0 adjusted with $HCO_2H$); flow rate: 0.2 mL/min. $t_R$ = 33.270 min, $t_S$ = 35.262 min.

(*rac*)-**2a** (commercially available standard):

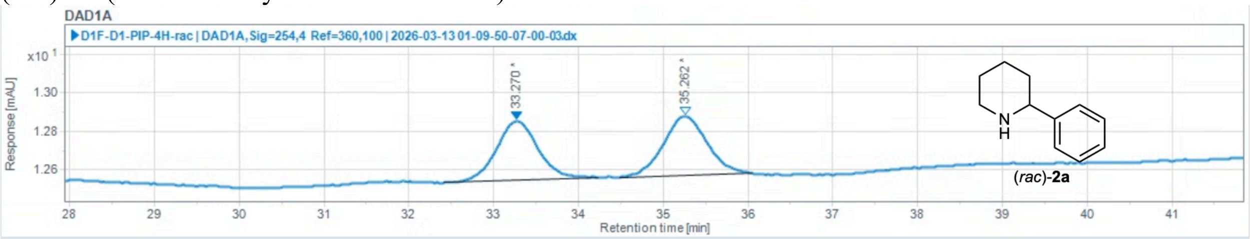

Injection Results

Peaks Summary

| # | Name | Signal description | RT (min) | Area (mAU·s) | Area% | Height (mAU) | Height% | Amount | Concentration | Start tim |
|---|---|---|---|---|---|---|---|---|---|---|
| 1 | | DAD1A,Sig=254,4 Ref=360,100 | 33.270 | 10.460 | 49.012 | 0.307 | 50.25 | | | |
| 2 | | DAD1A,Sig=254,4 Ref=360,100 | 35.262 | 10.881 | 50.988 | 0.304 | 49.75 | | | |

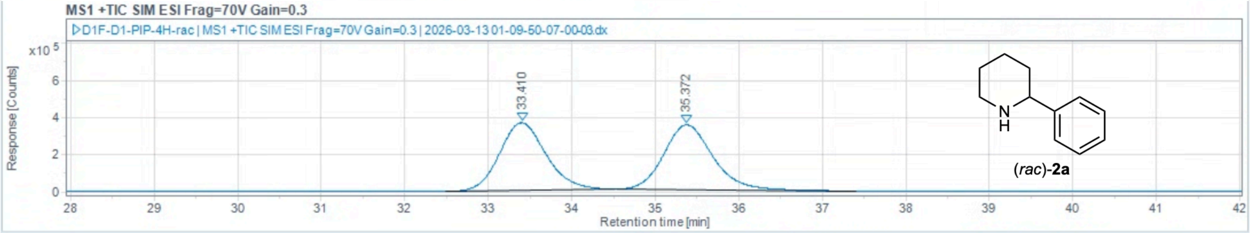

Injection Results

Peaks Summary

| # | Name | Signal description | RT (min) | Area (Counts·s) | Area% | Height (Counts) | Height% | Amount | Concentration | Start tim |
|---|---|---|---|---|---|---|---|---|---|---|
| 1 | | MS1 +TIC SIM ESI Frag=70V Gain=0.3 | 33.410 | 14206364.978 | 49.736 | 365616.260 | 51.07 | | | |
| 2 | | MS1 +TIC SIM ESI Frag=70V Gain=0.3 | 35.372 | 14356974.653 | 50.264 | 350287.991 | 48.93 | | | |

(*S*)-**2a** (commercially available standard):

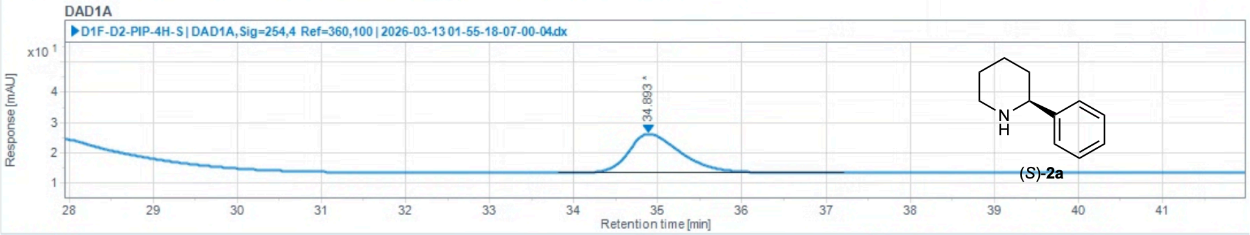

Injection Results

Peaks Summary

| # | Name | Signal description | RT (min) | Area (mAU·s) | Area% | Height (mAU) | Height% | Amount | Concentration | Start tim |
|---|---|---|---|---|---|---|---|---|---|---|
| 1 | | DAD1A,Sig=254,4 Ref=360,100 | 34.893 | 567.528 | 100.000 | 12.937 | 100.00 | | | |

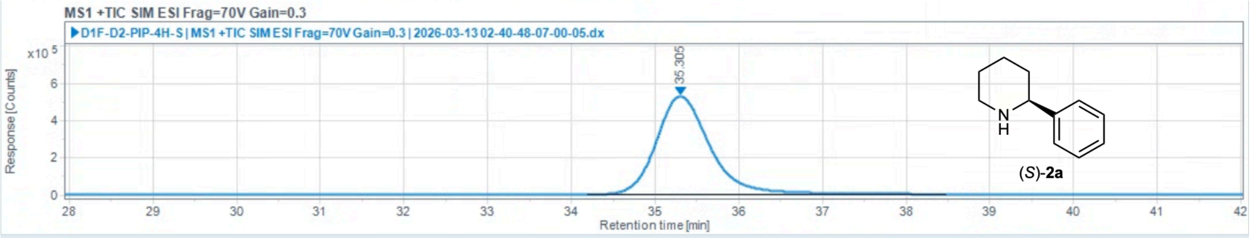

Injection Results

Peaks Summary

| # | Name | Signal description | RT (min) | Area (Counts·s) | Area% | Height (Counts) | Height% | Amount | Concentration | Start tim |
|---|---|---|---|---|---|---|---|---|---|---|
| 1 | | MS1 +TIC SIM ESI Frag=70V Gain=0.3 | 35.305 | 23513403.082 | 100.000 | 527373.776 | 100.00 | | | |

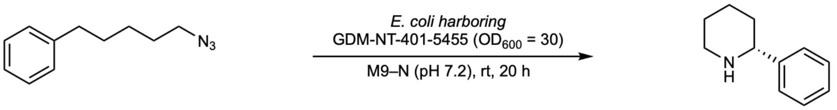

**1a** **2a**

Reaction performed on analytical scale using **GDM-NT-401-5455**: 95% ee favoring (*R*)-**2a**.

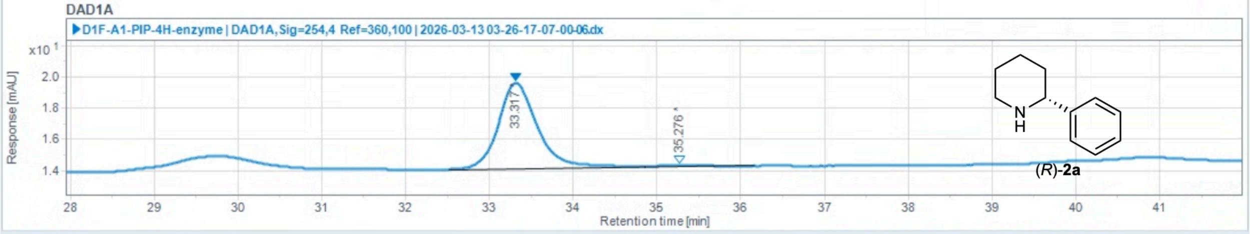

Injection Results

Peaks Summary

| # | Name | Signal description | RT (min) | Area (mAU·s) | Area% | Height (mAU) | Height% | Amount | Concentration | Start tim |
|---|---|---|---|---|---|---|---|---|---|---|
| 1 | | DAD1A,Sig=254,4 Ref=360,100 | 33.317 | 179.212 | 97.929 | 5.481 | 98.32 | | | |
| 2 | | DAD1A,Sig=254,4 Ref=360,100 | 35.276 | 3.790 | 2.071 | 0.094 | 1.68 | | | |

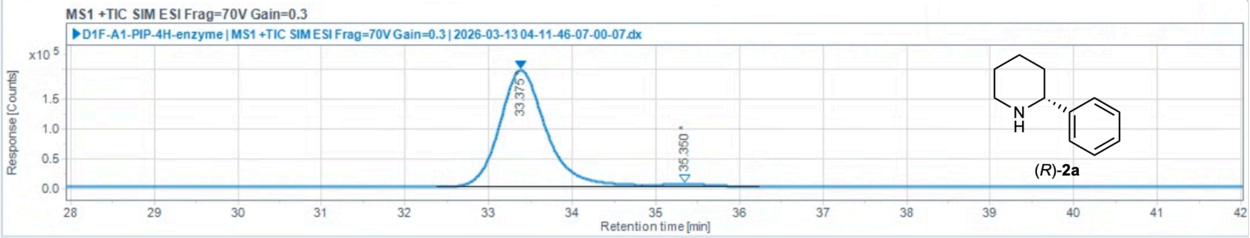

Injection Results

Peaks Summary

| # | Name | Signal description | RT (min) | Area (Counts·s) | Area% | Height (Counts) | Height% | Amount | Concentration | Start tim |
|---|---|---|---|---|---|---|---|---|---|---|
| 1 | | MS1 +TIC SIM ESI Frag=70V Gain=0.3 | 33.375 | 7585259.311 | 97.548 | 194831.248 | 97.90 | | | |
| 2 | | MS1 +TIC SIM ESI Frag=70V Gain=0.3 | 35.350 | 190639.788 | 2.452 | 4182.668 | 2.10 | | | |

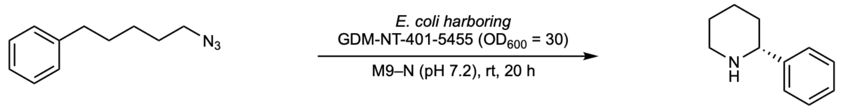

**1a** **2a**

Reaction performed on 1.0 mmol scale using **GDM-NT-401-5455**: 95% ee favoring (*R*)-**2a**.

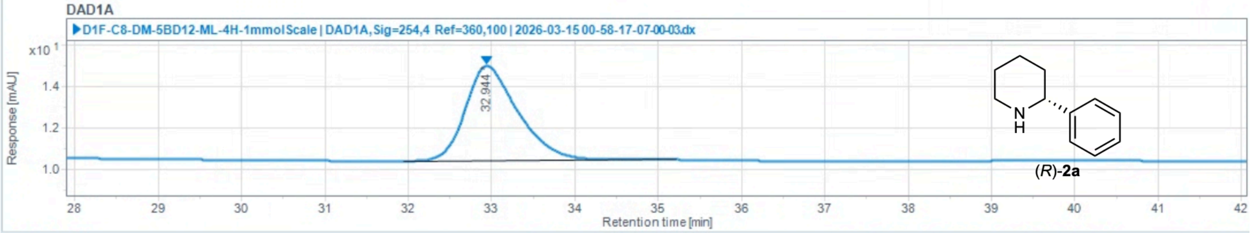

#### Injection Results

Peaks Summary

| # | Name | Signal description | RT (min) | Area (mAU·s) | Area% | Height (mAU) | Height% | Amount | Concentration | Start tim |
|---|---|---|---|---|---|---|---|---|---|---|
| 1 | | DAD1A,Sig=254,4 Ref=360,100 | 32.944 | 201.221 | 100.000 | 4.544 | 100.00 | | | |

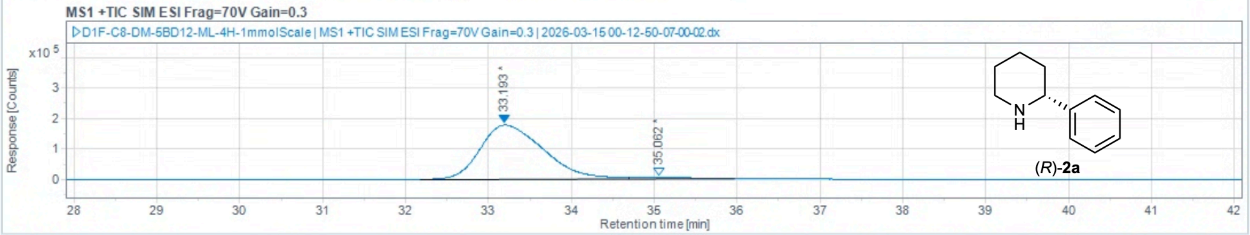

#### Injection Results

Peaks Summary

| # | Name | Signal description | RT (min) | Area (Counts·s) | Area% | Height (Counts) | Height% | Amount | Concentration | Start tim |
|---|---|---|---|---|---|---|---|---|---|---|
| 1 | | MS1 +TIC SIM ESI Frag=70V Gain=0.3 | 33.193 | 9410385.178 | 97.643 | 177249.125 | 97.51 | | | |
| 2 | | MS1 +TIC SIM ESI Frag=70V Gain=0.3 | 35.062 | 227126.251 | 2.357 | 4525.898 | 2.49 | | | |

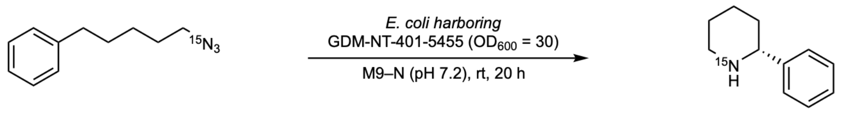

**1b** **2b**

Reaction performed on analytical scale using **GDM-NT-401-5455**: 95% ee favoring (*R*)-**2b**.

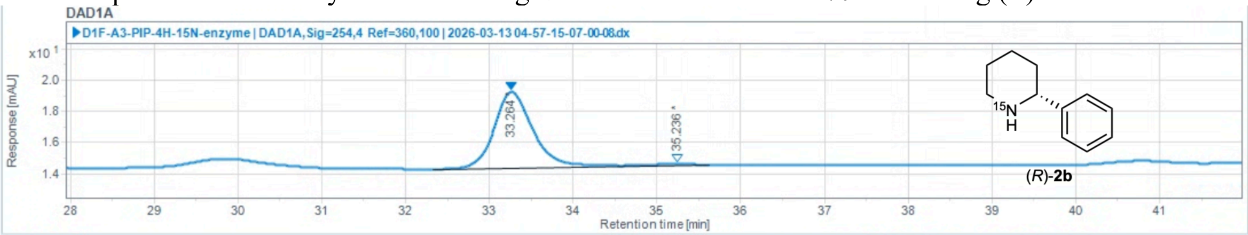

Injection Results

Peaks | Summary

| # | Name | Signal description | RT (min) | Area (mAU·s) | Area% | Height (mAU) | Height% | Amount | Concentration | Start tim |
|---|---|---|---|---|---|---|---|---|---|---|
| 1 | | DAD1A,Sig=254,4 Ref=360,100 | 33.264 | 157.519 | 97.708 | 4.878 | 98.17 | | | |
| 2 | | DAD1A,Sig=254,4 Ref=360,100 | 35.236 | 3.695 | 2.292 | 0.091 | 1.83 | | | |

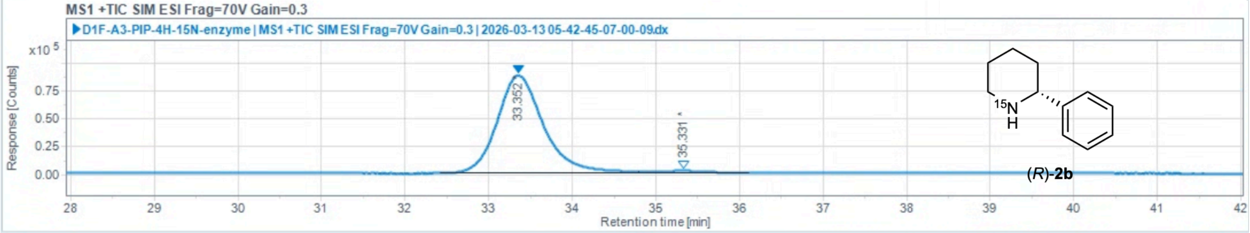

Injection Results

Peaks | Summary

| # | Name | Signal description | RT (min) | Area (Counts·s) | Area% | Height (Counts) | Height% | Amount | Concentration | Start tim |
|---|---|---|---|---|---|---|---|---|---|---|
| 1 | | MS1 +TIC SIM ESI Frag=70V Gain=0.3 | 33.352 | 3398754.648 | 97.660 | 87738.341 | 97.96 | | | |
| 2 | | MS1 +TIC SIM ESI Frag=70V Gain=0.3 | 35.331 | 81450.521 | 2.340 | 1829.352 | 2.04 | | | |

**2-(4-methoxyphenyl)piperidine (2c)**
Chiral separation method: polar ionic HPLC-MS using *dual* Agilent InfinityLab Poroshell 120 Chiral-V, 2.7 µm, 2.1 × 150 mm; eluent: 40% MeOH (0.2 w% $NH_4CO_2H$), 60% $H_2O$ (20 mM $NH_4CO_2H$, pH 4.0 adjusted with $HCO_2H$); flow rate: 0.2 mL/min. $t_R$ = 33.460 min, $t_S$ = 36.557 min.

(*rac*)-**2c** (commercially available standard):

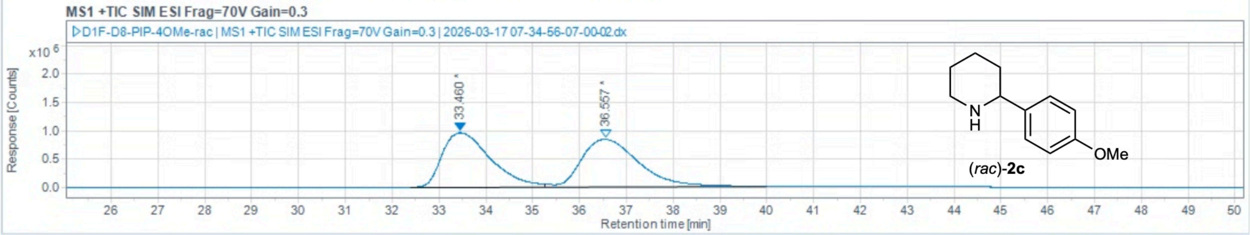

Injection Results

Peaks Summary

| # | Name | Signal description | RT (min) | Area (Counts·s) | Area% | Height (Counts) | Height% | Amount | Concentration | Start tim |
|---|---|---|---|---|---|---|---|---|---|---|
| 1 | | MS1 +TIC SIM ESI Frag=70V Gain=0.3 | 33.460 | 68734223.408 | 49.186 | 956143.898 | 53.39 | | | |
| 2 | | MS1 +TIC SIM ESI Frag=70V Gain=0.3 | 36.557 | 71008360.767 | 50.814 | 834764.032 | 46.61 | | | |

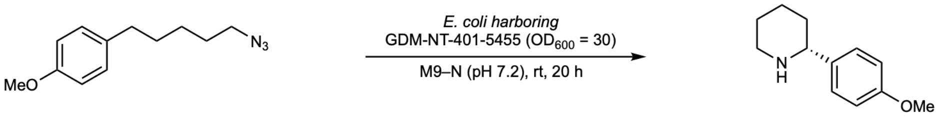

**1c** **2c**

Reaction performed on analytical scale using **GDM-NT-401-5455**: 90% ee favoring (*R*)-**2c**.

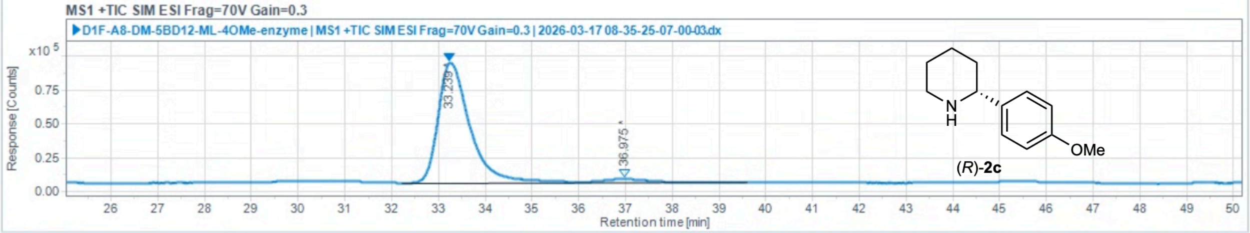

Injection Results

Peaks Summary

| # | Name | Signal description | RT (min) | Area (Counts·s) | Area% | Height (Counts) | Height% | Amount | Concentration | Start tim |
|---|---|---|---|---|---|---|---|---|---|---|
| 1 | | MS1 +TIC SIM ESI Frag=70V Gain=0.3 | 33.239 | 4186469.225 | 94.999 | 88813.374 | 96.89 | | | |
| 2 | | MS1 +TIC SIM ESI Frag=70V Gain=0.3 | 36.975 | 220403.147 | 5.001 | 2854.616 | 3.11 | | | |

**2-(p-tolyl)piperidine (2d)**

Chiral separation method: polar ionic HPLC-MS using *dual* Agilent InfinityLab Poroshell 120 Chiral-V, 2.7 µm, 2.1 × 150 mm; eluent: 40% MeOH (0.2 w% $NH_4CO_2H$), 60% $H_2O$ (20 mM $NH_4CO_2H$, pH 4.0 adjusted with $HCO_2H$); flow rate: 0.2 mL/min. $t_R$ = 31.902 min, $t_S$ = 38.191 min.

(*rac*)-**2d** (commercially available standard):

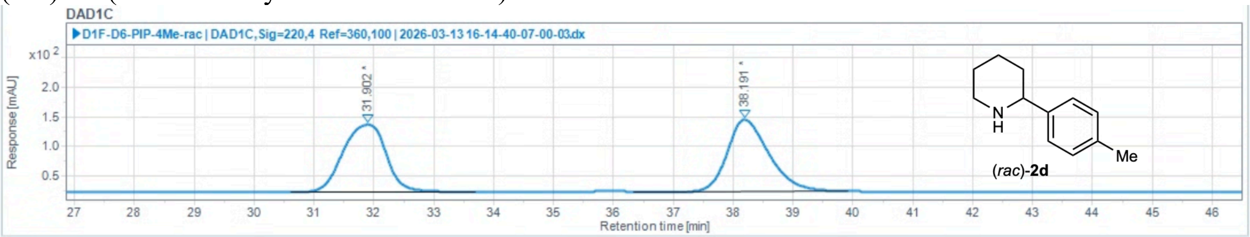

Injection Results

Peaks | Summary

| # | Name | Signal description | RT (min) | Area (mAU·s) | Area% | Height (mAU) | Height% | Amount | Concentration | Start tim |
|---|---|---|---|---|---|---|---|---|---|---|
| 1 | | DAD1C,Sig=220,4 Ref=360,100 | 31.902 | 5669.445 | 49.424 | 113.074 | 48.45 | | | |
| 2 | | DAD1C,Sig=220,4 Ref=360,100 | 38.191 | 5801.655 | 50.576 | 120.325 | 51.55 | | | |

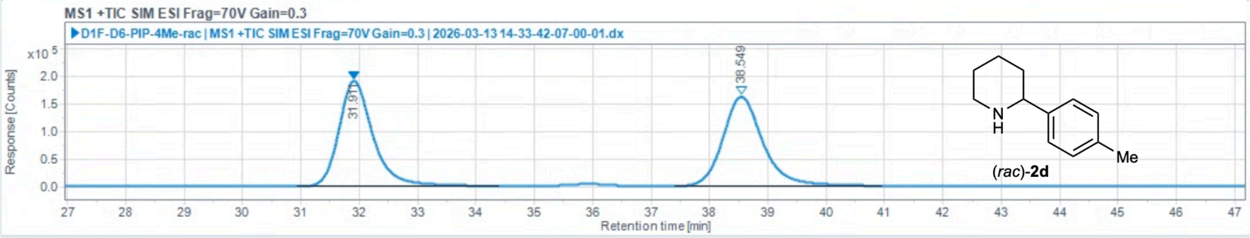

Injection Results

Peaks | Summary

| # | Name | Signal description | RT (min) | Area (Counts·s) | Area% | Height (Counts) | Height% | Amount | Concentration | Start tim |
|---|---|---|---|---|---|---|---|---|---|---|
| 1 | | MS1 +TIC SIM ESI Frag=70V Gain=0.3 | 31.911 | 7679986.394 | 50.051 | 190124.084 | 54.17 | | | |
| 2 | | MS1 +TIC SIM ESI Frag=70V Gain=0.3 | 38.549 | 7664386.118 | 49.949 | 160841.833 | 45.83 | | | |

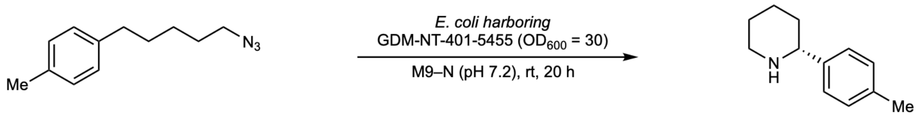

**1d** **2d**

Reaction performed on analytical scale using **GDM-NT-401-5455**: 95% ee favoring (*R*)-**2d**.

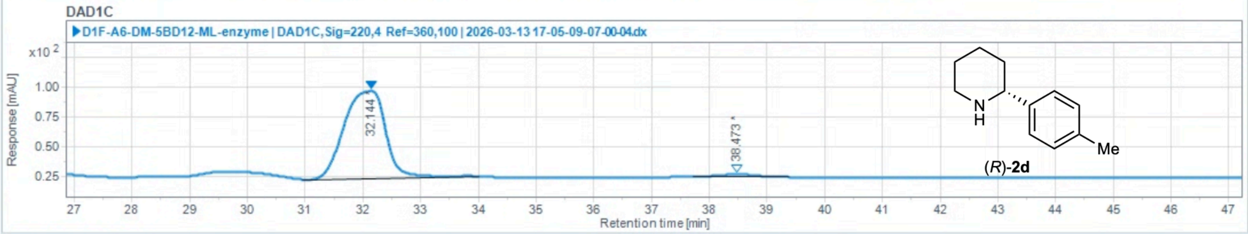

#### Injection Results

Peaks | Summary

| # | Name | Signal description | RT (min) | Area (mAU·s) | Area% | Height (mAU) | Height% | Amount | Concentration | Start tim |
|---|---|---|---|---|---|---|---|---|---|---|
| 1 | | DAD1C,Sig=220,4 Ref=360,100 | 32.144 | 3663.540 | 97.501 | 73.382 | 97.22 | | | |
| 2 | | DAD1C,Sig=220,4 Ref=360,100 | 38.473 | 93.883 | 2.499 | 2.096 | 2.78 | | | |

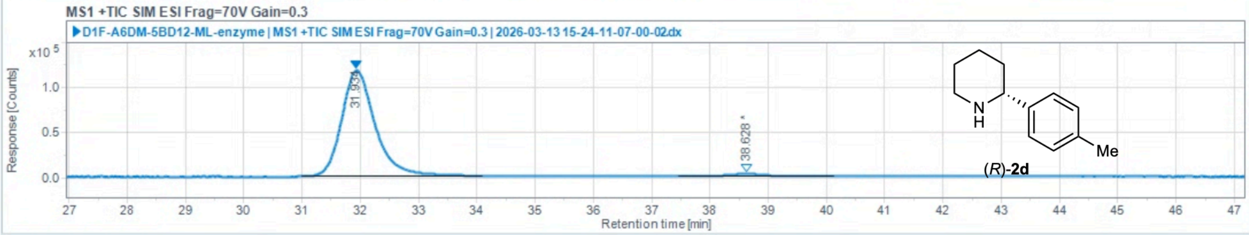

#### Injection Results

Peaks | Summary

| # | Name | Signal description | RT (min) | Area (Counts·s) | Area% | Height (Counts) | Height% | Amount | Concentration | Start tim |
|---|---|---|---|---|---|---|---|---|---|---|
| 1 | | MS1 +TIC SIM ESI Frag=70V Gain=0.3 | 31.934 | 4689084.199 | 97.331 | 117454.964 | 97.69 | | | |
| 2 | | MS1 +TIC SIM ESI Frag=70V Gain=0.3 | 38.628 | 128569.878 | 2.669 | 2780.380 | 2.31 | | | |

**2-(4-chlorophenyl)piperidine (2e)**

Chiral separation method: polar ionic HPLC-MS using *dual* Agilent InfinityLab Poroshell 120 Chiral-V, 2.7 µm, 2.1 × 150 mm; eluent: 40% MeOH (0.2 w% $NH_4CO_2H$), 60% $H_2O$ (20 mM $NH_4CO_2H$, pH 4.0 adjusted with $HCO_2H$); flow rate: 0.2 mL/min. $t_R$ = 39.843 min, $t_S$ = 43.259 min.

(*rac*)-**2e** (commercially available standard):

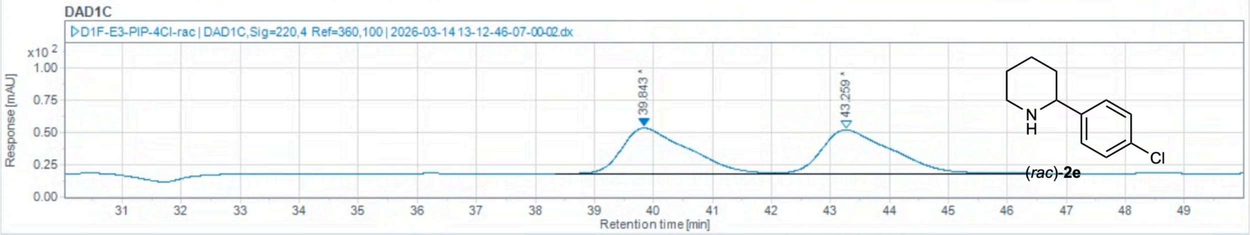

Injection Results

Peaks | Summary

| # | Name | Signal description | RT (min) | Area (mAU·s) | Area% | Height (mAU) | Height% | Amount | Concentration | Start tim |
|---|---|---|---|---|---|---|---|---|---|---|
| 1 | | DAD1C,Sig=220,4 Ref=360,100 | 39.843 | 2763.003 | 49.872 | 35.547 | 51.10 | | | |
| 2 | | DAD1C,Sig=220,4 Ref=360,100 | 43.259 | 2777.230 | 50.128 | 34.023 | 48.90 | | | |

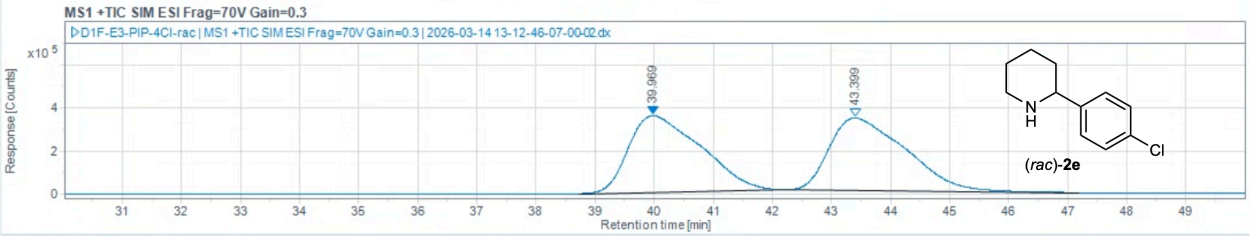

Injection Results

Peaks | Summary

| # | Name | Signal description | RT (min) | Area (Counts·s) | Area% | Height (Counts) | Height% | Amount | Concentration | Start tim |
|---|---|---|---|---|---|---|---|---|---|---|
| 1 | | MS1 +TIC SIM ESI Frag=70V Gain=0.3 | 39.969 | 30180876.610 | 49.930 | 355181.664 | 51.36 | | | |
| 2 | | MS1 +TIC SIM ESI Frag=70V Gain=0.3 | 43.399 | 30265091.243 | 50.070 | 336422.630 | 48.64 | | | |

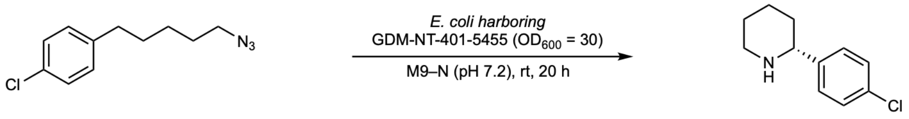

**1e** **2e**

Reaction performed on analytical scale using **GDM-NT-401-5455**: 98% ee favoring (*R*)-**2d**.

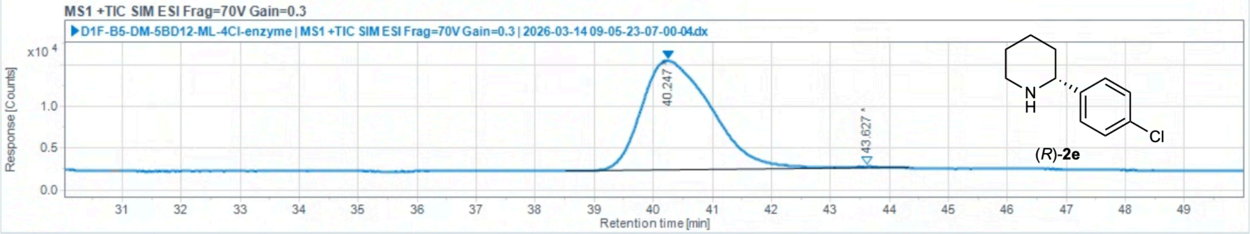

Injection Results

Peaks | Summary

| # | Name | Signal description | RT (min) | Area (Counts·s) | Area% | Height (Counts) | Height% | Amount | Concentration | Start tim |
|---|---|---|---|---|---|---|---|---|---|---|
| 1 | | MS1 +TIC SIM ESI Frag=70V Gain=0.3 | 40.247 | 1077781.535 | 99.164 | 13085.857 | 98.56 | | | |
| 2 | | MS1 +TIC SIM ESI Frag=70V Gain=0.3 | 43.627 | 9089.772 | 0.836 | 191.822 | 1.44 | | | |

**2-(4-fluorophenyl)piperidine (2f)**

Chiral separation method: polar ionic HPLC-MS using *dual* Agilent InfinityLab Poroshell 120 Chiral-V, 2.7 µm, 2.1 × 150 mm; eluent: 40% MeOH (0.2 w% $NH_4CO_2H$), 60% $H_2O$ (20 mM $NH_4CO_2H$, pH 4.0 adjusted with $HCO_2H$); flow rate: 0.2 mL/min. $t_R$ = 30.163 min, $t_S$ = 31.511 min.

(*rac*)-**2f** (commercially available standard):

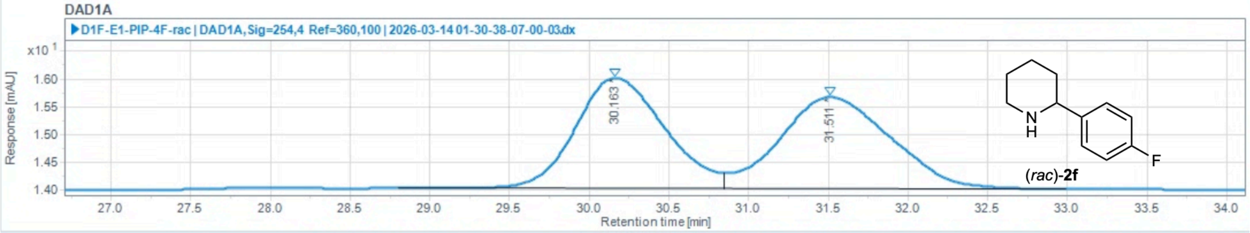

Injection Results

Peaks | Summary

| # | Name | Signal description | RT (min) | Area (mAU·s) | Area% | Height (mAU) | Height% | Amount | Concentration | Start tim |
|---|---|---|---|---|---|---|---|---|---|---|
| 1 | | DAD1A,Sig=254,4 Ref=360,100 | 30.163 | 77.217 | 49.283 | 1.982 | 54.66 | | | |
| 2 | | DAD1A,Sig=254,4 Ref=360,100 | 31.511 | 79.465 | 50.717 | 1.644 | 45.34 | | | |

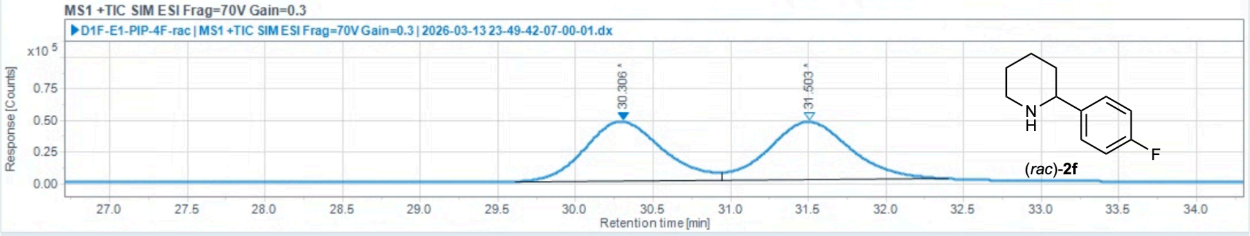

Injection Results

Peaks | Summary

| # | Name | Signal description | RT (min) | Area (Counts·s) | Area% | Height (Counts) | Height% | Amount | Concentration | Start tim |
|---|---|---|---|---|---|---|---|---|---|---|
| 1 | | MS1 +TIC SIM ESI Frag=70V Gain=0.3 | 30.306 | 1644114.358 | 49.372 | 47123.121 | 50.63 | | | |
| 2 | | MS1 +TIC SIM ESI Frag=70V Gain=0.3 | 31.503 | 1685944.497 | 50.628 | 45949.804 | 49.37 | | | |

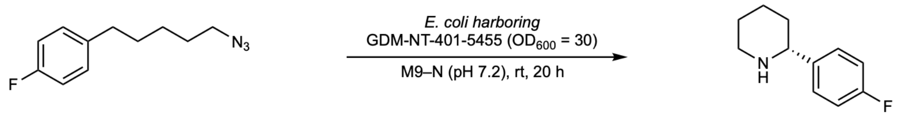

**1f** **2f**

Reaction performed on analytical scale using **GDM-NT-401-5455**: 89% ee favoring (*R*)-**2f**.

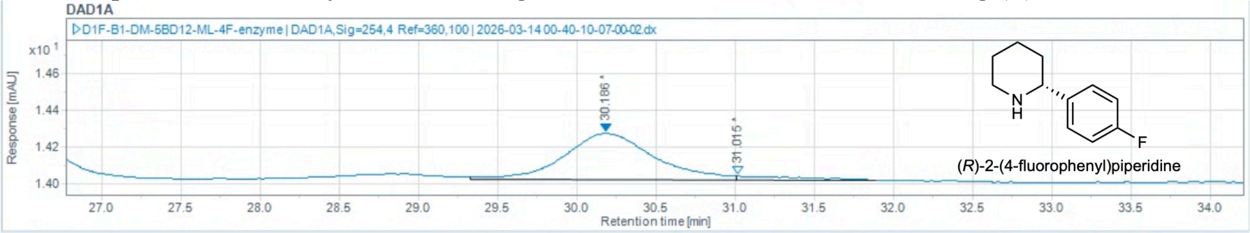

#### Injection Results

Peaks | Summary

| # | Name | Signal description | RT (min) | Area (mAU·s) | Area% | Height (mAU) | Height% | Amount | Concentration | Start tim |
|---|---|---|---|---|---|---|---|---|---|---|
| 1 | | DAD1A,Sig=254,4 Ref=360,100 | 30.186 | 10.046 | 94.637 | 0.252 | 92.17 | | | |
| 2 | | DAD1A,Sig=254,4 Ref=360,100 | 31.015 | 0.569 | 5.363 | 0.021 | 7.83 | | | |

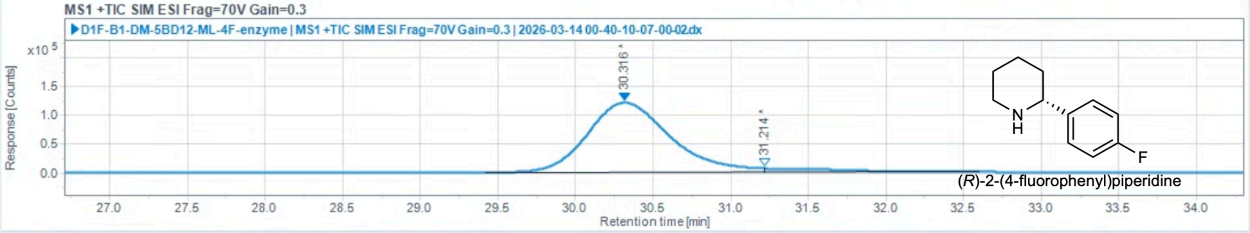

#### Injection Results

Peaks | Summary

| # | Name | Signal description | RT (min) | Area (Counts·s) | Area% | Height (Counts) | Height% | Amount | Concentration | Start tim |
|---|---|---|---|---|---|---|---|---|---|---|
| 1 | | MS1 +TIC SIM ESI Frag=70V Gain=0.3 | 30.316 | 4526151.842 | 95.489 | 121277.039 | 94.89 | | | |
| 2 | | MS1 +TIC SIM ESI Frag=70V Gain=0.3 | 31.214 | 213842.548 | 4.511 | 6533.229 | 5.11 | | | |

**2-(4-(trifluoromethyl)phenyl)piperidine (2g)**
Chiral separation method: chiral GC using chiral column Cyclosil-B (S/N USC696124H). $t_S$ = 23.409 min, $t_R$ = 23.486 min.

(*rac*)-**2g** (commercially available standard):

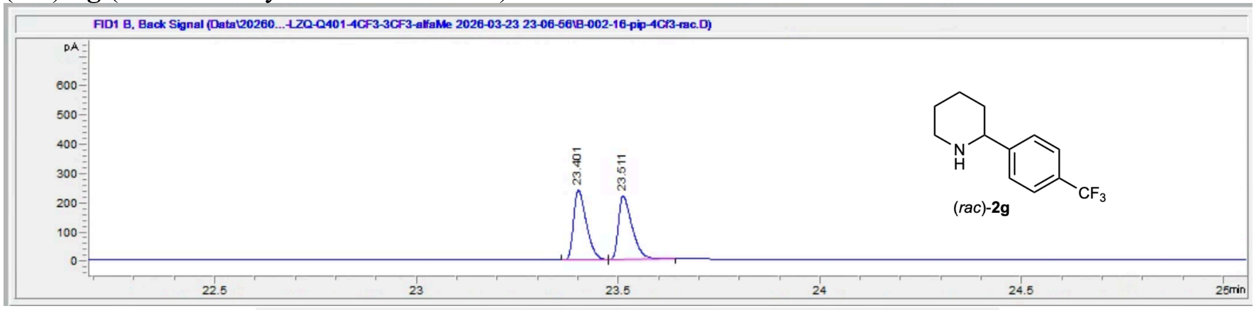

| # | Time | Type | Area | Height | Width | Area% | Symmetry |
|---|---|---|---|---|---|---|---|
| 1 | 23.401 | BB | 523.1 | 237.6 | 0.0329 | 50.779 | 0.6 |
| 2 | 23.511 | BB | 507.1 | 216.9 | 0.0361 | 49.221 | 0.504 |

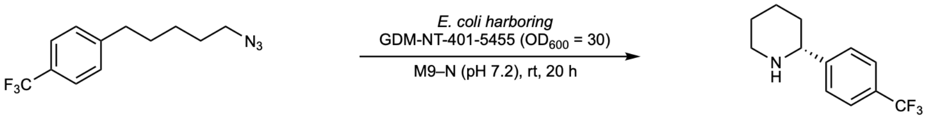

**1g** **2g**

Reaction performed on analytical scale using **GDM-NT-401-5455**: 87% ee favoring (*R*)-**2g**.

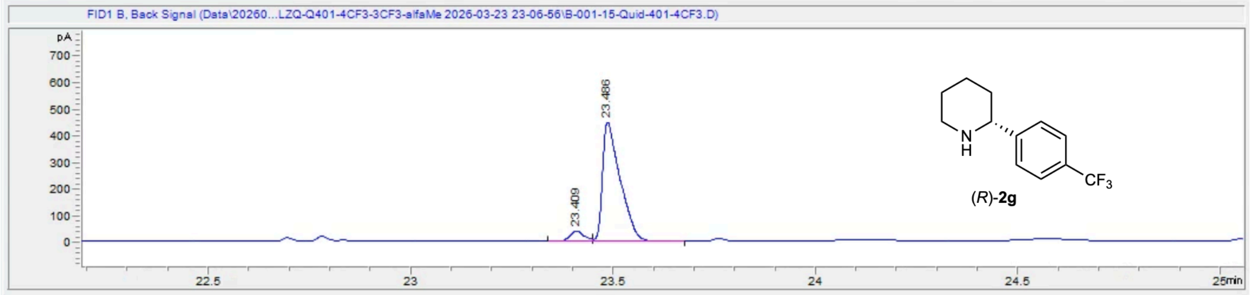

| # | Time | Type | Area | Height | Width | Area% | Symmetry |
|---|---|---|---|---|---|---|---|
| 1 | 23.409 | BV | 93.4 | 39 | 0.0352 | 6.474 | 0.854 |
| 2 | 23.486 | VB | 1349 | 445.6 | 0.0425 | 93.526 | 0.388 |

**2-(3-methoxyphenyl)piperidine (2h)**

Chiral separation method: normal phase HPLC using Daicel chiralpak IC column; eluent: hexanes:*i*-Pr-OH (90:10) with 0.1 v% diethylamine as additive; flowrate: 1.0 mL/min. $t_R$ = 9.144 min, $t_S$ = 10.291 min.

(*rac*)-**2h** (commercially available standard):

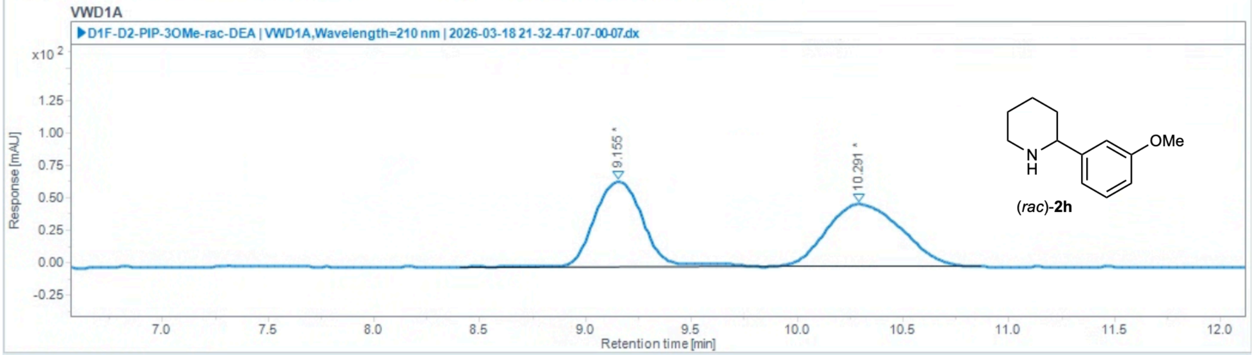

Injection Results

Peaks Summary

| Name | Signal description | RT (min) | Area (mAU·s) | Area% | Height (mAU) | Height% | Start time (min) | End time (min) |
|---|---|---|---|---|---|---|---|---|
| | VWD1A,Wavelength=210 nm | 9.155 | 1040.382 | 47.008 | 65.339 | 57.87 | 8.407 | 9.912 |
| | VWD1A,Wavelength=210 nm | 10.291 | 1172.823 | 52.992 | 47.576 | 42.13 | 9.912 | 10.869 |

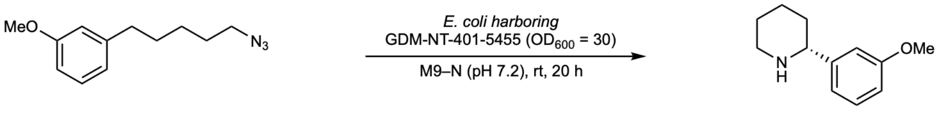

**1h** **2h**

Reaction performed on analytical scale using **GDM-NT-401-5455**: >99% ee favoring (*R*)-**2h**.

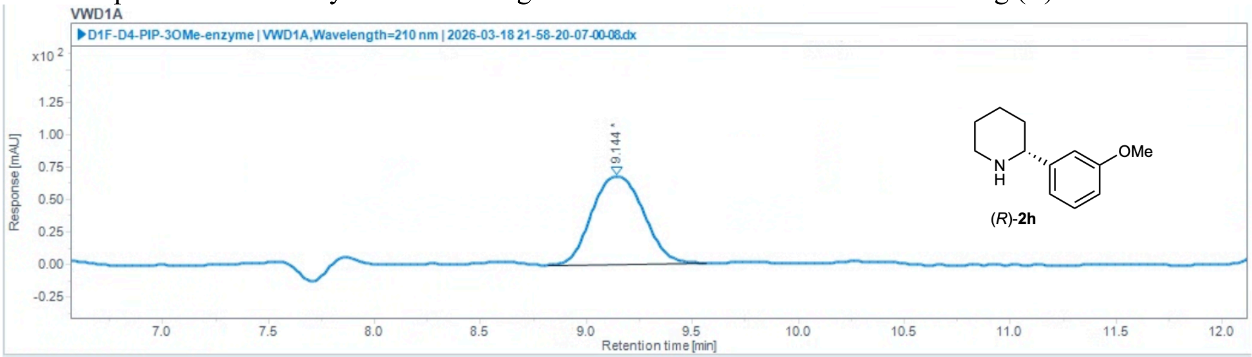

Injection Results

Peaks Summary

| Name | Signal description | RT (min) | Area (mAU·s) | Area% | Height (mAU) | Height% | Start time (min) | End time (min) |
|---|---|---|---|---|---|---|---|---|
| | VWD1A,Wavelength=210 nm | 9.144 | 1149.552 | 100.000 | 67.522 | 100.00 | 8.817 | 9.567 |

**2-(3-methylphenyl)piperidine (2i_Ts)**
Chiral separation method: normal phase HPLC using Daicel chiralpak IB column; eluent: hexanes:*i*-Pr-OH (95:5); flowrate: 0.6 mL/min. $t_S$ = 13.353 min, $t_R$ = 14.563 min.

(*rac*)-**2i_Ts** (commercially available standard):

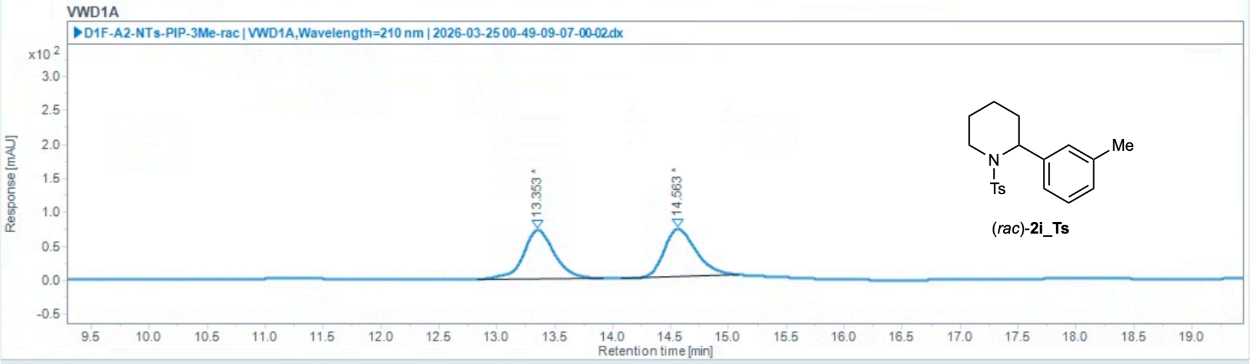

Injection Results

Peaks Summary

| Name | Signal description | RT (min) | Area (mAU·s) | Area% | Height (mAU) | Height% | Start time (min) | End time (min) |
|---|---|---|---|---|---|---|---|---|
| | VWD1A,Wavelength=210 nm | 13.353 | 1353.378 | 49.925 | 72.110 | 50.61 | 12.838 | 13.924 |
| | VWD1A,Wavelength=210 nm | 14.563 | 1357.431 | 50.075 | 70.361 | 49.39 | 14.079 | 15.093 |

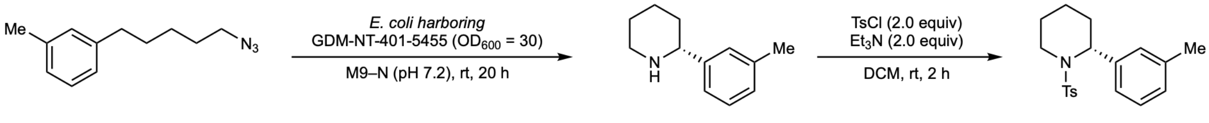

**1i** **2i** **2i_Ts**

Reaction performed on analytical scale using **GDM-NT-401-5455**: 94% ee favoring (*R*)-**2i**.

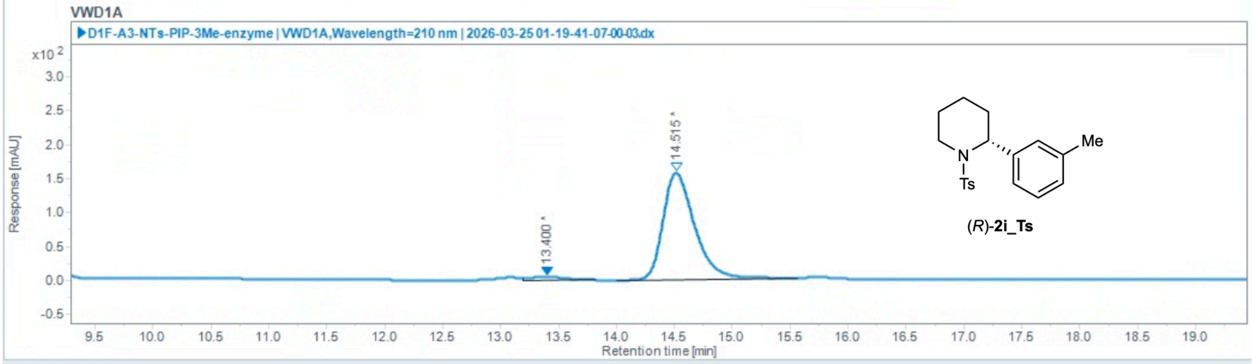

Injection Results

Peaks Summary

| Name | Signal description | RT (min) | Area (mAU·s) | Area% | Height (mAU) | Height% | Start time (min) | End time (min) |
|---|---|---|---|---|---|---|---|---|
| | VWD1A,Wavelength=210 nm | 13.400 | 94.867 | 3.104 | 4.703 | 2.92 | 13.200 | 13.821 |
| | VWD1A,Wavelength=210 nm | 14.515 | 2961.189 | 96.896 | 156.314 | 97.08 | 14.007 | 15.569 |

**2-(3-fluorophenyl)piperidine (2j)**

Chiral separation method: polar ionic HPLC-MS using *dual* Agilent InfinityLab Poroshell 120 Chiral-V, 2.7 µm, 2.1 × 150 mm; eluent: 40% MeOH (0.2 w% $NH_4CO_2H$), 60% $H_2O$ (20 mM $NH_4CO_2H$, pH 4.0 adjusted with $HCO_2H$); flow rate: 0.2 mL/min. $t_R$ = 29.974 min, $t_S$ = 31.838 min.

(*rac*)-**2j** (commercially available standard):

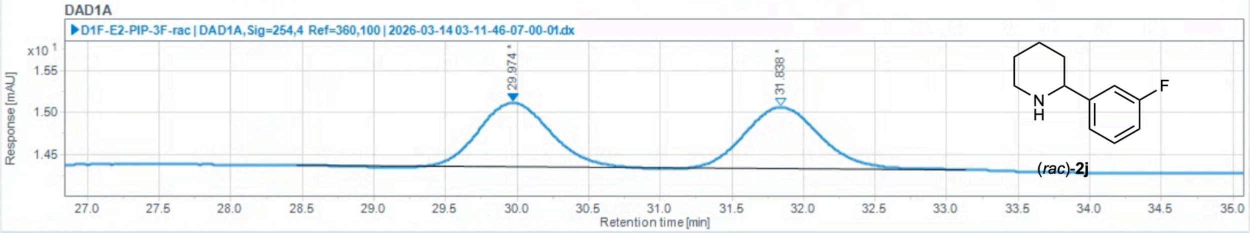

Injection Results

Peaks | Summary

| # | Name | Signal description | RT (min) | Area (mAU·s) | Area% | Height (mAU) | Height% | Amount | Concentration | Start tim |
|---|---|---|---|---|---|---|---|---|---|---|
| 1 | | DAD1A,Sig=254,4 Ref=360,100 | 29.974 | 26.382 | 49.431 | 0.766 | 51.17 | | | |
| 2 | | DAD1A,Sig=254,4 Ref=360,100 | 31.838 | 26.990 | 50.569 | 0.731 | 48.83 | | | |

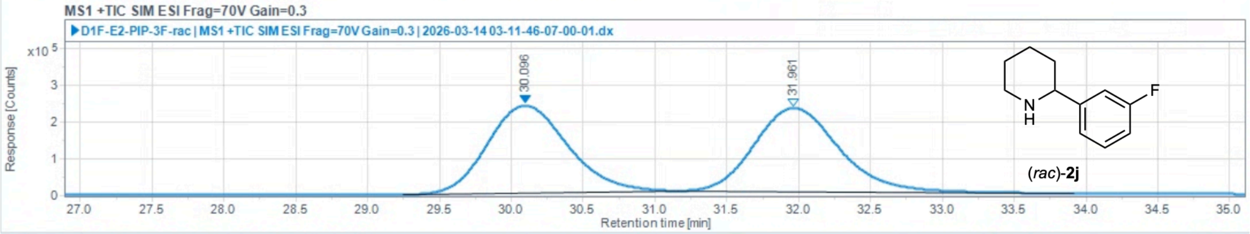

Injection Results

Peaks | Summary

| # | Name | Signal description | RT (min) | Area (Counts·s) | Area% | Height (Counts) | Height% | Amount | Concentration | Start tim |
|---|---|---|---|---|---|---|---|---|---|---|
| 1 | | MS1 +TIC SIM ESI Frag=70V Gain=0.3 | 30.096 | 9312011.716 | 49.689 | 238722.520 | 51.08 | | | |
| 2 | | MS1 +TIC SIM ESI Frag=70V Gain=0.3 | 31.961 | 9428684.231 | 50.311 | 228584.034 | 48.92 | | | |

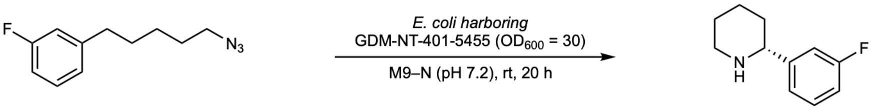

**1j** **2j**

Reaction performed on analytical scale using **GDM-NT-401-5455**: >99% ee favoring (*R*)-**2j**.

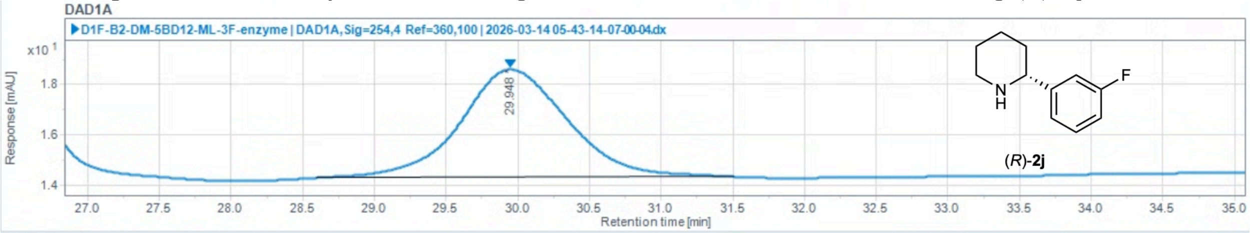

#### Injection Results

Peaks Summary

| # | Name | Signal description | RT (min) | Area (mAU·s) | Area% | Height (mAU) | Height% | Amount | Concentration | Start tim |
|---|---|---|---|---|---|---|---|---|---|---|
| 1 | | DAD1A,Sig=254,4 Ref=360,100 | 29.948 | 224.422 | 100.000 | 4.294 | 100.00 | | | |

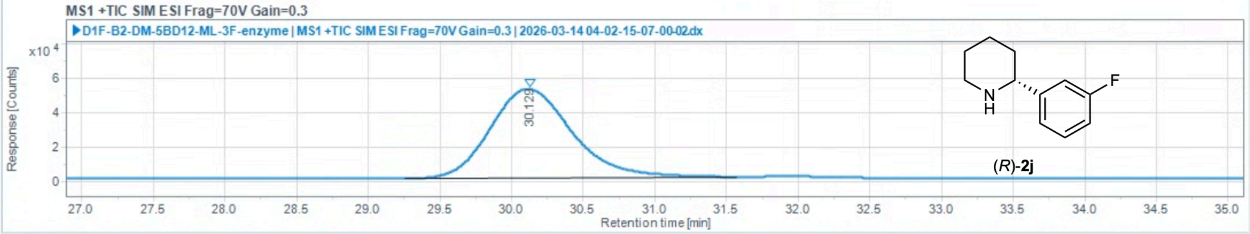

#### Injection Results

Peaks Summary

| # | Name | Signal description | RT (min) | Area (Counts·s) | Area% | Height (Counts) | Height% | Amount | Concentration | Start tim |
|---|---|---|---|---|---|---|---|---|---|---|
| 1 | | MS1 +TIC SIM ESI Frag=70V Gain=0.3 | 30.129 | 2061064.255 | 100.000 | 51659.638 | 100.00 | | | |

**2-(3-(trifluoromethyl)phenyl)piperidine (2k)**
Chiral separation method: chiral GC using chiral column Cyclosil-B (S/N USC696124H). $t_S$ = 22.681 min, $t_R$ = 22.751 min.

(*rac*)-**2k** (commercially available standard):

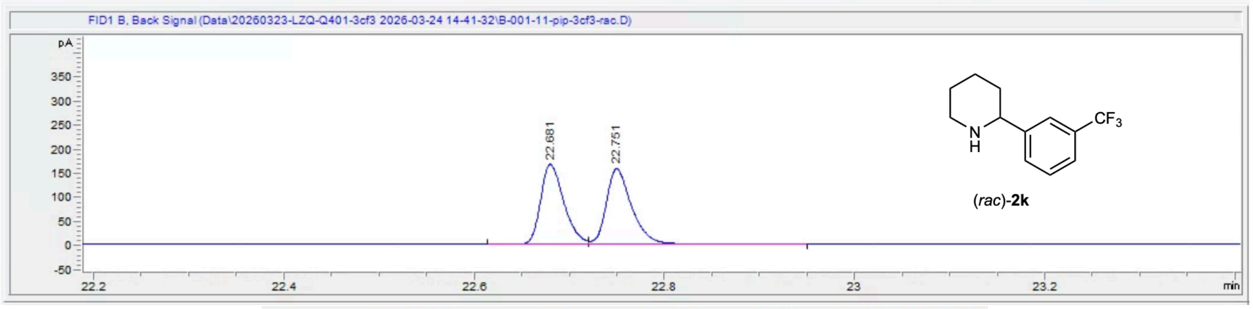

| # | Time | Type | Area | Height | Width | Area% | Symmetry |
|---|---|---|---|---|---|---|---|
| 1 | 22.681 | BV | 279.8 | 164.5 | 0.027 | 49.337 | 0.735 |
| 2 | 22.751 | VB | 287.4 | 156.2 | 0.0287 | 50.663 | 0.687 |

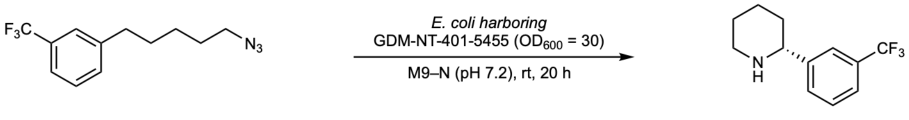

Reaction performed on analytical scale using **GDM-NT-401-5455**: 95% ee favoring (*R*)-**2k**.

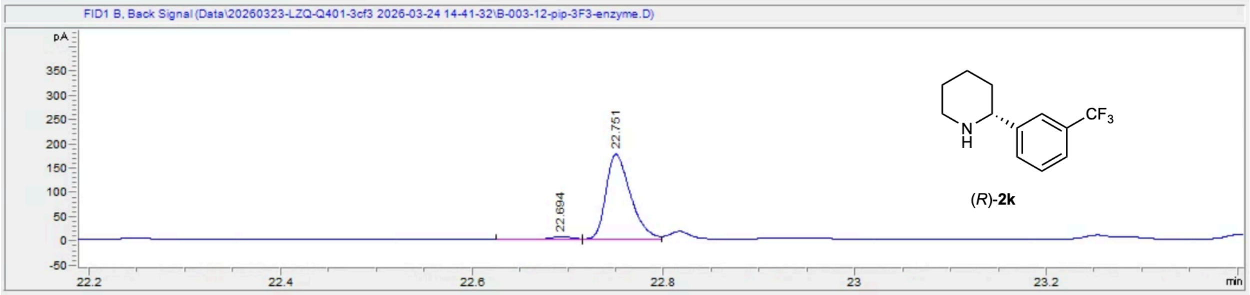

| # | Time | Type | Area | Height | Width | Area% | Symmetry |
|---|---|---|---|---|---|---|---|
| 1 | 22.694 | BV | 8.2 | 5.5 | 0.0238 | 2.552 | 0.999 |
| 2 | 22.751 | VV | 313.2 | 174.9 | 0.0281 | 97.448 | 0.694 |

**2-(naphthalen-2-yl)piperidine (2l)**

Chiral separation method: polar ionic HPLC-MS using *dual* Agilent InfinityLab Poroshell 120 Chiral-V, 2.7 µm, 2.1 × 150 mm; eluent: 40% MeOH (0.2 w% $NH_4CO_2H$), 60% $H_2O$ (20 mM $NH_4CO_2H$, pH 4.0 adjusted with $HCO_2H$); flow rate: 0.2 mL/min. $t_R$ = 66.624 min, $t_S$ = 78.949 min.

(*rac*)-**2l** (commercially available standard):

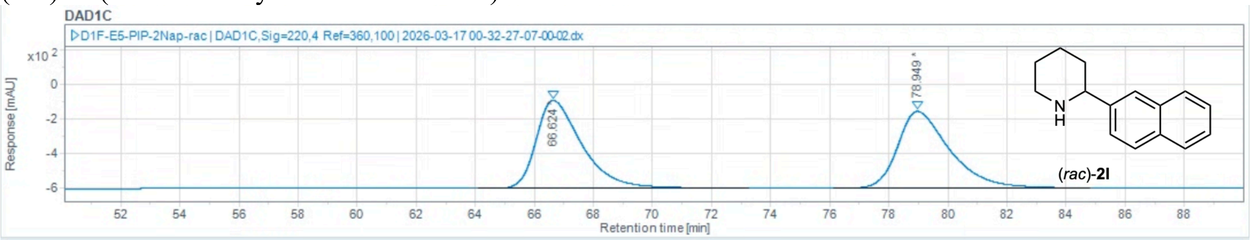

Injection Results

Peaks | Summary

| # | Name | Signal description | RT (min) | Area (mAU·s) | Area% | Height (mAU) | Height% | Amount | Concentration | Start tim |
|---|---|---|---|---|---|---|---|---|---|---|
| 1 | | DAD1C,Sig=220,4 Ref=360,100 | 66.624 | 54113.691 | 50.018 | 504.433 | 53.37 | | | |
| 2 | | DAD1C,Sig=220,4 Ref=360,100 | 78.949 | 54073.986 | 49.982 | 440.685 | 46.63 | | | |

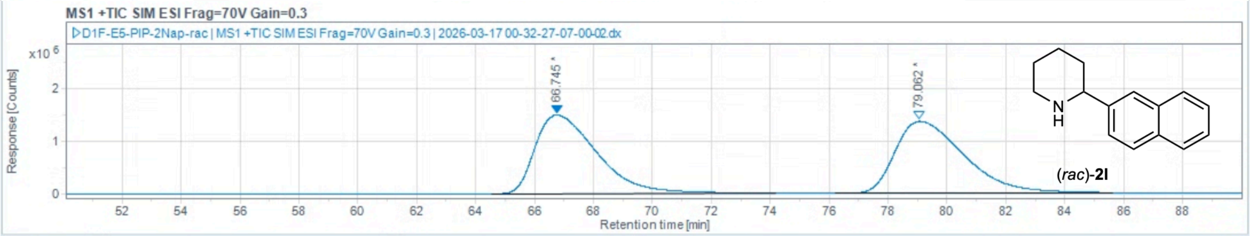

Injection Results

Peaks | Summary

| # | Name | Signal description | RT (min) | Area (Counts·s) | Area% | Height (Counts) | Height% | Amount | Concentration | Start tim |
|---|---|---|---|---|---|---|---|---|---|---|
| 1 | | MS1 +TIC SIM ESI Frag=70V Gain=0.3 | 66.745 | 218866336.508 | 49.596 | 1497041.905 | 52.19 | | | |
| 2 | | MS1 +TIC SIM ESI Frag=70V Gain=0.3 | 79.062 | 222436009.551 | 50.404 | 1371175.621 | 47.81 | | | |

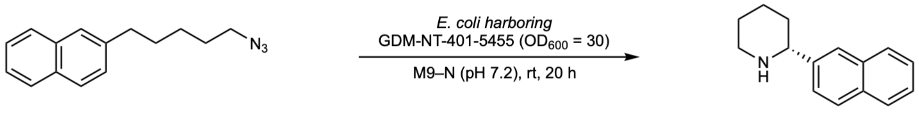

**1l** **2l**

Reaction performed on analytical scale using **GDM-NT-401-5455**: 89% ee favoring (*R*)-**2l**.

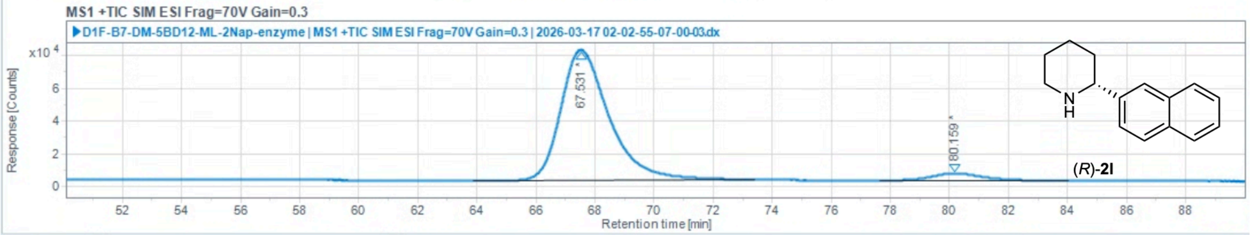

Injection Results

Peaks | Summary

| # | Name | Signal description | RT (min) | Area (Counts·s) | Area% | Height (Counts) | Height% | Amount | Concentration | Start tim |
|---|---|---|---|---|---|---|---|---|---|---|
| 1 | | MS1 +TIC SIM ESI Frag=70V Gain=0.3 | 67.531 | 8982932.124 | 94.481 | 79526.386 | 94.72 | | | |
| 2 | | MS1 +TIC SIM ESI Frag=70V Gain=0.3 | 80.159 | 524691.414 | 5.519 | 4429.400 | 5.28 | | | |

**2-phenylazepane (2m)**
Chiral separation method: polar ionic HPLC-MS using *dual* Agilent InfinityLab Poroshell 120 Chiral-V, 2.7 µm, 2.1 × 150 mm; eluent: 40% MeOH (0.2 w% $NH_4CO_2H$), 60% $H_2O$ (20 mM $NH_4CO_2H$, pH 4.0 adjusted with $HCO_2H$); flow rate: 0.2 mL/min. $t_1$ = 31.105 min, $t_2$ = 43.372 min.

(*rac*)-**2m** (commercially available standard):

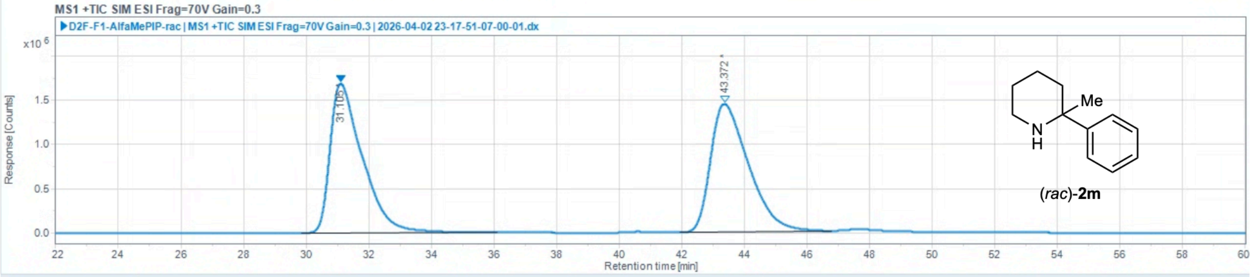

Injection Results

Peaks | Summary

| # | Name | Signal description | RT (min) | Area (Counts·s) | Area% | Height (Counts) | Height% | Amount | Concentration | Start time (min) | End time (min) |
|---|---|---|---|---|---|---|---|---|---|---|---|
| 1 | | MS1 +TIC SIM ESI Frag=70V Gain=0.3 | 31.105 | 120652233.554 | 49.777 | 1689689.656 | 53.87 | | | 29.846 | 36.086 |
| 2 | | MS1 +TIC SIM ESI Frag=70V Gain=0.3 | 43.372 | 121734017.317 | 50.223 | 1447104.868 | 46.13 | | | 41.952 | 46.763 |

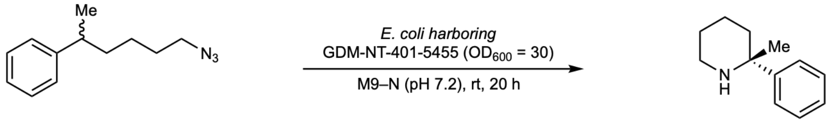

**1m** **2m**

Reaction performed on analytical scale using **GDM-NT-401-5455**: 36% ee favoring (*R*)-**2m**.

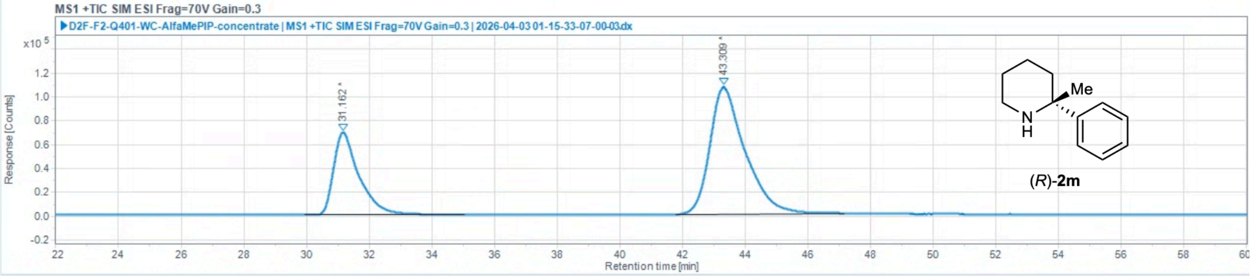

Injection Results

Peaks | Summary

| # | Name | Signal description | RT (min) | Area (Counts·s) | Area% | Height (Counts) | Height% | Amount | Concentration | Start time (min) | End time (min) |
|---|---|---|---|---|---|---|---|---|---|---|---|
| 1 | | MS1 +TIC SIM ESI Frag=70V Gain=0.3 | 31.162 | 3895077.488 | 31.888 | 69437.582 | 39.39 | | | 29.952 | 35.026 |
| 2 | | MS1 +TIC SIM ESI Frag=70V Gain=0.3 | 43.309 | 8319617.524 | 68.112 | 106846.025 | 60.61 | | | 41.797 | 47.144 |

**2-phenylazepane (2n)**

Chiral separation method: polar ionic HPLC-MS using *dual* Agilent InfinityLab Poroshell 120 Chiral-V, 2.7 µm, 2.1 × 150 mm; eluent: 40% MeOH (0.2 w% $NH_4CO_2H$), 60% $H_2O$ (20 mM $NH_4CO_2H$, pH 4.0 adjusted with $HCO_2H$); flow rate: 0.2 mL/min. $t_S$ = 34.593 min, $t_S$ = 36.125 min.

(*rac*)-**2n** (commercially available standard):

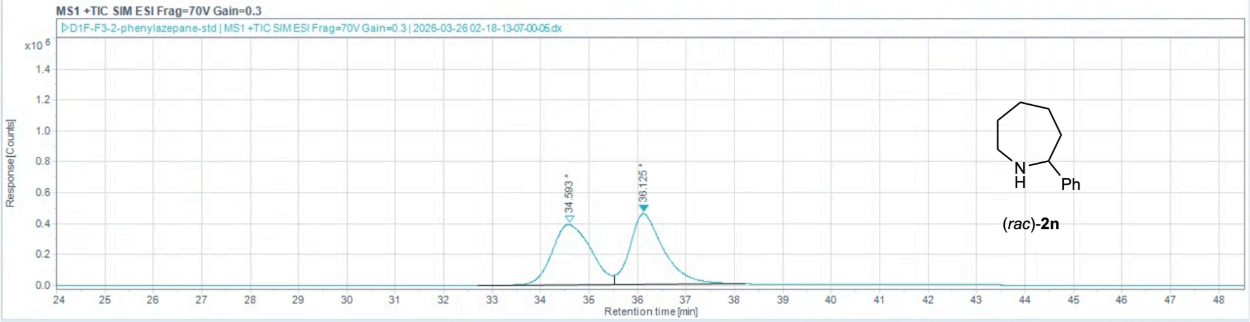

Injection Results

Peaks | Summary

| # | Name | Signal description | RT (min) | Area (Counts·s) | Area% | Height (Counts) | Height% | Amount | Concentration | Start time (min) | End time (min) |
|---|---|---|---|---|---|---|---|---|---|---|---|
| 1 | | MS1 +TIC SIM ESI Frag=70V Gain=0.3 | 34.593 | 21532441.948 | 49.263 | 391394.568 | 45.92 | | | 32.711 | 35.519 |
| 2 | | MS1 +TIC SIM ESI Frag=70V Gain=0.3 | 36.125 | 22176390.220 | 50.737 | 460855.590 | 54.08 | | | 35.519 | 38.228 |

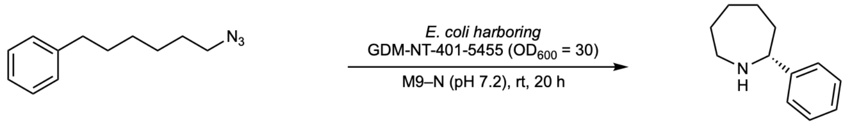

**1n** **2n**

Reaction performed on analytical scale using **GDM-NT-401-5455**: 98% ee

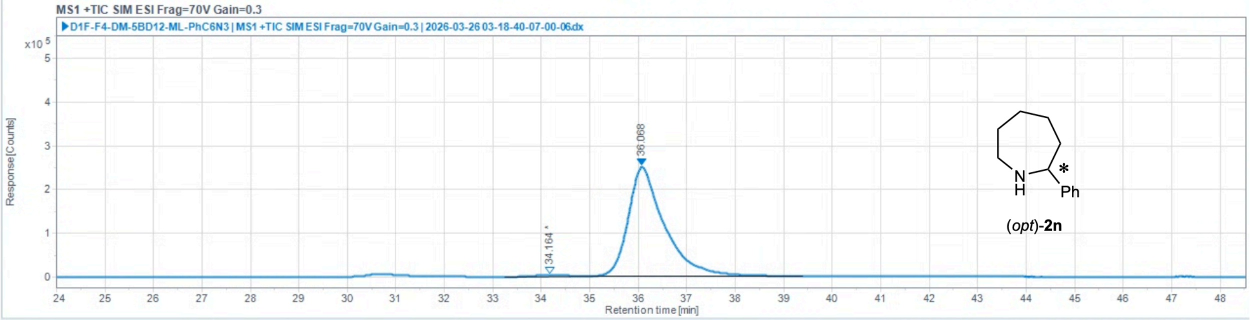

Injection Results

Peaks | Summary

| # | Name | Signal description | RT (min) | Area (Counts·s) | Area% | Height (Counts) | Height% | Amount | Concentration | Start time (min) | End time (min) |
|---|---|---|---|---|---|---|---|---|---|---|---|
| 1 | | MS1 +TIC SIM ESI Frag=70V Gain=0.3 | 34.164 | 136310.449 | 1.046 | 3405.375 | 1.35 | | | 33.236 | 35.065 |
| 2 | | MS1 +TIC SIM ESI Frag=70V Gain=0.3 | 36.068 | 12893372.628 | 98.954 | 249166.032 | 98.65 | | | 35.065 | 39.387 |

**2-phenylpyrrolidine (2o)**

Chiral separation method: polar ionic HPLC-MS using *dual* Agilent InfinityLab Poroshell 120 Chiral-V, 2.7 µm, 2.1 × 150 mm; eluent: 40% MeOH (0.2 w% $NH_4CO_2H$), 60% $H_2O$ (20 mM $NH_4CO_2H$, pH 4.0 adjusted with $HCO_2H$); flow rate: 0.2 mL/min. $t_1$ = 32.092 min, $t_2$ = 33.260 min.

(*rac*)-**2o** (commercially available standard):

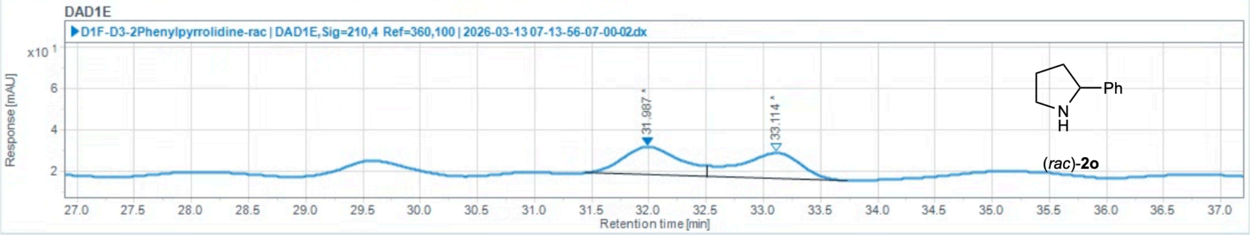

Injection Results

Peaks | Summary

| # | Name | Signal description | RT (min) | Area (mAU·s) | Area% | Height (mAU) | Height% | Amount | Concentration | Start tim |
|---|---|---|---|---|---|---|---|---|---|---|
| 1 | | DAD1E,Sig=210,4 Ref=360,100 | 31.987 | 479.985 | 50.853 | 13.589 | 52.20 | | | |
| 2 | | DAD1E,Sig=210,4 Ref=360,100 | 33.114 | 463.884 | 49.147 | 12.442 | 47.80 | | | |

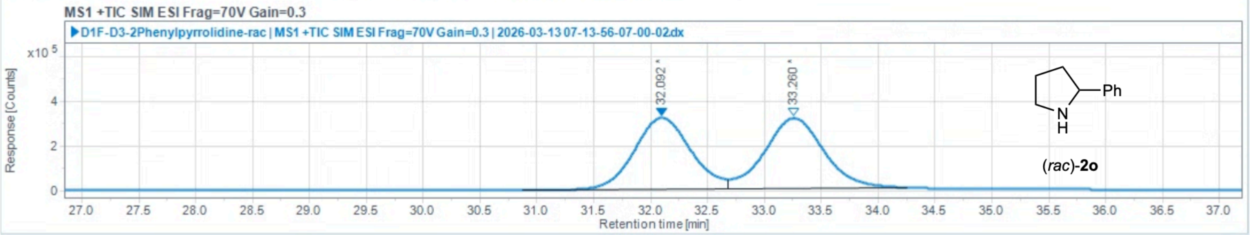

Injection Results

Peaks | Summary

| # | Name | Signal description | RT (min) | Area (Counts·s) | Area% | Height (Counts) | Height% | Amount | Concentration | Start tim |
|---|---|---|---|---|---|---|---|---|---|---|
| 1 | | MS1 +TIC SIM ESI Frag=70V Gain=0.3 | 32.092 | 10958013.430 | 49.032 | 319228.485 | 50.38 | | | |
| 2 | | MS1 +TIC SIM ESI Frag=70V Gain=0.3 | 33.260 | 11390788.639 | 50.968 | 314351.664 | 49.62 | | | |

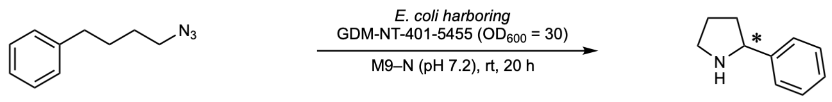

**1o** **2o**

Reaction performed on analytical scale using **GDM-NT-401-5455**: 15% ee

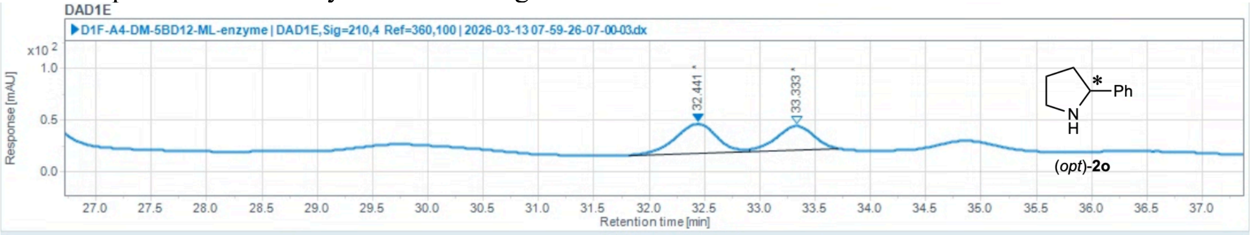

#### Injection Results

Peaks | Summary

| # | Name | Signal description | RT (min) | Area (mAU·s) | Area% | Height (mAU) | Height% | Amount | Concentration | Start tim |
|---|---|---|---|---|---|---|---|---|---|---|
| 1 | | DAD1E,Sig=210,4 Ref=360,100 | 32.441 | 726.046 | 57.676 | 28.015 | 55.00 | | | |
| 2 | | DAD1E,Sig=210,4 Ref=360,100 | 33.333 | 532.782 | 42.324 | 22.920 | 45.00 | | | |

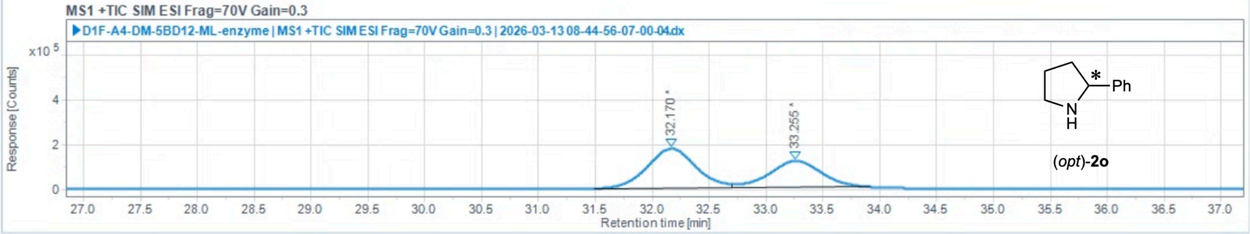

#### Injection Results

Peaks | Summary

| # | Name | Signal description | RT (min) | Area (Counts·s) | Area% | Height (Counts) | Height% | Amount | Concentration | Start tim |
|---|---|---|---|---|---|---|---|---|---|---|
| 1 | | MS1 +TIC SIM ESI Frag=70V Gain=0.3 | 32.170 | 5292778.016 | 58.222 | 176851.000 | 59.96 | | | |
| 2 | | MS1 +TIC SIM ESI Frag=70V Gain=0.3 | 33.255 | 3797956.252 | 41.778 | 118108.134 | 40.04 | | | |
